# Uridine Bisphosphonates Differentiate Phosphoglycosyl Transferase Superfamilies

**DOI:** 10.1101/2023.09.19.558431

**Authors:** Leah M. Seebald, Pouya Haratipour, Michaela R. Jacobs, Hannah M. Bernstein, Boris A. Kashemirov, Charles E. McKenna, Barbara Imperiali

## Abstract

Complex bacterial glycoconjugates are essential for bacterial survival, and drive interactions between pathogens and symbionts, and their human hosts. Glycoconjugate biosynthesis is initiated at the membrane interface by phosphoglycosyl transferases (PGTs), which catalyze the transfer of a phosphosugar from a soluble uridine diphospho-sugar (UDP-sugar) substrate to a membrane-bound polyprenol-phosphate (Pren-P). Two distinct superfamilies of PGT enzymes, denoted as polytopic and monotopic, carry out this reaction but show striking differences in structure and mechanism. With the goal of creating non-hydrolyzable mimics (UBP-sugars) of the UDP-sugar substrates as chemical probes to interrogate critical aspects of these essential enzymes, we designed and synthesized a series of uridine bisphosphonates (UBPs), wherein the diphosphate bridging oxygen of the UDP and UDP-sugar is replaced by a substituted methylene group (CXY; X/Y = F/F, Cl/Cl, (*S*)-H/F, (*R*)-H/F, H/H, CH_3_/CH_3_). These compounds, which incorporated as the conjugating sugar an *N*-acetylglucosamine (GlcNAc) substituent at the β-phosphonate, were evaluated as inhibitors of a representative polytopic PGT (WecA from *Thermotoga maritima*) and a monotopic PGT (PglC from *Campylobacter jejuni*). Although CHF-BP most closely mimics pyrophosphate with respect to its acid/base properties, the less basic CF_2_-BP conjugate most strongly inhibited PglC, whereas the more basic CH_2_-BP analogue was the strongest inhibitor of WecA. These surprising differences indicate different modes of ligand binding for the different PGT superfamilies implicating a modified P–O^−^ interaction with the structural Mg^2+^, consistent with their catalytic divergence. Furthermore, at least for the monoPGT superfamily example, this was not the sole determinant of ligand binding: the two diastereomeric CHF-BP conjugates, which feature a chiral center at the P_α_-CHF-P_β_ carbon, exhibited strikingly different binding affinities and the inclusion of GlcNAc with the native α-anomer configuration significantly improved binding affinity. UBP-sugars are a valuable tool for elucidating the structures and mechanisms of the distinct PGT superfamilies and offer a promising scaffold to develop novel antibiotic agents for the exclusively prokaryotic monoPGT superfamily.

**TABLE OF CONTENTS GRAPHIC:** 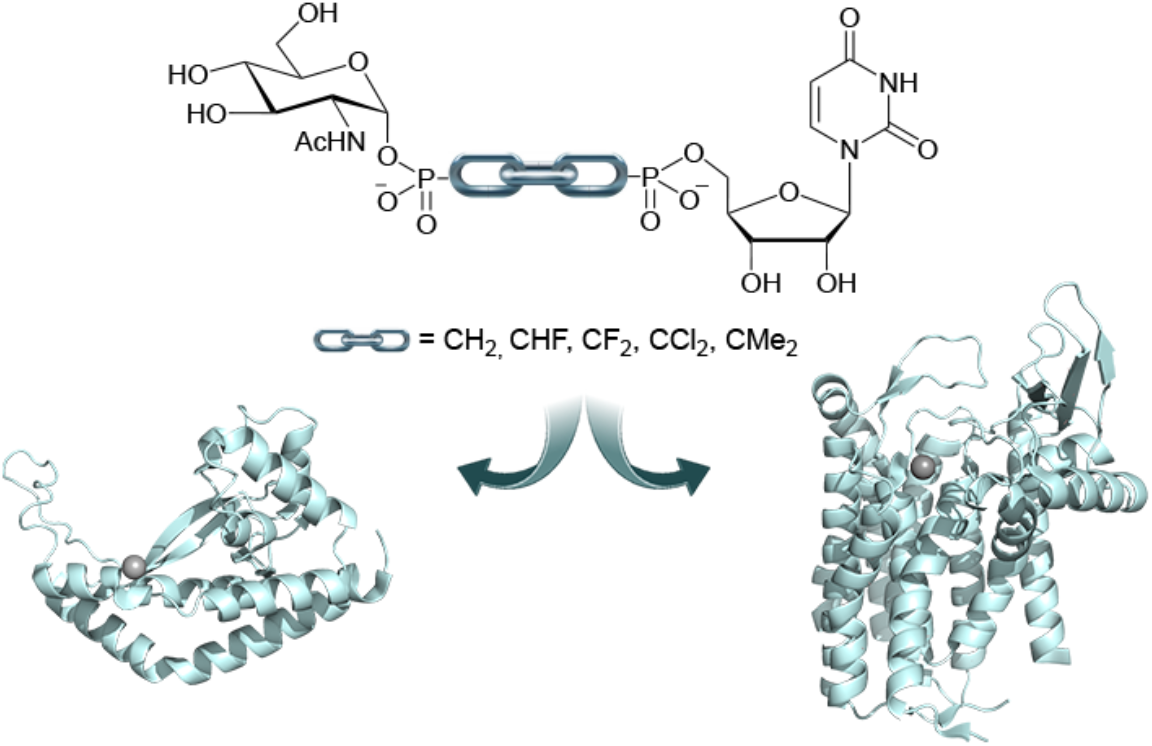

---

Nucleoside diphosphate sugar (NDP-sugar) substrates are involved in a variety of essential cellular processes and serve key roles as glycosyl and phosphoglycosyl donor substrates for the biosynthesis of complex glycoconjugates.^1^ An important strategy for glycoconjugate assembly involves a membrane-associated initiation step catalyzed by a phosphoglycosyl transferase (PGT). PGTs mediate the transfer of a phosphosugar from an NDP-sugar substrate to a membrane-anchored polyprenol phosphate (Pren-P) acceptor (**Figure 1**).^2^ The resulting PGT product, a Pren-PP-sugar, is then further elaborated by stepwise addition of sugars from NDP-sugar donors, catalyzed by the sequential action of a set of glycosyl transferases (GTs).^3^

**Figure 1.**
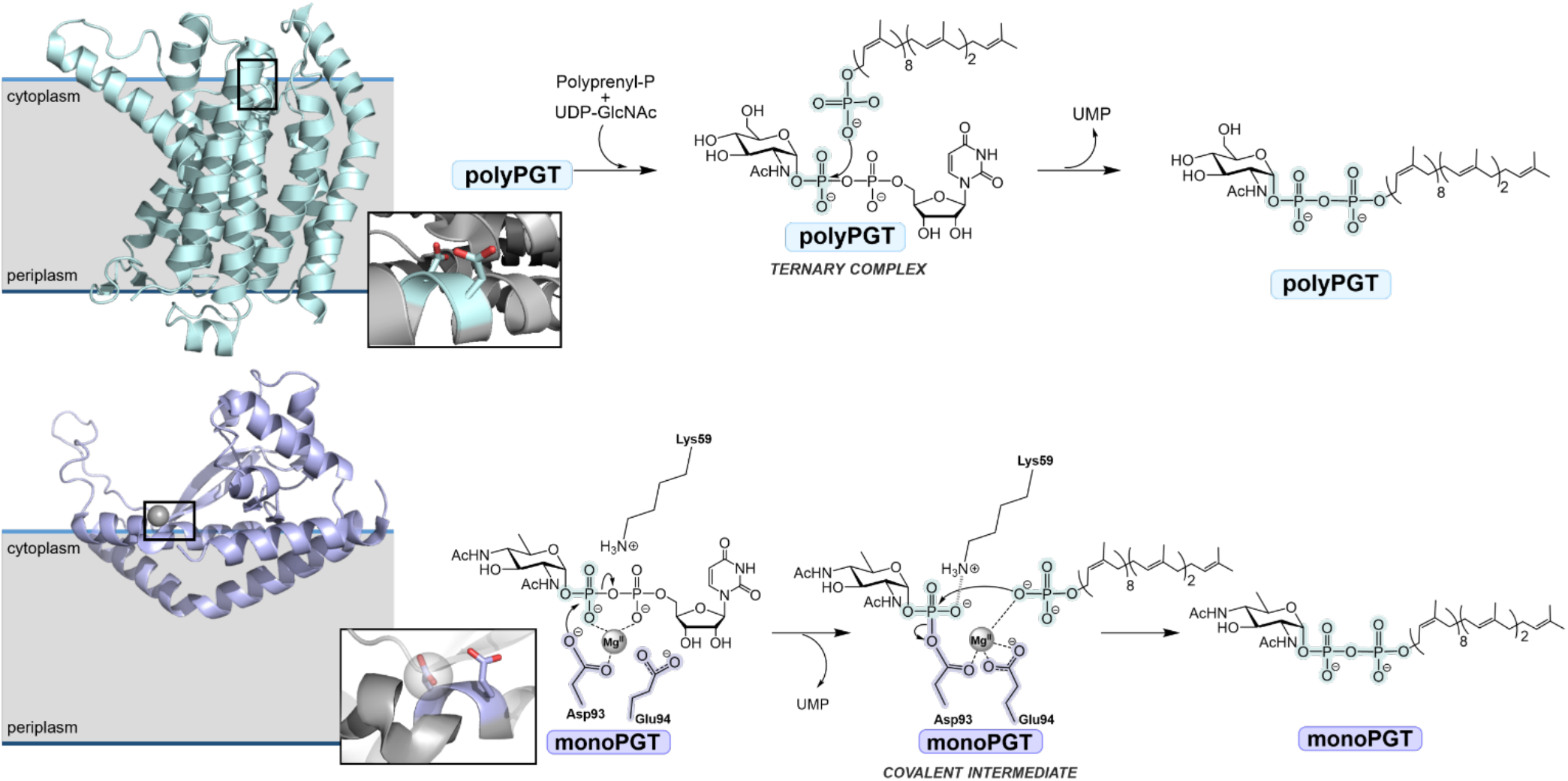
**A**. Ribbon diagram of polyPGT MraY from *Aquifex aeolicus* (PDB 5CKR). Insert shows close-up view of the MraY active site. PolyPGTs invoke a mechanism involving a ternary complex intermediate and feature a conserved Gly-Xaa-Xaa-Asp-Asp motif. **B**. Ribbon diagram of monoPGT PglC from *Campylobacter concisus* (PDB 5W7L), insert shows the close-up view of the PglC active site Asp-Glu catalytic dyad. MonoPGTs follow a ping-pong mechanism, wherein a covalent phosphoglycosyl-Asp intermediate is formed.

There is emerging interest in PGTs due in part to the surprising mechanistic and structural dichotomy between the two known PGT superfamilies.^4^ These superfamilies include either a polytopic or a monotopic functional domain,^5^ and, although the different PGTs catalyze chemically equivalent transformations, thus far, it has been shown that polytopic PGTs (polyPGTs) proceed through a ternary complex mechanism,^6-8^ and monotopic PGTs (monoPGTs) invoke a ping-pong mechanism (**Figure 1AB**).^9^ Although polyPGTs are observed across all domains of life, extensive bioinformatics analyses have revealed that the monoPGT superfamily is exclusive to prokaryotes.^10,11^ The biological importance of PGTs in the first step of glycoconjugate assembly in bacterial pathogens and symbionts highlights the significance of structural and mechanistic studies to provide new insight into the determinants of ligand specificity and the key drivers of catalysis as the foundation for inhibitor and chemical probe discovery.

Progress in our understanding of the structures and mechanisms of PGTs is challenged by the fact that these enzymes are integral membrane proteins. To date, structures of two polyPGTs have been reported. These are MraY (*Aquifex aeolicus*)^12^ the essential PGT in the biosynthesis of the bacterial cell wall peptidoglycan and the human GlcNAc-1-P-transferase or GPT (DPAGT1), which catalyzes the initiating step in the dolichol pathway for *N*-linked protein glycosylation.^8, 13, 14^ The structures are intriguing as several show complexes of the polyPGTs with bisubstrate analogues of natural product origin (e.g. tunicamycin and muraymycin D2).^12, 14, 15^ The polyPGT superfamily features a conserved Gly-Xaa-Xaa-Asp-Asp (GXXDD) motif with the conserved adjacent aspartic acid residues proposed to be required for substrate orientation and essential for activity in these and other polyPGTs.^16,17^ Although many of the inhibitor- and substrate-bound complexes lack the Mg^2+^ co-factor, two structures of the human GPT with tunicamycin^14^ and the UDP-GlcNAc/Mg^2+^ complex^13^ provide insight into polyPGT-small molecule binding.

For the monoPGT superfamily, there is a single X-ray crystal structure of PglC (from *Campylobacter concisus*).^18^ Structural and mechanistic studies support a mechanism involving a catalytic Asp-Glu (DE) dyad, wherein the aspartic acid carboxylate attacks at the β-phosphate of the NDP-sugar substrate forming a covalent phosphosugar intermediate.^9^ In the structure, the essential Mg^2+^ cofactor is coordinated to the Asp carboxyl group and a phosphate ion.^18^ In this case, the two-step ping-pong mechanism of the monoPGTs precludes determination of the structure of the Michaelis complex in the presence of the NDP-sugar substrate bound and Mg^2+^, as the covalent intermediate forms even in the absence of the Pren-P acceptor substrate. At first glance, the GXXDD motif of the polyPGT superfamily appears to bear resemblance to the Asp-Glu (DE) catalytic dyad in the monoPGT family. However, based on the crystal structures of the PGTs, the geometry of these motifs are quite distinct; the conserved adjacent aspartic acid residues of the polyPGTs are presented on a canonical α-helix,^19^ whereas the DE catalytic-dyad of the monoPGT superfamily has side chains with co-facial positioning along the 3^10^ helix that scaffolds the active site (**Figure 1**, inserts).^18^

Despite the biological significance of PGTs, the mechanistic tools available for probing these crucial enzymes in glycoconjugate assembly are limited. Furthermore, many of the existing small molecule probes are derived from complex nucleoside natural products, which are mostly proposed to bind to the polyPGTs as bisubstrate analogues.^15, 20^ As a complement to the natural product inhibitors and the synthetic analogues that they inspire,^21^ non-cleavable NDP-sugar CXY-bisphosphonate analogues (NBPs) can potentially provide insight into the structure and function of enzymes, such as the PGTs, that act on NDP-sugar substrates. Importantly, NBP-sugars could provide critical information for structure determination,^22, 23^ as inhibitor scaffolds for identifying structure-activity relationships (SAR),^21^ including inhibitor binding modes^24^ and, in the case of monoPGTs, as leads for agents with antibiotic activity.

With these opportunities in mind, we describe here the modular syntheses of a series of uridine 5’-bisphosphonate (CXY-UBP) and uridine 5’-bisphosphonate-*N*-acetylglucosamine (GlcNAc-CXY-UBP) analogues (**Figure 2**) in which the central diphosphate (P-O-P) oxygen is replaced by a substituted methylene (P-CXY-P), wherein X/Y = H/H, F/F, Cl/Cl, CH_3_/CH_3_, (*S*)-H/F and (*R*)-H/F. The α,β-CXY-and β,γ-bisphosphonate analogues of deoxynucleotides^25-29^ have been previously used as probes of nucleic acid polymerase structure and function.^30-33^ In addition to introduction of a non-hydrolyzable bisphosphonate mimicking the natural diphosphate, the CXY substitution allows variation in such properties as POH/PO^−^ acidity, P-O/P-C bond lengths, P-O-P/P-C-P bond angles,^34^ and the effect of steric perturbation adjacent to the CXY group.^35^ Finally, for BPs where X ≠ Y, as in CHF, the individual diastereomeric analogues will have nearly equivalent POH/PO^−^ acidity but different orientations of the C-X and C-Y substituents within the binding site. Herein, the PGT substrate analogues are applied in comparative inhibition studies of representative members of the two PGT superfamilies.

**Figure 2.**
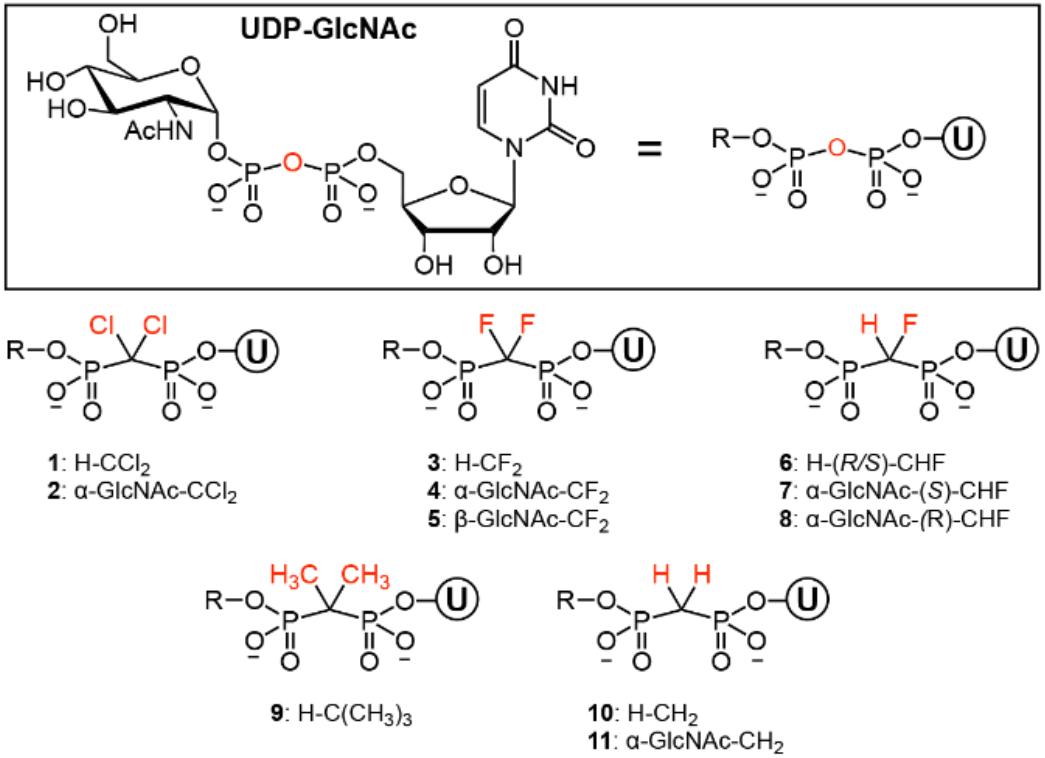
Non-hydrolyzable uridine 5’-bisphosphonate analogues (H-CXY-UBP) and uridine 5’-bisphosphonate-*N*-acetylglucosamine analogues (GlcNAc-CXY-UBP) synthesized and analyzed in this study. The diphosphate bridging oxygen of the UDP is replaced by a panel of substituted methylene groups (CXY; X/Y = F/F, Cl/Cl, (*R*)-H/F, (*S*)-F/H, H/H, CH_3_/CH_3_). Uridine is denoted as a circled “U”. The stereochemical assignments of **7** and **8** are (*S*)-CHF and (*R*)-CHF, respectively.

## Results

### Synthesis of the CXY-UBP and GlcNAc-CXY-UBP analogues

We developed two synthetic routes to the nucleotide analogues in this study (**Figure 2, 1**-**11**). The routes differ in the initial conjugation partner for the bisphosphonate moiety (uridine or GlcNAc). **Scheme 1A**, exemplified by the synthesis of α-GlcNAc-CH_2_ (**11**) (11% overall yield), benefits from the use of unprotected uridine but requires two demethylation steps due to the use of trimethyl ester bisphosphonate. In contrast, **Scheme 1B** is more modular, facilitating diversification of the bisphosphonate components, and affords higher overall yields. **Scheme 1B** was utilized to synthesize five R-CXY-UBPs (R=H, CYX: CCl_2_ (**1**), CF_2_ (**3**), (*R*/*S*)-CHF (**6**), C(CH_3_)_3_ (**9**), CH_2_ (**10**) (71-90% overall yields) and five GlcNAc-CXY-UBP derivatives α-GlcNAc-CCl_2_ (**2**), α-GlcNAc-CF_2_ (**4**), β-GlcNAc-CF_2_ (**5**), α-GlcNAc-(*S*)-CHF (**7**), α-GlcNAc-(*R*)-CHF (**8**) (∼25% overall yields). All compounds were characterized by ^31^P NMR, ^1^H NMR, ^13^C NMR, ^19^F NMR (for CHF and CF_2_ analogues), COSY, HSQC, and high-resolution mass spectrometry (HRMS). To assign the CHF stereocenter in α-GlcNAc-(*S*)-CHF (**7**) and α-GlcNAc-(*R*)-CHF (**8**), a modified synthesis incorporating a sterochemically-defined bisphosphonate intermediate derivatized with a chiral auxiliary was deployed (**Scheme 2**).

**Scheme 1.**
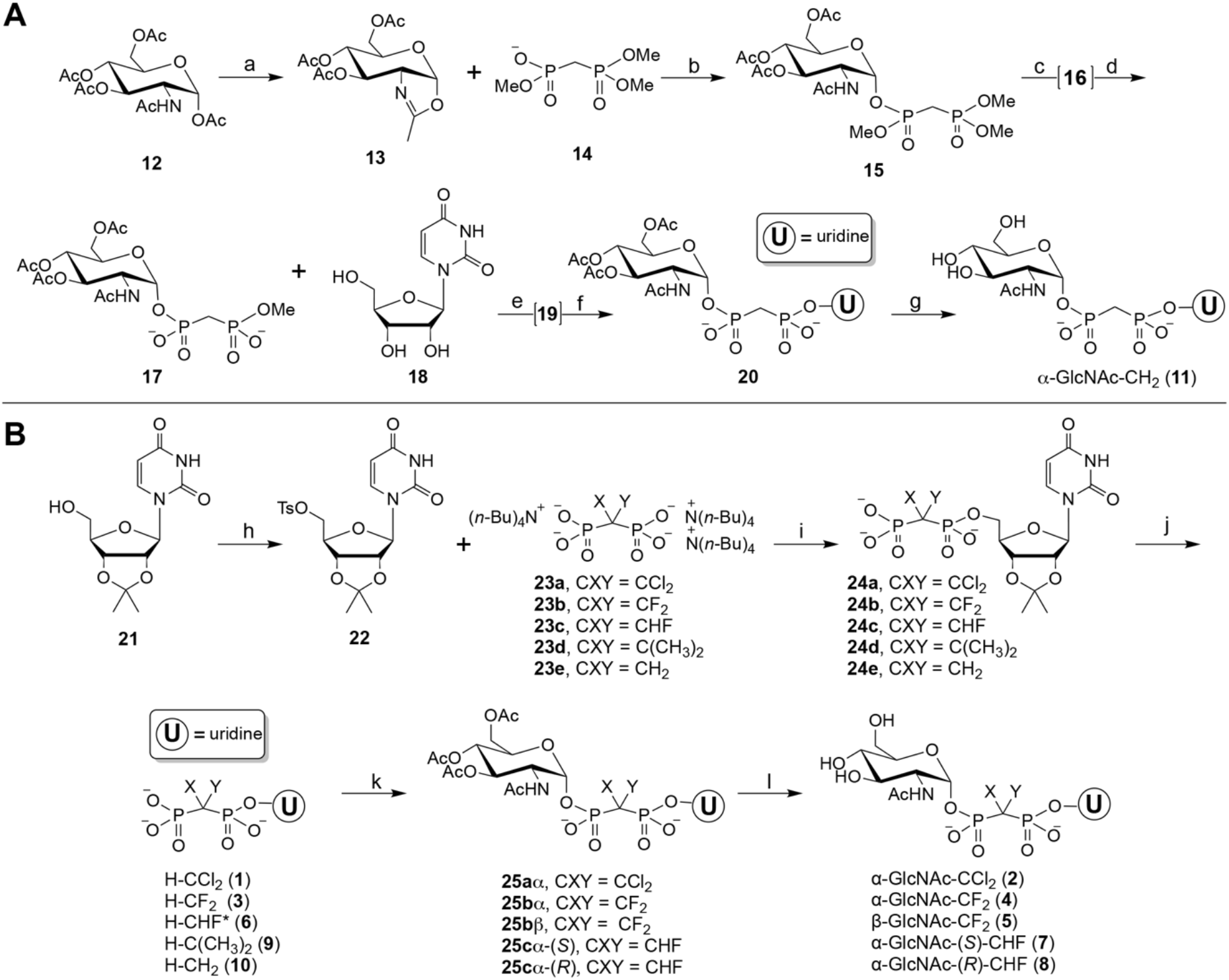
**A**. Reagents and conditions: (a) Triflic acid (HOTf), CH_2_Cl_2_, rt, inert atmosphere, 2 h (quant. yield of **13**); (b) **14** (from tetramethyl ester refluxed 5.5 h in MeCN with excess TEA, monoacid generated with DOWEX H^+^(90%)) and **13**, dioxane, 55 °C, inert atmosphere, 18 h (95%); (c) NaI, acetone, rt, 18 h (quant. yield of symmetrical disodium salt (**16**); (d) DOWEX H^+^, MeOH, H_2_O, 0 °C (quant. yield of **17**) (e) **18** (uridine), dioxane, DMF, PPh_3_, DIAD, rt, inert atmosphere, 18 h (yield of **19** not measured); (f) PhSH, DIPEA, NMP, 40 °C, inert atmosphere, 48 h (20% yield of **20** from **17**); (g) H_2_O, NH_4_OH (pH 11), rt, 1 h (70%). **B**. Reagents and conditions: (h) **21** (2’,3’-*O*-isopropylidene uridine), *p*-TsCl, pyridine, rt, inert atmosphere, 18 h (**22**, 95%); (i) general procedure: **23** [from tetraacid, H_2_O, EtOH, 3 equiv. (n-Bu)_4_NOH, pH ∼7.5, rt, 5 min], MeCN, rt, inert atmosphere (**23a-23e**, 75-95%); (j) aq. HCl pH ∼0.5, rt, 2 h (quant. yield of **24a-e**); (k) 1) DOWEX H^+^, H_2_O; 2) **2** (GlcNAc oxazoline), DMF, 45 °C, inert atmosphere, 48 h (49-70%); to obtain β-anomer: 75 °C, 3 h (75%)), the mixture of (*R*/*S*)-CHF diastereomers was cleanly separated by preparative reverse-phase HPLC; (l) aq. NH_4_OH (pH 10.5-11.5), rt, 2.5 h (quant. yield). α-GlcNAc-(*S*)-CHF (**7**) and α-GlcNAc-(*R*)-CHF (**8**) were established as (*S*)-CHF and (*R*)-CHF diastereomers, respectively.

**Scheme 2.**
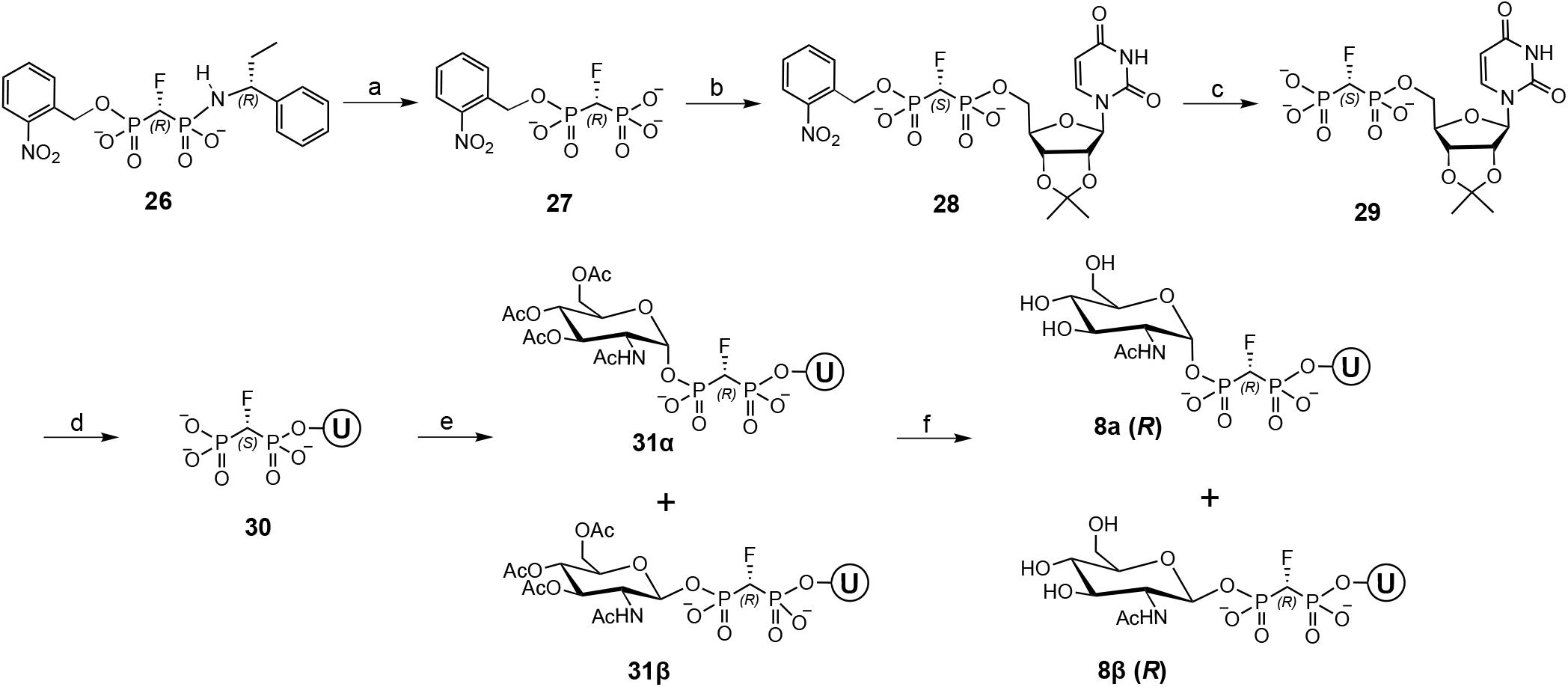
Reagents and conditions: (a) i. 1 M HCl, rt, ii. DOWEX H^+^, H_2_O, MeOH (quant. yield of **27**) (b) **22**, DMF, 50 °C, inert atmosphere (44% yield of **28**), (c) H_2_O, UV (λ = 365 nm), 48 h (76% yield of **29**) (d) 1 M HCl, rt, 1 h (quant. yield of **30**) (e) [bisphosphonate preparation: 1:1 H_2_O, MeOH, 3 equiv. (n-Bu)_4_NOH, pH ∼8.3, rt], DMF, GlcNAc oxazoline = **13**, 45 °C, inert atmosphere, 5.5 h (57% yield of **31α** and **31β**) (f) H_2_O, NH_4_OH pH 10.5, 28 °C, overnight (31% yield of **8α** and **8β**). Note: The designation of chirality at the -CHF-center changes through the scheme due to the change in Cahn-Ingold-Prelog priority of the phosphate ester substituents.

**Scheme 1A** begins with peracetyl-GlcNAc, **12**, which is converted to the corresponding oxazoline (**13**) following known procedures.^36^ Compound **13**, in the presence of trimethyl methylenebis(phosphonate) (**14**) resulted in the bisphosphonate ester **15**. The bisphosphonate ester (**15**) was symmetrically di-demethylated with sodium iodide to obtain the intermediate **16**, which was converted to **17** by passage through DOWEX H^+^. The Mitsunobu coupling of **17** and uridine (**18**), which favored attack at the less sterically hindered β-P atom provided intermediate **19**, which was then demethylated with DIPEA/PhSH to yield **20**.^37, 38^ The *O*-acetyl groups in **20** were removed with aqueous ammonium hydroxide solution to afford the desired product α-GlcNAc-CH_2_ (**11**).

The second route (**Scheme 1B**) begins with activation of 2’,3’-*O*-isopropylidene uridine (**21**) as the 5’-tosyl ester **22**,^39^ followed by reaction with the tris(tetrabutylammonium) salt of the selected methylene-bisphosphonates (**23a-e**) and then ion exchange to afford **24a-e**. The 2’,3’-isopropylidene protecting group was then removed under acidic conditions to obtain five H-CXY-UBPs: H-CCl_2_ (**1**), H-CF_2_ (**3**), H-(*R*/*S*)-CHF (**6**), H-C(CH_3_)_3_ (**9**), H-CH_2_ (**10**). Selected H-CXY-UBP products (**1, 3, 6**) were reacted with **13** to afford *O*-acetyl protected GlcNAc-CXY-UBPs (**25a-c)**. In summary, the reaction of **1** with **13** at 45 °C, provided **25aα**. Under similar conditions, UBP **3** provided **25bα**. In addition, when **3** was subjected to modified conditions, including higher temperatures, the glycosidic linkage was epimerized to the β-anomer at C-1 designated as **25bβ**.^36^ Finally, the H-(*R*/*S*)-CHF mixture, **6\***, was converted to **25cα\*** by reaction with **13** using the standard 45 °C conditions and individual (*R*/*S*)-CHF diastereomers were separated by RP-HPLC. *O*-acetyl deprotection of the GlcNAc under basic conditions produced α-GlcNAc-CCl_2_ (**2**), α-GlcNAc-CF_2_ (**4**), β-GlcNAc-CF_2_ (**5**), and the individual stereoisomer α-GlcNAc-(*S*)-CHF (**7**) and α-GlcNAc-(*R*)-CHF (**8**). For **7** and **8**, the *O*-acetylated GlcNAc substituent was crucial to this separation as the precursor H-(*R*/*S*)-CHF (**6**) could not be resolved by RP-HPLC, and thus was studied as an epimeric mixture **6\***. Isolation of each *O*-acetylated GlcNAc containing CHF-epimer **(25cα)** was carried out by RP-HPLC at the penultimate step in **Scheme 1B**, by separating **25cα\***. This purification yielded individual, but not yet stereochemically-defined, epimers of *S*-**25cα** and *R*-**25cα**, primed for the final synthetic step.^25^ Compound **9** was not converted to the corresponding α-GlcNAc-C(CH_3_)_2_ derivative due to the weak inhibition of the PGTs observed for this UBP precursor.

The CHF-bisphosphonate intermediate (**23c**) is prochiral, thus, the product mono- and di-phosphonate esters with uridine and GlcNAc, such as **7** and **8**, were obtained as a 1:1 mixture of diastereomers. These related isomers could be cleanly separated by preparative HPLC. However, as presented later, the diastereomeric α-GlcNAc-(*S/R*)-CHF compounds showed different inhibitory activity with the mono- and polyPGTs thus motivating elucidation of the absolute stereochemistry at the CHF chiral center. To accomplish this, we prepared CHF bisphosphonate derivatives modified with an enantiopure chiral auxiliary, (*R*)-(+)-α-ethylbenzylamine, and a photolabile protecting group (**26**) that allowed the correlation of chromatographic elution and NMR properties with absolute stereochemistry as defined by X-ray crystallographic analysis (**Scheme 2**, shown for a selected isomer).^25^ The chiral auxiliary was removed from **26** under acidic conditions producing **27**, which was then coupled to the tosyl-activated uridine isopropylidine (**22**) to give **28**. Photochemical deprotection at 365 nm yielded **29**, followed by uridine deprotection (to **30**) and coupling to the GlcNAc oxazoline (**13**) afforded the acetyl protected α- and β-nucleotide analogues **31α** and **31β**. After deacetylation of **31α**, chromatographic analysis was conducted as in the initial synthesis of **7** and **8**, and the HPLC retention times correlated α-GlcNAc-(*S*)-CHF (**7**) as the (*S*)-isomer and α-GlcNAc-(*R*)-CHF (**8**) as the (*R*)-isomer. Detailed NMR analysis was also carried out to support the absolute assignment of configuration (**S186 and S187**)

### Enzyme targets and biochemical analysis

The bisphosphonate analogues were assessed for inhibition of representative members of each of the two PGT superfamilies. For the monoPGT analysis we screened two PglC orthologs; one from *Campylobacter jejuni* (PglC (*Cj*), which is a human food-borne enteropathogen that is a significant cause of gastroenteritis worldwide^40^ and the other from *C. concisus* (72% sequence homology with *Cj*).^41^ The PglC (*Cc*) was included as it is the target of the only successful monoPGT X-ray structure determination,^18^ however as the orthologs showed similar trends and as the majority of the mechanistic studies had been carried out with PglC (*Cj*),^9^ we pursued detailed analysis of the *Cj* ortholog. The PglCs from *Campylobacter* catalyzes the first membrane-committed step in the biosynthesis of *N*-linked glycoproteins.^42^ The monoPGT data are compared with that obtained for the polyPGT WecA from *Thermotoga maritima* (WecA (*Tm*)).^6^ WecA is an integral membrane protein that initiates the O-antigen and enterobacterial common antigen biosynthesis pathways by catalyzing the transfer of GlcNAc-1-phosphate to undecaprenyl phosphate (Und-P) to produce Und-PP-GlcNAc.^17, 43^ WecA natively uses UDP-GlcNAc as substrate.

Biochemical analysis has shown that the biochemically-preferred substrate for the *Campylobacter* PglCs is UDP-*N,N*ʹ-diacetylbacillosamine (UDP-diNAcBac) (**Fig. S1**).^3^ However, for these studies, *N*-acetylglucosamine (GlcNAc) was included in the structures of the UBP analogues rather than diNAcBac to simplify and expedite synthesis. GlcNAc shares structural features with diNAcBac, including similar stereochemistry and the C2-*N*-acetamido moiety. UDP-GlcNAc is accepted as a substrate by the PglC enzymes included in this analysis although in general the K_m_ values are significantly higher (∼20-fold) than those for UDP-diNAcBac.

Using the Promega UMP/CMP-Glo™ system, PGT steady-state kinetics were studied under initial linear rate conditions with <10% UDP-diNAcBac turnover. The UMP/CMP-Glo™ assay quantifies PGT activity through detection of released UMP by enzyme-mediated conversion of the UMP by-product to a luciferase substrate that can be monitored by luminescence. In all cases, a correction to account for any off-target inhibition of the UMP Glo™ assay reagent enzymes was included in all analyses.^44^ Due to off-target interactions with assay components when analyzing the simple (non-glycosylated) UBP analogues, inhibition studies were also carried out applying an orthogonal radioactivity-based assay which monitors transfer of a radiolabeled-phosphosugar from UDP-[^3^H]-diNAcBac to the unlabeled Und-P acceptor; substrate conversion is quantified by scintillation counting after liquid/liquid extraction.^3^

### Inhibition of PGTs by selected H-CXY-UBP and GlcNAc-CXY-UBP analogues

The simple UBP analogues including **1** (H-CCl_2_), **3** (H-CF_2_), **6** (H-(*R*/*S*)-CHF), **9** (H-C(CH_3_)_3_), and **10** (H-CH_2_), (**Figure 2**, R=H), were first screened at 100 µM with the PGT enzymes to identify the bisphosphonate bridging atoms that conferred the best binding (**Figure S2**). Compound (**3**) H-CF_2_ demonstrated modest inhibition towards each PGT, with 33% ± 6% inhibition of PglC (*Cj*), 14% ± 11% inhibition of PglC (*Cc*), and 19% ± 10% inhibition of WecA (*Tm*). Inhibition by the other UBP analogues was negligible and in some cases the limitations of the UMP-Glo assay precluded conclusive interpretation at the R=H UBP stage.

To develop the probes, we then investigated modification of **3** with GlcNAc featuring either α- or β-anomer linkages (α-GlcNAc-CF_2_ (**4**) and β-GlcNAc-CF_2_ (**5**)). The inclusion of both anomers was valuable for determining whether the GlcNAc-UBPs bound in a substrate-like mode, as the PGTs in this analysis are known to act on the α-anomers of bacterial UDP-sugars. Inhibition studies at 100 µM with PglC (*Cj*) and WecA (*Tm*), comparing UBP H-CF_2_ (**3**) with α-GlcNAc-CF_2_ (**4**), and β-GlcNAc-CF_2_ (**5**) are illustrated in **Figure 3A**. With PglC (*Cj*), comparison of H-CF_2_ (**3**) with α-GlcNAc-CF_2_ (**4**) showed that incorporation of α-GlcNAc into the bisphosphonate analogue enhances binding and increases inhibition from 33% ± 6% to 76% ± 13%. In contrast, β-GlcNAc-CF_2_ (**5**) shows < 5% inhibition under identical conditions, confirming that the inclusion of a sugar with the native stereochemistry promotes more efficient binding. Inhibition of PglC (*Cj*) by **3** and **4** is concentration dependent (**Fig. 3B**), and the IC_50_ of **4** for PglC (*Cj*) was determined to be 32 µM ± 5.3 µM (**Fig. 3C**) under assay conditions with competing UDP-diNAcBac at 20 μM. In contrast, α-GlcNAc-CF_2_ (**4**) showed minimal inhibition of the polyPGT WecA (*Tm*) relative to the parent UBP (**3**) (**Fig. 3A**), consistent with the hypothesis that the UBP-CXY moiety supports a different mode of binding for polyPGTs and highlighting the catalytic divergence between PGT superfamilies.^8, 9^

**Figure 3.**
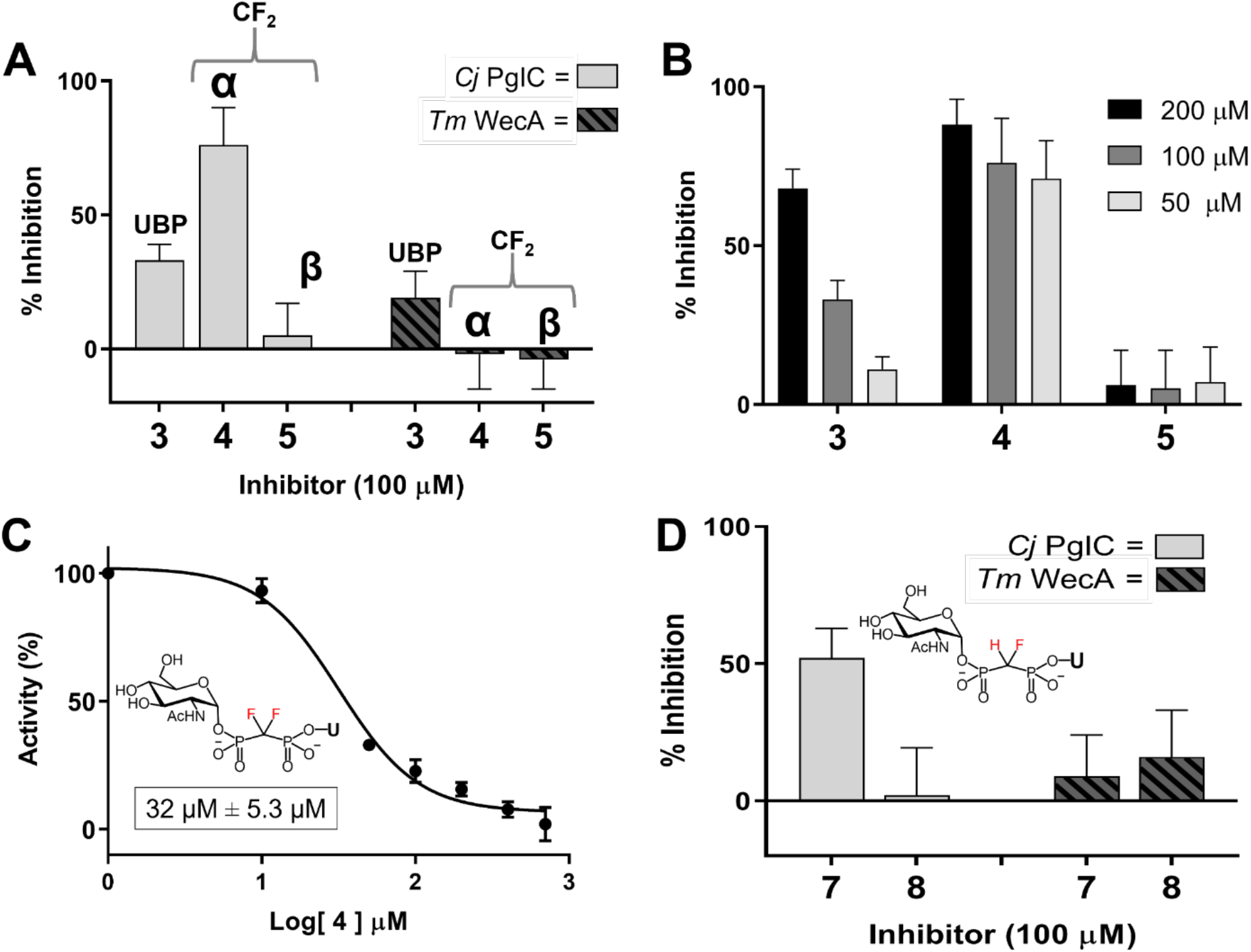
Activity of PglC (*Cj*) and WecA (*Tm*) measured after incubation with UBP probes followed by a reaction with UDP-diNAcBac (for PglC) and UDP-GlcNAc (for WecA), in reference to a control with no inhibitor. Solid bars represent the inhibition of PglC (*Cj*) activity. Striped bars represent the inhibition of WecA (*Tm*) activity. **A**. Inhibition of UBPs comprising CXY = F/F with R substituent = H (**3**), α-GlcNAc (**4**) and β-GlcNAc (**5**). **B**. Varied inhibitor concentration with activity of PglC (*Cj*) measured by incubation with the UBP probes comprising CXY = F/F, followed by a reaction with UDP-sugar substrate in reference to a control with no inhibitor. Black bars represent 200 µM inhibitor, dark grey bars represent 100 µM inhibitor, and light grey bars represent 50 µM inhibitor. Error bars indicate mean ± SD; n = 3. **C**. The IC_50_ value of **4** with PglC (*Cj*) was measured after 10 min incubation with **4** at varied concentrations (0 µM – 700 µM), followed by a reaction with UDP-diNAcBac substrate (20 µM) and quenched after 10% of the UMP product is formed (15 min). The IC_50_ value was found to be 32 µM ± 5.3 µM. **D**. Inhibition of UBPs comprising CXY = H/F chiral center with R substituent = α-GlcNAc (compounds **7** and **8**, defined as (*S*)-CHF and (*R*)-CHF, respectively). Data was plotted using GraphPad Prism as percentage remaining activity compared to no inhibitor, Error bars indicate mean ± SD; n = 3.

The micromolar IC_50_ of α-GlcNAc-CF_2_ (**4**) prompted an investigation into the specific effect of the fluorine atoms in the asymmetric environment of the PGT active sites. The contribution from each fluorine atom in the CXY moiety was parsed out using stereochemically-defined UBP analogues with the R substituent as α-GlcNAc, namely: α-GlcNAc-(*S*)-CHF (**7**) and α-GlcNAc-(*R*)-CHF (**8**). **Figure 3D** shows the comparison of the percent inhibition of α-GlcNAc-(*S*)-CHF (**7**) and α-GlcNAc-(*R*)-CHF (**8**) at 100 µM towards PglC (*Cj*) and WecA (*Tm*). The results demonstrate a strong preference for the (*S*)-CHF stereochemistry of α-GlcNAc-(*S*)-CHF (**7**) for PglC (*Cj*) (52% ± 10% inhibition) *vs* (*R*)-CHF of α-GlcNAc-(*R*)-CHF (**8**) (<2% inhibition). In contrast, WecA (*Tm*) demonstrates distinct binding preferences for these UBP probes, with α-GlcNAc-(*R*)-CHF (**8**) showing very slightly preferential inhibition of α-GlcNAc-(*R*)-CHF (**8**) relative to α-GlcNAc-(*S*)-CHF (**7**) (**Fig. 3D**). Notably, with respect to PglC (*Cj*), the percent inhibition of α-GlcNAc-(*S*)-CHF (**7**) (52% ± 10% inhibition) is not additive with total percent inhibition from α-GlcNAc-CF_2_ (**4**) (76% ± 13%), suggesting additional physicochemical contributions, for example effects on the pK_a_s, from the more electronegative CF_2_ moiety that may drive binding beyond a specific interaction of the C-F moiety with the enzyme.

Due to the improved binding of GlcNAc-modified UBPs, such as **4**, relative to the simple UBP analogues, we investigated PGT inhibition with analogues having a methylene bridge, CXY = H/H. The BP analogue H-CH_2_ (**10**), with no R substituent, shows very limited inhibition of PglC (*Cj*) and WecA (*Tm*). However, comparison of H-CH_2_ (**10**) with the α-GlcNAc modification (**11**), reinforces that the combination of CXY moiety with an α-linked sugar also supports a substrate-like binding mode, in this case, for a polyPGT. α-GlcNAc-CH_2_ (**11**) is the best inhibitor of WecA (*Tm*) with 48% ± 7% inhibition, compared to 28% ± 12% inhibition with PglC (*Cj*) at 100 μM. The IC_50_ value of α-GlcNAc-CH_2_ (**11**) with WecA (*Tm*) was found to be 41 µM ± 10 µM (**Figure S3**).

Overall, the inhibition of PglC and WecA by H-CCl_2_ (**1**), α-GlcNAc-CCl_2_ (**2**), H-CF_2_ (**3**), α-GlcNAc-CF_2_ (**4**), α-GlcNAc-(*S*)-CHF (**7**), α-GlcNAc-(*R*)-CHF (**8**), H-CH_2_ (**10**), and α-GlcNAc-CH_2_ (**11**) are graphed and compared (**Figure S4**). The relative affinities of all the UBP-sugar analogues are summarized in **Figure 4**.

**Figure 4.**
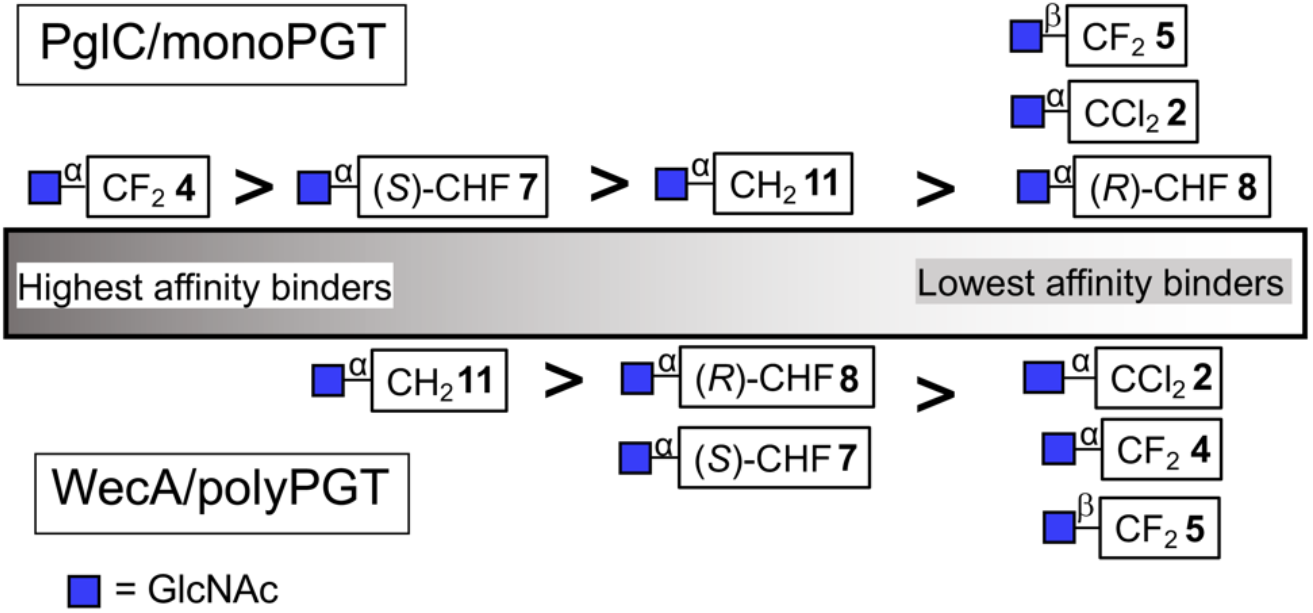
Comparison of inhibition by GlcNAc-UBP derivatives.

## Discussion

The synthetic chemistry towards bisphosphonate analogues of NDP-sugars presented here is modular and can be adapted to prepare many other nucleotide-sugar mimics with tunable P-CXY-P bisphosphonate moieties, different nucleobases and various carbohydrates or carbohydrate bioisosteres.

The relative inhibitory properties of the UBP-sugar analogues with the two PGT superfamilies are illustrated graphically in **Figure 4**. In general, although the simple uridine 5’-bisphosphonates (CXY-UBP) showed limited inhibition of both PGT superfamilies, the initial analysis with these analogues served as a guide to prioritize further elaboration into GlcNAc-modified nucleoside analogues. Elaboration of the CF_2_-UBP with an α-linked GlcNAc provides the best inhibitor of the monoPGT PglC (*Cj*) (IC_50_ 32 µM ± 5.3 µM, **Fig. 3C**). The improvement in inhibition contrasts the complete loss of activity with the β-linked GlcNAc analog, supporting that substrate mimicry is essential for binding. This observation highlights opportunities for improving monoPGT inhibition, for example through integration of diNAcBac, which is the carbohydrate in the native PglC substrate in place of GlcNAc. Although UDP-GlcNAc is a far poorer substrate for PglC than UDP-diNAcBac with a K_M_ value ∼20 fold higher, the manipulation of diNAcBac is synthetically more challenging and there are no commercial sources for this rare sugar or any of its derivatives. Alternatively, the modular synthetic approach readily enables synthesis of modified UBP conjugates that include moieties that substitute for the sugar. For example, in glycan-binding proteins recognition of sugar substrates is often promoted by electrostatic interactions wherein an electropositive C−H in carbohydrates interacts with electron-rich aromatic amino acids yielding CH−π interactions.^45^ Thus a carbohydrate substitute could mimic these CH-π interactions while also affording simplified synthetic routes. When occupying the sugar binding site, aryl groups can act as sugar bioisosteres by participating in similarly favorable π - π stacking interactions.

As illustrated in **Figure 4**, the mono- and polyPGTs show distinct preferences for different UBP-sugar analogues and although GlcNAc-CF_2_-UBP (**4**) is the preferred inhibitor for the monoPGT PglC (*Cj*), GlcNAc-CH_2_-UBP (**11**) is the preferred inhibitor for the polyPGT WecA (*Tm*). Both GlcNAc-CCl_2_-UBP (**2**) and GlcNAc(C(CH_3_)_2_)-UBP (**9**) show very weak inhibition of the PGTs, however, we do not explicitly discuss these analogues in the trends due to the additional steric burden imposed at the bridging site of the bisphosphonate. Analysis of the trends for WecA (**11**>**8**>**7**>**4**), show that there is a correlation between the increasing basicity of the bisphosphonate moiety and the strength of inhibitor binding. This trend becomes evident when referencing the pK_a_ values of the CXY-BP moieties: CF_2_-BP has the lowest pK_a_ values, corresponding to the lowest relative basicity in its anionic forms, while CH_2_-BP and C(CH_3_)_2_-BP have the highest pK_a_ values and therefore the highest relative basicity. The IC_50_ of **11** for WecA (*Tm*) is 41 µM ± 10 µM (**Fig. S3**). The preferential binding to **11** suggests that the primary factor responsible for GlcNAc-UBP analogue binding is the structural Mg^2+^ cofactor, which may principally be responsible for the stability of the ternary complex by playing a role in substrate binding and orientation. Specifically, increasing the basicity of the bisphosphonate moiety will strengthen the ionic interactions between the phosphonate oxygen anions and the Mg^2+^. The P−C bonds of bisphosphonates are ca. 0.2 A longer than the P–O bonds in pyrophosphate, however this is compensated by the P_α_−C−P_β_ angles, that are more acute than the P_α_−O−P_β_ angle of phosphates, resulting in similar distances between P_α_ and P_β_ atoms in pyrophosphate and bisphosphonates. This conclusion, based on a study of dNTP β,γ-bisphosphonates,^34^ should be valid for P,P’ diesters (data not shown). Thus, these structural differences should not significantly alter the Mg^2+^ coordination geometry.

Analysis of the structures of GPT (the human polyPGTs), which also features the conserved DD motif (**Fig. S190**) and uses UDP-GlcNAc as substrate, either in the presence of a UDP-GlcNAc-Mg^2+^ complex,^13^ or tunicamycin, a potent bisubstrate analog inhibitor^14^ (**Fig. S191A-D**) provide some insight. The structures show the UDP-GlcNAc-Mg^2+^ and tunicamycin coinciding in the bound state, but do not show the Mg^2+^ bound to the conserved aspartates to support a role in substrate orientation, but not a direct role in catalysis. This rationale is both consistent with its known function^17^ and our previously reported results where a similar correlation was observed between the basicity of bisphosphonate analogues and the binding affinity to Pol β.^46^

In contrast to the polyPGT, PglC (*Cj*) inhibition follows a different trend (**4**>**7**>**11**>**8**), with the highest inhibition (IC_50_ 32 µM ± 5.3 µM, **Fig. 3C**) observed with the least basic analogue (**4**). This alludes to the possibility that the role of Mg^2+^ in the monoPGT system is most important for catalysis in the ping-pong mechanism and plays a less significant role in substrate orientation. In proteins, such as Pol β, catalytic Mg^2+^ is known to exhibit a lower affinity to the protein in the active site compared to structural Mg^2+^.^46^ This suggests that the substrate orientations in complex with PglC are nonsuperimposable and ground state conformations of this complex are mainly defined by protein-ligand interactions.

The binding of **4** to PglC supports that the CF_2_ group is the best replacement of the oxygen atom in the parent UDP-sugar. However, this does not give information on the contribution of each C-F bond and whether there may be specific interactions with the target enzyme, or, if the effect is principally one of modulating the electronic and structural properties of the analog relative to the parent UDP-GlcNAc. To address this issue, we investigated the individual effects of the diastereotopic C-F bonds in **7** and **8** by assessing the inhibitory activity with stereochemically-defined α-GlcNAc-CHF-UBPs. The significant difference between (*S*)-CHF (**7**) and (*R*)-CHF (**8**) (**7**>>**8**) is noteworthy. We considered two hypotheses. First, that the C-F in **7** may form a specific interaction upon binding to PglC. In this case, a possible amino acid candidate is PglC residue Lys59 (**Fig. 1B**). Lys59 is a catalytically-essential residue, which in the current mechanistic proposal, based on the structure (PDB: 5WL7), would be close to the diphosphate oxygen to protonate the UMP leaving group upon formation of the covalent intermediate.^9, 18^ Alternatively, the (*S*)-CHF (**7**) and (*R*)-CHF (**8**) isomers might preferentially adopt different solution state conformations. The solution-state conformations of UDP-GlcNAc, (*S*)-CHF (**7**), and (*R*)-CHF (**8**) in water in the presence of MgCl_2_ were assessed by nuclear Overhauser effect (NOE) analysis (SI NMR summary and **Figs. S188** and **S189**). Under these conditions, all three compounds have uracil protons in an observable NOE range (< 5 Å) to many of the GlcNAc protons, indicating a collapsed structure. The key difference between these three structures arises with the interproton distances between uracil and the *N*-acetyl protons on the GlcNAc. In UDP-GlcNAc and (*S*)-CHF (**7**) the *N*-acetyl protons are within range to observe clear NOEs with the uracil C6 proton. The (*R*)-CHF (**8**) also has the acetyl group protons within range, albeit with a much weaker signal. This difference is significant enough to conclude that **7** and **8** have different favored conformations in solution, and that **7** behaves similarly to UDP-GlcNAc in solution. A possible outcome of conformational differences between **7** and **8** may be that Mg^2+^-bound **7**, like UDP-GlcNAc, may be preorganized into a favorable state for binding to the monoPGT leading to improved affinity relative to **8**. In contrast to the results with PglC, there is no significant difference between the inhibitory activity of **7** and **8** with WecA. Future structural and computational studies will be applied to further investigate the observed variations in binding to the PGT superfamilies.

## Conclusions

Nucleoside diphosphate sugar (NDP-sugar) substrates feature in innumerable cellular processes and there is considerable interest in the development and study of non-hydrolyzable analogues for applications in structural biology, mechanistic enzymology, and as leads for inhibitor development. In this study we have focused on phosphoglycosyl transferases, which catalyze the first membrane-committed step in many glycoconjugate assembly pathways. PGTs play pivotal roles in initiating production of diverse glycans that are essential for bacterial survival and pathogen-host interactions. In bacteria, PGTs and their NDP-sugar substrates are far more varied than in eukaryotes, reflecting the greater diversity of glycoconjugates in unicellular microorganisms and the potential for selective antibiotic and probe development.

We have presented the synthesis and biochemical analysis of a panel of uridine 5’-bisphosphonate (CXY-UBP) and uridine 5’-bisphosphonate-*N*-acetylglucosamine (GlcNAc-CXY-UBP) analogues of the UDP-sugar substrates of phosphoglycosyl transferases. The analogues feature a central substituted methylene group that can be tuned to modify the steric and electronic properties of the bridging bisphosphonate. The two PGT superfamilies are differentiated by mechanism and involve either - the direct attack by UndP on the β-phosphate of a UDP-sugar for the polyPGTs, or attack by the conserved aspartic acid residue on the UDP-sugar for the monoPGTs. The polyPGT and monoPGT superfamilies are also distinguished by highly divergent 3D structures. Together, the studies presented here underscore the mechanistic dichotomy of the PGT superfamilies and provide a pathway towards selective inhibition of either the prokaryotic monoPGT superfamily or the polyPGT superfamily found across domains of life. As the monoPGTs are exclusively prokaryotic,^11^ these enzymes represent potential new targets for the development of antibiotic and antivirulence agents due to the pivotal roles played by the complex glycoconjugates that are biosynthesized in pathways that are initiated by monoPGTs in bacteria.

## ASSOCIATED CONTENT

### Supporting Information

Materials and methods including enzyme expression and purification, biochemical assays, detailed synthetic procedures and full structural characterization of all new compounds, summary of NOE studies, and sequence and structure comparison of polyPGTs.

## AUTHOR INFORMATION

### Author Contributions

‡These authors contributed equally.

## NOTES

The authors declare no competing financial interest.

## ACKNOWLEDGMENTS

The authors thank Christine Arbour (MIT) and Inah Kang (USC) for assistance with reviewing and editing the manuscript and Walt Massefski and Bruce Adams of the MIT Department of Chemistry Instrumentation Facility (DCIF) for assistance with NOE studies.

## ABBREVIATIONS

PGT: phosphoglycosyl transferase
monoPGT: monotopic PGT
polyPGT: polytopic PGT
NDP: nucleoside diphosphate
UDP: uridine diphosphate
BP: bisphosphonate
UBP: Uridine bisphosphonate
*Cj*: *Campylobacter jejuni*
*Cc*: *Campylobacter concisus*
*Tm*: *Thermotoga maritima*
GlcNAc: *N*-acetyl glucosamine
Und-P: undecaprenyl phosphate
diNAcBac: diacetylbacillosamine
Pgl: protein glycosylation

