## Supplementary Material for "Uridine Bisphosphonates Differentiate Phosphoglycosyl Transferase Superfamilies"

#### Materials and methods

Uridine (99%) and D-(+)-glucosamine hydrochloride (>98%) were purchased from Alfa-Aesar. All other reagents were purchased from Sigma-Aldrich, TCI chemicals, Fluka or Oakwood Chemical (reagent grade) and used as received. All phosphonic esters and bisphosphonic acids were prepared according to the literature.[1-3] Chiral synthons were isolated and characterized according to our previously published methods.[4] Purifications of tetraalkyl bis(phosphonate) esters were performed using an ISCO Combi*Flash*Rf+ Lumen flash chromatography system equipped with an ELSD detector. $^{1}H$, $^{31}P$, $^{13}C$, $^{19}F$ NMR, HSQCAD and COSY spectra were obtained on a Varian 400-MR, VNMRS-500 or VNMRS-600 spectrometer. All $^{1}H$ and $^{13}C$ peak assignments were verified by COSY and HSQCAD, respectively. $^{31}P$ NMR spectra were proton-decoupled. Multiplicities are quoted as singlet (s), doublet (d), triplet (t), unresolved multiplet (m), doublet of doublets (dd), doublet of doublet of doublets (ddd), doublet of triplets (dt), triplet of doublets (td), and broad (br). All chemical shifts (δ) are reported in parts per million (ppm) relative to residual $CHD_2OD$ in $CD_3OD$ (δ 3.34, $^{1}H$ NMR), $CHCl_3$ in $CDCl_3$ (δ 7.26, $^{1}H$ NMR), HDO in $D_2O$ (δ 4.80, $^{1}H$ NMR), external 85% $H_3PO_4$ (δ 0.00, $^{31}P$ NMR), external $C_6F_6$ (δ -164.9, $^{19}F$ NMR) or external $Si(CH_3)_4$ (δ 0.00, $^{13}C$ NMR). NMR samples in $D_2O$ were adjusted to pH 10 or higher (using sodium carbonate) which is critical to obtain high resolution $^{31}P$ NMR spectra[5] (unless stated otherwise). The pH meter measurements were calibrated at three different pH values (4, 7, and 10) using standard buffers. NMR spectra processing was performed with MestReNova 11.0.2. Preparative HPLC was performed using a Shimadzu Prominence instrument equipped with a Shimadzu SPD-20A UV detector (0.5 mm path length) with detection at 260 nm for uridine bisphosphonate derivatives (CXY-UBP) and all GlcNAc-CXY-UBP. Strong Anion Exchange (SAX) HPLC was performed on a Macherey Nagel 21.4 mm × 150 mm SP15/25 Nucleogel column. Reversed phase HPLC was performed on a Hamilton PRP-1 21.2 mm × 250 mm (7 µm) column. High-resolution mass spectrometry (HRMS) was performed on a Water Synapt G2-Si ESI spectrometer (performed at the School of Chemical Sciences Mass Spectrometry Laboratory (MSL) at the University of Illinois) and low-resolution mass spectrometry on a Finnigan LCQ Deca XP Max mass spectrometer equipped with an ESI source, both in the negative ion mode. MS m/z values were calculated using ChemDraw 15.0.0.106 or iMass 1.3. Compound IUPAC names were assigned using MarvinSketch 16.12.12. The molar yields of the final products (GlcNAc-CXY-UBP and CXY-UBP) were estimated by UV absorbance referenced to the extinction coefficient of GlcNAc-UDP at 262 nm ($\varepsilon = 9780\ M^{-1}\ cm^{-1}$, pH 7).[6] The 'slow' and 'fast' HPLC peak descriptors reflect elution order of individual diastereomers (*R/S*)-α-GlcNAc(OAc)$_3$-CHF-UBP, **25c-α1** (*S*)-CHF isomer/**25c-α2** (*R*)-CHF isomer, on the RP-HPLC column.

#### Enzyme expression and purification

##### Expression and purification of PglCs

The expression and purification of each homolog of PglC was carried following the protocol as described previously.[7] Presence of protein was verified by SDS-PAGE.

Sequence of His$_6$-SUMO-PglC from *Campylobacter concisus* (*Cc*):

MGHHHHHHGSLQDSEVNQEAKPEVKPEVKPETHINLKVSDGSSEIFFKIKKTTPLRRL-
MEAFAKRQGKEMDSLRFLYDGIRIQADQAPEDLDMEDNDIIEAHREQIGGSGSGMYRNFLKRVIDILGALFLLILTSPIIIA
TAIFIYFKVSRDVIFTQARPGLNEKIFKIYKFKTMSDERDAN-
GELLPDDQRLGKFGKLIRSLSLDELPQLFNVLKGDMSFIGPRPLLVEYLPIYNETQKHRHDVRPGITGLAQVNGRNAISW
EKKFEYDVYYAKNLSFMLDVKIALQTIEKVLKRSGVSKEGQATTEKFNGKN

Sequence of His$_6$-SUMO-PglC from *Campylobacter jejuni* (*Cj*):

MGHHHHHHGSLQDSEVNQEAKPEVKPEVKPETHINLKVSDGSSEIFFKIKKTTPLRRL-
MEAFAKRQGKEMDSLRFLYDGIRIQADQAPEDLDMEDNDIIEAHREQIGGGGGG MYEKVFKRIFDFILALVLLVLFSPV
ILITALLLKITQGSVIFTQNRPGLDEKIFKIYKFKTMSDERDEKGELLSDELRLKAF-
GKIVRSLSLDELLQLFNVLKGDMSFVGPRPLLVEYLPLYNKEQKLRHKVRPGITGWAQVNGRNAISWQKKFELDVYYVK
NISFLLDLKIMFLTALKVLKRSGVSKEGHVTTEKFNGKN

##### Expression and purification of WecA

The expression and purification of WecA from *Thermotoga maritima* was carried following the protocol as described previously.[7] All the fractions were confirmed by SDS-PAGE. Given the challenges associated with the purification of proteins containing multiple TMHDs only partial purity was achieved for WecA. This was found to be consistent with previous literature.[7]

Sequence of GB1-WecA-His$_6$ from *Thermotoga maritima*:

MQYKLALNGKTLKGETTTEAVDAATAEKVFKQYANDNGVDGEWTYDDAT-
KTFTVTEGSMWEAIISFFLTSVLSVFAKKTEFLDRPDSRKSHGRAVPPVGGVSIFLTLLIFERDNPFFLFSIPLFLLGLLDDLF
DLSYRIKLAVTALVAVWFSTAVTIEVSIFGARIHPVFFVIWFVGMVNAFNVVDGLD-
GLLSGISLFSSLMIGERSLAFSIIGFLPWNLPDAKVFLGNSGSFLLGAYLSTASVVFFEGDLGYATLFLGFPFYEIVFSFVRRL
VVKKNPFSPDEKHTHHVFSRKIGKWKTLLILVSFSLMFNLLGLSQKFYFIFLYVVLCCVLLFTYCVLQR
GNGNLKLLEHHHHHH

#### Biochemical assays

**UMP Glo™ biochemical assays.** PGT assays were performed using the UMP-Glo™ assay [Promega cat. VA1130], which detects UMP release from the phosphoglycosyl transferase reactions and provides a quantitative luminescence output. Off-target inhibition of the UMP-Glo™ reagent enzymes by the inhibitors was tested and values were mathematically corrected. To correct for off-target inhibition, the UMP-Glo™ assay control experiments were first conducted in the presence and absence of inhibitors at 1 µM UMP. These concentrations of UMP represent the amount of nucleotide released in a typical assay. Using this information, the percent of background inhibition was used to adjust the luminescence readout. The quenching solution was prepared as described by Promega specifications. Specific assay conditions for each enzyme are described below.

Biochemical validation of substrate selection for PglC (*Cj*) and PglC (*Cc*) has been previously described.[8, 9]

**Cc and Cj PglC inhibition**: The assay conditions included 50 mM HEPES, 100 mM NaCl, pH 7.5, 5 mM $MgCl_2$ and 0.1% triton X-100. The reactions were performed with 20 µM Und-P, 20 µM UDP-diNAcBac and (0.3 nM PglC (*Cc*), 5 nM PglC (*Cj)*). The volume of the assay was 10 µL and conducted at ambient room temperature. Catalytically active PglC followed previously reported activity under the same assay conditions.[7] Inhibitors were added at a final concentration of 200 µM, 100 µM, and 50 µM with a final DMSO concentration of 10%. PglC was pre-incubated in the reaction mixture lacking UDP-diNAcBac for 10 min at RT. Upon the addition of UDP-diNAcBac, the reaction was allowed to proceed for a time within the predetermined linear range before the addition of an equal volume of quenching solution (*Cc* 10 min, *Cj* 4.5 min). The reaction mixture was transferred to a 96-well plate (white, nonbinding surface, Corning). The luminescence measurements were carried out using a SynergyH1 multi-mode plate reader (Bio Tek). The 96-well plate maintained at 25 $^{0}$C and shaken in the double orbital mode at 237 cpm for 16 min followed by incubation for 44 min, after which time the luminescence was measured. Luminescence units were converted to UMP amounts from a standard curve that was made following by the manufacturer's protocol.

**WecA inhibition**: The assay conditions included 100 mM Tris•HCl, pH 8, 10 mM $MgCl_2$, and 0.1% triton X-100. The reactions were performed with 500 µM Und-P, 100 µM UDP-GlcNAc, and 15 nM WecA. The volume of the assay was 10 µL and conducted at 40 °C. Catalytically active WecA demonstrated similar activity with previously reported assay conditions.[7, 10] Inhibitors were added at a final concentration of 200 µM, 100 µM, and 50 µM with a final DMSO concentration of 10%. WecA was pre-incubated in the reaction mixture lacking UDP-GlcNAc for 10 min at 40 °C. Upon the addition of UDP-GlcNAc, the reaction was allowed to proceed for a time within the predetermined linear range (15 min) before the addition of and equal volume of quenching solution. The reaction mixture was transferred to a 96-well plate (white, nonbinding surface, Corning). The luminescence measurements were carried out using a SynergyH1 multi-mode plate reader (Bio Tek). The 96-well plate maintained at 25 $^{o}$C and shaken in the double orbital mode at 237 cpm for 16 min followed by incubation for 44 min, after which time the luminescence was measured. Luminescence units were converted to UMP amounts from a standard curve that was made following by the manufacturer's protocol.

**Inhibition studies using radioactivity-based assay**

UDP-[$^3$H]diNAcBac synthesis and purification has been previously described.[11, 12] PglC (*Cj*) activity in the presence and absence of inhibitors was measured using a radioactive extraction-based assay.[13] Briefly, reactions contained 20 µM UndP, 3 µM UDP-[$^3$H]diNAcBac (500 dpm/pmol), and 6 nM SUMO-PglC (*Cj*) in a final volume of 100 µL of assay buffer (50 mM HEPES, pH 7.5, 100 mM NaCl, 0.1 % Triton X-100, 5 mM $MgCl_2$, and 10% DMSO). Inhibitors were added to a final concentration of 100 µM. PglC (*Cj*), UndP, and inhibitors were pre-incubated in assay buffer for 5 minutes. Reactions were started by addition of UDP-[$^3$H]diNAcBac and 15 µL aliquots were quenched into 1 mL of 2:1 $CHCl_3$:MeOH at 0.5, 1, 2, 3, 5, and 7 minutes. The organic layers were washed three times with 500 µL PSUP (Pure Solvent Upper Phase = 15 mL $CHCl_3$, 240 mL MeOH, 1.83 g KCl, 235 mL $H_2O$). The organic layers were mixed with 5 mL Opti-Fluor O (PerkinElmer), the combined aqueous layers were mixed with 5 mL EcoLite Liquid Scintillation Cocktail (MP Biomedicals), and all layers were analyzed on a Beckman Coulter LS6500 scintillation counting system. PglC (*Cj*) activity is reported as percentage of dpm in the organic layer normalized to the total amount of dpm per quench point. The percentage of dpm in the organic layer was also calculated in the same manner for a no-enzyme control, and this background value was subtracted from all results.

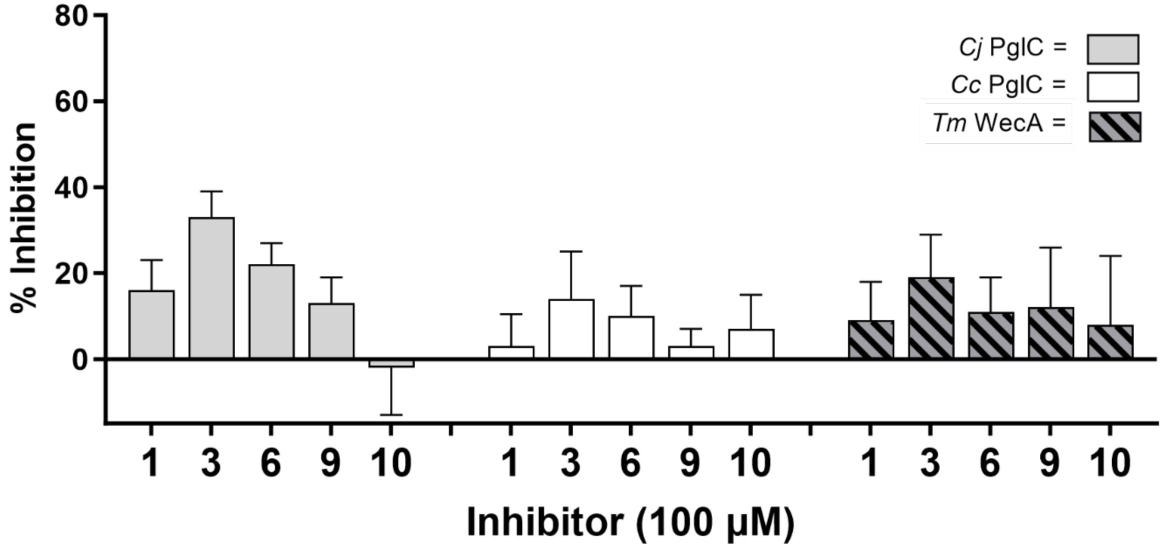

**Figure S1**. Activity of PglC (*Cj*), PglC (*Cc*) and WecA (*Tm*) measured by incubation with the UBP probes comprising CXY with the minimal substituent R = H, followed by a reaction with UDP-sugar substrate in reference to a control with no inhibitor. Solid bars represent the inhibition of each PglC activity. Striped bars represent the inhibition of WecA activity. Results for **3** were corroborated with the orthogonal radiation assay due to strong off-target inhibition of the assay enzymes.

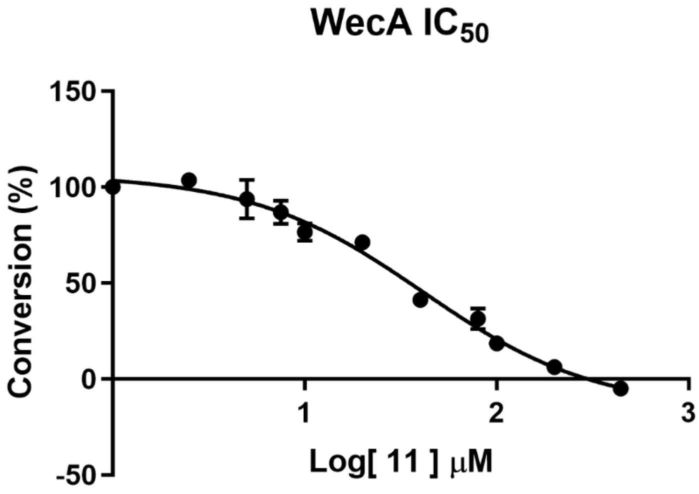

**Figure S2**. The $IC_{50}$ value of **11** with WecA (*Tm*) was measured after 10 min incubation with **11** at varied concentrations (0 μM – 450 μM), followed by a reaction with UDP-GlcNAc substrate (100 μM) and quenched after 10% of the UMP product is formed. The $IC_{50}$ value was found to be 41 μM ± 11 μM.

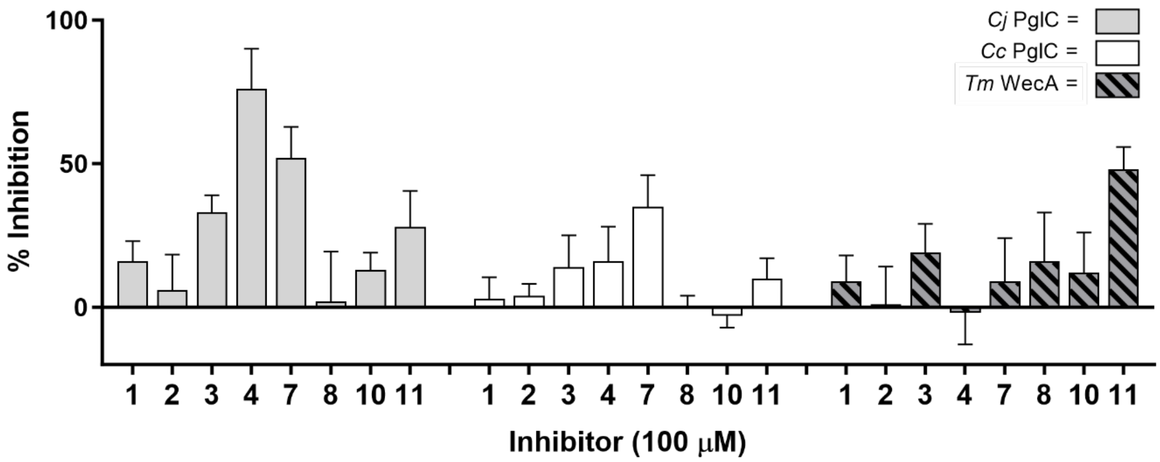

**Figure S3**. Activity of PglC (*Cj*), PglC (*Cc*) and WecA (*Tm*) measured by incubation with select UBPs, followed by a reaction with UDP-sugar substrate in reference to a control with no inhibitor. Solid bars represent the inhibition of each PglC activity. Striped bars represent the inhibition of WecA (*Tm*) activity.

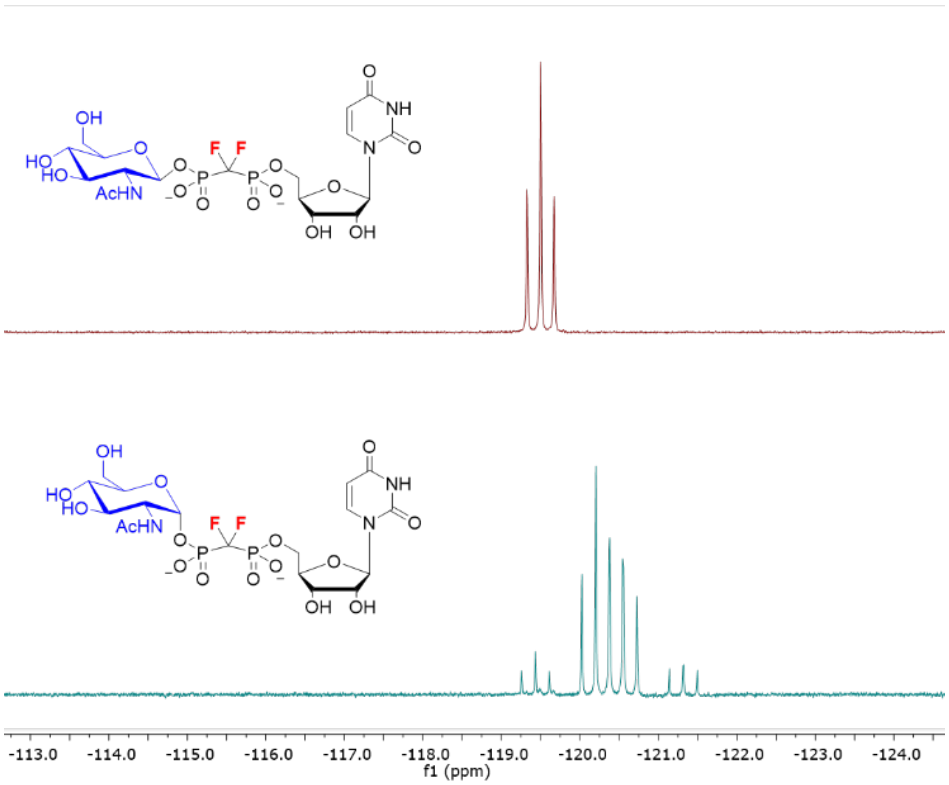

**Figure S4**. $^{19}$F NMR (470 MHz, $D_2O$, pH 7.5) of **4** and **5**.

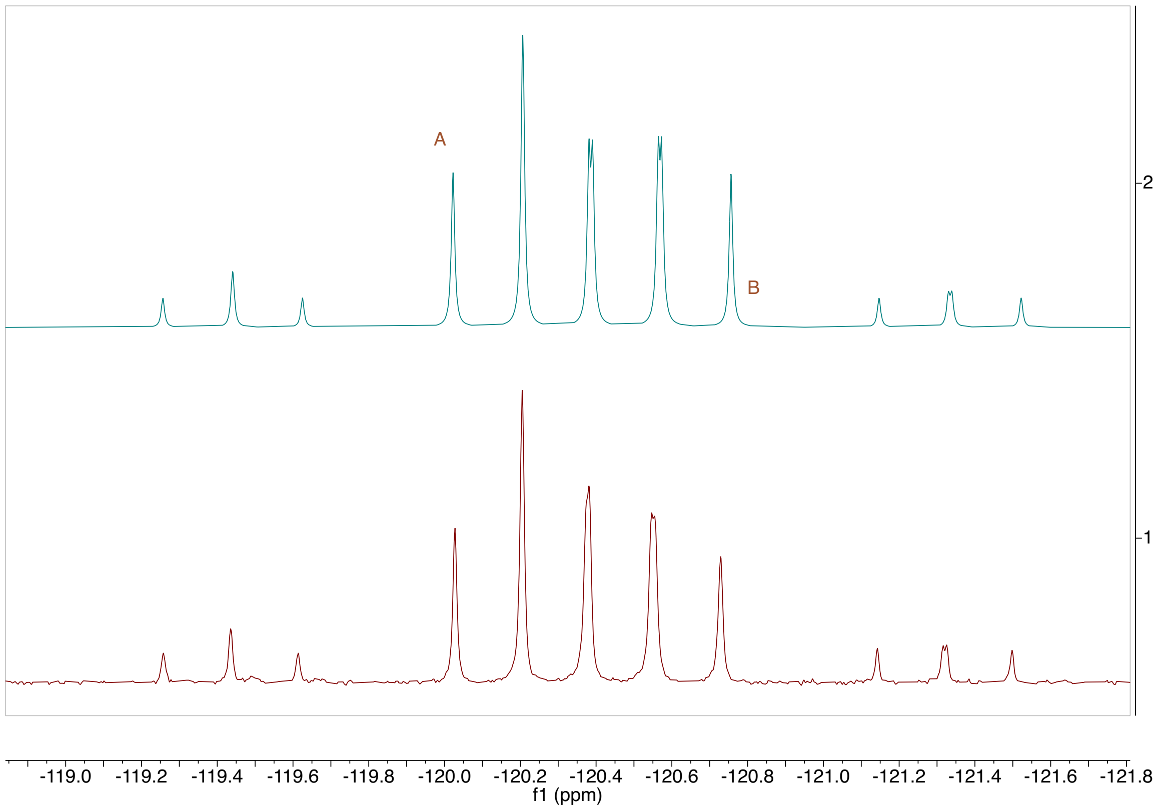

**Figure S5**. MestReNova simulated $^{19}$F NMR versus previously collected $^{19}$F NMR of **5**. Top is simulated. Bottom is the collected NMR spectrum.

Individual $^{19}$F NMR spectrum calculated using the following parameters:

A (-119.998 ppm): $J_{F1P1}$ = 86.0 Hz, $J_{F1P2}$ = 86.0 Hz, $J_{F1F2}$ = 360.0 Hz

B (-120.816 ppm): $J_{F2P1}$ = 86.5 Hz, $J_{F2P2}$ = 91 Hz

#### Synthesis of α-GlcNAc-$CH_2$-UBP analogue (11)

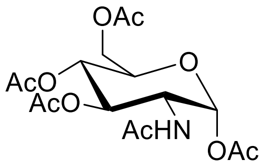

**[(3*S*,6*R*)-3,4,6-tris(acetyloxy)-5-acetamidooxan-2-yl]methyl acetate, GlcNAc(OAc)$_4$ 12.** To a solution of D-(+)-glucosamine hydrochloride (11.0 g, 51 mmol, 1 equiv) in anhydrous pyridine (50 mL) was added $Ac_2O$ (47.7 mL, 52.1 g, 510 mmol, 10 equiv) and stirred at room temperature for 12 h. After completion ($R_f$ = 0.23, hexane/EtOAc (1:3)) volatiles were removed under reduced pressure and the crude mixture was purified by automated flash column chromatography (equipped with ELSD detector) using stepwise gradient elution (0%−100% ethyl acetate in hexane) to give **12** as beige powder (23.7 g, 48.5 mmol, 95%). $^1$H NMR (400 MHz, $CDCl_3$) δ 6.17 (d, *J* = 3.8 Hz, 1H, H1'), 5.55 (br, 1H, NH), 5.29 – 5.16 (m, 2H, H3' and H4'), 4.52 – 4.45 (m, 1H, H2'), 4.25 (ABX system with $H_b$6' and H5'; appears as dd, *J* = 12.5, 4.1 Hz, 1H, $H_a$6'), 4.07 (ABX system with $H_a$6' and H5'; appears as dd, *J* = 12.5, 2.4 Hz, 1H, $H_b$6'), 4.02 – 3.95 (m, 1H, H5'), 2.19 (s, 3H, $CH_3$ in Ac), 2.09 (s, 3H, $CH_3$ in Ac), 2.06 (s, 3H, $CH_3$ in Ac), 2.05 (s, 3H, $CH_3$ in Ac), 1.94 (s, 3H, $CH_3$ in Ac). $^{13}$C NMR (101 MHz, $CDCl_3$) δ 171.53 (NC=O), 170.62 (OC=O), 170.00 (OC=O), 169.06 (OC=O), 168.63 (OC=O), 90.62 (C1'), 70.58 (C5'), 69.65 (C4'), 67.52 (C3'), 61.51 (C6'), 50.95 (C2'), 22.93 ($CH_3$), 20.86 ($CH_3$), 20.64 (2 $CH_3$), 20.51 ($CH_3$). Characterization data is consistent with the literature.[14]

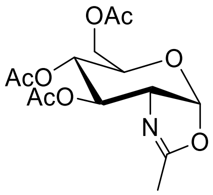

**[(3a*R*,6*S*)-6,7-bis(acetyloxy)-2-methyl-3aH,5H,6H,7H,7aH-pyrano[3,2-d][1,3]oxazol-5-yl]methyl acetate, Glc(OAc)$_3$-oxazoline 13.** To a solution of [(3*S*,6*R*)-3,4,6-tris(acetyloxy)-5-acetamidooxan-2-yl]methyl acetate **12** (2 g, 5.1 mmol, 1 equiv) in anhydrous DCM (10 mL) was added trifluoromethanesulfonic acid (HOTf, 1.35 mL, 2.3 g, 15.3 mmol, 3 equiv) and stirred at room temperature for 2 h under inert $N_2$ gas (colorless solution converted to dark solution). After completion (monitored by TLC, oxidation visualization) the solution was cooled in ice bath and TEA (3 mL) was added dropwise to neutralize the solution. The mixture was diluted with 10 mL DCM and washed with water (10 mL), phosphate buffer pH 7 (10 mL), and again water (4 × 10 mL) and dried over $MgSO_4$. DCM was removed under reduced pressure to obtain **13** as yellow oil (1.68 g, 5.1 mmol, quant.) and used freshly to the next step without purification. $^1$H NMR (500 MHz, $CDCl_3$) δ 5.98 (d, *J* = 7.3 Hz, 1H, H1'), 5.29 – 5.26 (m, 1H, H3'), 4.94 – 4.93 (m, 1H, H4'), 4.19 –

4.17 (m, 2H, H6’), 4.16 – 4.12 (m, 1H, H2’), 3.62 (dt, *J* = 9.0, 4.3 Hz, 1H, H5’), 2.12 (s, 3H, $CH_3$ in Ac), 2.11 (s, 3H, $CH_3$ in Ac), 2.10 (s, 3H, $CH_3$ in Ac), 2.09 (s, 3H, $CH_3$ in Ac). MS (ESI) m/z: $[M + H]^+$ calcd for $C_{14}H_{20}NO_8^+$ 330.1; found 330.0.

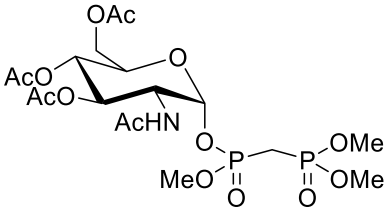

**[(3*S*,6*R*)-3,4-bis(acetyloxy)-6-({[(dimethoxyphosphoryl)methyl](methoxy)phosphoryl}oxy)-5-acetamidooxan-2-yl]methyl acetate, GlcNAc(OAc)$_3$-CH$_2$-Me$_3$BP 15.** Dimethyl [(dimethoxyphosphoryl)methyl]phosphonate ($Me_4$-$CH_2$-BP, 200 mg, 0.86 mmol) was dissolved in 1 mL of acetonitrile, followed by addition of TEA (5 mL, excess). The solution was allowed to reflux for 5.5 h and monitored by $^{31}P$ NMR (see Figure S16, 94% by NMR). The volatiles were removed under reduced pressure and residue was dissolved in 10 mL of $H_2O$ and unreacted tetramethyl ester BP was extracted with DCM (3 × 5 mL). The aqueous phase was passed through DOWEX $H^+$, dried under reduced pressure and co-evaporated with anhydrous dioxane (3 × 5 mL) to yield 168 mg (0.77 mmol, 90%) of **14** as pale-yellow oil. $^{31}P$ NMR (162 MHz, $CDCl_3$) δ 29.84 (d, *J* = 6.7 Hz), 8.64 (d, *J* = 6.7 Hz).

In a separate flask, freshly made **13** (425 mg, 1.29 mmol, 1.5 equiv) was co-evaporated with anhydrous dioxane (3 × 5 mL), dissolved in 3 mL anhydrous dioxane and added to **14** (168 mg, 0.77 mmol, 1 equiv). (Note: this reaction is moisture sensitive in which each molecule of water deactivates two oxazoline intermediates causing the formation of disaccharide via O-glycosidic bond; observed by MS). This mixture was stirred overnight at 55 °C under nitrogen gas. The progress of reaction monitored by ($^{31}P$ NMR, a mixture of two diastereomers (3:1) appear as four doublets: δ 18.2-21.8). After completion, crude mixture was concentrated under reduced pressure and purified by automated flash column chromatography (equipped with ELSD detector) using stepwise gradient elution (see **Figure S12**; 0 to 5 min, 0%–100% EtOAc in hexane; 5 to 20 min, 100% EtOAc; 20 to 30 min, 10% MeOH in EtOAc) to give **15** (398 mg, 0.73 mmol, 95%) as a mixture of two diastereomers a/b (3:1). $^1H$ NMR (400 MHz, $CDCl_3$) δ 8.01 (d, *J* = 9.7 Hz, 1H, NH of isomer a), 7.64 (d, *J* = 9.6 Hz, 1H, NH of isomer b), 5.86 – 5.81 (m, 1H, H1’ of isomer a), 5.67 – 5.61 (m, 1H, H1’ of isomer b), 5.34 – 5.25 (m, 1H, H3’ of both isomer a and b), 5.21 – 5. 14 (m, 1H, H4’ of both isomer a and b), 4.64 – 4.55 (m, 1H, H2’ of isomer a), 4.55 – 4.47 (m, 1H, H2’ of isomer b), 4.36 – 4.22 (m, 1H , H5’ of both isomer a and b), 4.19 – 4.08 (m, 2H, H6’ of both isomer a and b), 3.91 – 3.76 (m, 9H, 3 $POCH_3$), 2.65 – 2.36 (m, 2H, $pCH_2p$ of isomer a and b), 2.09 – 1.97 (m, 12H, 4 $CH_3$ in Ac). $^{31}P$ NMR (162

MHz, $CDCl_3$) δ 21.78 (d, *J* = 8.6 Hz), 21.53 (d, *J* = 2.3 Hz), 21.35 (d, *J* = 2.3 Hz), 18.24 (d, *J* = 8.6 Hz). MS (ESI) m/z: $[M + Na]^+$ calcd for $C_{18}H_{31}NNaO_{14}P_2^+$ 570.1; found 570.2.

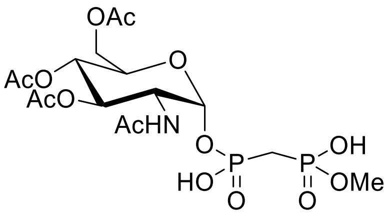

**[({[(2*R*,5*S*)-4,5-bis(acetyloxy)-6-[(acetyloxy)methyl]-3-acetamidooxan-2yl]oxy}(hydroxy)phosphoryl)methyl](methoxy)phosphinic acid, GlcNAc(OAc)$_3$-CH$_2$-MeBP 17.** To the solution of **15** (353 mg, 0.65 mmol, 1 equiv) in 4 mL anhydrous acetone was added 205 mg sodium iodide (1.36 mmol, 2.1 equiv; more NaI would be problematic in the next step DOWEX $H^+$ passage via conversion to HI which cleaves sugar-BP bond) and stirred overnight under $N_2$. The completion of the reaction was monitored by $^{31}P$ NMR and solvent was removed under reduced pressure to yield **16** as disodium salt (366 mg, 0.65 mmol, quant.). (Note: to remove unreacted sugar or disaccharide from the previous step (if any), the residue should be dissolved in 10 mL water and washed by EtOAc). $^1H$ NMR (400 MHz, $D_2O$) δ 5.47 (α-anomer, dd, *J* = 7.6 Hz to P, 3.5 Hz to H2', 1H, H1'), 5.24 (dd, *J* = 10.5, 9.8 Hz, 1H, H3'), 5.01 (t, *J* = 9.8 Hz, 1H, H4'), 4.38 – 4.30 (m, 2H, H6'), 4.23 (ddd, *J* = 10.5 Hz to H3', 3.5 Hz to H1', 1.8 Hz to P, 1H, H2'), 4.11 – 4.04 (m, 1H, H5'), 3.50 (d, *J* = 10.9 Hz, 3H, $POCH_3$), 2.19 – 2.05 (m, 2H, $pCH_2p$), 2.02 (s, 3H, $CH_3$ in Ac), 1.98 (s, 3H, $CH_3$ in Ac), 1.93 (s, 3H, $CH_3$ in Ac), 1.91 (s, 3H, $CH_3$ in Ac). $^{31}P$ NMR (202 MHz, $D_2O$, pH 7) δ 18.22 (d, *J* = 11.8 Hz, 1P), 16.44 (d, *J* = 11.8 Hz , 1P).

Conversion of **16** to the corresponding free acid **17** was carried outby passage through a column of DOWEX $H^+$ using cold MeOH:$H_2O$ (9:1), dried under reduced pressure and cold condition (in an ice bath). Note: since the sugar-phosphorous ester bond is fragile in acidic condition (confirmed by $^{31}P$ NMR), the DOWEX $H^+$ should be prewashed with DI water to completely remove the excess of HCl used for DOWEX $H^+$ preparation.

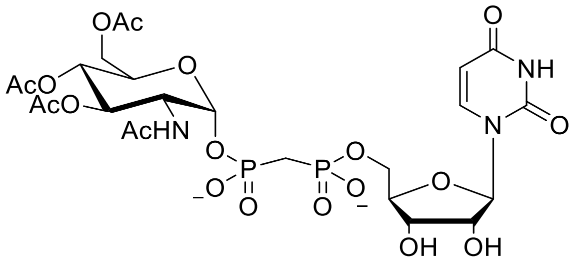

**[(2*R*,5*R*)-5-(2,4-dioxo-1,2,3,4-tetrahydropyrimidin-1-yl)-3,4-dihydroxyoxolan-2-yl]methyl {[(2*R*,5*S*)-4,5-bis(acetyloxy)-6-[(acetyloxy)methyl]-3-acetamidooxan-2-yl phosphono]methyl}phosphonate, Glc-NAc(OAc)$_3$-CH$_2$-UBP 20.** Compound **17** (230 mg, 0.44 mmol, 1 equiv) dried under reduced pressure by co-evaporation with anhydrous dioxane (3 × 2 mL) and left under reduced pressure until the total weight of the flask remained constant. Then it was redissolved in 4 mL of anhydrous dioxane and 0.5 mL anhydrous DMF, followed by sequential addition of uridine (**18**, 119 mg, 0.49 mmol, 1.1 equiv) and $PPh_3$ (176 mg, 0.67 mmol, 1.5 equiv). Distilled anhydrous diisopropyl azodicarboxylate (DIAD) (131 uL, 135 mg, 0.67 mmol, 1.5 equiv) was dissolved in 0.5 mL of anhydrous dioxane and added to the reaction mixture drop-wise under $N_2$. The reaction mixture was stirred under $N_2$ at rt overnight and monitored by MS (ESI) m/z: $[M-H]^-$ calcd for $C_{25}H_{36}N_3O_{19}P_2^-$ 744.2; found 744.3. Note: if necessary, more $PPh_3$ and DIAD should be added for completion of the reaction. After completion, volatiles were removed under reduced pressure to afford **19** which was used in the next step without further purification. Demethylation of **19** was carried out via addition of excess thiophenol (1.43 g, 1.34 mL, 13 mmol, 20 equiv) and DIPEA (1.68 g, 2.27 mL, 13 mmol, 20 equiv) mixture in 2 mL anhydrous *N*-methyl-2-pyrrolidone (NMP) and stirring for two days at 40 °C under $N_2$ atmosphere. When the reaction was completed (monitored by MS), the crude mixture was purified by dual-pass preparative HPLC: (1) Macherey-Nagel Nucleogel SAX 1000−10 (25 mm × 15 cm) column, using gradient mode (A/ 10% acetonitrile in $H_2O$ and B/ 10% acetonitrile in 0.5 M triethylammonium bicarbonate, pH 7.5; 0−7 min, 100% A; 7−10 min, 0%-25% B; 10−12.5 min, 25% B; 12.5−16 min, 25%-35% B; 16−20 min, 35% B; 20−25 min, 35%-75% B) at a flow rate of 8 mL/min (retention time = 17 min) with a UV detection (261 nm); (2) Hamilton PRP-1 column (7 µm, 250 mm × 21 mm) with 14% acetonitrile in 0.1 M triethylammonium bicarbonate, pH 7.5, at a flow rate of 8.0 mL/min using an isocratic mode (retention time = 17 min) with a UV detection (261 nm) to give desired product **20** (64.37 mg, 0.088 mmol, 20% two steps) as bis(triethylammonium) salts.

$^{31}$P NMR (162 MHz, $D_2O$, pH 7) δ 16.91 (d, *J* = 12.4 Hz, 1P), 16.00 (d, *J* = 12.4 Hz, 1P). $^1$H NMR (400 MHz, $D_2O$) δ 7.72 (d, *J* = 7.7 Hz, 1H, H6 in U), 5.87 (d, *J* = 4.2 Hz, 1H, H1’ in U), 5.73 (d, *J* = 7.7 Hz, 1H, H5 in U), 5.46 (α-anomer, dd, *J* = 7.7 Hz to P, 3.7 Hz to H2”, 1H, H1”), 5.26 – 5.20 (m, 1H, H3”), 5.00 (appear as t, *J* =

9.8 Hz, 1H, H4”), 4.40 – 3.97 (m, 9H), 2.23 – 2.11 (m, 2H, p$CH_2$p), 2.01 (s, 3H, $CH_3$ in Ac), 1.97 (s, 3H, $CH_3$ in Ac), 1.92 (s, 3H, $CH_3$ in Ac), 1.90 (s, 3H, $CH_3$ in Ac).

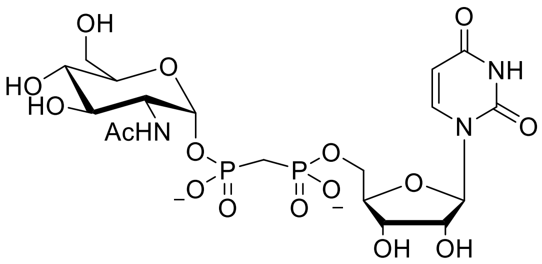

**(2*R*,5*S*)-3-acetamido-4,5-dihydroxy-6-(hydroxymethyl)oxan-2-yl ({[(2*R*,5*R*)-5-(2,4-dioxo-1,2,3,4-tetrahydropyrimidin-1-yl)-3,4-dihydroxyoxolan-2-yl]methyl phosphono}methyl)phosphonate, GlcNAc-$CH_2$-UBP 11.** To remove *O*-acetyl groups, compound **20** (26.3 mg, 0.036 mmol, defined by UV), was dissolved in 5 mL water, pH was adjusted to 11 with ammonium hydroxide ($NH_4OH$) and stirred for 1 h (monitored by MS). Then, final product was concentrated under reduced pressure and purified by RP-HPLC using a Hamilton PRP-1 column (7 µm, 250 mm × 21 mm) with 4% acetonitrile in 0.1 M triethylammonium bicarbonate, pH 7.5, at a flow rate of 8.0 mL/min using isocratic mode (retention time = 10 min) with a UV detection (261 nm) to give desired product **11** as bis(triethylammonium) salts (15 mg, defined by UV, 0.025 mmol, 70%).

$^{31}$P NMR (202 MHz, $D_2O$, pH 7) δ 16.34 (AB system appear as dd, *J* = 11.3 Hz). $^{1}$H NMR (500 MHz, $D_2O$) δ 7.72 (d, *J* = 7.7 Hz, 1H, H6 in U), 5.88 (d, *J* = 4.1 Hz, 1H, H1’), 5.76 (d, *J* = 7.7 Hz, 1H, H5 in U), 5.37 (α-anomer, dd, *J* = 7.9 Hz to P, 3.4 Hz to H2”, 1H, H1”), δ 4.25 – 4.21 (m, 2H, H2’ and H3’), 4.14 – 4.10 (m, 1H, H4’), 4.09 – 3.99 (m, 2H, H5’), 3.89 – 3.81 (m, 2H, H2” and H3”), 3.77 – 3.64 (m, 3H, H5” and H6”), 3.39 (t, *J* = 9.6 Hz, 1H, H4”), 2.09 (ABX system appears as tq, *J* = 19.9, 15.2 Hz, 2H, p$CH_2$p), 1.96 (s, 3H, $CH_3$ in Ac). HRMS (ESI-TOF) m/z: $[M - H]^-$ calcd for $C_{18}H_{28}N_3O_{16}P_2^-$ 604.0945; found 604.0942.

#### Synthesis of CXY-UBP analogues (CXY = $CH_2$, $CF_2$, $CCl_2$, $C(CH_3)_2$, (*R*/*S*)-CHF)

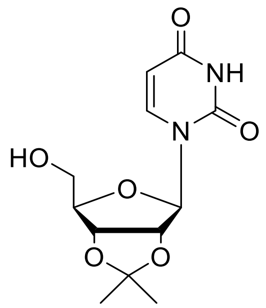

**1-[(4*R*,6*R*)-6-(hydroxymethyl)-2,2-dimethyl-tetrahydro-2H-furo[3,4-d][1,3]dioxol-4-yl]-1,2,3,4-tetrahydropyrimidine-2,4-dione, 2′,3′-*O*-isopropylidene-uridine 21.** The preparation of **21** was performed according to a published procedure with minor modifications.[15] 60 mg of *p*-toluene sulfonic acid monohydrate (*p*-TsOH) was coevaporated with anhydrous acetone (3 × 1 mL) and added to a solution of **18** (2 g, 8.19 mmol, 1 equiv) and 2,2-dimethoxypropane (8 mL, 64 mmol, 7.8 equiv) in 60 mL anhydrous acetone and refluxed at 85 °C. Yellow solution become dark during the progress of reaction. Completion of the reaction monitored by TLC (20% MeOH in DCM, uridine Rf: 0.3; **21** Rf: 0.73, UV visualization) and after 90 min volatiles were removed under reduced pressure. The crude mixture was purified by automated flash column chromatography using stepwise gradient elution (silica, solid load, 0%–30% methanol in DCM, **21** eluted out at 10% MeOH in DCM) to give the title compound as a pale yellow powder (2.3 g, 8.09 mmol, 98%).

$^1$H NMR (400 MHz, $CDCl_3$) δ 7.36 (d, *J* = 8.1 Hz, 1H, H6 in U), 5.73 (d, *J* = 8.1 Hz, 1H, H5 in U), 5.57 (d, *J* = 3.1 Hz, 1H, H1′), 5.00 (ABXX′ system appears as ddd, *J* = 31.9, 6.5, 3.2 Hz, 2H, H2′ and H3′), 4.29 (appears as q, *J* = 3.3 Hz, 1H, H4′), 3.98 – 3.72 (m, 2H, H6′), 1.58 (s, 3H, $CH_3$), 1.36 (s, 3H, $CH_3$). $^{13}$C NMR (101 MHz, $CDCl_3$) δ 162.57 (C4), 150.11 (C2), 142.80 (C6), 114.46 (OCO), 102.65 (C5), 96.12 (C1′), 86.78 (C4′), 83.51 (C2′), 80.26 (C3′), 62.69 (C6′), 27.26 ($CH_3$), 25.25 ($CH_3$). MS (ESI) m/z: $[M - H]^-$ calcd for $C_{12}H_{15}N_2O_6^-$ 283.1; found 283.1. Characterization data is consistent with the literature ($^{13}$C NMR was not provided).[16]

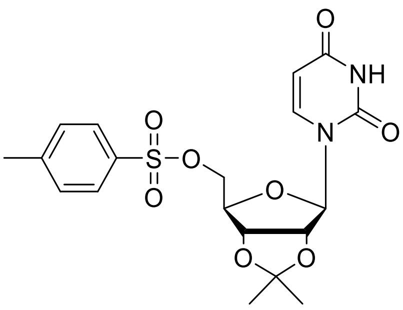

**[(4*R*,6*R*)-6-(2,4-dioxo-1,2,3,4-tetrahydropyrimidin-1-yl)-2,2-dimethyl-tetrahydro-2H-furo[3,4-d][1,3]dioxol-4-yl]methyl 4-methylbenzene-1-sulfonate, 2',3'-*O*-isopropylidene-5'-*O*-toluenesulfonyluridine 22.** The preparation of **22** was performed according to a published procedure with minor modifications.[17] To the solution of **21** (320 mg, 1.12 mmol, 1 equiv) in anhydrous pyridine (3 mL) at 0 °C was added freshly crystalized *p*-toluenesulfonyl chloride (*p*-TsCl, 279 mg, 1.46 mmol, 1.3 equiv) The initial yellow solution turned red during the progress of the reaction. Completion of reaction monitored by TLC (10% MeOH in DCM, **22** Rf: 0.65; **21** Rf: 0.36; UV visualization). Volatiles were removed under reduced pressure and the crude mixture was purified by automated flash column chromatography using stepwise gradient elution (silica, solid load, 0%–20% methanol in DCM, **22** eluted out at 5% MeOH in DCM; see Figure S32) to give the title compound as a pale yellow powder (464.8 mg, 1.06 mmol, 95%).

$^{1}$H NMR (500 MHz, $CDCl_3$) δ 9.49 (br, 1H, NH), 7.76 (d, *J* = 7.5 Hz, 2H, H2 in *p*-Ts), 7.32 (d, *J* = 7.4 Hz, 2H, H3 in *p*-Ts), 7.22 (d, *J* = 8.1 Hz, 1H, H6 in U), 5.70 (d, *J* = 8.1 Hz, 1H, H5 in U), 5.64 (br, 1H, H1'), 4.93 (appears as bd, *J* = 6.1 Hz, 1H, H2'), 4.80 – 4.75 (m, 1H, H3'), 4.36 – 4.31 (m, 1H, H5'), 4.27 (s, 2H, , H5' and H4'), 2.43 (s, 3H, $CH_3$ in *p*-Ts), 1.53 (s, 3H, $CH_3$ in 2',3'-*O*-isopropylidene), 1.32 (s, 3H, $CH_3$ in 2',3'-*O*-isopropylidene). $^{13}$C NMR (126 MHz, $CDCl_3$) δ 163.37 (C4 in U), 150.03 (C2 in U), 145.27 (C1 in *p*-Ts), 142.29 (C6 in U), 132.49 (C4 in *p*-Ts), 129.89 (2C, C3 in *p*-Ts), 127.95 (2C, C2 in *p*-Ts), 114.64 (OCO in 2',3'-*O*-isopropylidene), 102.71 (C5 in U), 94.88 (C1'), 85.06 (C4'), 84.33 (C2'), 80.83 (C3'), 69.33 (C5'), 27.01 ($CH_3$ in 2',3'-*O*-isopropylidene), 25.18 ($CH_3$ in 2',3'-*O*-isopropylidene), 21.66 ($CH_3$ in *p*-Ts). MS (ESI) m/z: $[M + H]^+$ calcd for $C_{19}H_{23}N_2O_8S^+$ 439.1; found 439.1. Characterization data is consistent with the literature.[18]

**General Method 1:** A general procedure for preparation of 2',3'-*O*-isopropylidene-uridine-5'-(methylene)bisphosphonate derivatives, **24**. The preparation of CXY-UBP analogues was performed according to a published procedure with minor modifications.[19] High concentrations of **22** (~1 M) and bisphosphonates are necessary for the reactions to proceed at reasonable rates. The coupling reactions were performed in rubber septum sealed vials (5-10 mL) equipped with magnetic stirring under nitrogen atmosphere at room temperature. A solution of appropriate methylenebis(phosphonic acid) derivative **23** in water/EtOH (1:1) was treated with tetra-*n*-butylammonium hydroxide dropwise until pH turned to 7-8.

The mixture was stirred at room temperature for 5 min. The solvents were removed under reduced pressure, co-evaporated with anhydrous acetonitrile (3 x 2 mL) and left under reduced pressure overnight for complete dryness. To the well-dried appropriate tris(tetra-*n*-butylammonium) methylenebis(phosphonate) (3 equiv) was added a solution of **22** (1 to 2 M, 1 equiv) in anhydrous acetonitrile. This viscous solution was stirred under nitrogen atmosphere at room temperature. The progress of the reaction was monitored by $^{31}$P NMR. Upon completion, volatiles were removed under reduced pressure, the residue was re-dissolved in water and purified by dual-pass preparative RP-HPLC, Hamilton PRP-1 column (7 µm, 250 mm × 21 mm), isocratic mode, using (13% - 15%) acetonitrile in 0.1 M triethylammonium bicarbonate, pH 7.5, at a flow rate of 8.0 mL/min and UV detection (261 nm).

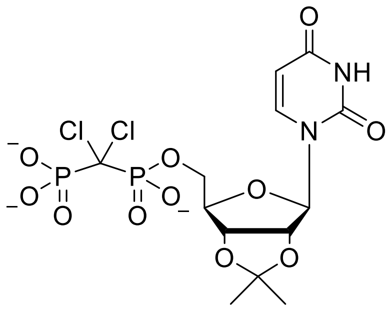

**[(4*R*,6*R*)-6-(2,4-dioxo-1,2,3,4-tetrahydropyrimidin-1-yl)-2,2-dimethyl-tetrahydro-2H-furo[3,4-d][1,3]dioxol-4-yl]methyl [dichloro(phosphono)methyl]phosphonate, 2’,3’-*O*-isopropylidene-$CCl_2$-UBP 24a.** According to General Method 1, to well-dried tris(tetra-*n*-butylammonium) hydrogen (dichloromethylene)bisphosphonate **23a** (1.8 mmol, 3 equiv) was added **22** (265 mg, 0.6 mmol, 1 equiv)in 600 uL anhydrous acetonitrile. The solution was stirred for 2 d (~ 85% conversion as identified by $^{31}$P NMR). The crude mixture was purified by dual-pass preparative RP-HPLC, Hamilton PRP-1 column (7 µm, 250 mm × 21 mm), isocratic mode, using 13.5% acetonitrile in 0.1 M triethylammonium bicarbonate, pH 7.5, at a flow rate of 8.0 mL/min and UV detection (261 nm). The title compound eluted at 12.4 min. After removal of solvents under reduced pressure and co-evaporation with water (3 x 2 mL), the desired product **24a** was obtained as bis(triethylammonium) salts (0.49 mmol, 81% defined by UV).

$^1$H NMR (400 MHz, $D_2O$) δ 7.83 (d, *J* = 8.1 Hz, 1H, H6 in U), 5.85 (d, *J* = 3.2 Hz, 1H, H1’), 5.80 (d, *J* = 8.1 Hz, 1H, H5 in U), 5.01 – 4.91 (ABXX’ system appears as m, 2H, H2’ and H3’), 4.46 – 4.42 (m, 1H, H4’), 4.25 – 4.20 (m, 2H, H5’), 1.49 (s, 3H, $CH_3$), 1.29 (s, 3H, $CH_3$). $^{31}$P NMR (162 MHz, $D_2O$, pH 10) δ 9.90 (d, *J* = 16.0 Hz, 1P), 7.69 (d, *J* = 16.0 Hz, 1P). MS (ESI) m/z: $[M – H]^–$ calcd for $C_{13}H_{17}Cl_2N_2O_{11}P_2^-$ 509.0; found 509.0.

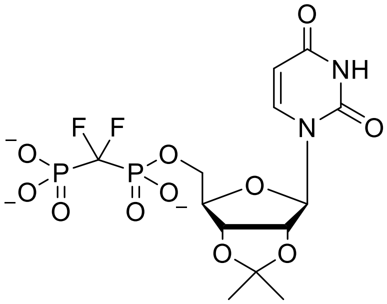

**[(4*R*,6*R*)-6-(2,4-dioxo-1,2,3,4-tetrahydropyrimidin-1-yl)-2,2-dimethyl-tetrahydro-2H-furo[3,4-d][1,3]dioxol-4-yl]methyl [difluoro(phosphono)methyl]phosphonate, 2',3'-*O*-isopropylidene-$CF_2$-UBP 24b.** According to General Method 1, to well-dried tris(tetra-*n*-butylammonium) hydrogen (difluoromethylene)bisphosphonate **23b** (1.8 mmol, 3 equiv) was added **22** (265 mg, 0.6 mmol, 1 equiv) in 600 uL anhydrous acetonitrile. The solution was stirred for 2 d (~ 80% conversion as identified by $^{31}$P NMR). The crude mixture was purified by dual-pass preparative RP-HPLC, Hamilton PRP-1 column (7 µm, 250 mm × 21 mm), isocratic mode, using 13% acetonitrile in 0.1 M triethylammonium bicarbonate, pH 7.5, at a flow rate of 8.0 mL/min and UV detection (261 nm). The title compound eluted at 11 min. After removal of solvents under reduced pressure and co-evaporation with water (3 x 2 mL), the desired product **24b** was obtained as bis(triethylammonium) salts (0.45 mmol, 75% defined by UV).

$^{1}$H NMR (400 MHz, $D_2O$) δ 7.77 (d, *J* = 8.1 Hz, 1H, H6 in U), 5.83 (d, *J* = 2.9 Hz, 1H, H1'), 5.79 (d, *J* = 8.1 Hz, 1H, H5 in U), 4.96 – 4.90 (ABXX' system appears as m, 2H, H2' and H3'), 4.45 – 4.41 (m, 1H, H4'), 4.17 – 4.12 (m, 2H, H5'), 1.48 (s, 3H, $CH_3$), 1.29 (s, 3H, $CH_3$). $^{31}$P NMR (202 MHz, $D_2O$, pH 7.5) δ 5.84 (td, *J* = 87.0, 53.2 Hz, 1P), 3.32 (td, *J* = 75.0, 53.2 Hz, 1P). $^{19}$F NMR (376 MHz, $D_2O$, pH 7.5) δ -117.41 (dd, *J* = 87.0, 75.0 Hz). MS (ESI) m/z: [M – H]$^-$ calcd for $C_{13}H_{17}F_2N_2O_{11}P_2^-$ 477.0; found 477.1.

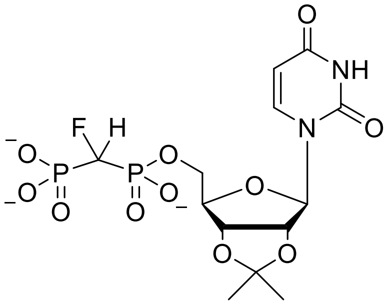

**[(4R,6*R*)-6-(2,4-dioxo-1,2,3,4-tetrahydropyrimidin-1-yl)-2,2-dimethyl-tetrahydro-2H-furo[3,4-d][1,3]dioxol-4-yl]methyl [fluoro(phosphono)methyl]phosphonate, 2',3'-*O*-isopropylidene-(*R*/*S*)-CHF-UBP 24c.** According to General Method 1, to well-dried tris(tetra-*n*-butylammonium) hydrogen (monofluoromethylene)bisphosphonate **23c** (1.8 mmol, 3 equiv) was added **22** (265 mg, 0.6 mmol, 1 equiv) in 600 uL anhydrous acetonitrile. The solution was stirred for 2 d (~ 85% conversion as identified by $^{31}$P NMR). The crude mixture was purified by dual-pass preparative RP-HPLC, Hamilton PRP-1 column (7 µm, 250 mm × 21 mm), isocratic mode, using 13% acetonitrile in 0.1 M triethylammonium bicarbonate, pH 7.5, at a flow rate of 8.0 mL/min and UV detection (261 nm). The title compound eluted at 11 min. After removal of solvents under reduced pressure and co-evaporation with water (3 x 2 mL), the desired product **24c** was obtained as bis(triethylammonium) salts (0.47 mmol, 79% defined by UV).

$^{1}$H NMR (500 MHz, $D_2O$) mixture of two diastereomers (1:1), δ 7.72 (d, *J* = 8.0 Hz, 1H, H6 in U), 5.87 (d, *J* = 2.5 Hz, 1H, H1'), 5.78 (d, *J* = 8.0 Hz, 1H, H5 in U), 4.97 – 4.90 (ABXX' system appears as m, 2H, H2' and H3'), 4.53 and 4.62 (each appears as td, *J* = 12.5, 5.6 Hz, 1H, pCHFp), 4.40 (appears as q, *J* = 3.4 Hz, 1H, H4'), 4.14 – 4.06 (m, 2H, H5'), 1.50 (s, 3H, $CH_3$), 1.30 (s, 3H, $CH_3$). $^{19}$F NMR (470 MHz, $D_2O$, pH 7.5) δ -217.32 (mixture of two isomers (1:1) appears as dddd, *J* = 63.8, 56.0, 45.4, 36.2 Hz). $^{31}$P NMR (202 MHz, $D_2O$, pH 7.5) mixture of two diastereomers (1:1), δ 13.79 (isomer 1: dd, *J* = 15.6, 12.1 Hz, 1P), 13.47 (isomer 2: dd, *J* = 15.7, 11.9 Hz, 1P), 6.94 (isomer 1: dd, *J* = 12.1, 5.5 Hz, 1P), 6.67 (isomer 2: dd, *J* = 11.9, 5.5 Hz, 1P). MS (ESI) m/z: $[M – H]^-$ calcd for $C_{13}H_{18}FN_2O_{11}P_2^-$ 459.0; found 459.2.

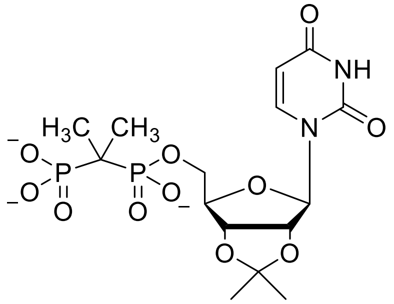

**[(4*R*,6*R*)-6-(2,4-dioxo-1,2,3,4-tetrahydropyrimidin-1-yl)-2,2-dimethyl-tetrahydro-2H-furo[3,4-d][1,3]dioxol-4-yl]methyl (2-phosphonopropan-2-yl)phosphonate, 2',3'-*O*-isopropylidene-C($CH_3$)$_2$-UBP 24d.** According to General Method 1, to well-dried tris(tetra-*n*-butylammonium) hydrogen (dimethylmethylene)bisphosphonate **23d** (1.8 mmol, 3 equiv) was added **22** (265 mg, 0.6 mmol, 1 equiv) in 600 uL anhydrous acetonitrile. The solution was stirred for 2 d (~ 100% conversion as identified by $^{31}$P NMR). The crude mixture was purified by dual-pass preparative RP-HPLC, Hamilton PRP-1 column (7 µm, 250 mm × 21 mm), isocratic mode, using 15% acetonitrile in 0.1 M triethylammonium bicarbonate, pH 7.5, at a flow rate of 8.0 mL/min and UV detection (260 nm). The title compound eluted at 7.9 min (see Figure S52). After removal of solvents under reduced pressure and co-evaporation with water (3 x 2 mL), the desired product 24 d was obtained as bis(triethylammonium) salts (0.57 mmol, 95% defined by UV).

$^{1}$H NMR (500 MHz, $D_2O$) δ 7.70 (d, *J* = 8.1 Hz, 1H, H6 in U), 5.80 (d, *J* = 2.6 Hz, 1H, H1'), 5.77 (d, *J* = 8.1 Hz, 1H, H5 in U), 4.97 – 4.88 (ABXX' system appears as m, 2H, H2' and H3'), 4.44 – 4.38 (m, 1H, H4'), 4.08 – 4.02 (m, 2H, H5'), 1.48 (s, 3H, $CH_3$), 1.28 (s, 3H, $CH_3$), 1.19 – 1.16 (m, 6H, pC($CH_3$)$_2$p). $^{13}$C NMR (126 MHz,

$D_2O$) δ 166.35 (C4 in U), 151.34 (C2 in U), 142.66 (C6 in U), 114.57 (OCO in 2',3'-isopropylidene), 101.96 (C5 in U), 92.41 (C1'), 85.35 (d, $^3J_{PC}$ = 6.5 Hz, C4'), 84.16 (C2'), 80.59 (C3'), 64.53 (d, $^2J_{PC}$ = 6.2 Hz, C5'), 35.95 (dd, $^1J_{PC}$ = 125.6, 121.3 Hz, bridging C in $pC(CH_3)_2p$), 26.00 ($CH_3$ in 2',3'-isopropylidene), 24.30 ($CH_3$ in 2',3'-isopropylidene), 19.65 (ABXX' system appears as q, $J$ = 4.6 Hz, $CH_3$ in $pC(CH_3)_2p$). $^{31}P$ NMR (202 MHz, $D_2O$, pH 7.5) δ 27.47 (br, 1P), 24.32 (br, 1P). MS (ESI) m/z: $[M - H]^-$ calcd for $C_{15}H_{23}N_2O_{11}P_2^-$ 469.1; found 469.1. HRMS (ESI-TOF) m/z: $[M - H]^-$ calcd for $C_{15}H_{23}N_2O_{11}P_2^-$ 469.0777; found 469.0774.

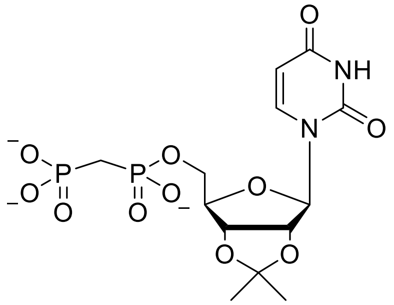

**[(4*R*,6*R*)-6-(2,4-dioxo-1,2,3,4-tetrahydropyrimidin-1-yl)-2,2-dimethyl-tetrahydro-2H-furo[3,4-d][1,3]dioxol-4-yl]methyl (phosphonomethyl)phosphonate, 2',3'-*O*-isopropylidene-$CH_2$-UBP 24e.** According to General Method 1, to well-dried tris(tetra-*n*-butylammonium) hydrogen (methylene)bisphosphonate **23e** (1.8 mmol, 3 equiv) was added **22** (265 mg, 0.6 mmol, 1 equiv) in 600 uL anhydrous acetonitrile. The solution was stirred for 2 d (~ 90% conversion as identified by $^{31}P$ NMR). The crude mixture was purified by dual-pass preparative RP-HPLC, Hamilton PRP-1 column (7 µm, 250 mm × 21 mm), isocratic mode, using 13% acetonitrile in 0.1 M triethylammonium bicarbonate, pH 7.5, at a flow rate of 8.0 mL/min and UV detection (261 nm). The title compound eluted at 9.8 min (see Figure S61). After removal of solvents under reduced pressure and co-evaporation with water (3 x 2 mL), the desired product **24e** was obtained as bis(triethylammonium) salts (0.5 mmol, 83% defined by UV).

$^1H$ NMR (400 MHz, $D_2O$) δ 7.77 (d, $J$ = 8.1 Hz, 1H, H6 in U), 5.81 (d, $J$ = 2.5 Hz, 1H, H1'), 5.79 (d, $J$ = 8.1 Hz, 1H, H5 in U), 4.96 – 4.90 (ABXX' system appears as m, 2H, H2' and H3'), 4.50 – 4.42 (m, 1H, H4'), 4.07 – 3.93 (m, 2H, H5'), 1.99 (t, $J$ = 19.5 Hz, 2H, $pCH_2p$), 1.49 (s, 3H, $CH_3$), 1.29 (s, 3H, $CH_3$). $^{31}P$ NMR (202 MHz, $D_2O$ pH 7.5) δ 21.95 (d, $J$ = 8.5 Hz, 1P), 11.30 (d, $J$ = 8.5 Hz, 1P). MS (ESI) m/z: $[M - H]^-$ calcd for $C_{13}H_{19}N_2O_{11}P_2^-$ 441.0; found 441.1.

**General Method 2:** A general procedure for 2',3'-*O*-isopropylidene removal to obtain the corresponding uridine-5'-(methylene)bisphosphonate derivatives (**1, 3, 6, 9,** and **10**).[4] Appropriate 2',3'-*O*-isopropylidene-uridine-5'-(methylene)bisphosphonate **24**(**a, b, c, e**) (0.57-0.45 mmol) was dissolved in 2 mL water and pH was adjusted to 0.2-0.5 with 1 N HCl solution and stirred at room temperature in a sealed flask.

The progress of the deprotection reaction was monitored by MS. After 2 h, the mixture is evaporated under reduced pressure (room temperature) and co-evaporated with water (2 x 2 mL) to remove the excess HCl followed by neutralizing the solution by 1 N $Na_2CO_3$. Then, volatiles were removed under reduced pressure, the residue was re-dissolved in water purified by RP-HPLC, Hamilton PRP-1 column (7 µm, 250 mm × 21 mm) with (4%-4.3%) acetonitrile in 0.1 M triethylammonium bicarbonate, pH 7.5, at a flow rate of 8.0 mL/min using an isocratic mode, with a UV detection (261 nm).

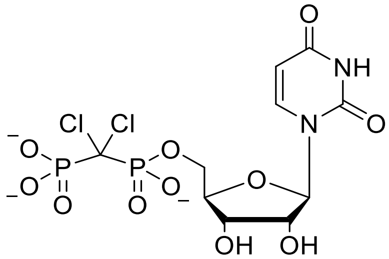

**[(2*R*,5*R*)-5-(2,4-dioxo-1,2,3,4-tetrahydropyrimidin-1-yl)-3,4-dihydroxyoxolan-2-yl]methyl [dichloro-(phosphono)methyl]phosphonate, $CCl_2$-UBP 1.** According to General Method 2, 0.49 mmol of **24a** was deprotected in acidic condition and purified by preparative RP-HPLC, Hamilton PRP-1 column (7 µm, 250 mm × 21 mm), isocratic mode, using 4.3% acetonitrile in 0.1 M triethylammonium bicarbonate, pH 7.5, at a flow rate of 8.0 mL/min and UV detection (261 nm). The title compound eluted at 9.3 min. After removal of solvents under reduced pressure and co-evaporation with water (3 x 2 mL), the desired product **1** was obtained as bis(triethylammonium) salts; colorless film (0.49 mmol, quant., defined by UV).

$^1H$ NMR (400 MHz, $D_2O$) δ 7.83 (d, $J$ = 8.1 Hz, 1H, H6 in U), 5.85 (d, $J$ = 3.2 Hz, 1H, H1'), 5.80 (d, $J$ = 8.1 Hz, 1H, H5 in U), 5.01 – 4.91 (ABXX' system appears as m, 2H, H2' and H3'), 4.46 – 4.42 (m, 1H, H4'), 4.25 – 4.20 (m, 2H, H5'), 1.49 (s, 3H, $CH_3$), 1.29 (s, 3H, $CH_3$). $^{31}P$ NMR (162 MHz, $D_2O$, pH 10) δ 9.90 (d, $J$ = 16.0 Hz, 1P), 7.69 (d, $J$ = 16.0 Hz, 1P). LRMS (ESI) m/z: $[M – H]^–$ calcd for $C_{10}H_{13}Cl_2N_2O_{11}P_2^-$ 468.9; found 469.0. HRMS (ESI-TOF) m/z: $[M – H]^–$ calcd for $C_{10}H_{13}Cl_2N_2O_{11}P_2^-$ 468.9372; found 468.9370.

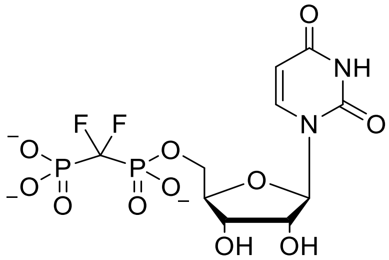

**[(2*R*,5*R*)-5-(2,4-dioxo-1,2,3,4-tetrahydropyrimidin-1-yl)-3,4-dihydroxyoxolan-2-yl]methyl [difluoro-(phosphono)methyl]phosphonate, $CF_2$-UBP 3.** According to General Method 2, 0.45 mmol of **24b** was deprotected in acidic condition and purified by preparative RP-HPLC, Hamilton PRP-1 column (7 µm, 250 mm × 21 mm), isocratic mode, using 4.3% acetonitrile in 0.1 M triethylammonium bicarbonate, pH 7.5, at a flow rate of 8.0 mL/min and UV detection (261 nm). The title compound eluted at 9.3 min. After removal of solvents under reduced pressure and co-evaporation with water (3 x 2 mL), the desired product **3** was obtained as bis(triethylammonium) salts; colorless film (0.45 mmol, quant., defined by UV).

$^{1}$H NMR (500 MHz, $D_2O$) δ 7.88 (d, *J* = 8.1 Hz, 1H, H6 in U), 5.88 (d, *J* = 4.5 Hz, 1H, H1'), 5.84 (d, *J* = 8.1 Hz, 1H, H5 in U), 4.28 – 4.24 (m, 2H, H2' and H3'), 4.22 – 4.14 (m, 3H, H4' and H5'). $^{19}$F NMR (470 MHz, $D_2O$, pH 7.5) δ -119.85 (ABXX' system appears as t, *J* = 83.1 Hz). $^{31}$P NMR (202 MHz, $D_2O$, pH 7.5) δ 5.34 – 1.55 (ABXX' system appears as m, 2P). LRMS (ESI) m/z: [M – H]$^-$ calcd for $C_{10}H_{13}F_2N_2O_{11}P_2^-$ 437.0; found 437.1. HRMS (ESI-TOF) m/z: [M – H]$^-$ calcd for $C_{10}H_{13}F_2N_2O_{11}P_2^-$ 436.9963; found 436.9962.

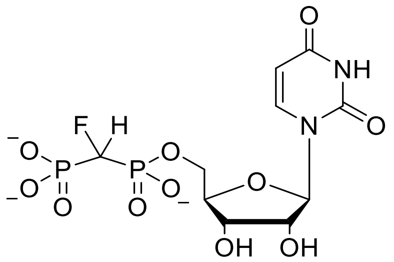

**[(2*R*,5*R*)-5-(2,4-dioxo-1,2,3,4-tetrahydropyrimidin-1-yl)-3,4-dihydroxyoxolan-2-yl]methyl [fluoro(phosphono)methyl]phosphonate, (*R*/*S*)-CHF-UBP 6.** According to General Method 2, 0.47 mmol of **24c** was deprotected in acidic condition and purified by preparative RP-HPLC, Hamilton PRP-1 column (7 µm, 250 mm × 21 mm), isocratic mode, using 4.2% acetonitrile in 0.1 M triethylammonium bicarbonate, pH 7.5, at a flow rate of 8.0 mL/min and UV detection (261 nm). The title compound eluted at 9.2 min. After removal of solvents under reduced pressure and co-evaporation with water (3 x 2 mL), the desired product **6** was obtained as bis(triethylammonium) salts; colorless film (0.47 mmol, quant., defined by UV).

$^{1}$H NMR (500 MHz, $D_2O$) mixture of two diastereomers (1:1), δ 7.84 (d, *J* = 8.1 Hz, 1H, H6 in U), 5.85 (d, *J* = 4.4 Hz, 1H, H1'), 5.81 (appears as dd, *J* = 8.1, 1.1 Hz, 1H, H5 in U), 4.90 and 4.81 (two (*R*/*S*)-pCHFp chemical shifts, each appears as td, *J* = 12.7, 4.1 Hz, 1H), 4.26 – 4.21 (m, 2H, H2' and H3'), 4.20 – 4.07 (m, 3H, H4' and H5'). $^{19}$F NMR (470 MHz, $D_2O$, pH 7.5) mixture of two diastereomers (1:1), δ -217.10 (ddd, *J* = 63.1, 56.2, 45.6 Hz, 1F, isomer 1), -217.57 (ddd, *J* = 62.7, 55.9, 45.4 Hz, 1F, isomer 2). $^{31}$P NMR (202 MHz, $D_2O$, pH 7.5) mixture of two diastereomers (1:1), δ 13.62 (appears as dd, *J* = 63.6, 11.7 Hz, 1P), 6.83 (appear as dd, *J* = 56.7, 11.5 Hz, 1P). LRMS (ESI) m/z: [M – H]$^-$ calcd for $C_{10}H_{14}FN_2O_{11}P_2^-$ 419.0; found 419.2. HRMS (ESI-TOF) m/z: [M – H]$^-$ calcd for $C_{10}H_{14}FN_2O_{11}P_2^-$ 419.0057; found 419.0052.

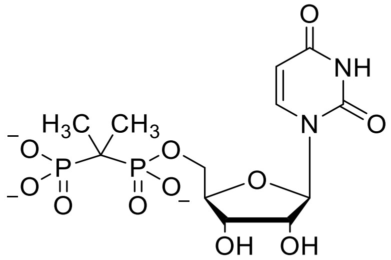

**[(2*R*,5*R*)-5-(2,4-dioxo-1,2,3,4-tetrahydropyrimidin-1-yl)-3,4-dihydroxyoxolan-2-yl]methyl (2-phosphonopropan-2-yl)phosphonate, $C(CH_3)_2$-UBP 9.** According to General Method 2, 0.57 mmol of **24d** was deprotected in acidic condition and purified by preparative RP-HPLC, Hamilton PRP-1 column (7 µm, 250 mm × 21 mm), isocratic mode, using 4.2% acetonitrile in 0.1 M triethylammonium bicarbonate, pH 7.5, at a flow rate of 8.0 mL/min and UV detection (260 nm). The title compound eluted at 9.3 min. After removal of solvents under reduced pressure and co-evaporation with water (3 x 2 mL), the desired product **9** was obtained as bis(triethylammonium) salts; colorless film (0.57 mmol, quant., defined by UV).

$^1$H NMR (400 MHz, $D_2O$) δ 7.88 (d, *J* = 8.1 Hz, 1H, H6 in U), 5.82 (d, *J* = 4.8 Hz, 1H, H1'), 5.79 (d, *J* = 8.1 Hz, 1H, H5 in U), 4.28 – 4.21 (m, 2H, H2' and H3'), 4.12 – 4.06 (m, 3H, H4' and H5'), 1.33 – 1.17 (m, 6H, $pC(CH_3)_2p$). $^{31}$P NMR (202 MHz, $D_2O$, pH 7.5) δ 27.34 (br, 1P), 24.65 (br, 1P). $^{13}$C NMR (126 MHz, $D_2O$) δ 166.13 (C4 in U), 151.72 (C2 in U), 141.96 (C6 in U), 102.46 (C5 in U), 88.31 (C1'), 83.53 (d, $^3J_{PC}$ = 6.8 Hz, C4'), 73.64 (C2'), 69.59 (C3'), 64.05 (d, $^2J_{PC}$ = 6.1 Hz, C5'), 35.93 (dd, $^1J_{PC}$ = 125.7, 122.0 Hz, bridging C in $pC(CH_3)_2p$), 19.64 (ABXX' system appear as q, 2 x $CH_3$ in $pC(CH_3)_2p$). LRMS (ESI) m/z: [M – H]$^-$ calcd for $C_{12}H_{19}N_2O_{11}P_2^-$ 429.0; found 429.1. HRMS (ESI-TOF) m/z: [M – H]$^-$ calcd for $C_{12}H_{19}N_2O_{11}P_2^-$ 429.0464; found 429.0456.

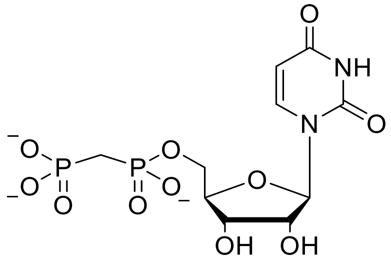

**[(2*R*,5*R*)-5-(2,4-dioxo-1,2,3,4-tetrahydropyrimidin-1-yl)-3,4-dihydroxyoxolan-2-yl]methyl (phosphonomethyl)phosphonate, $CH_2$-UBP 10.** According to General Method 2, 0.5 mmol of **24e** was deprotected in acidic condition and purified by preparative RP-HPLC, Hamilton PRP-1 column (7 µm, 250 mm × 21 mm), isocratic mode, using 4% acetonitrile in 0.1 M triethylammonium bicarbonate, pH 7.5, at a flow rate of 8.0 mL/min and UV detection (261 nm). The title compound eluted at 8.6 min. After removal of solvents under reduced pressure and co-evaporation with water (3 x 2 mL), the desired product **10** was obtained as bis(triethylammonium) salts; colorless film (0.5 mmol, quant., defined by UV).

$^1H$ NMR (500 MHz, $D_2O$) δ 7.87 (d, $J$ = 8.0 Hz, 1H, H6 in U), 5.83 (d, $J$ = 4.2 Hz, 1H, H1'), 5.81 (d, $J$ = 8.0 Hz, 1H, H5 in U), 4.28 – 4.22 (m, 2H, H2' and H3'), 4.13 (br, 1H, H4'), 4.08 – 3.97 (m, 2H, H5'), 2.05 (t, $J$ = 19.7 Hz, 2H, $pCH_2p$). $^{31}P$ NMR (162 MHz, $D_2O$, pH 11.7) δ 22.23 (d, $J$ = 9.4 Hz, 1P), 10.97 (d, $J$ = 9.4 Hz, 1P). $^{13}C$ NMR (126 MHz, $D_2O$) δ 166.15 (C4 in U), 151.72 (C2 in U), 141.79 (C6 in U), 102.47 (C5 in U), 88.37 (C1'), 83.33 (d, $^3J_{PC}$ = 7.8 Hz, C4'), 73.67 (C2'), 69.55 (C3'), 63.22 (d, $^2J_{PC}$ = 4.8 Hz, C5'), 27.37 (t, $^1J_{PC}$ = 124.3 Hz, $pCH_2p$). LRMS (ESI) m/z: $[M-H]^-$ calcd for $C_{10}H_{15}N_2O_{11}P_2^-$ 401.0; found 401.1. HRMS (ESI-TOF) m/z: $[M-H]^-$ calcd for $C_{10}H_{15}N_2O_{11}P_2^-$ 401.0151; found 401.0147.

#### Synthesis of GlcNAc-CXY-UBP analogues (CXY = $CH_2$, $CF_2$, $CCl_2$, (*R*)-CHF, (*S*)-CHF)

**General Method 3:** A general procedure for coupling **13** with **1**, **3**, and **6** to obtain the corresponding α-GlcNAc$(OAc)_3$-CXY-UBP, **25a-c**. The triacids products of appropriate CXY-UBP **1**, **3**, or **6** (0.05 mmol) were obtained by passage through a pipette column of DOWEX $H^+$ using water as eluent. The solvent was removed under reduced pressure, co-evaporated by anhydrous DMF (3 × 1 mL) and left under reduced pressure until the total weight of the flask remained constant. Then, well-dried oxazoline **13** (0.1 mmol, 2 equiv) in 1 mL anhydrous DMF was added to the corresponding free acid CXY-UBP and stirred under $N_2$ gas at 45 °C for 2-3 d and the reaction was monitored by MS. (Note: Higher temperature results β-anomer in anomeric center C1"). Each molecule of water in the reaction mixture deactivates two oxazoline intermediates **13** causing the formation of a glycosidic bond (disaccharide) which was detected by MS. In this case, more oxazoline **13** can be added to move the reaction forward. After completion, the crude material was concentrated under reduced pressure and purified by dual-pass preparative HPLC: (1) Macherey-Nagel Nucleogel SAX 1000−10 (25 mm × 15 cm) column, using gradient mode (A/ 10% acetonitrile in $H_2O$ and B/ 10% acetonitrile in 0.5 M triethylammonium bicarbonate, pH 7.5; 0−7 min, 100% A; 7−10 min, 0%-25% B; 10−12.5 min, 25% B; 12.5−16 min, 25%-35% B; 16−20 min, 35% B; 20−25 min, 35%-75% B; see Figure S109) at a flow rate of 8 mL/min with a UV detection (261 nm); (2) Hamilton PRP-1 column (7 µm, 250 mm × 21 mm) with 14% acetonitrile in 0.1 M triethylammonium bicarbonate, pH 7.5, at a flow rate of 8.0 mL/min using an isocratic mode with a UV detection (254/260 nm) to give desired product **25a-c** as bis(triethylammonium) salts.

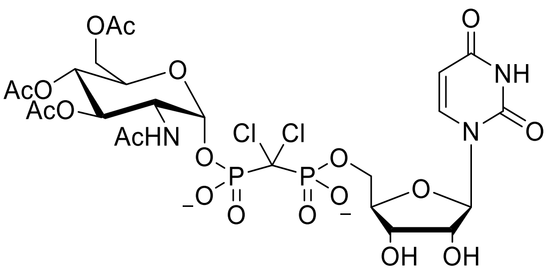

**[(2*R*,5*R*)-5-(2,4-dioxo-1,2,3,4-tetrahydropyrimidin-1-yl)-3,4-dihydroxyoxolan-2-yl]methyl {[(2*R*,5*S*)-4,5-bis(acetyloxy)-6-[(acetyloxy)methyl]-3-acetamidooxan-2-yl phosphono]dichloromethyl}phosphonate, α-GlcNAc(OAc)$_3$-CCl$_2$-UBP, 25a.** According to General Method 3, 0.05 mmol of **1** was converted to free acid form and coupled with **13** (0.1 mmol, 2 equiv) at 45 °C for 2 d. After dual-pass preparative HPLC purification (SAX, RP-HPLC), the title compound **25a** was obtained as bis(triethylammonium) salts; colorless film (pure α-anomer, 0.031 mmol, 63%, defined by UV).

SAX-HPLC purification retention time = 16.6 min (see Figure S104 for detail gradient condition); RP-HPLC purification retention time = 12.9 min. $^1$H NMR (500 MHz, D$_2$O) δ 7.88 (d, *J* = 7.5 Hz, 1H, H6 in U), 5.91 (d, *J* = 4.5 Hz, 1H, H1' in U), 5.81 (br, 1H, H5 in U), 5.60 – 5.56 (m, 1H, H1" in Glc), 5.19 (t, *J* = 9.9 Hz, 1H, H3" in Glc), 5.02 (t, *J* = 9.9 Hz, 1H, H4" in Glc), 4.37 – 4.23 (m, 7H, protons in both Glc and U), 4.17 – 4.15 (m, 1H, H6" in Glc), 4.10 – 4.08 (m, 1H, H6" in Glc), 2.02 (s, 3H, Ac), 1.97 (s, 3H, Ac), 1.92 (s, 6H, Ac). $^{31}$P NMR (202 MHz, D$_2$O pH 7.5) δ 7.59 (d, *J* = 15.5 Hz, 1P), 6.08 (d, *J* = 15.5 Hz, 1P). MS (ESI) m/z: [M – H]$^-$ calcd for $C_{24}H_{32}Cl_2N_3O_{19}P_2^-$ 798.0; found 798.1.

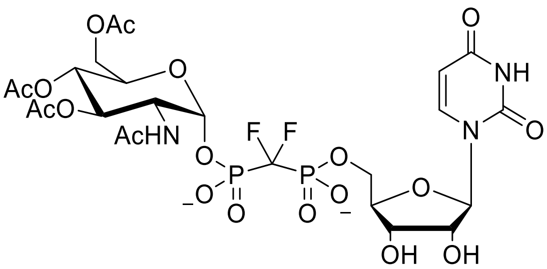

**[(2*R*,5*R*)-5-(2,4-dioxo-1,2,3,4-tetrahydropyrimidin-1-yl)-3,4-dihydroxyoxolan-2-yl]methyl {[(2*R*,5*S*)-4,5-bis(acetyloxy)-6-[(acetyloxy)methyl]-3-acetamidooxan-2-yl phosphono]difluoromethyl}phosphonate, α-GlcNAc(OAc)$_3$-CF$_2$-UBP, 25b-α.** According to General Method 3, 0.05 mmol of **3** was converted to the free acid form and coupled with **13** (0.1 mmol, 2 equiv) at 45 °C for 3 d. After dual-pass preparative HPLC purification (SAX, RP-HPLC), the title compound **25b-α** was obtained as bis(triethylammonium) salts; colorless film (pure α-anomer, 0.024 mmol, 49%, defined by UV).

SAX-HPLC purification retention time = 17.9 min (see Figure S109); RP-HPLC purification retention time = 13.3 min. $^1$H NMR (500 MHz, D$_2$O) δ 7.73 (d, *J* = 7.7 Hz, 1H, H6 in U), 5.91 (d, *J* = 5.1 Hz, 1H, H1' in U), 5.75

(d, $J$ = 7.6 Hz, 1H, H5 in U), 5.52 (dd, $J$ = 6.6, 3.5 Hz, 1H, H1" in Glc), 5.20 (dd, $J$ = 10.4, 9.6 Hz, 1H, H3" in Glc), 5.01 (t, $J$ = 9.8 Hz, 1H, H4" in Glc), 4.35 – 4.19 (m, 7H, protons in both Glc and U), 4.16 – 4.13 (m, 1H, H6" in Glc), 4.09 – 4.04 (m, 1H, H6" in Glc), 1.96 (s, 3H, Ac), 1.95 (s, 3H, Ac), 1.92 (s, 3H, Ac) , 1.90 (s, 3H, Ac). $^{31}$P NMR (202 MHz, $D_2O$ pH 7.5) δ 4.31 – 1.46 (ABXX' system appears as m, 2P). $^{19}$F NMR (470 MHz, $D_2O$, pH 7.5) δ -120.01 (ABXX' system appears as ddd, $J$ = 86.9, 81.5, 20.1 Hz). MS (ESI) m/z: [M – H]$^-$ calcd for $C_{24}H_{32}F_2N_3O_{19}P_2^-$ 766.1; found 766.2.

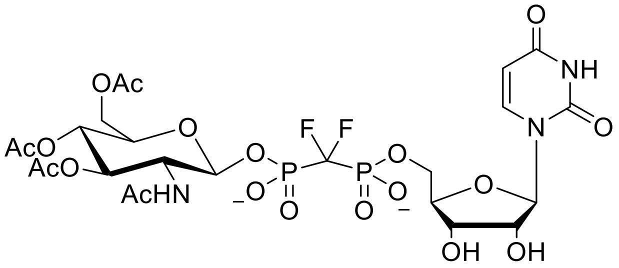

**[(2*R*,5*R*)-5-(2,4-dioxo-1,2,3,4-tetrahydropyrimidin-1-yl)-3,4-dihydroxyoxolan-2-yl]methyl {[(2*S*,5*S*)-4,5-bis(acetyloxy)-6-[(acetyloxy)methyl]-3-acetamidooxan-2-yl phosphono]difluoromethyl}phosphonate, β-GlcNAc(OAc)$_3$-CF$_2$-UBP, 25b-β.** To obtain the β-anomer of the title compound, the coupling reaction was performed at the higher temperature. According to General Method 3, 0.05 mmol of **3** was converted to the free acid form, dissolved in 0.5 mL anhydrous DMF and the solution stirred at 75 °C for 5 min. To this solution, **13** (0.1 mmol, 2 equiv) in 0.5 mL anhydrous DMF was added dropwise and the mixture was left at 75 °C for 3 h. After dual-pass preparative HPLC purification (SAX-, RP-HPLC), the title compound 25b-β was obtained as bis(triethylammonium) salts; colorless film (pure β-anomer, 0.038 mmol, 75%, defined by UV). (Note: To elaborate separation of α-anomer and β-anomer, a mixture of both epimers were separated by Hamilton PRP-1 column (7 µm, 250 mm × 21 mm) under General Method 3 conditions (see **Figure S120**).

SAX-HPLC purification retention time = 17.9 min; RP-HPLC purification retention time = 14.3 min. $^1$H NMR (500 MHz, $D_2O$) δ 7.89 (d, $J$ = 8.1 Hz, 1H, H6 in U), 5.88 (d, $J$ = 4.4 Hz, 1H, H1' in U), 5.84 (d, $J$ = 8.1 Hz, 1H, H5 in U), 5.22 (d, $J$ = 8.1 Hz, 1H H1" in Glc), 5.18 (dd, $J$ = 10.5, 9.3 Hz, 1H, H3" in Glc), δ 4.98 (t, $J$ = 9.7 Hz, 1H, H4" in Glc). 4.33 (dd, $J$ = 12.7, 3.6 Hz, 1H, H6" in Glc), 4.29 – 4.12 (m, 6H, protons in U), 4.09 (dd, $J$ = 12.7, 2.2 Hz, 1H, H6" in Glc), 4.00 – 3.91 (m, 2H, H2" and H5" in Glc), 2.01 (s, 3H, Ac), 1.97 (s, 3H, Ac), 1.94 (s, 3H, Ac), 1.86 (s, 3H, Ac). $^{31}$P NMR (202 MHz, $D_2O$, pH 7.5) δ 2.53 (ABXX' system appears as dtd $J$ = 286.4, 82.2, 59.9 Hz, 2P). $^{19}$F NMR (470 MHz, $D_2O$, pH 7.5) δ -119.52 (ABXX' system appears as td, $J$ = 82.1, 36.2 Hz). MS (ESI) m/z: [M – H]$^-$ calcd for $C_{24}H_{32}F_2N_3O_{19}P_2^-$ 766.1; found 766.2.

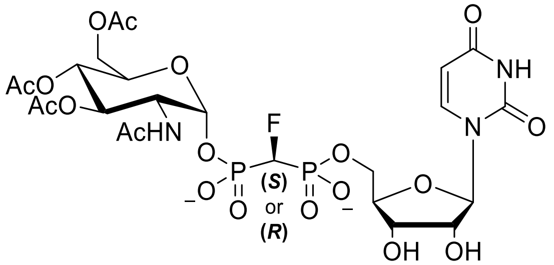

**[(2*R*,5*R*)-5-(2,4-dioxo-1,2,3,4-tetrahydropyrimidin-1-yl)-3,4-dihydroxyoxolan-2-yl]methyl [(*S*)-[(2*R*,5*S*)-4,5-bis(acetyloxy)-6-[(acetyloxy)methyl]-3-acetamidooxan-2-yl phosphono](fluoro)methyl]phosphonate and its (*R*) isomer, (*S*) or (*R*) α-GlcNAc(OAc)$_3$-CHF-UBP, 25c-α1/25c-α2.** According to General Method 3 (with modified RP-HPLC purification condition), 0.05 mmol of **6** was converted to the free acid form and coupled with **13** (0.1 mmol, 2 equiv) at 45 °C for 2 d. After preparative SAX-HPLC purification (retention time = 17.5 min), the resulting 1:1 mixture 25c-α1/25c-α2 was separated by preparative RP-HPLC (see **Figure S123**), Hamilton PRP-1 column (7 µm, 250 mm × 21 mm) with 8.3% acetonitrile in 0.24 M ammonium acetate ($NH_4OAc$), pH 6.5, at a flow rate of 8.0 mL/min using an isocratic mode with a UV detection (261 nm) to obtain title compounds **25c-α1** (fast eluting isomer, (*S*)-CHF) and **25c-α2** (slow eluting isomer, (*R*)-CHF) as ammonium salts; colorless crystals.

**25c-α1** (((*S*)-CHF, fast isomer, pure α-anomer, 0.017 mmol, 70%, defined by UV): RP-HPLC purification retention time = 16.2 min. $^{19}$F NMR (376 MHz, $D_2O$, pH 7.5) δ -221.88 (td, $^2J_{PF}$ = 61.7 Hz, $^2J_{HF}$ = 45.3 Hz, 1F). $^{31}$P NMR (202 MHz, $D_2O$, pH 7.5) δ 9.58 (dd, $^2J_{PF}$ = 61.7 Hz, $^2J_{PP}$ = 11.5 Hz, 1P), 8.66 (dd, $^2J_{PF}$ = 61.7 Hz, $^2J_{PP}$ = 11.5 Hz, 1P). MS (ESI) m/z: [M – H]$^-$ calcd for $C_{24}H_{33}FN_3O_{19}P_2^-$ 748.1; found 748.1.

**25c-α2** ((*R*)-CHF, slow isomer, pure α-anomer, 0.016 mmol, 66%, defined by UV): RP-HPLC purification retention time = 18.0 min. $^{19}$F NMR (470 MHz, $D_2O$, pH 7.5) δ -220.50 (td, $^2J_{PF}$ = 61.1 Hz, $^2J_{HF}$ = 45.8 Hz, 1F). $^{31}$P NMR (202 MHz, $D_2O$, pH 7.5) δ 9.35 (dd, $^2J_{PF}$ = 61.1 Hz, $^2J_{PP}$ = 12.2 Hz, 1P), 8.60 (dd, $^2J_{PF}$ = 61.1 Hz, $^2J_{PP}$ = 12.2 Hz, 1P). MS (ESI) m/z: [M – H]$^-$ calcd for $C_{24}H_{33}FN_3O_{19}P_2^-$ 748.1; found 748.1.

**General Method 4:** A general procedure for *O*-acetyl removal to obtain the final corresponding GlcNAc-CXY-UBP (**2, 4, 5, 7,** and **8**). Compound **25a-c** were dissolved in 2 mL water and pH was adjusted to 11.5 by adding $NH_4OH$ solution. (Note: To avoid racemization on the bridging pCHFp, for **25c-α1** and **25c-α2** pH was adjusted to 10.5). The solution was stirred at room temperature for 2-3 h in a sealed flask. The progress of the reaction was monitored by MS. Upon completion, the solvent was removed the under reduced pressure, the residue was re-dissolved in water and purified by preparative RP-HPLC, Hamilton PRP-1 column (7 µm, 250 mm × 21 mm), isocratic mode, using (4%) acetonitrile in 0.1 M triethylammonium bicarbonate, pH 7.5, at a flow rate of 8.0 mL/min and UV detection (261 nm) to yield desired product (2, 4, 5, 7, and 8) as bis(triethylammonium) salts.

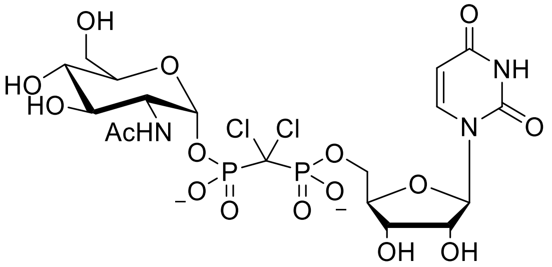

**[(2*R*,5*R*)-5-(2,4-dioxo-1,2,3,4-tetrahydropyrimidin-1-yl)-3,4-dihydroxyoxolan-2-yl]methyl {dichloro-[(2*R*,5*S*)-3-acetamido-4,5-dihydroxy-6-(hydroxymethyl)oxan-2-yl phosphono]methyl}phosphonate, α-GlcNAc-$CCl_2$-UBP 2.** According to General Method 4, 0.02 mmol of **25a** was deprotected under basic conditions (pH 11.5) and purified by preparative RP-HPLC to obtain **2** as bis(triethylammonium) salts; colorless film (pure α-anomer, 0.02 mmol, quant., defined by UV).

RP-HPLC purification retention time = 12.2 min. $^1$H NMR (400 MHz, $D_2O$) δ 7.93 (d, *J* = 8.1 Hz, 1H, H6 in U), δ 5.88 (d, *J* = 4.6 Hz, 1H, H1′ in U), 5.82 (d, *J* = 8.1 Hz, 1H, H5 in U), 5.48 (dd, $^3J_{PH}$ = 5.9 Hz, $^3J_{HH}$ = 3.4 Hz, 1H, H1″ in Glc), 4.32 – 4.20 (m, 4H, H2′, H3′, H4′, and H5′ in U), 4.19 – 4.12 (m, 1H, H5′ in U), 3.89 (dt, *J* = 10.4, 3.0 Hz, 1H, H2″ in Glc), 3.86 – 3.80 (m, 1H, H5″ in Glc), 3.80 – 3.64 (m, 3H, H3″ and H6″ in Glc), 3.40 (t, *J* = 9.6 Hz, 1H, H4″ in Glc), 1.96 (s, 3H, Ac). $^{31}$P NMR (162 MHz, $D_2O$, pH 7.5) δ 7.48 (d, $^2J_{PP}$ = 14.6 Hz, 1P), 5.95 (d, $^2J_{PP}$ = 14.6 Hz, 1P). LRMS (ESI) m/z: [M – H]$^-$ calcd for $C_{18}H_{26}Cl_2N_3O_{16}P_2^-$ 672.0; found 672.0. HRMS (ESI-TOF) m/z: [M – H]$^-$ calcd for $C_{18}H_{26}Cl_2N_3O_{16}P_2^-$ 672.0165; found 672.0170.

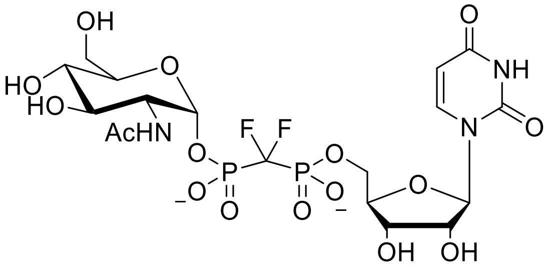

**[(2*R*,5*R*)-5-(2,4-dioxo-1,2,3,4-tetrahydropyrimidin-1-yl)-3,4-dihydroxyoxolan-2-yl]methyl {[(2*R*,5*S*)-3-acetamido-4,5-dihydroxy-6-(hydroxymethyl)oxan-2-yl phosphono]difluoromethyl}phosphonate, α-Glc-NAc-$CF_2$-UBP 4.** According to General Method 4, 0.02 mmol of **25b-α** was deprotected under basic conditions (pH 11.5) and purified by preparative RP-HPLC to obtain **4** as bis(triethylammonium) salts; colorless film (pure α-anomer, 0.02 mmol, quant., defined by UV).

RP-HPLC purification retention time = 10.0 min. $^1$H NMR (400 MHz, $D_2O$) δ 7.76 (d, *J* = 7.9 Hz, 1H, H6 in U), δ 5.88 (d, *J* = 5.1 Hz, 1H, H1′ in U), 5.77 (d, *J* = 7.9 Hz, 1H, H5 in U), 5.40 (dd, $^3J_{PH}$ = 6.5 Hz, $^3J_{HH}$ = 3.3 Hz, 1H, H1″ in Glc), 4.28 – 4.07 (m, 5H, H2′, H3′, H4′, and H5′ in U), 3.86 (dt, *J* = 10.5, 2.8 Hz, 1H, H2″ in Glc), 3.81 (ddd, *J* = 10.1, 4.7, 2.3 Hz, 1H, H5″ in Glc), 3.77 – 3.64 (m, 3H, H3″ and H6″ in Glc), 3.40 (dd, *J* = 10.1, 9.1

Hz, 1H, H4" in Glc), 1.93 (s, 3H, Ac). $^{19}$F NMR (470 MHz, $D_2O$, pH 7.5) δ -119.26 – -121.50 (ABXX' system appears as m). $^{31}$P NMR (202 MHz, $D_2O$, pH 7.5) δ 3.80 – 1.78 (ABXX' system appears as m). LRMS (ESI) m/z: $[M-H]^-$ calcd for $C_{18}H_{26}F_2N_3O_{16}P_2^-$ 640.1; found 640.2. HRMS (ESI-TOF) m/z: $[M-H]^-$ calcd for $C_{18}H_{26}F_2N_3O_{16}P_2^-$ 640.0756; found 640. 0756.

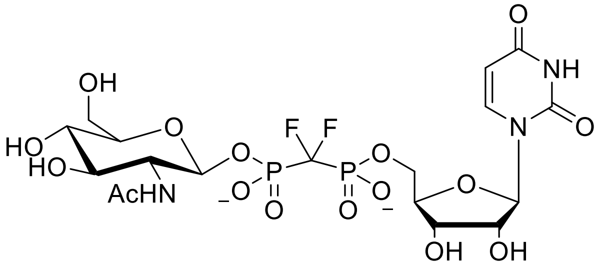

**[(2*R*,5*R*)-5-(2,4-dioxo-1,2,3,4-tetrahydropyrimidin-1-yl)-3,4-dihydroxyoxolan-2-yl]methyl {[(2*S*,5*S*)-3-acetamido-4,5-dihydroxy-6-(hydroxymethyl)oxan-2-yl phosphono]difluoromethyl}phosphonate, β-Glc-NAc-$CF_2$-UBP 5.** According to General Method 4, 0.01 mmol of **25b-β** was deprotected under basic conditions (pH 11.5) and purified by preparative RP-HPLC to obtain **5** as bis(triethylammonium) salts; colorless film (pure β-anomer, 0.01 mmol, quant., defined by UV).

RP-HPLC purification retention time = 9.2 min. $^1$H NMR (400 MHz, $D_2O$) δ 7.72 (d, $J$ = 7.7 Hz, 1H, H6 in U), δ 5.90 (d, $J$ = 5.2 Hz, 1H, H1' in U), 5.75 (d, $J$ = 6.9 Hz, 1H, H5 in U), 5.01 (dd, $^3J_{HH}$ = 8.3 Hz, $^3J_{PH}$ = 7.6 Hz, 1H, H1" in Glc), 4.26 – 4.08 (m, 5H, H2', H3', H4', and H5' in U), 3.85 – 3.57 (m, 3H, H2", H3" and H4" in Glc), 3.51 – 3.29 (m, 3H, H5" and H6" in Glc), 1.89 (s, 3H, Ac). $^{19}$F NMR (470 MHz, $D_2O$, pH 7.5) δ -119.50 (ABXX' system appears as t, $J$ = 81.1 Hz, 2F). $^{31}$P NMR (202 MHz, $D_2O$, pH 7.5) δ 3.76 – 1.50 (ABXX' system appears as m). LRMS (ESI) m/z: $[M-H]^-$ calcd for $C_{18}H_{26}F_2N_3O_{16}P_2^-$ 640.1; found 640.2. HRMS (ESI-TOF) m/z: $[M-H]^-$ calcd for $C_{18}H_{26}F_2N_3O_{16}P_2^-$ 640.0756; found 640. 0764.

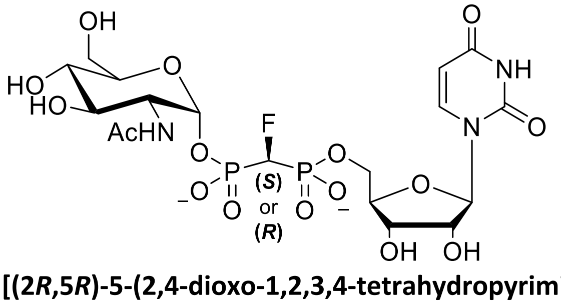

**[(2*R*,5*R*)-5-(2,4-dioxo-1,2,3,4-tetrahydropyrimidin-1-yl)-3,4-dihydroxyoxolan-2-yl]methyl [(*S*)-[(2*R*,5*S*)-3-acetamido-4,5-dihydroxy-6-(hydroxymethyl)oxan-2-yl phosphono](fluoro)methyl]phosphonate and its (*R*) isomer, (*S*)- or (*R*)-α-GlcNAc-CHF-UBP 7-α1/8-α2.** According to General Method 4, 0.01 mmol of individual **7-α1** and **8-α2** were deprotected under basic conditions (pH 10.5; higher pH causes racemization at the CHF bridging carbon) and purified by preparative RP-HPLC to obtain **7-α1** and **8-α2** as bis(triethylammonium) salts; colorless film.

**7-α1** ((*S*)-CHF, fast isomer, pure α-anomer, 0.01 mmol, quant., defined by UV): RP-HPLC purification retention time = 10.0 min. $^1$H NMR (400 MHz, $D_2O$) δ 7.67 (d, *J* = 7.7 Hz, 1H, H6 in U), δ 5.90 (d, *J* = 5.0 Hz, 1H, H1′ in U), 5.71 (d, *J* = 7.7 Hz, 1H, H5 in U), 5.38 (dd, $^3J_{PH}$ = 7.3 Hz, $^3J_{HH}$ = 3.3 Hz, 1H, H1″ in Glc), 4.84 (dt, $^2J_{HF}$ = 45.5 Hz, $^2J_{HP}$ = 12.1 Hz, 1H, pCHFp), 4.24 – 4.17 (m, 2H, H2′, and H3′ in U), 4.13 – 4.07 (m, 3H, H4′, and H5′ in U), 3.89 – 3.81 (m, 2H, H2″, and H5″ in Glc), 3.77 – 3.64 (m, 3H, H3″ and H6″ in Glc), 3.39 (dd, *J* = 10.1, 9.1 Hz, 1H, H4″ in Glc), 1.93 (s, 3H, Ac). $^{19}$F NMR (376 MHz, $D_2O$, pH 7.5) δ -221.89 (td, $^2J_{PF}$ = 61.8 Hz, $^2J_{HF}$ = 45.5 Hz). $^{31}$P NMR (202 MHz, $D_2O$, pH 7.5) δ 9.62 (dd, $^2J_{PF}$ = 61.8 Hz, $^2J_{PP}$ = 11.1 Hz), 8.75 (dd, $^2J_{PF}$ = 61.8 Hz, $^2J_{PP}$ = 11.1 Hz). LRMS (ESI) m/z: [M – H]$^-$ calcd for $C_{18}H_{27}FN_3O_{16}P_2^-$ 622.1; found 622.1. HRMS (ESI-TOF) m/z: [M – H]$^-$ calcd for $C_{18}H_{27}FN_3O_{16}P_2^-$ 622.0851; found 622.0851.

**8-α2** ((*R*)-CHF, slow isomer, pure α-anomer, 0.01 mmol, quant., defined by UV): RP-HPLC purification retention time = 9.8 min. $^1$H NMR (400 MHz, $D_2O$) δ 7.84 (d, *J* = 8.0 Hz, 1H, H6 in U), δ 5.86 (d, *J* = 3.8 Hz, 1H, H1′ in U), 5.81 (d, *J* = 8.0 Hz, 1H, H5 in U), 5.41 (dd, $^3J_{PH}$ = 7.2 Hz, $^3J_{HH}$ = 3.4 Hz, 1H, H1″ in Glc), 4.78 (dt, $^2J_{HF}$ = 45.5 Hz, $^2J_{HP}$ = 12.6 Hz, 1H, pCHFp), 4.29 – 4.21 (m, 2H, H2′, and H3′ in U), 4.17 – 4.08 (m, 3H, H4′, and H5′ in U), 3.88 – 3.79 (m, 2H, H2″, and H5″ in Glc), 3.77 – 3.63 (m, 3H, H3″ and H6″ in Glc), 3.40 (dd, *J* = 10.1, 9.1 Hz, 1H, H4″ in Glc), 1.94 (s, 3H, Ac). $^{19}$F NMR (470 MHz, $D_2O$, pH 7.5) δ -220.53 (td, $^2J_{PF}$ = 61.2 Hz, $^2J_{HF}$ = 45.5 Hz, 1F). $^{31}$P NMR (202 MHz, $D_2O$, pH 7.5) δ 9.63 (dd, $^2J_{PF}$ = 61.2 Hz, $^2J_{PP}$ = 12.5 Hz, 1P), 8.62 (dd, $^2J_{PF}$ = 61.2 Hz, $^2J_{PP}$ = 12.5 Hz, 1P). LRMS (ESI) m/z: [M – H]$^-$ calcd for $C_{18}H_{27}FN_3O_{16}P_2^-$ 622.1; found 622.1. HRMS (ESI-TOF) m/z: [M – H]$^-$ calcd for $C_{18}H_{27}FN_3O_{16}P_2^-$ 622.0851; found 622.0850.

Synthesis for the stereochemistry elucidation of **7** and **8**

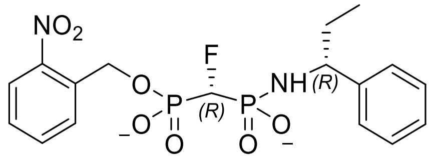

**[(*R*)-Fluoro[hydroxy({[(1*R*)-1-phenylpropyl]amino})phosphoryl]-methyl][(2-nitrophenyl)methoxy]phosphinic acid 26**. This compound was synthesized according to the literature with slight modifications.[4] (Fluoro(hydroxy((2-nitrobenzyl)oxy)phosphoryl)methyl)-phosphonic acid as bis(triethylammonium) salt (493 mg, 0.78 mmol, 1.0 eq) was dissolved in DMSO (2.6 mL). (*R*)-1-Phenyl- propan-1-amine (606 mg, 4.49 mmol, 5.7 eq), triphenylphosphine (390 mg, 1.49 mmol, 2.0 eq), and dithiopyridine (327 mg, 1.49 mmol, 2.0 eq) were added to the reaction mixture stirred at rt for 3h. The reaction was quenched with water. The solvents were removed under reduced pressure. The (*R*)-CHF-(*R*)-auxiliary isomer was isolated by dual pass RP-HPLC, Phenomenex Luna C18(2) HPLC column (5 µm, 250 mm × 10 mm), isocratic method, using 25% acetonitrile in 0.25 M ammonium acetate, pH 6.5, at a flow rate of 8.0 mL/min and UV detection (280 nm) (see **Figure S169**). After removal of solvents under reduced pressure and co-evaporation with water (3 x 2 mL), the desired product **26** was obtained as an ammonium salt; white solid (80 mg, 0.18 mmol, 46% assuming 1:1 isomer ratio).

The fast diastereomer,(*R*)-CHF-(*R*)-auxiliary26 was eluted at 14.6 min: $^1$H NMR (500 MHz, $CD_3OD$) δ 8.09 (dd, *J* = 8.3, 1.1 Hz, 1H), 7.98 (d, *J* = 7.9 Hz, 1H), 7.74 – 7.67 (m, 1H), 7.49 (t, *J* = 7.8 Hz, 1H), 7.35 (d, *J* = 7.0 Hz, 2H), 7.24 (t, *J* = 7.5 Hz, 2H), 7.11 (td, *J* = 7.4, 1.3 Hz, 1H), 5.33 (qd, *J* = 15.9, 7.2 Hz, 2H), 4.51 (dt, *J* = 46.8, 11.4 Hz, 1H), 4.22 (q, *J* = 7.3 Hz, 1H), 1.86 – 1.76 (m, 1H), 1.74 – 1.65 (m, 1H), 0.83 (t, *J* = 7.3 Hz, 3H). $^{19}$F NMR (470 MHz, $CD_3OD$) δ -219.23 (q, *J* = 56.8 Hz). $^{31}$P NMR (202 MHz, $CD_3OD$) δ 11.13 (d, *J* = 62.1 Hz), 9.93 (d, *J* = 58.4 Hz). Characterization data is consistent with the literature.[4]

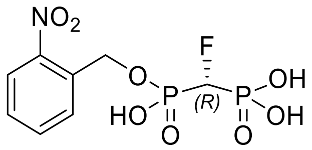

**[(*R*)-Fluoro({hydroxy[(2-nitrophenyl)methoxy]phosphoryl})-methyl]phosphonic acid 27.** This compound was synthesized according to the literature[4] on a reduced scale. (*R*)-CHF bisphosphonic acid derivative **26** (80 mg, 0.18 mmol) was dissolved in 0.1 M HCl (2.5 mL) and stirred at rt overnight. Solvent was removed under reduced pressure and co-evaporated with methanol (3 x 2 mL). **27** was obtained as the triacid product through passage of a pipet column of DOWEX $H^+$ using a mixture of MeOH/water (1:1) as eluent. The acid was treated with tetra-*n*-butylammonium hydroxide (3 eq) and brought to pH 8.3. After removal of solvents under reduced pressure and co-evaporation with water (3 x 2 mL), **27** was obtained as the tris(tetra-*n*-butylammonium) salt; white solid (59.2 mg, 0.18 mmol, quant.)

$^1$H NMR (400 MHz, $CD_3OD$) δ 8.16 (d, *J* = 7.9 Hz, 1H), 8.09 (d, *J* = 8.2 Hz, 1H), 7.73 (t, *J* = 7.6 Hz, 1H), 7.47 (t, *J* = 7.6 Hz, 1H), 5.51 (d, *J* = 7.2 Hz, 2H). $^{31}$P NMR (202 MHz, $D_2O$) δ 10.69 (d, *J* = 61.1 Hz), 8.35 (d, *J* = 62.3 Hz). Characterization data are consistent with the literature.[4]

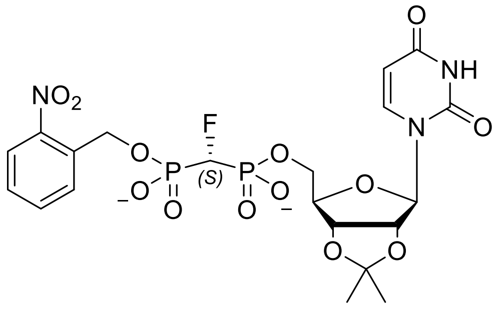

**(2-nitrophenyl)methyl [(*S*)-{[(4*R*,6*R*)-6-(2,4-dioxo-1,2,3,4-tetrahydropyrimidin-1-yl)-2,2-dimethyl-tetrahydro-2H-furo[3,4-d][1,3]dioxol-4-yl]methyl phosphonato}(fluoro)methyl]phosphonate 28.** According to General Method 1, **22** (47 mg, 0.1 mmol, 1.3 eq) was added to a mixture of **27** as the tris(tetra-*n*-butylammonium) salt (27.6 mg, 0.08 mmol, 1 eq) in DMF (130 μL). The reaction mixture was heated to 50˚C for 6 h. The crude mixture was purified by RP-HPLC, Luna Phenomenex C-18 column (5 μm, 250 mm × 21 mm) with a gradient method from 10 - 50% MeCN in 0.25 M ammonium acetate, pH 6.5, at a flow rate of 10.0 mL/min and UV detection (280 nm). After removal of solvents under reduced pressure and co-evaporation with water (3 x 2 mL), the desired product **28** was obtained as a yellow solid (22.1 mg 0.037 mmol, 46%).

RP-HPLC purification retention time = 16.19 min. $^1$H NMR (400 MHz, $CD_3OD$) δ 8.13 – 8.05 (m, 2H), 7.94 (d, *J* = 8.0 Hz, 1H), 7.73 (t, *J* = 7.6 Hz, 1H), 7.49 (t, *J* = 7.7 Hz, 1H), 5.96 (d, *J* = 3.4 Hz, 1H), 5.75 (d, *J* = 8.1 Hz, 1H), 5.48 (d, *J* = 6.7 Hz, 2H), 4.34 (d, *J* = 3.0 Hz, 1H), 4.23 (q, *J* = 3.0 Hz, 2H), 1.53 (s, 3H), 1.33 (s, 3H). $^{31}$P NMR (202 MHz, $CD_3OD$) δ 9.21 (d, *J* = 59.2 Hz).

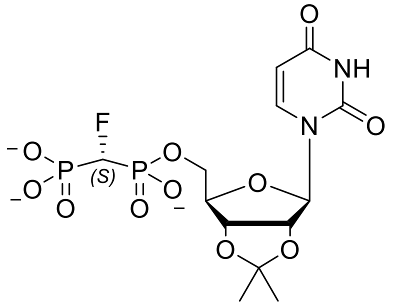

**[(4*R*,6*R*)-6-(2,4-dioxo-1,2,3,4-tetrahydropyrimidin-1-yl)-2,2-dimethyl-tetrahydro-2H-furo[3,4-d][1,3]dioxol-4-yl]methyl [(*S*)-fluoro(phosphonato)methyl]phosphonate 29.** In a quartz cuvette, 2-nitrobenzyl fluoromethylene uridine bisphosphonate **28** (22.1 mg, 0.037 mmol) was dissolved in water (2 mL) and irradiated (wavelength = 365nm) for 48 h. The reaction solution was extracted with DCM (3 x 5 mL), and the aqueous layer was collected. Solvents were removed under reduced pressure. the desired product **29** was obtained as a white solid (12.9 mg, 0.028 mmol, 76%).

$^1$H NMR (600 MHz, $D_2O$) δ 7.84 (d, *J* = 8.3 Hz, 1H), 5.88 (d, *J* = 2.5 Hz, 1H), 5.86 (d, *J* = 8.1 Hz, 1H), 5.01 (t, *J* = 2.9 Hz, 2H), 4.71 (t, *J* = 12.5 Hz, 1H), 4.53 (s, 1H), 4.24 – 4.13 (m, 2H), 1.57 (s, 3H), 1.37 (s, 3H). $^{31}$P NMR (243 MHz, $D_2O$) δ 9.21 (d, *J* = 59.2 Hz).

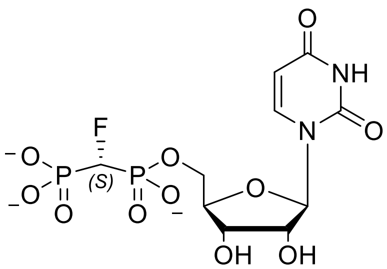

**[(2*R*,5*R*)-5-(2,4-dioxo-1,2,3,4-tetrahydropyrimidin-1-yl)-3,4-dihydroxyoxolan-2-yl]methyl [(*S*)-fluoro-(phosphono)methyl]phosphonate, (*S*)-CHF-UBP 30.** According to General Method 2, **29** (12.9 mg, 0.028 mmol) was deprotected in acidic conditions with 1 M HCl (1 mL) for 1h. The solvent was removed under reduced pressure and co-evaporated with water (3 x 2 mL) to afford 30 as a colorless film (11.8mg, 0.028 mmol, quant.).

$^{1}H$ NMR (600 MHz, $CD_3OD$) δ 7.86 (d, *J* = 8.1 Hz, 1H), 5.94 (s, 1H), 5.73 (d, *J* = 8.1 Hz, 1H), 5.13 (dt, *J* = 45.5, 13.1 Hz, 1H), 4.44 – 4.39 (m, 1H), 4.39 – 4.34 (m, 1H), 4.22 (t, *J* = 4.9 Hz, 1H), 4.18 (t, *J* = 5.2 Hz, 1H), 4.16 (s, 1H). $^{31}P$ NMR (243 MHz, $D_2O$) δ 10.22 (dd, *J* = 65.6, 12.6 Hz), 9.25 (dd, *J* = 61.0, 12.5 Hz).

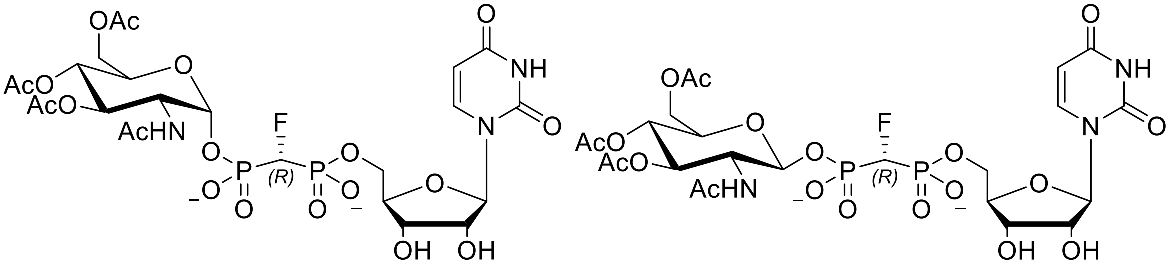

**[(2*R*,5*R*)-5-(2,4-dioxo-1,2,3,4-tetrahydropyrimidin-1-yl)-3,4-dihydroxyoxolan-2-yl]methyl [(*R*)-[(2*R*,5*S*)-4,5-bis(acetyloxy)-6-[(acetyloxy)methyl]-3-acetamidooxan-2-yl phosphono](fluoro)methyl] phosphonate and its β-anomer, (*R*) α/ β-GlcNAc(OAc)$_3$-CHF-UBP, α-anomer 31α and β-anomer 31β.** The addition of GlcNAc$_4$ was conducted According to General Method 3 with some modifications. **13** (39.8 mg, 0.12 mmol, 4 eq) was coupled with 30 (11.8 mg, 0.028 mmol, 1 eq) in anhydrous DMF (0.5 mL) by heating at 45˚C for 17h. The crude mixture was purified by RP-HPLC, Hamilton PRP-1 column (5 µm, 250 mm × 21 mm), 261 nm, isocratic mode, using 8.3% acetonitrile in 0.24 M triethylammonium bicarbonate, pH 6.5, at a flow rate of 8.0 mL/min. After removal of solvents under reduced pressure, the α and β anomers, **31α** and **31β**, were obtained as a colorless film (11.9mg, 0.016 mmol, 57%).

RP-HPLC purification retention time = 18.7 min. $^{1}H$ NMR (400 MHz, $D_2O$) δ 8.01 (d, *J* = 7.6 Hz, 1H), 5.95 (dd, *J* = 13.7, 6.0 Hz, 2H), 5.26 (dd, *J* = 10.6, 9.4 Hz, 2H), 4.45 – 4.11 (m, 8H), 4.07 – 3.98 (m, 2H), 2.10 (s, 3H), 2.06 (s, 3H), 2.02 (s, 3H), 1.95 (s, 3H). $^{19}F$ NMR (376 MHz, $D_2O$) δ -220.64 (β-anomer appears as q, *J* = 58.5 Hz), -229.05 (α-anomer appears as td, *J* = 67.1, 44.3 Hz).

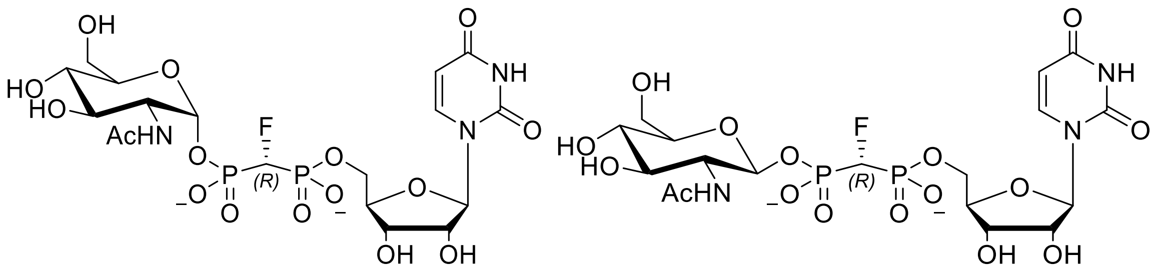

**[(2*R*,5*R*)-5-(2,4-dioxo-1,2,3,4-tetrahydropyrimidin-1-yl)-3,4-dihydroxyoxolan-2-yl]methyl [(*R*)-[(2*R*,5*S*)-4,5-bis(acetyloxy)-6-[(acetyloxy)methyl]-3-acetamidooxan-2-yl phosphono](fluoro)methyl]phosphonate, mixture of α- and β-anomers 8α (*R*) and 8β (*R*).** GlcNAc$_4$ FUBP **31α** and **31β** were deprotected under basic conditions According to General Method 4. The α- and β-anomers (11.9 mg, 0.016 mmol) were dissolved in water (2 mL), then the solution was brought to pH 10.5 with $NH_4OH$. Reaction mixture was heated to 27˚C and stirred overnight. The anomers were purified by RP-HPLC, Hamilton PRP-1 column (5 µm, 250 mm × 10 mm), isocratic mode, using 15% acetonitrile in 0.1 M triethylammonium bicarbonate, pH 7.5, at a flow rate of 3.0 mL/min and UV detection (261 nm). After removal of solvents under reduced pressure, the desired products **(*R*)-8α** and **(*R*)-8β** were obtained as a colorless film (3.1 mg, 0.005 mmol, 31%).

RP-HPLC purification retention time = 3.70 min. $^{1}$H NMR (500 MHz, $D_2O$) δ 8.04 (d, *J* = 8.2 Hz, 1H), 6.00 (d, *J* = 4.2 Hz, 1H), 5.97 (d, *J* = 8.1 Hz, 1H), 5.55 (α-anomer H1" in Glc appears as dd, *J* = 7.2, 3.2 Hz, 0H), 5.08 (β-anomer H1" in Glc appears as t, *J* = 8.4 Hz, 1H), 4.91 (t, *J* = 12.8 Hz, 1H), 4.42 (d, *J* = 4.0 Hz, 2H), 4.32 – 4.20 (m, 3H), 3.94 (dd, *J* = 12.5, 2.2 Hz, 1H), 3.85 – 3.72 (m, 2H), 3.64 – 3.44 (m, 3H), 2.06 (s, 3H). $^{19}$F NMR (470 MHz, $D_2O$) δ -219.60 (β-anomer appears as td, *J* = 60.7, 45.9 Hz), -220.49 (α-anomer appears as td, *J* = 61.3, 46.0 Hz).$^{31}$P NMR (202 MHz, $D_2O$) δ 9.61 (d, *J* = 60.1 Hz), 8.80 (d, *J* = 61.5 Hz).

### Spectra, chromatograms and other Figures

#### Spectra and chromatograms of GlcNAc-$CH_2$-UBP analogues and intermediates

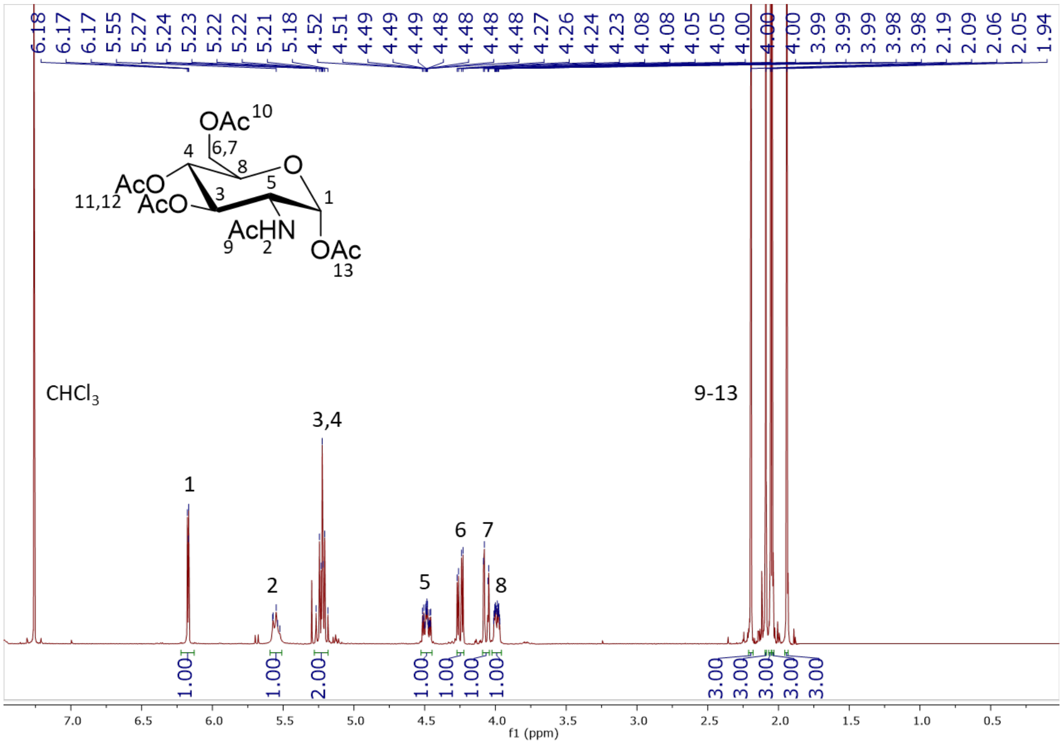

**Figure S6.** $^1H$ NMR (400 MHz, $CDCl_3$) of **12**

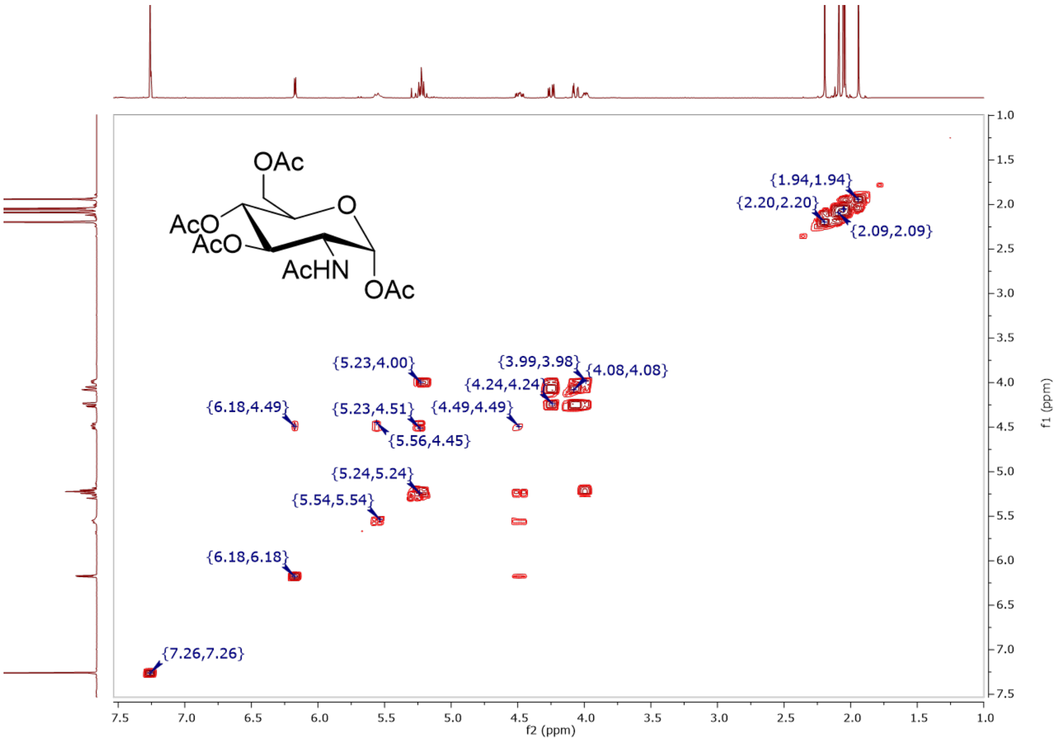

**Figure S7.** COSY (400 MHz, $CDCl_3$) of **12**

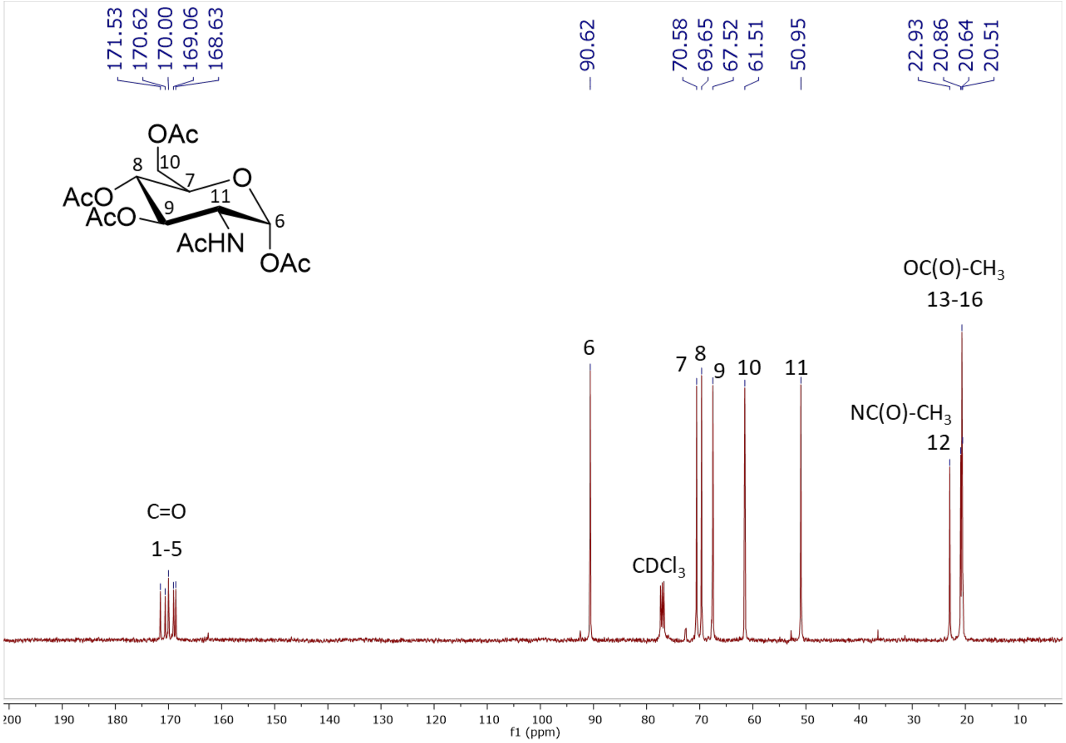

**Figure S8.** $^{13}$C NMR (101 MHz, CDCl$_3$) of **12**

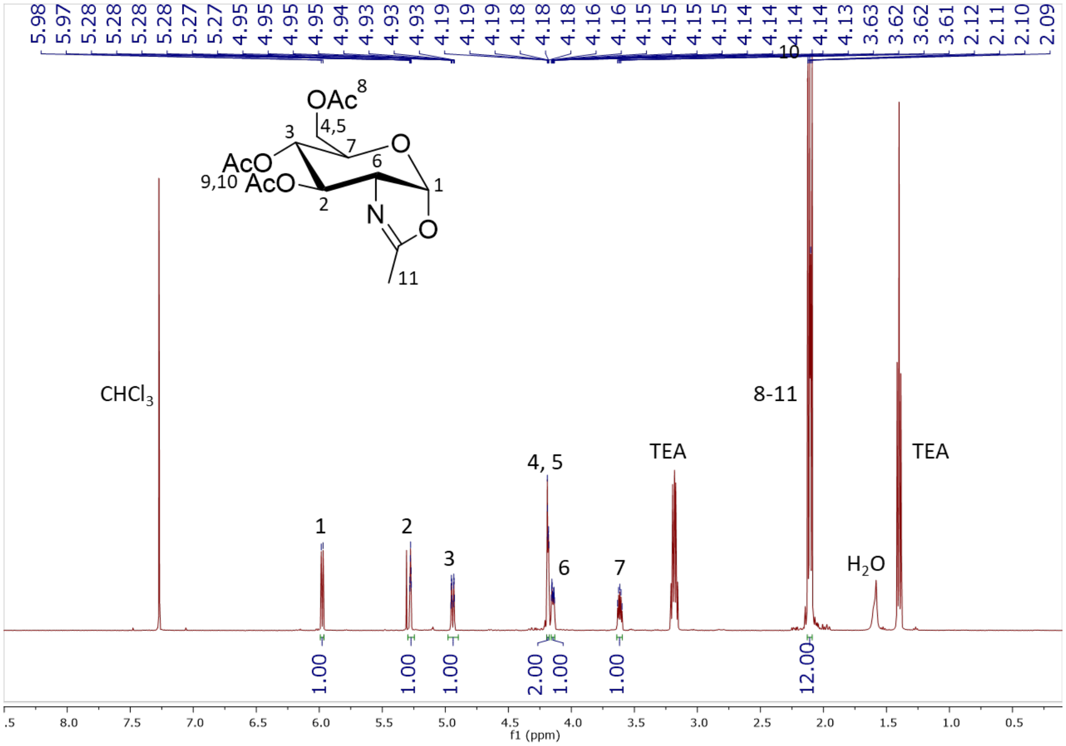

**Figure S9.** $^{1}$H NMR (500 MHz, CDCl$_3$) of **13**

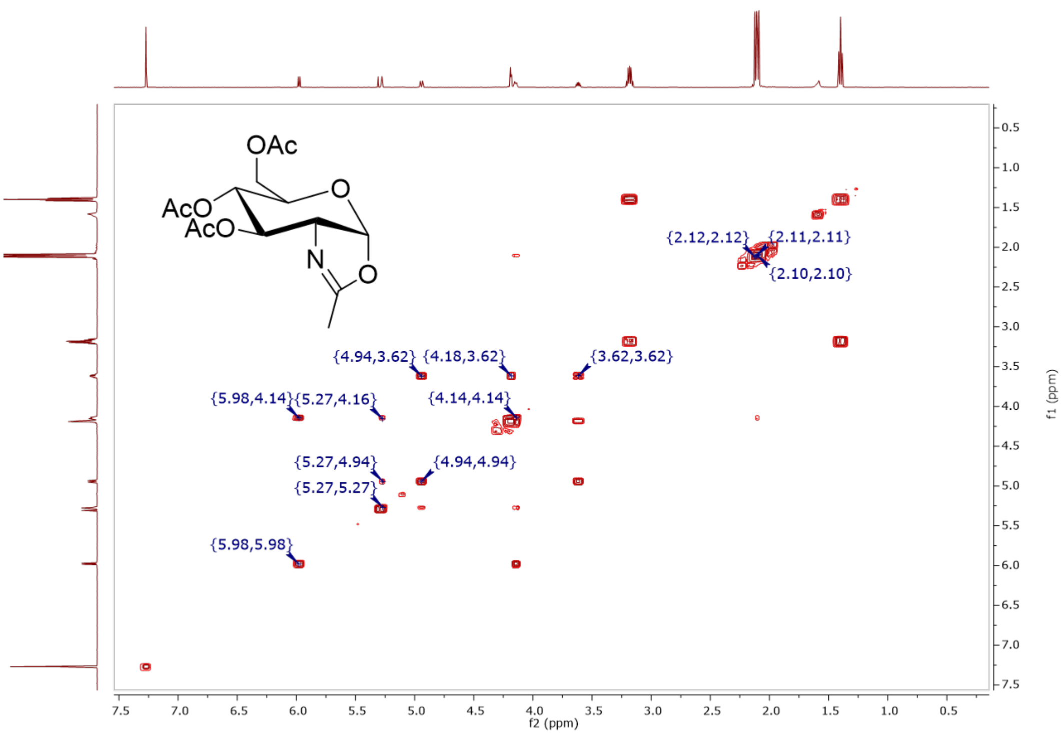

**Figure S10.** COSY (500 MHz, $CDCl_3$) of **13**

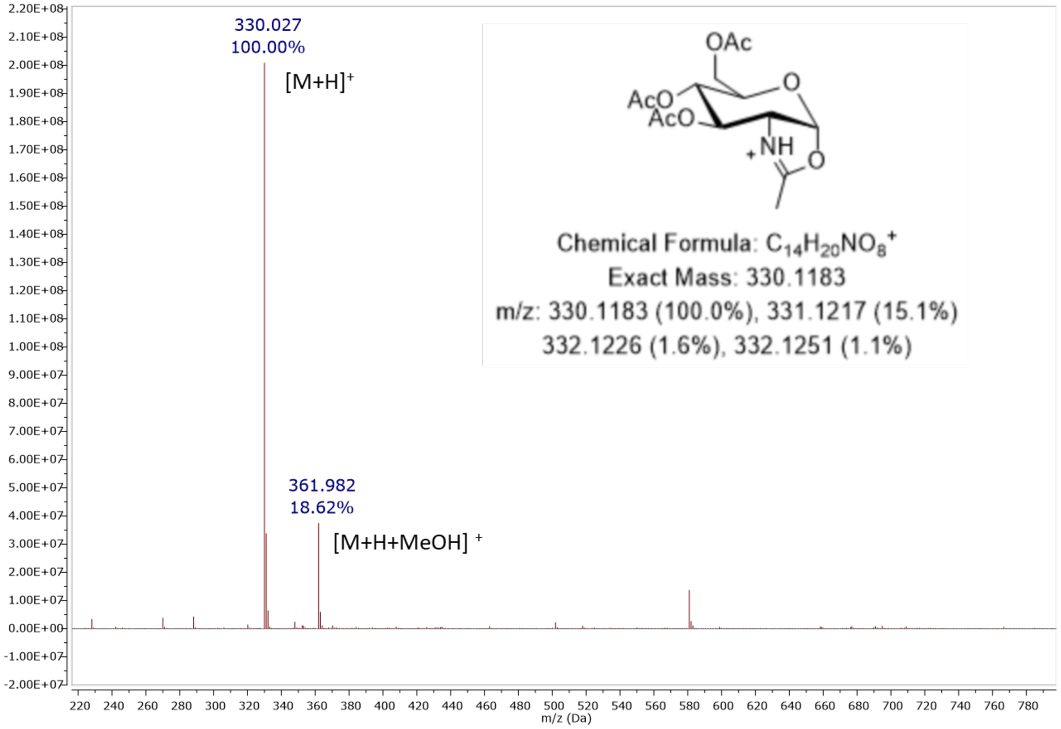

**Figure S11.** MS (ESI): $[M + H]^+$ of **13**

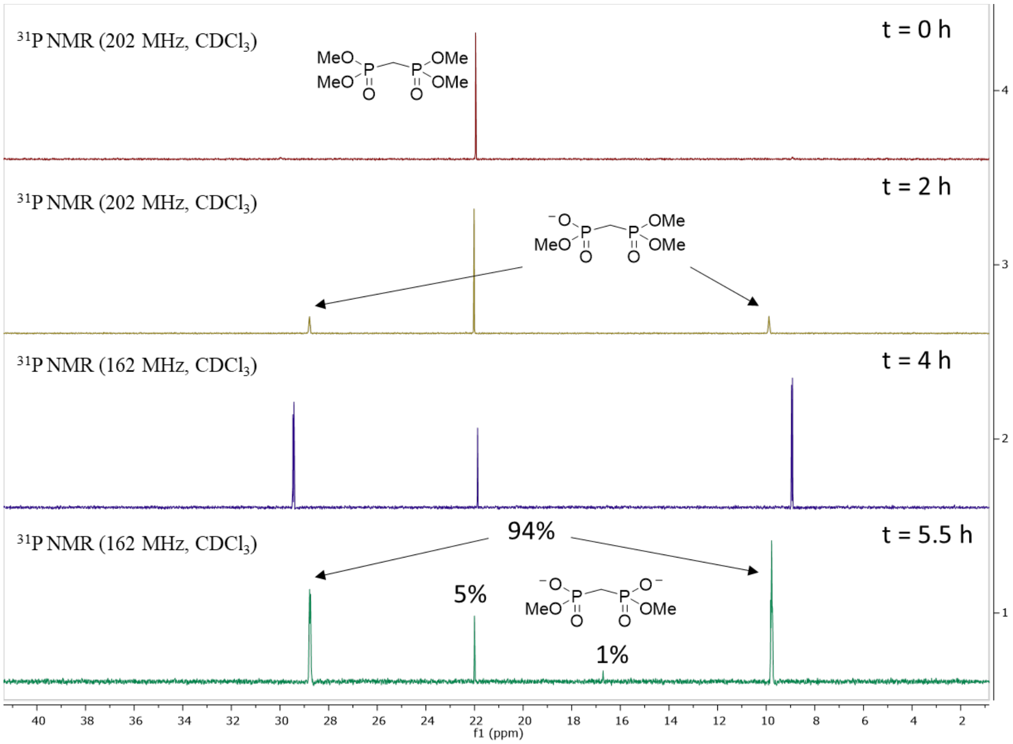

**Figure S12.** Progress of mono demethylation reaction over time by $^{31}$P NMR of **14**

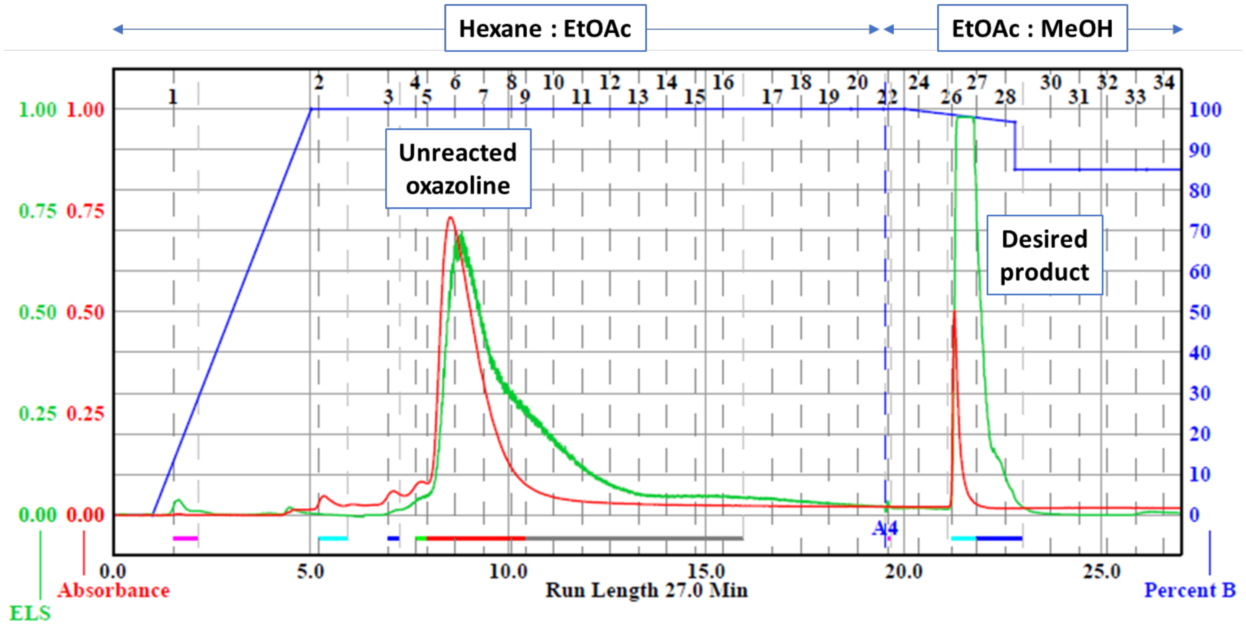

**Figure S13.** Purification of **15** using automated flash column chromatography

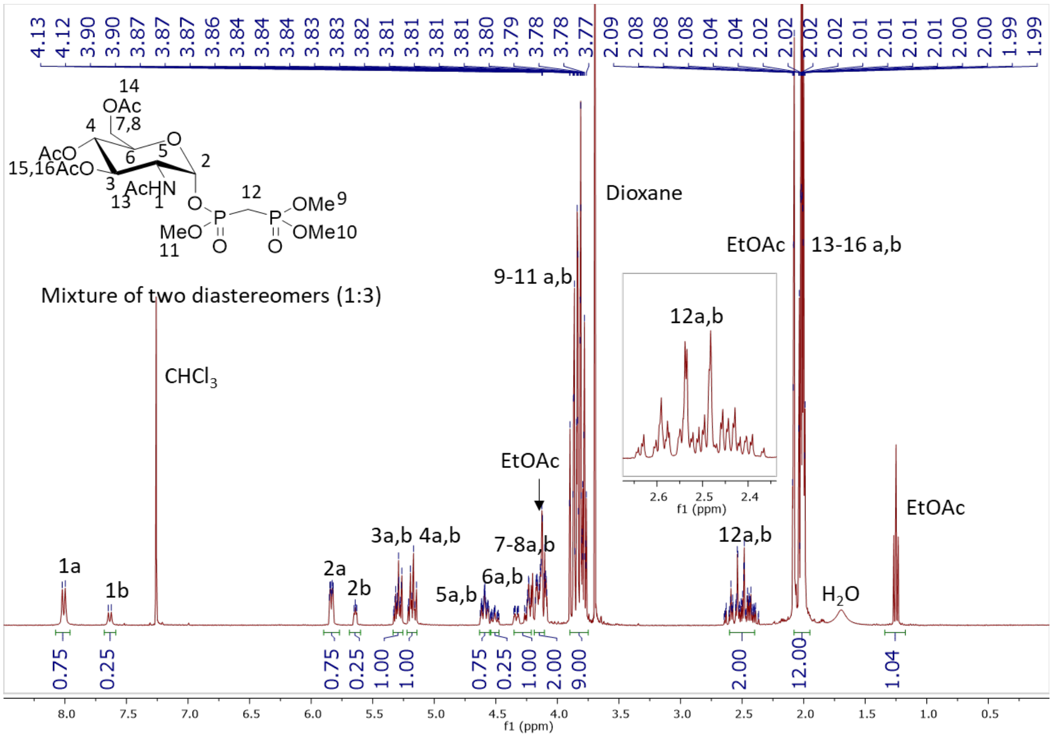

**Figure S14.** $^{1}H$ NMR (400 MHz, $CDCl_3$) of **15**

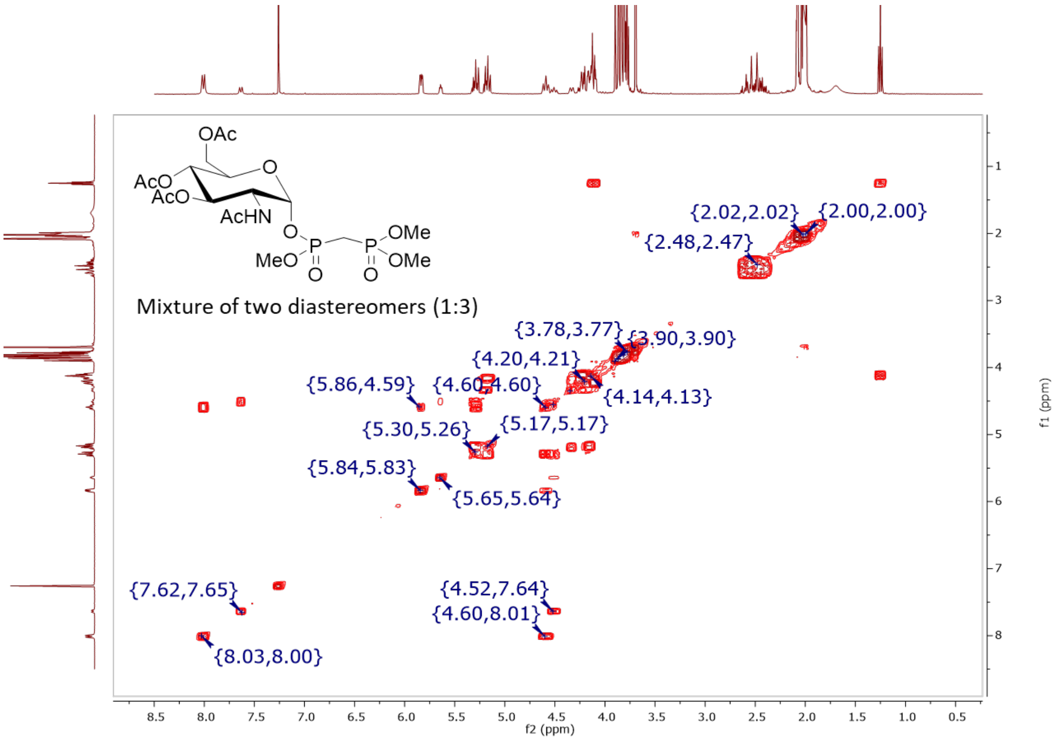

**Figure S15.** COSY (400 MHz, $CDCl_3$) of **15**

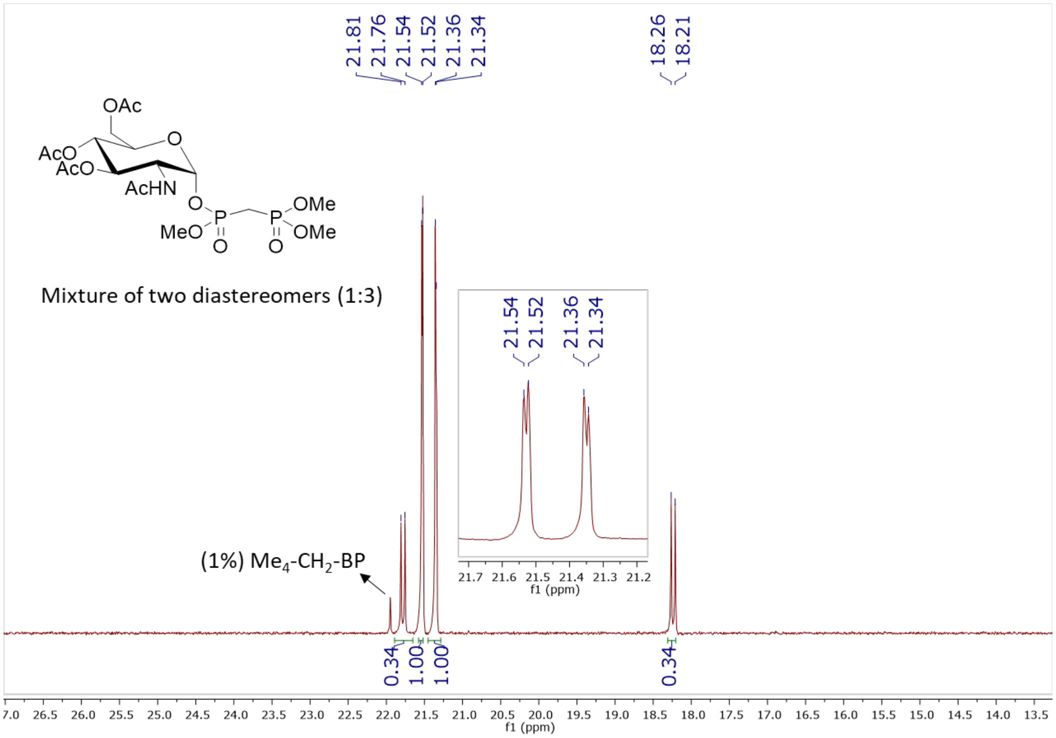

**Figure S16.** $^{31}$P NMR (162 MHz, $CDCl_3$) of **15**

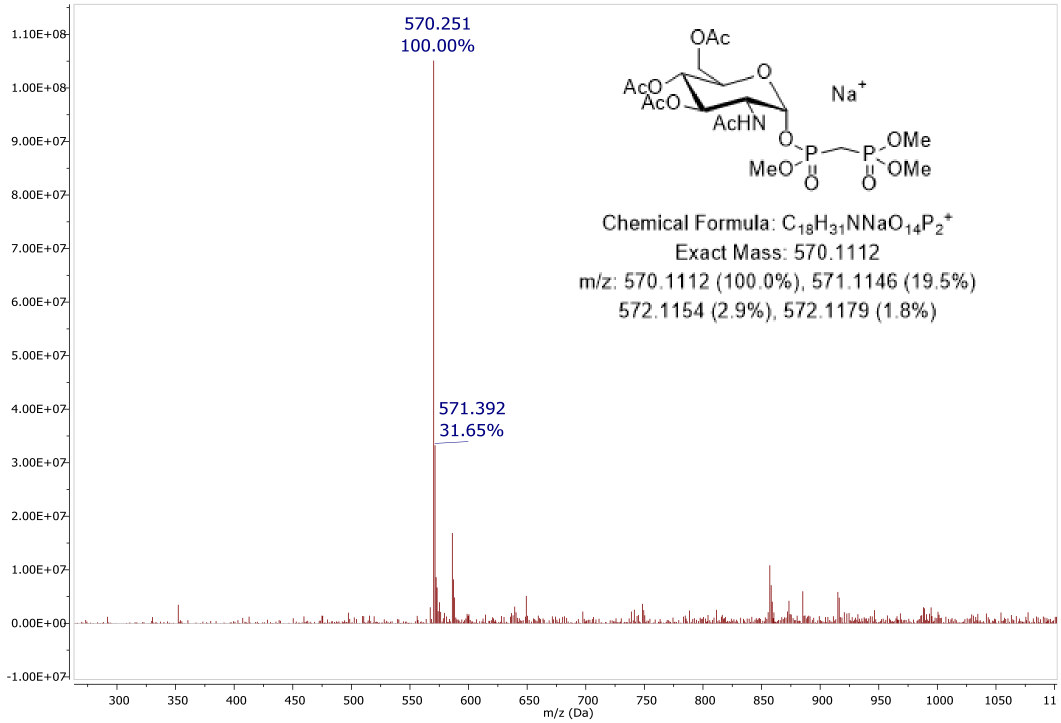

**Figure S17.** MS (ESI): $[M + Na]^+$ of **15**

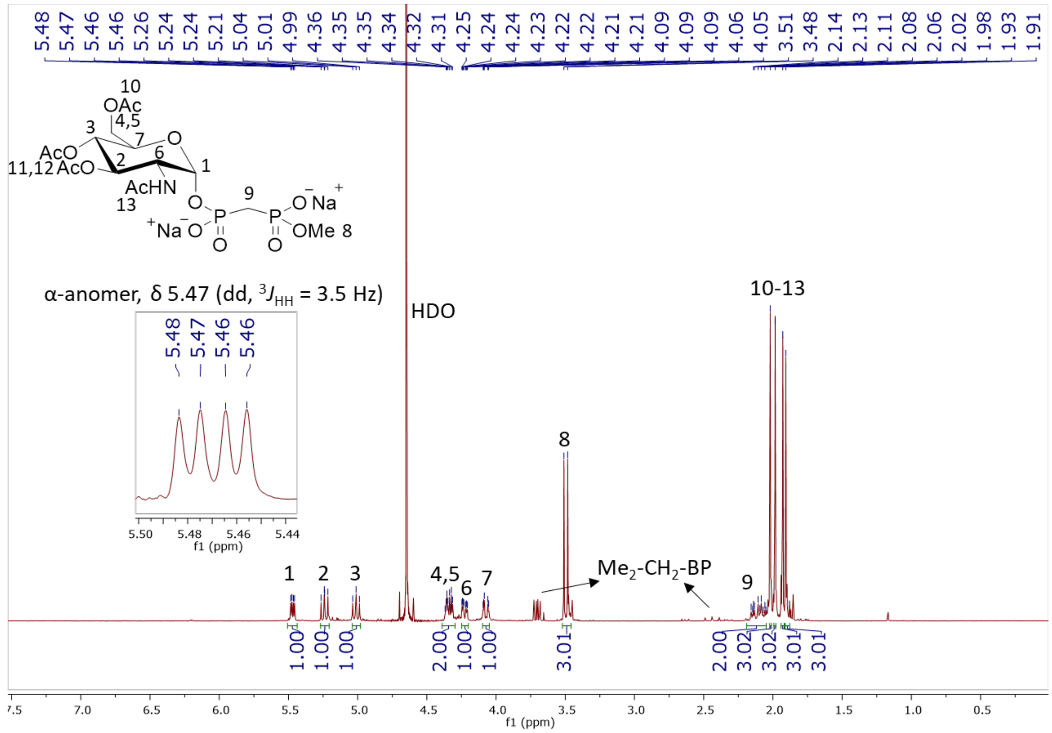

**Figure S18.** $^{1}H$ NMR (400 MHz, $D_2O$) of **16**

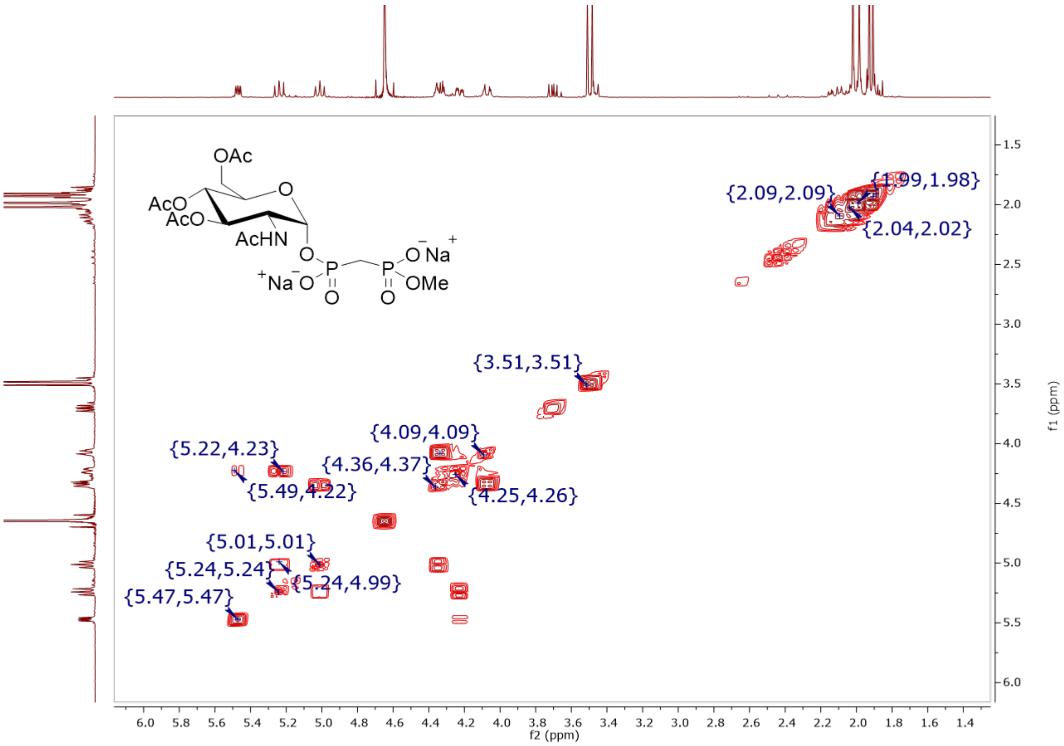

**Figure S19.** COSY (400 MHz, $D_2O$) of **16**

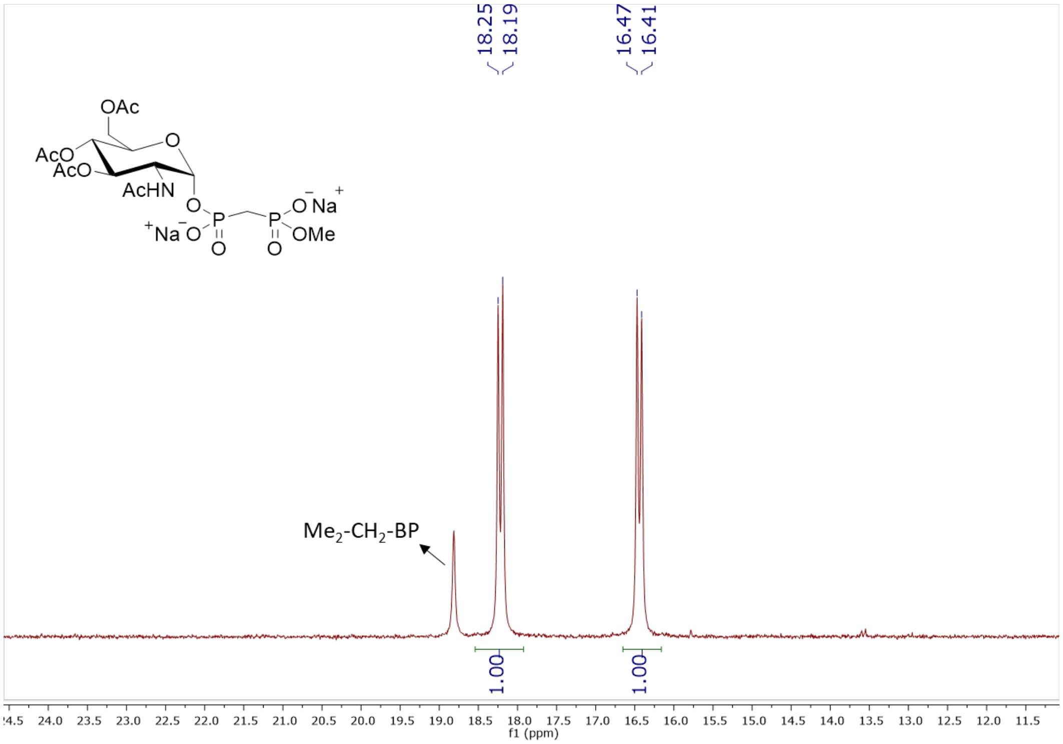

**Figure S20.** $^{31}P$ NMR (202 MHz, $D_2O$, pH 7) of **16**

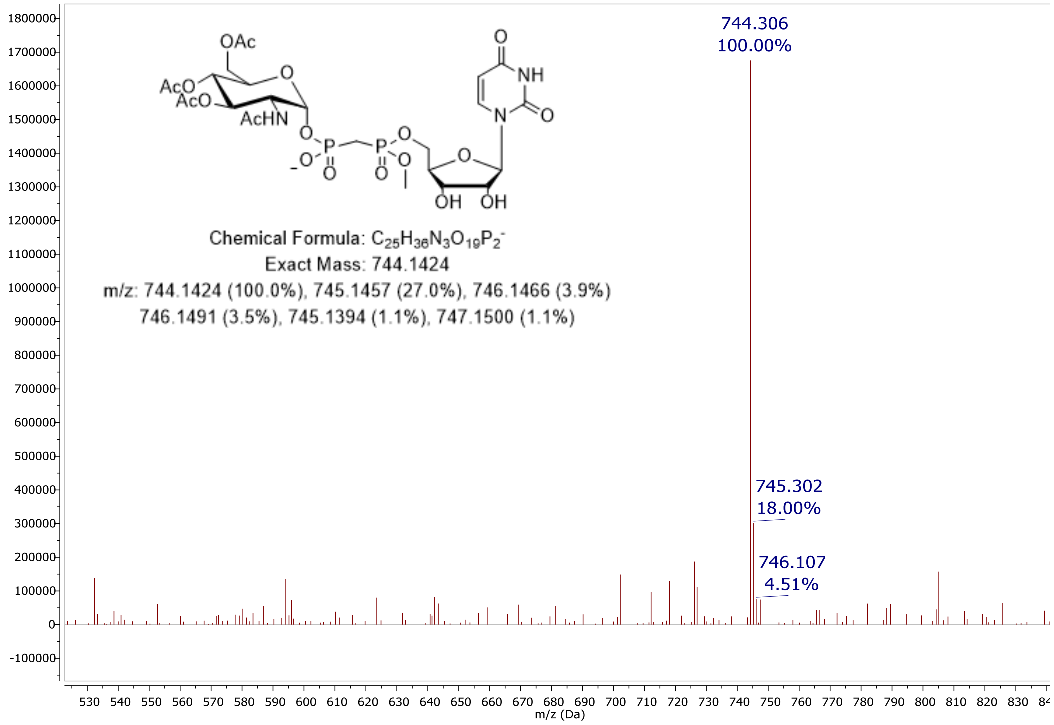

**Figure S21.** MS (ESI): [M – H]$^-$ of **19**

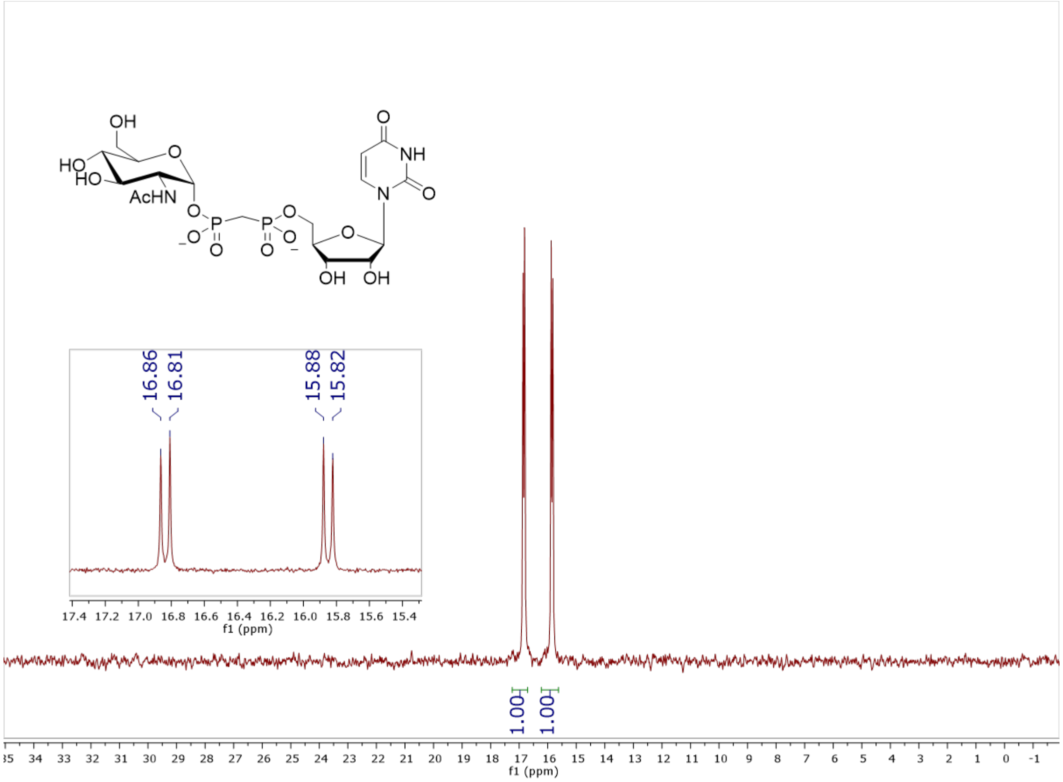

**Figure S22.** $^{31}$P NMR (202 MHz, $D_2O$, pH 7) of **11**

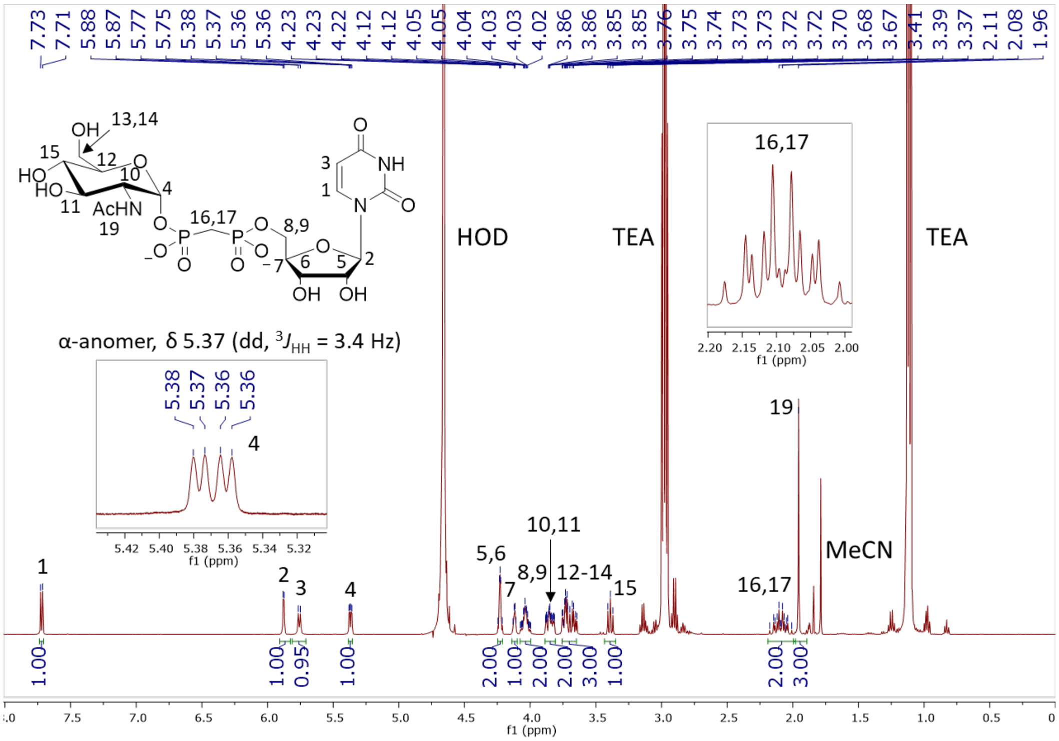

**Figure S23.** $^{1}$H NMR (500 MHz, $D_2O$) of **11**

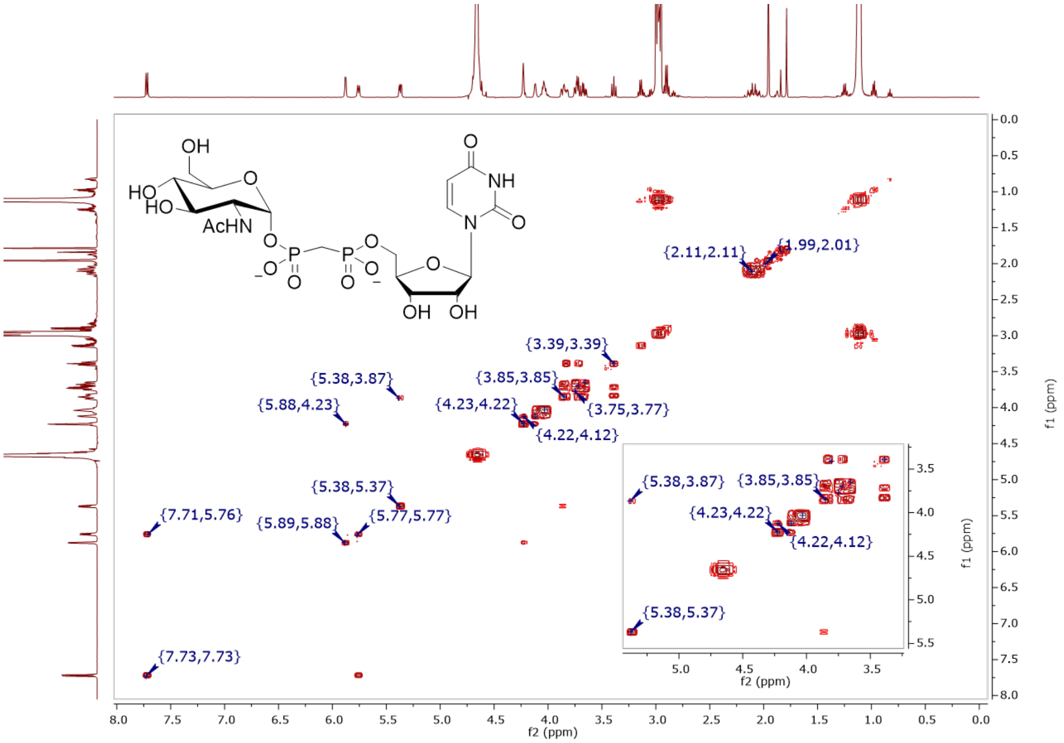

**Figure S24.** COSY (500 MHz, $D_2O$) of **11**

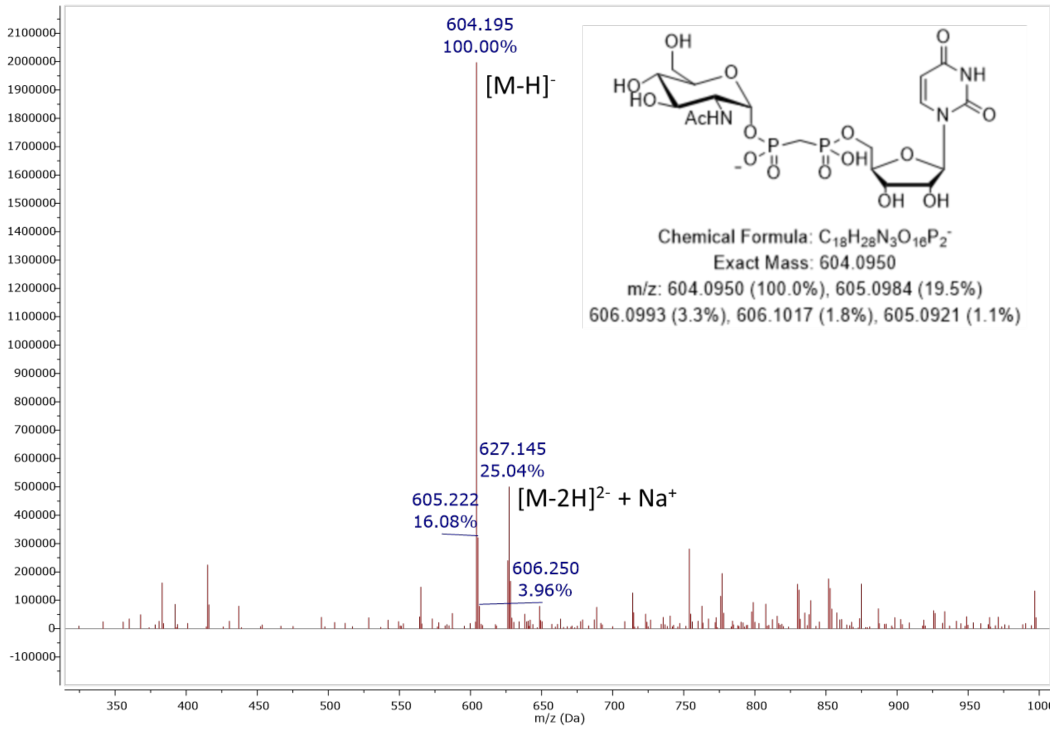

**Figure S25.** MS (ESI): $[M-H]^-$ of **11**

#### Elemental Composition Report

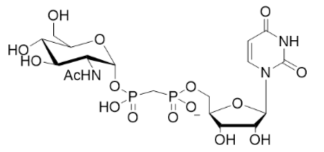

##### Single Mass Analysis

Tolerance = 5.0 PPM / DBE: min = -1.5, max = 50.0
Element prediction: Off
Number of isotope peaks used for i-FIT = 3

Monoisotopic Mass, Even Electron Ions
115 formula(e) evaluated with 1 results within limits (up to 50 closest results for each mass)
Elements Used:
C: 0-60 H: 0-100 N: 2-4 O: 15-17 P: 1-3
Pouya Haratipour, USCMCK12
Synapt_26237 21 (0.431) Cm (21-8)

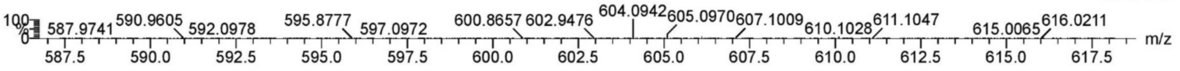

| | | | | | | | | |
|---|---|---|---|---|---|---|---|---|
| Minimum: | | | | -1.5 | | | | |
| Maximum: | | 5.0 | 5.0 | 50.0 | | | | |
| Mass | Calc. Mass | mDa | PPM | DBE | i-FIT | Norm | Conf(%) | Formula |
| 604.0942 | 604.0945 | -0.3 | -0.5 | 7.5 | 547.7 | n/a | n/a | C18 H28 N3 O16 P2 |

**Figure S26.** Elemental composition report from HRMS (ESI-TOF) [M – H]$^-$ of **11**

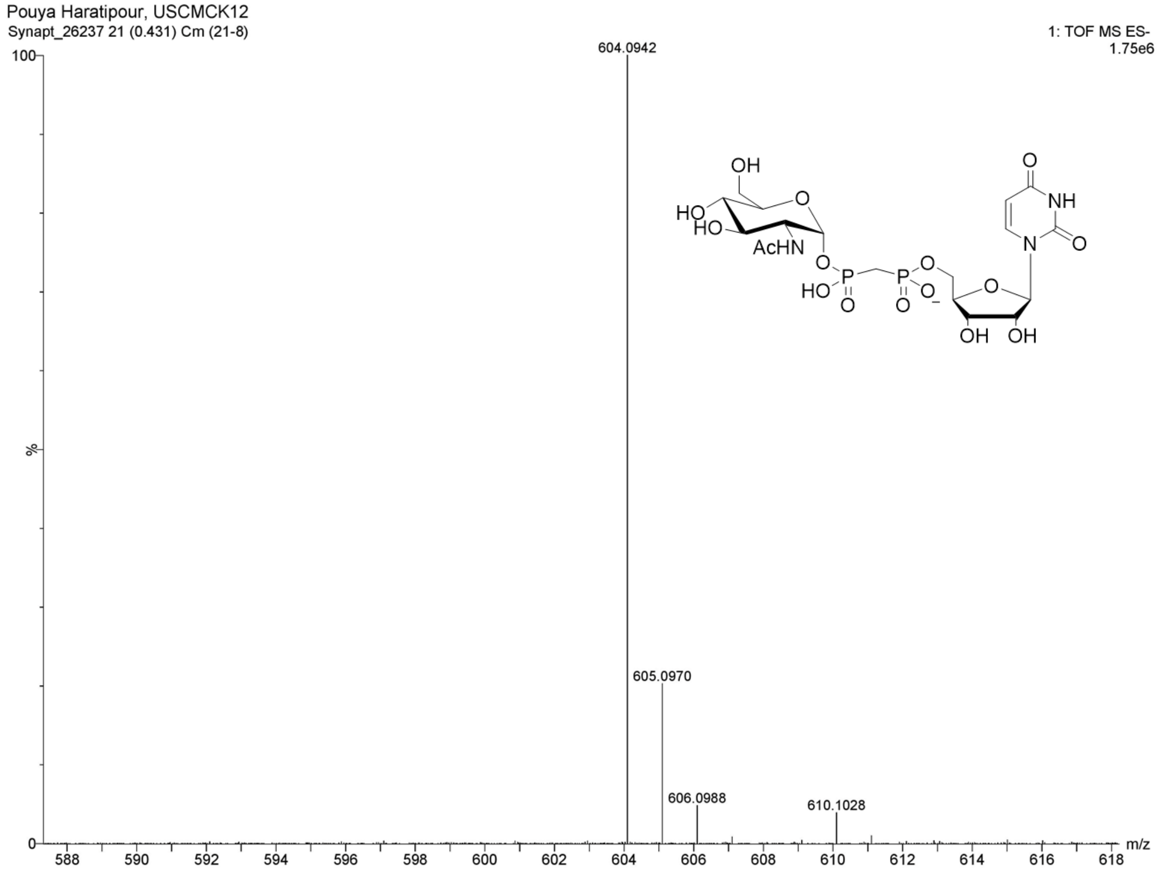

**Figure S27.** HRMS (ESI-TOF): [M – H]$^-$ of **11**

**Spectra and chromatograms of CXY-UBP analogues and intermediates**

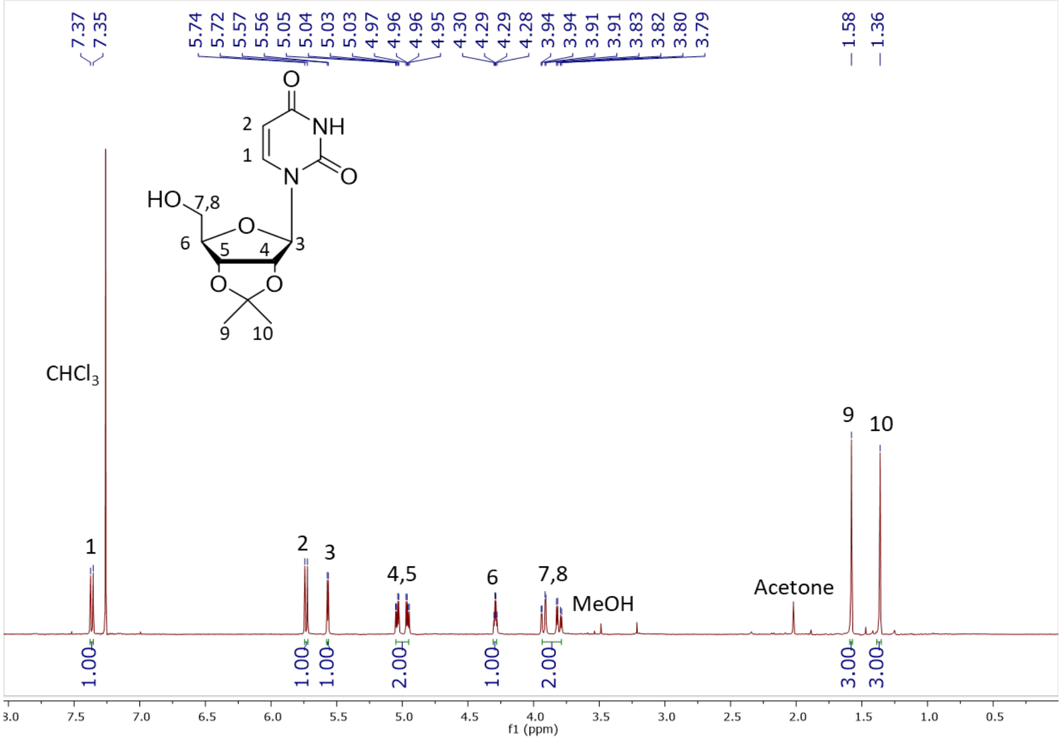

**Figure S28.** $^{1}H$ NMR (400 MHz, $CDCl_3$) of **21**

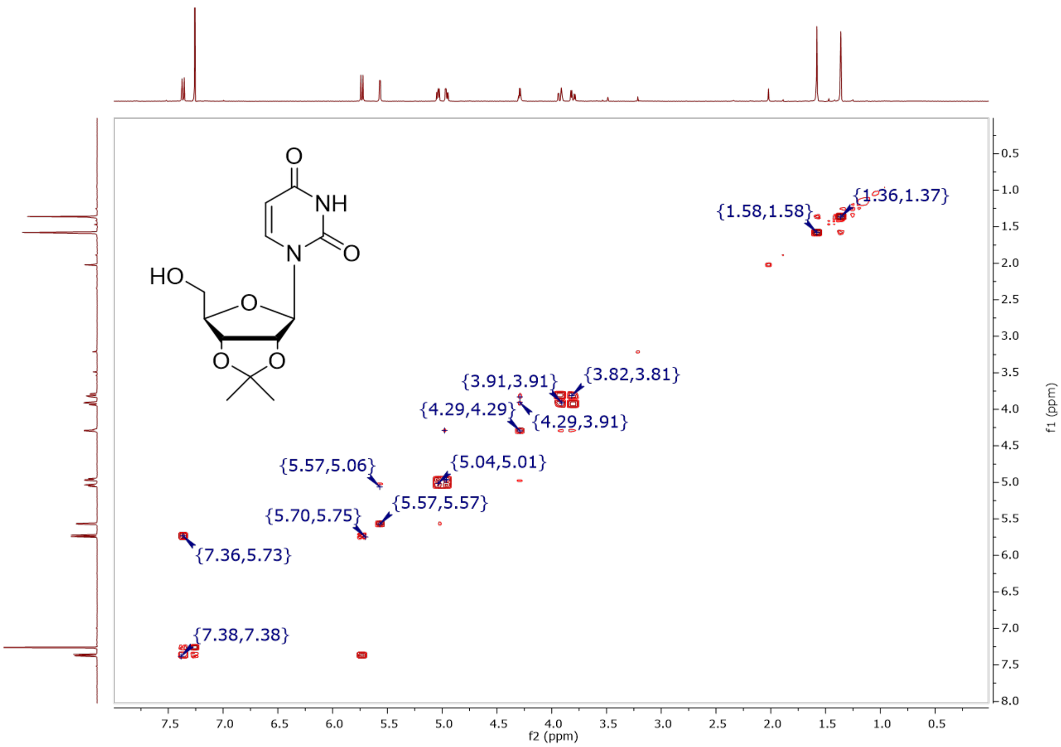

**Figure S29.** COSY (400 MHz, $CDCl_3$) of **21**

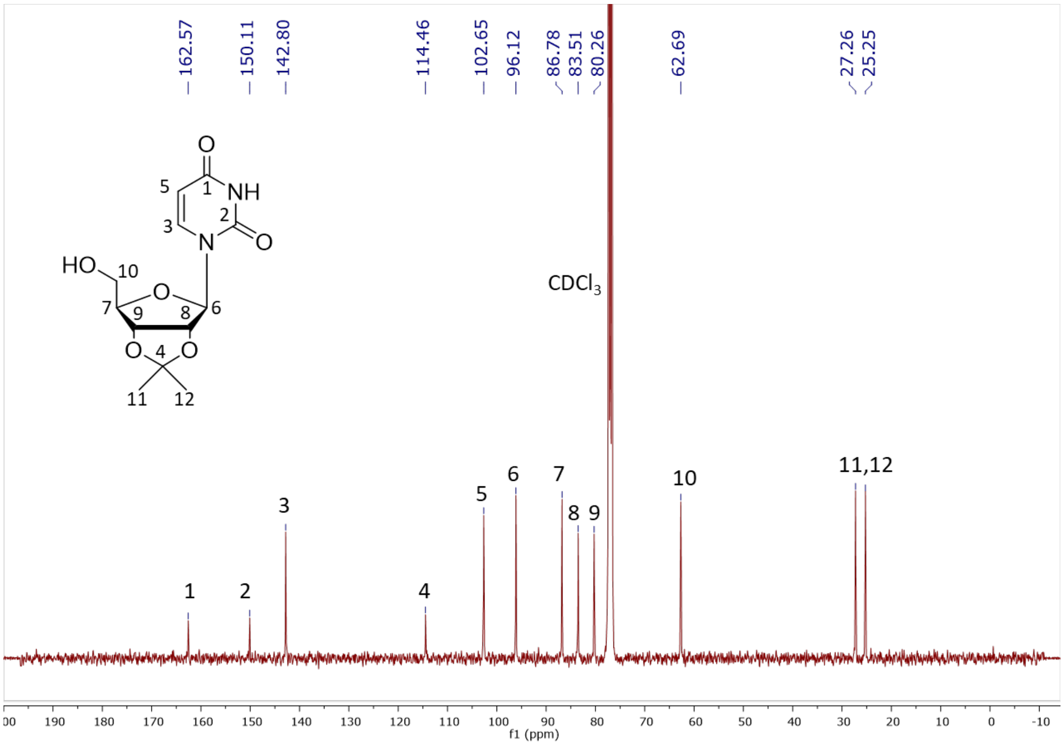

**Figure S30.** $^{13}C$ NMR (101 MHz, $CDCl_3$) of **21**

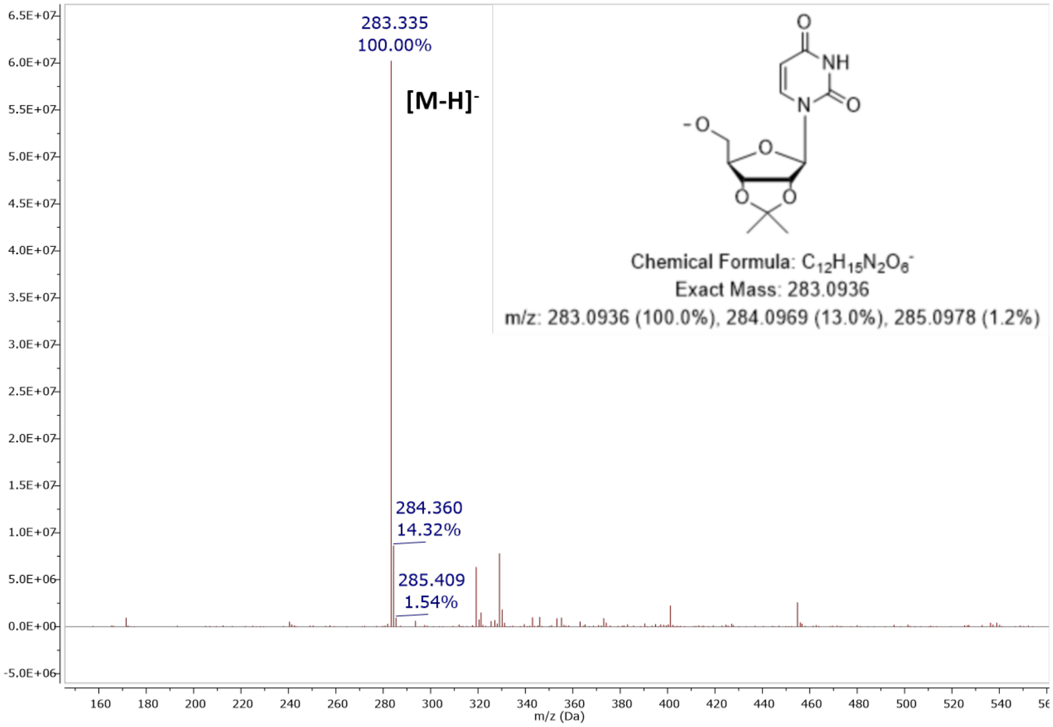

**Figure S31.** MS (ESI): $[M – H]^–$ of **21**

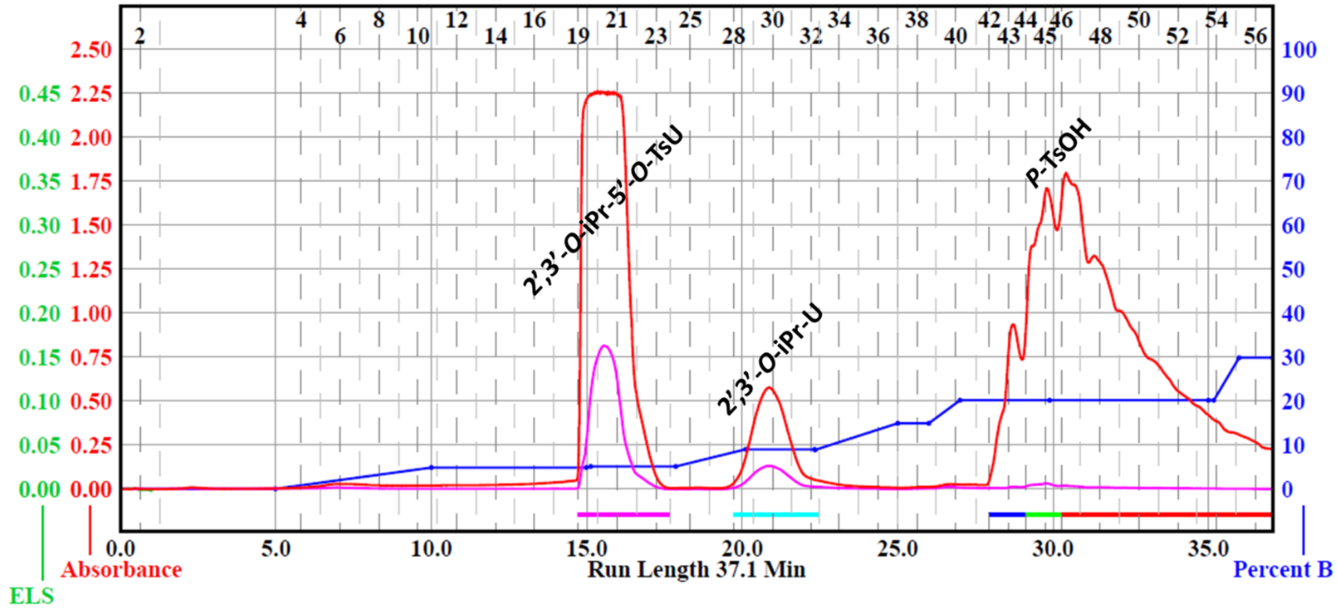

**Figure S32.** Purification of **22** using automated flash column chromatography

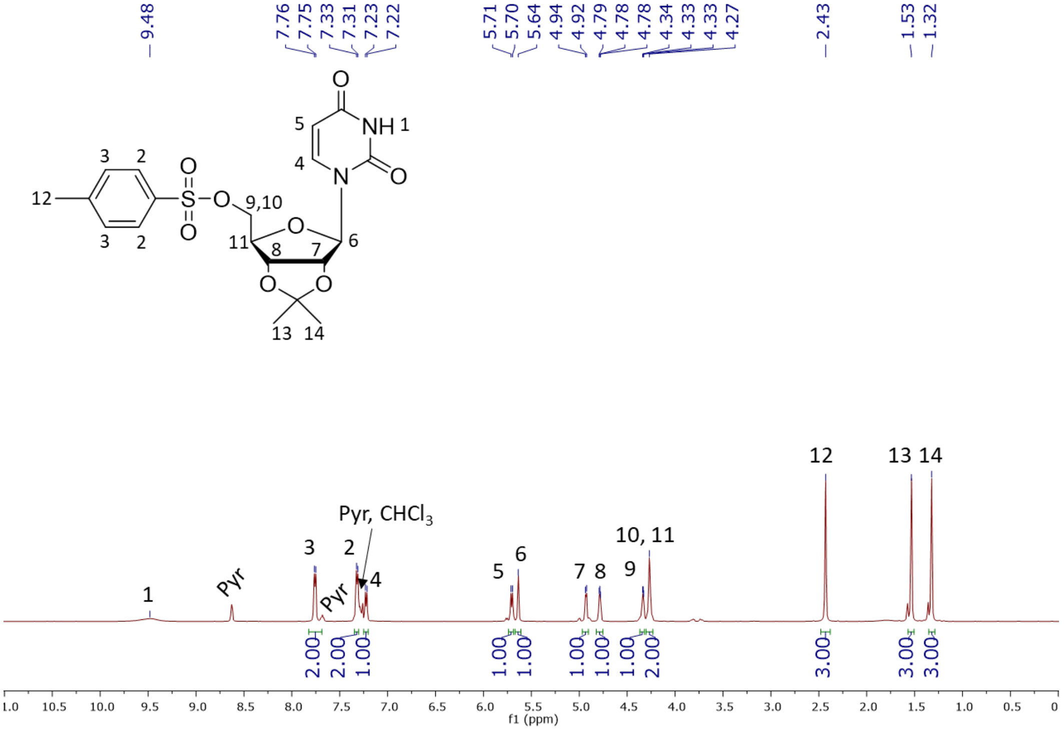

**Figure S33.** $^{1}$H NMR (500 MHz, CDCl$_3$) of **22**

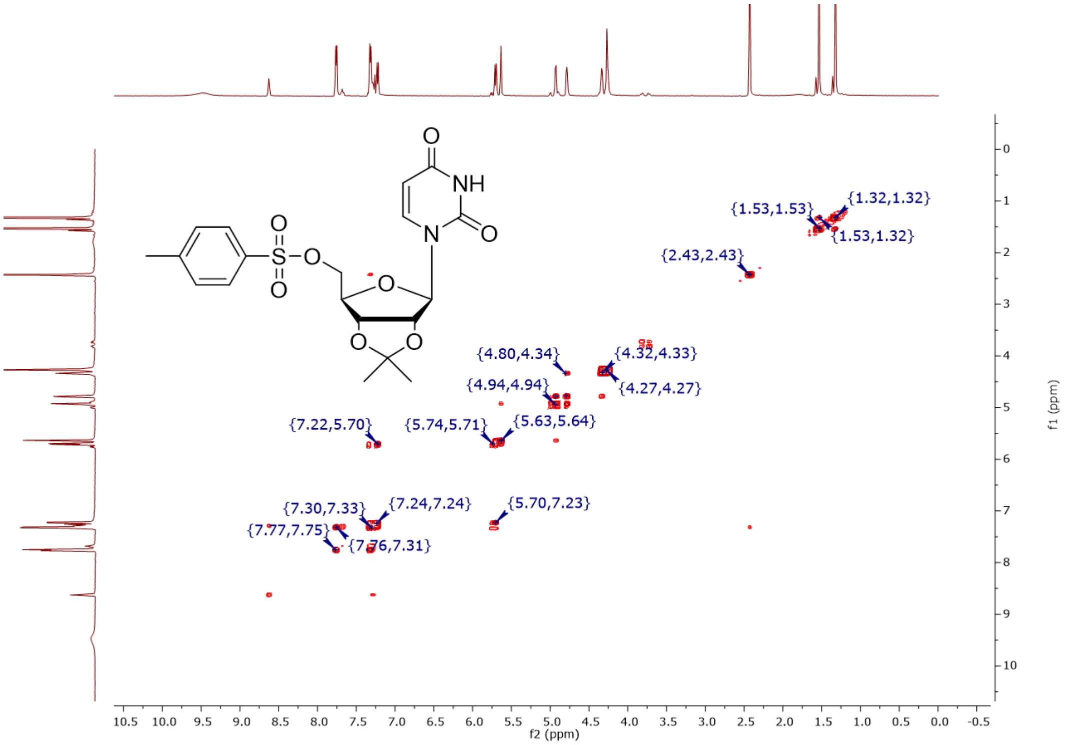

**Figure S34.** COSY (500 MHz, $CDCl_3$) of **22**

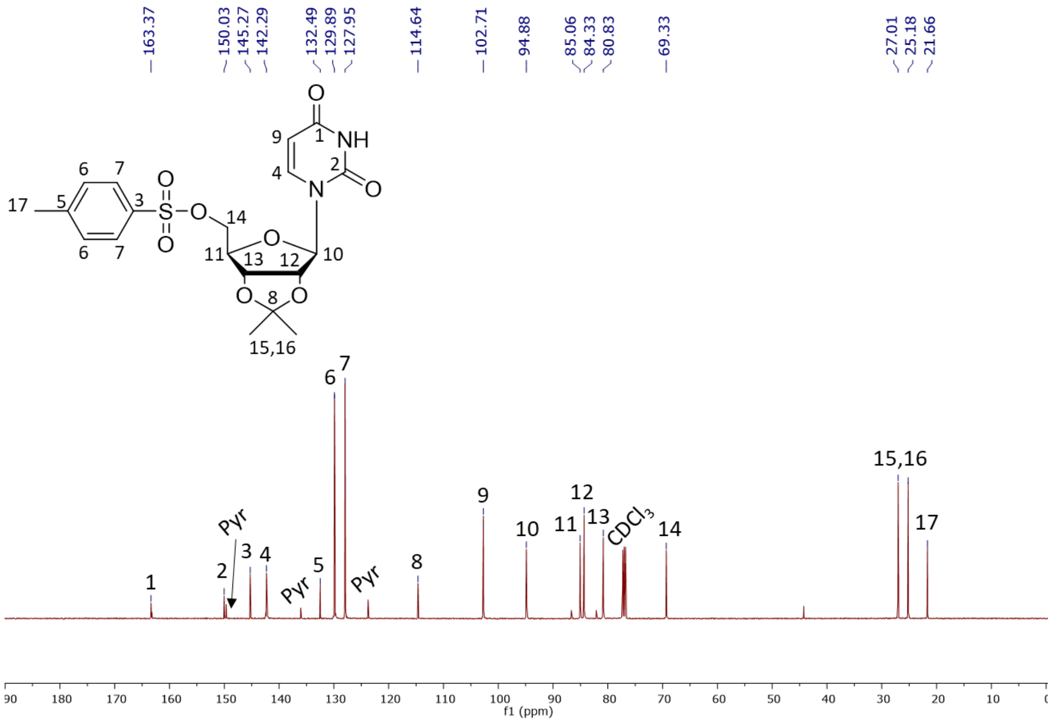

**Figure S35.** $^{13}C$ NMR (126 MHz, $CDCl_3$) of **22**

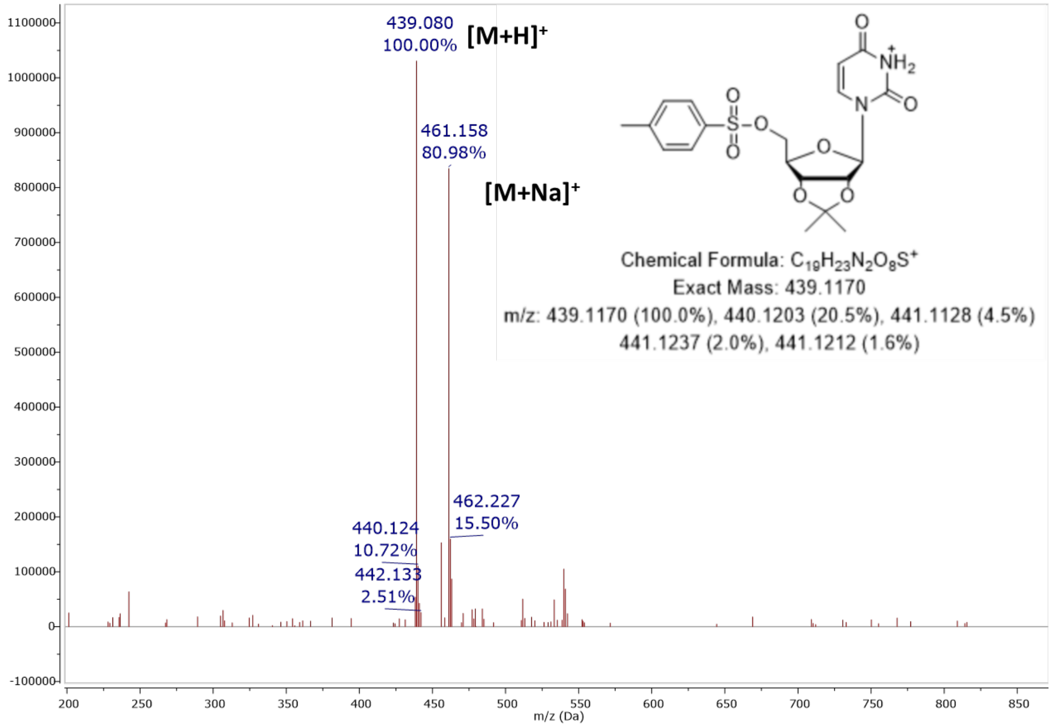

**Figure S36.** MS (ESI): $[M + H]^+$ of **22**

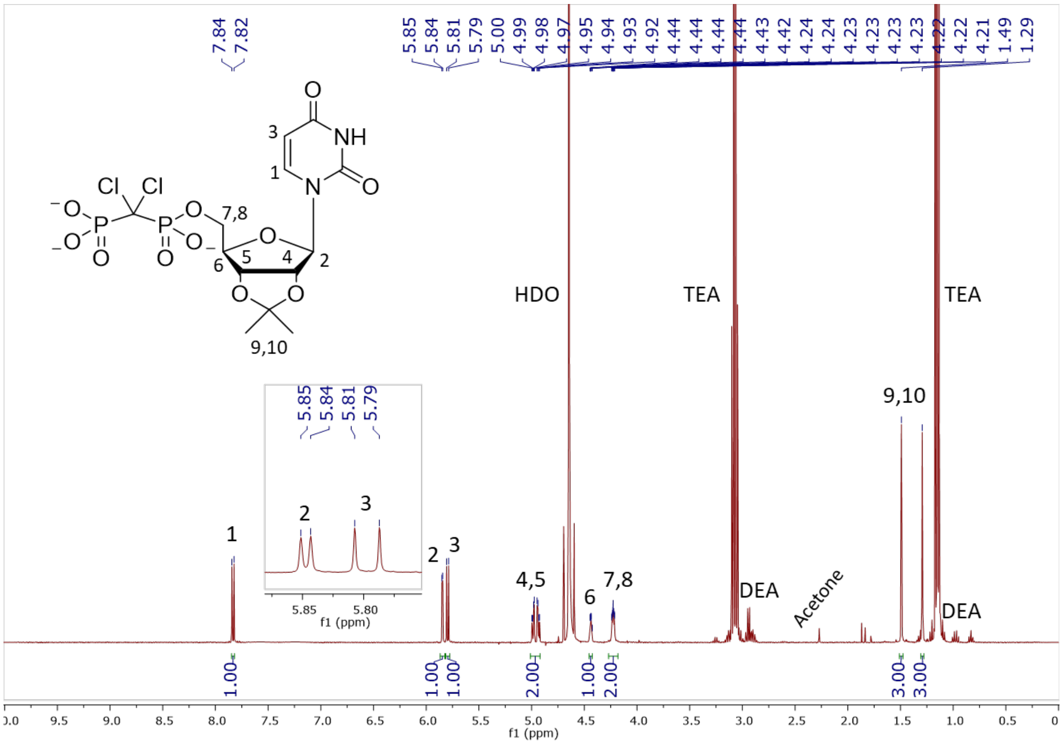

**Figure S37.** $^1H$ NMR (400 MHz, $D_2O$) of **24a**

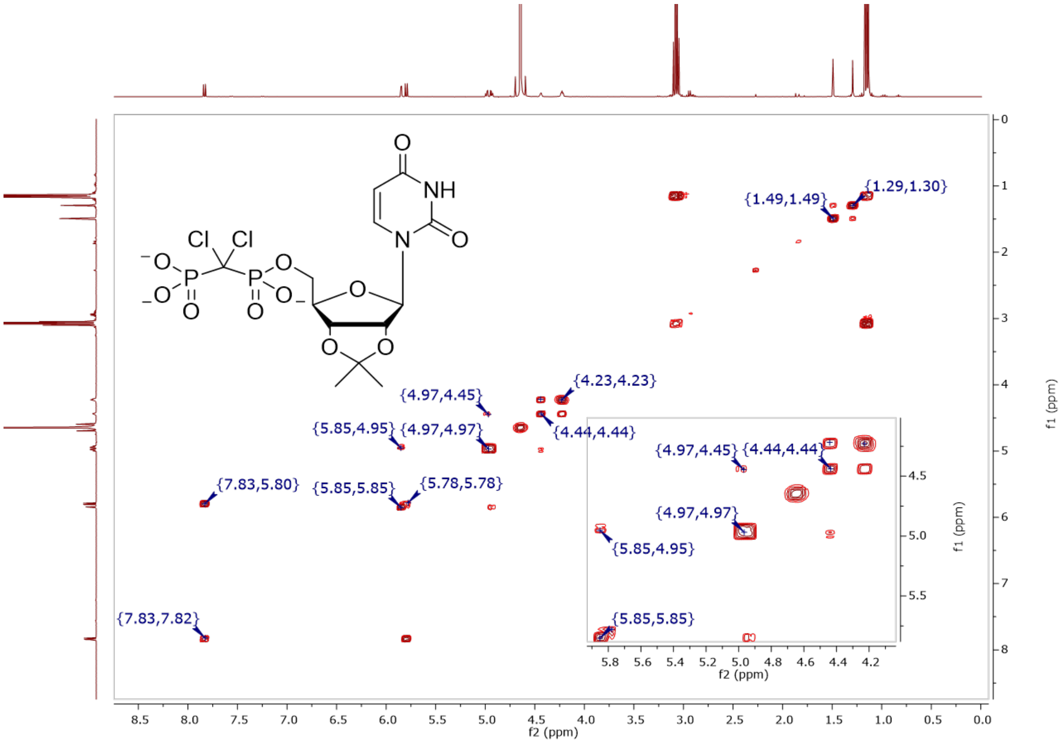

**Figure S38.** COSY (400 MHz, $D_2O$) of **24a**

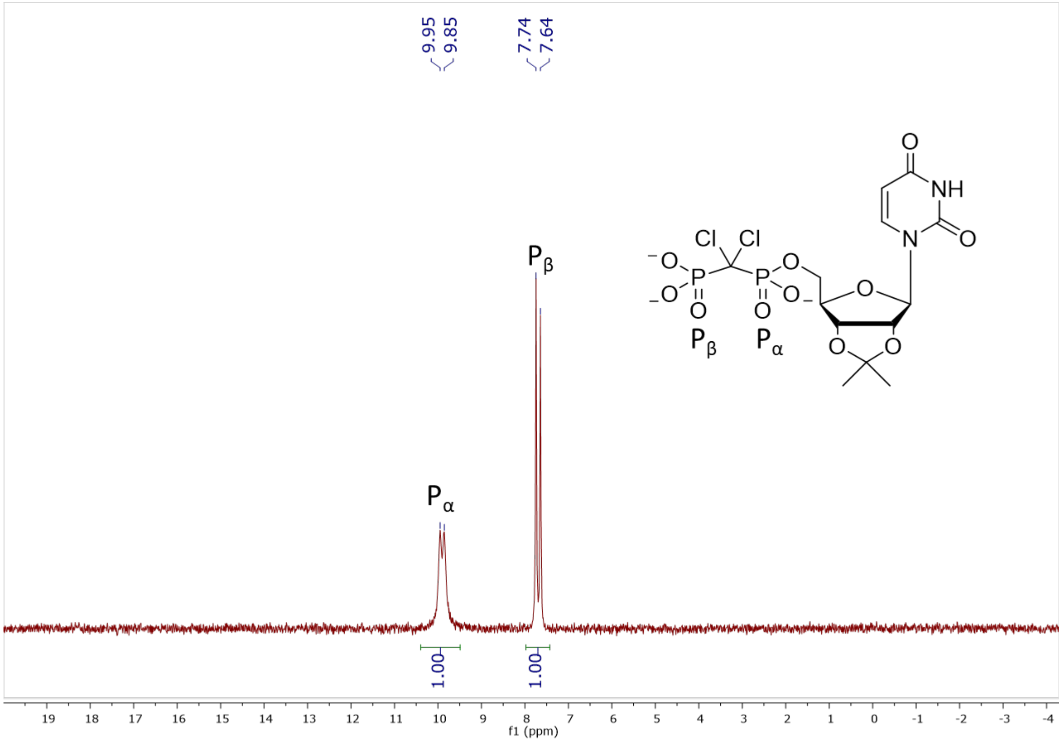

**Figure S39.** $^{31}$P NMR (162 MHz, $D_2O$, pH 10) of **24a**

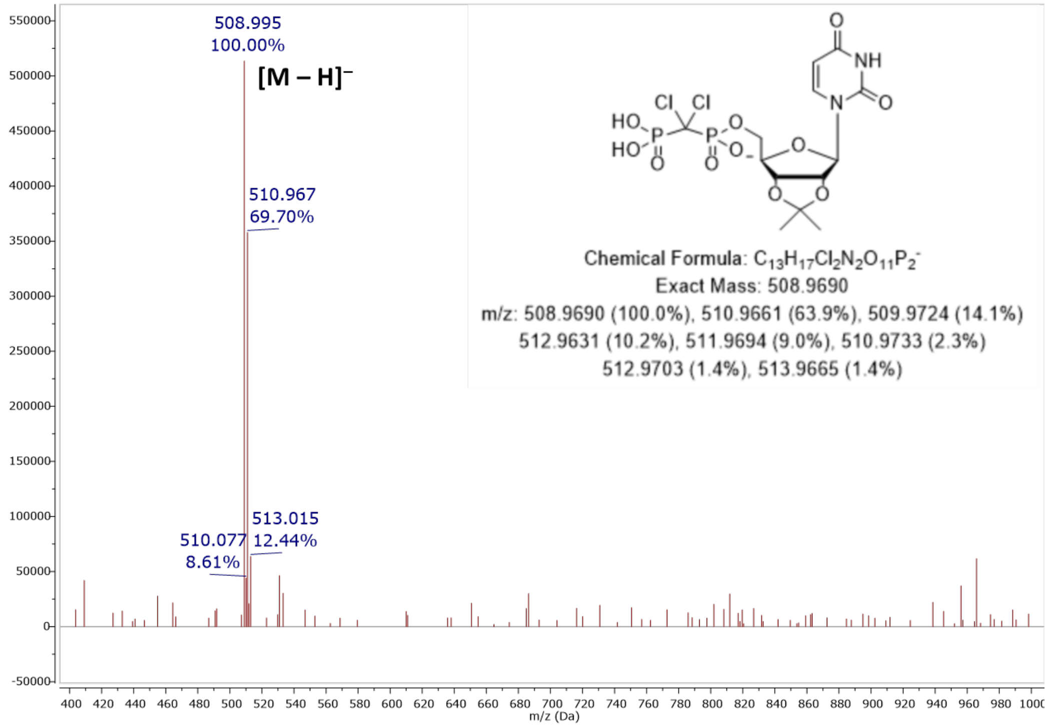

**Figure S40.** MS (ESI): [M – H]$^-$ of **24a**

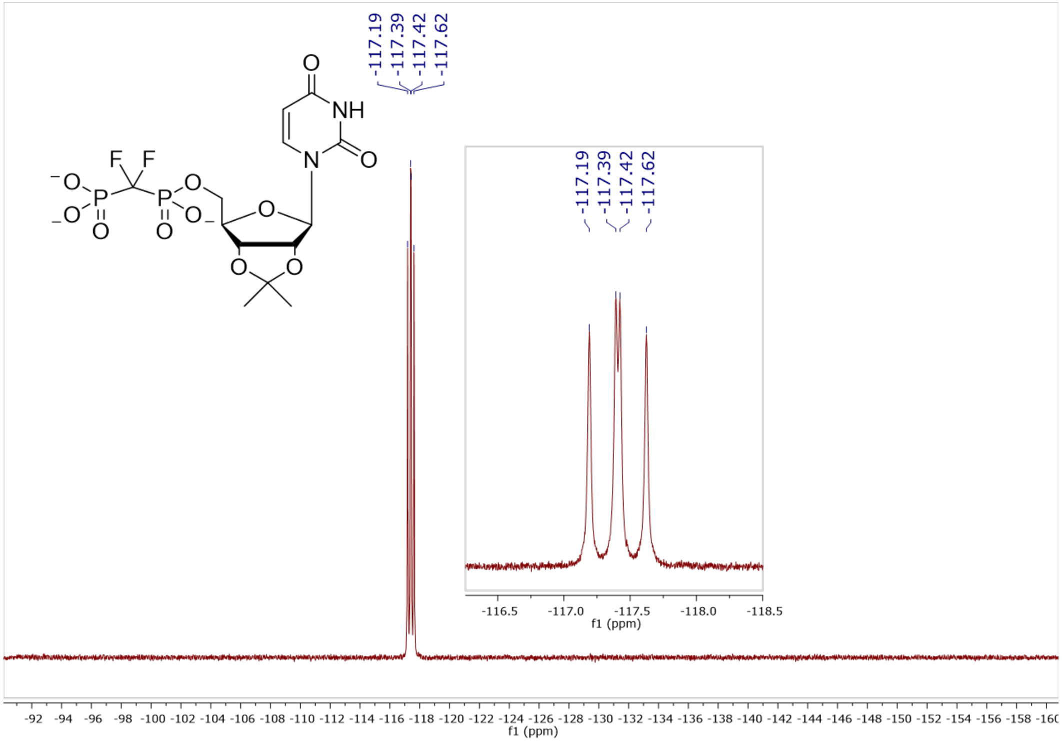

**Figure S41.** $^{19}$F NMR (376 MHz, $D_2O$, pH 7.5) of **24b**

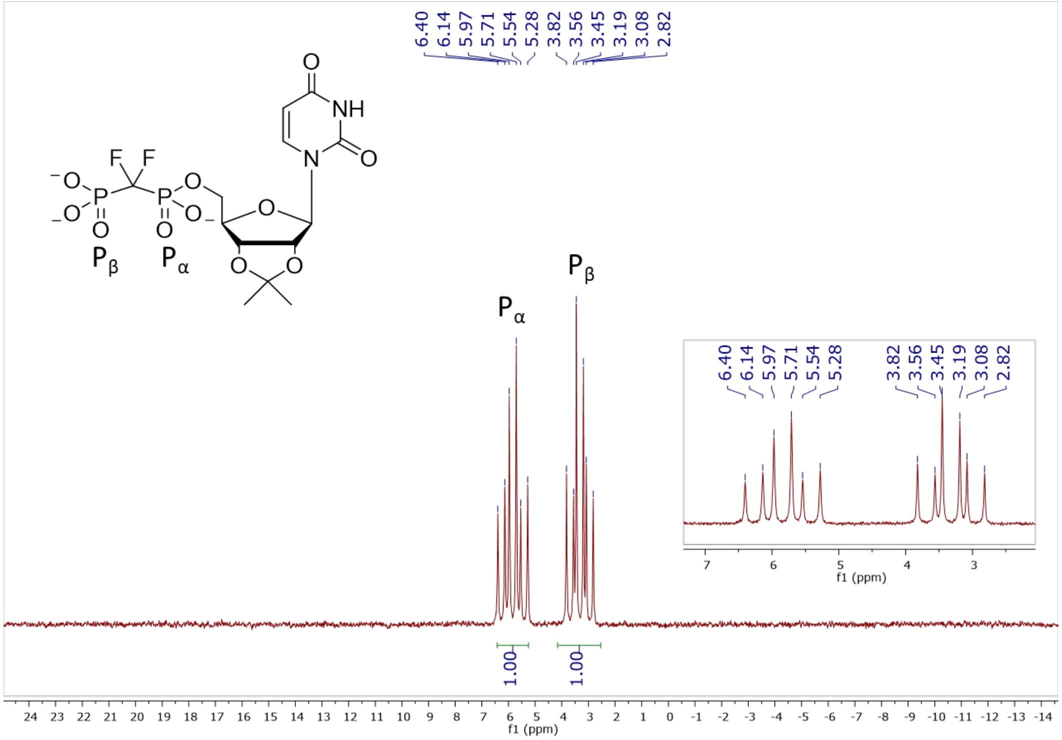

**Figure S42.** $^{31}P$ NMR (202 MHz, $D_2O$, pH 7.5) of **24b**

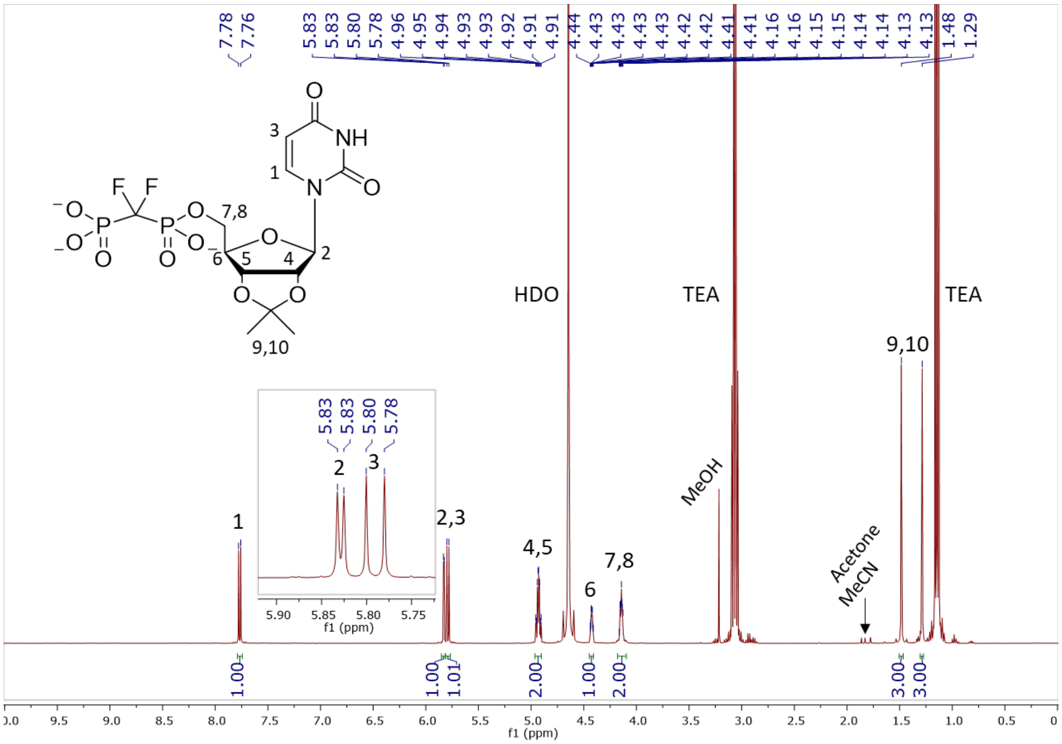

**Figure S43.** $^{1}H$ NMR (400 MHz, $D_2O$) of **24b**

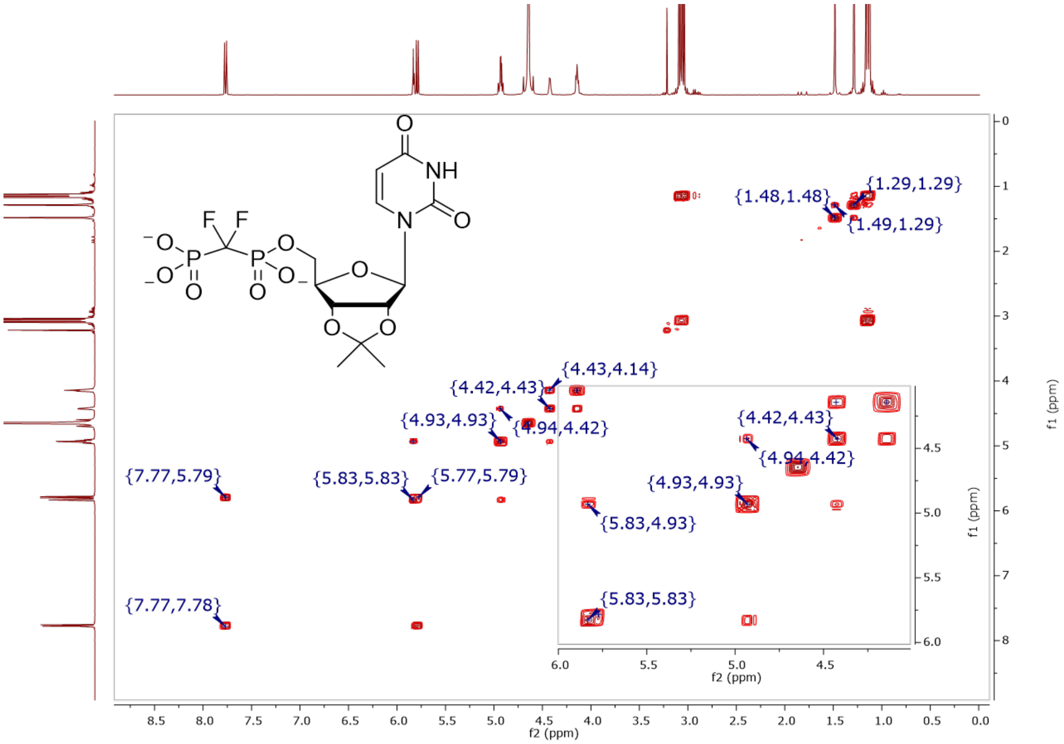

**Figure S44.** COSY (400 MHz, $D_2O$) of **24b**

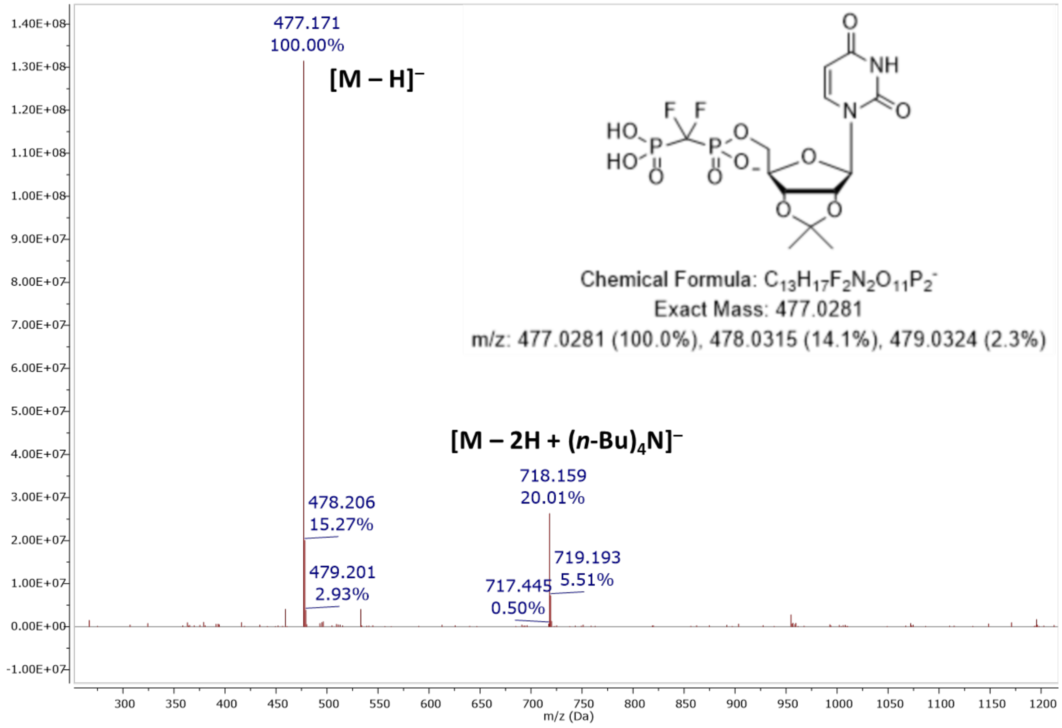

**Figure S45.** MS (ESI): $[M-H]^-$ of **24b**

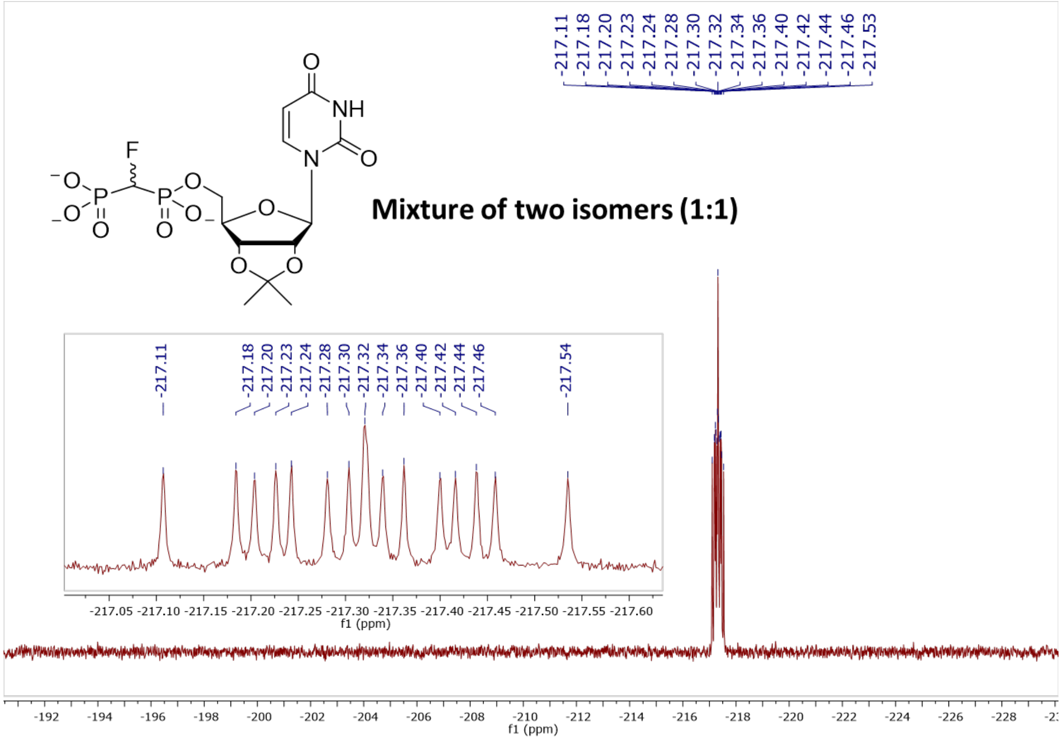

**Figure S46.** $^{19}F$ NMR (470 MHz, $D_2O$, pH 7.5) of **24c**

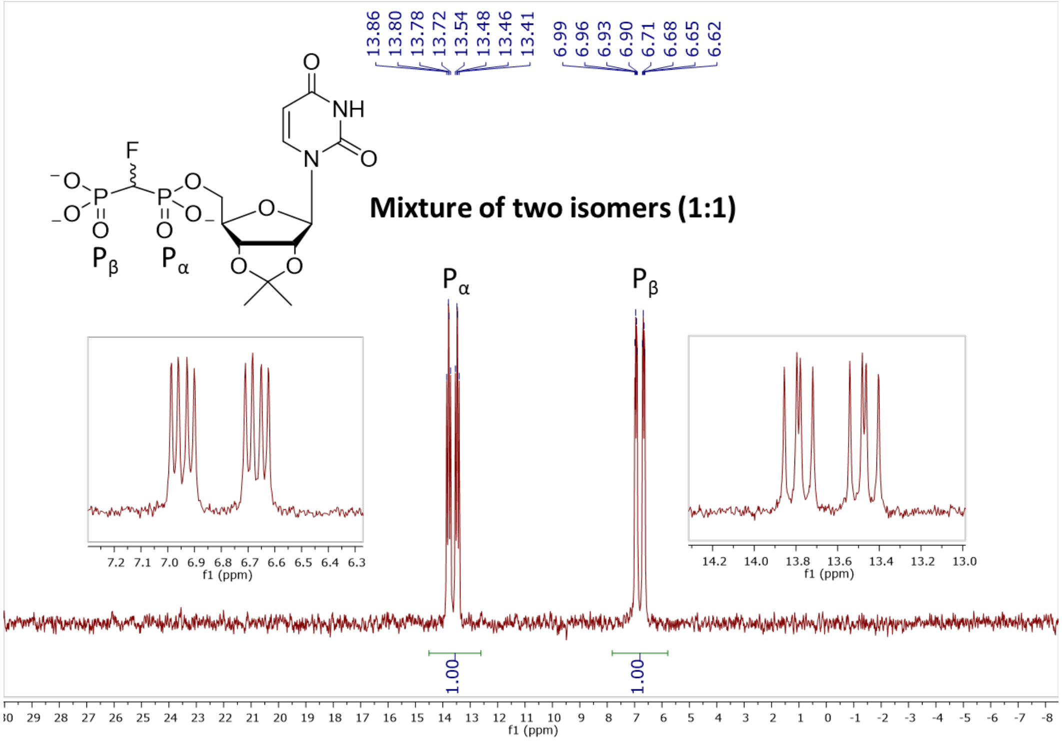

**Figure S47.** $^{31}P$ NMR (202 MHz, $D_2O$, pH 7.5) of **24c**

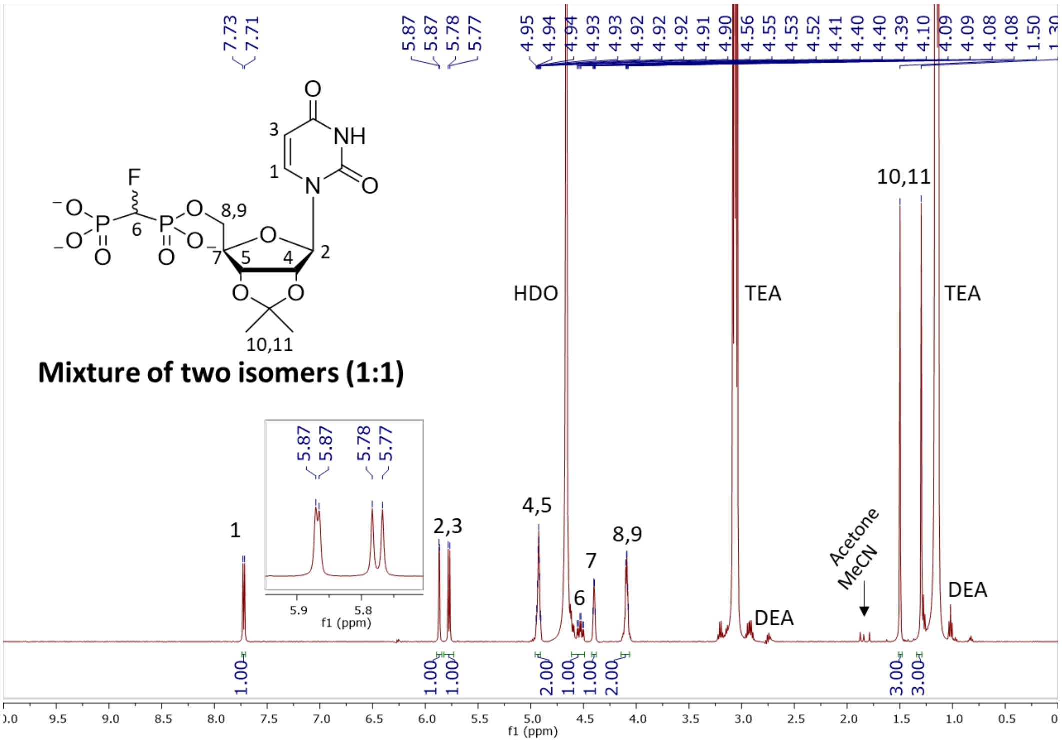

**Figure S48.** $^{1}H$ NMR (500 MHz, $D_2O$) of **24c**

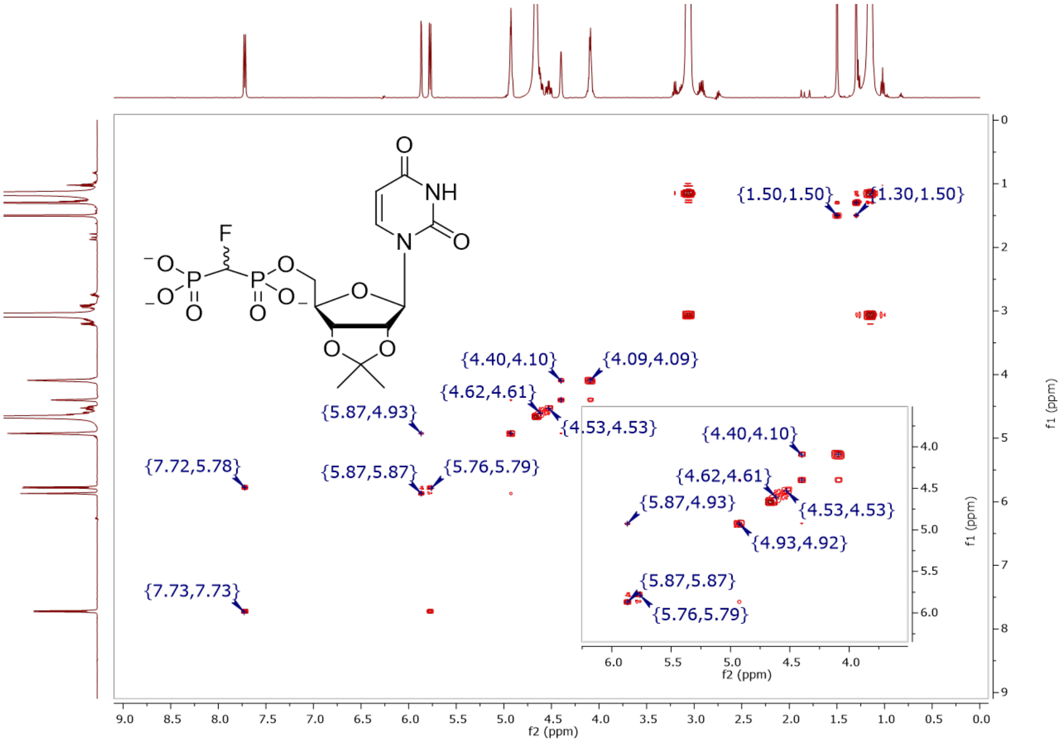

**Figure S49.** COSY (500 MHz, $D_2O$) of **24c**

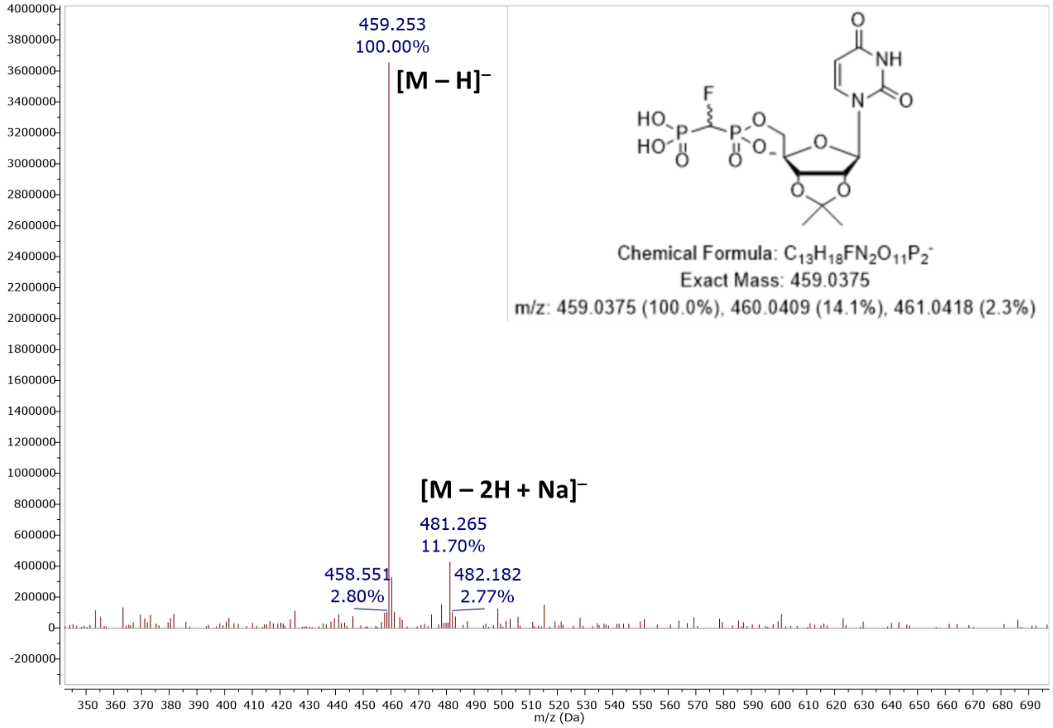

**Figure S50.** MS (ESI): [M – H]$^-$ of **24c**

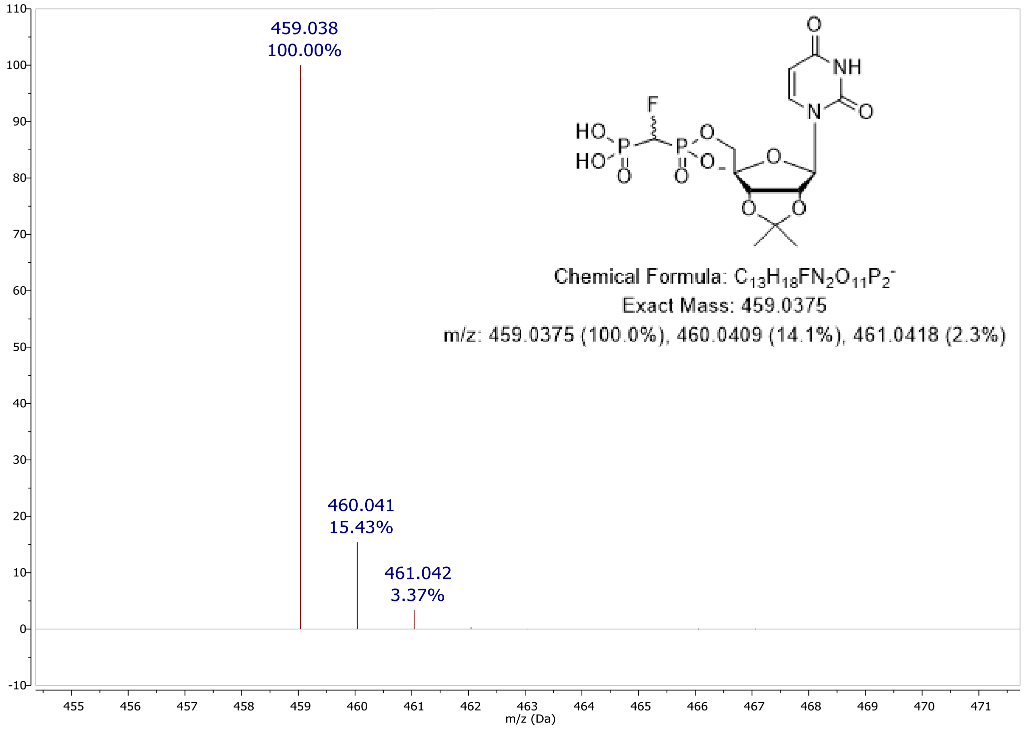

**Figure S51.** Predicted MS: [M – H]$^-$ of **24c**

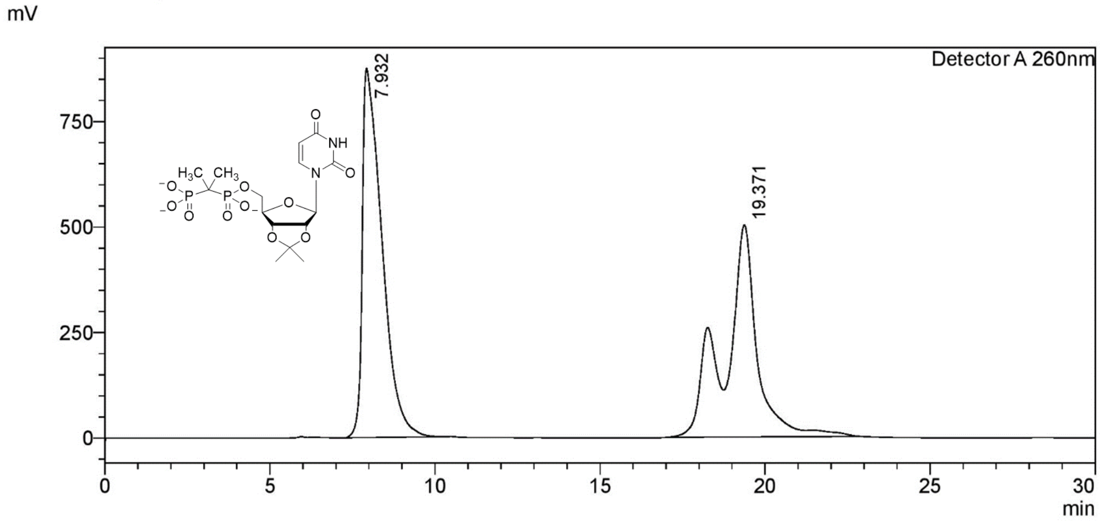

**Figure S52.** Preparative RP-HPLC purification of **24d**

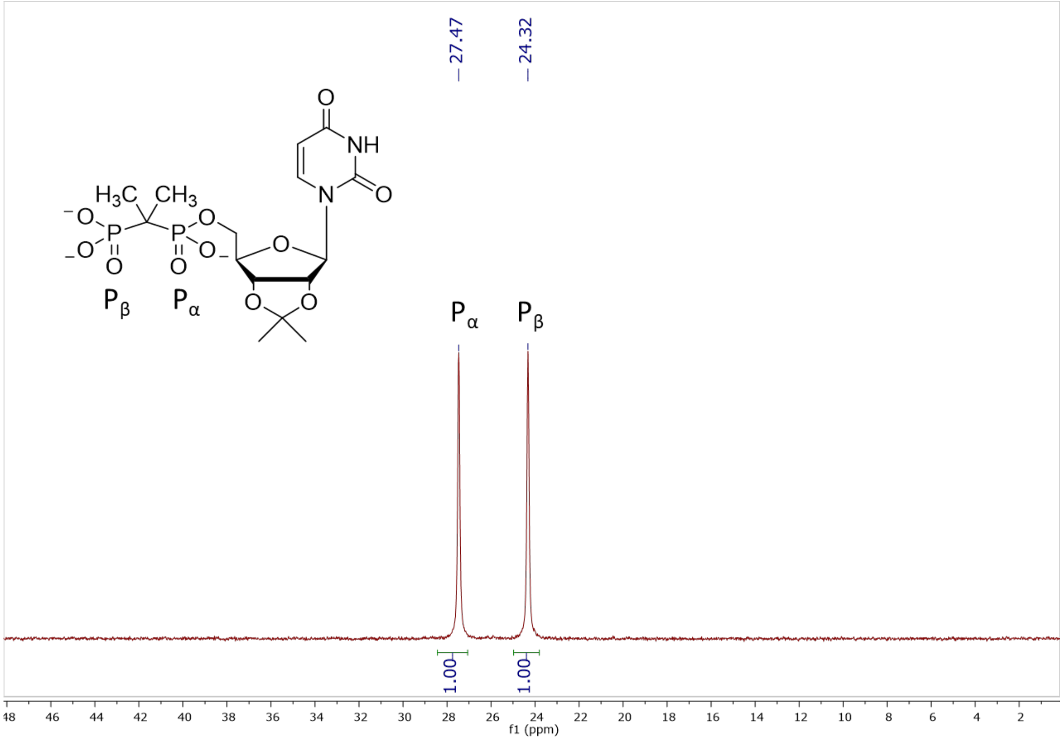

**Figure S53.** $^{31}$P NMR (202 MHz, $D_2O$, pH 7.5) of **24d**

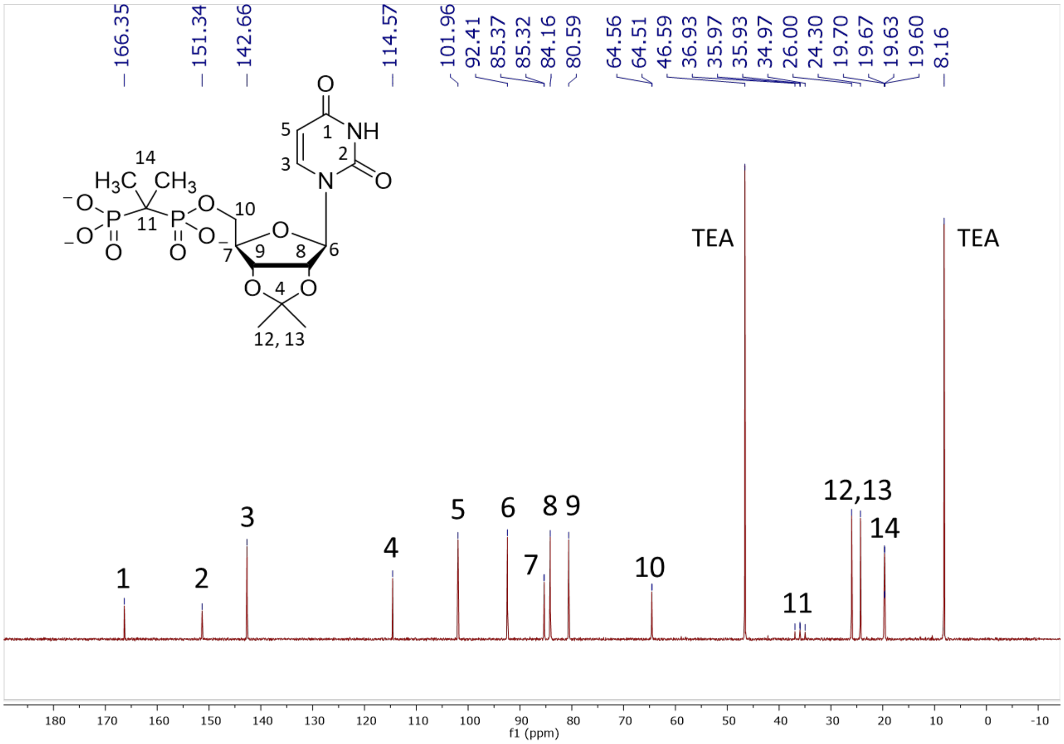

**Figure S54.** $^{13}C$ NMR (126 MHz, $D_2O$) of **24d**

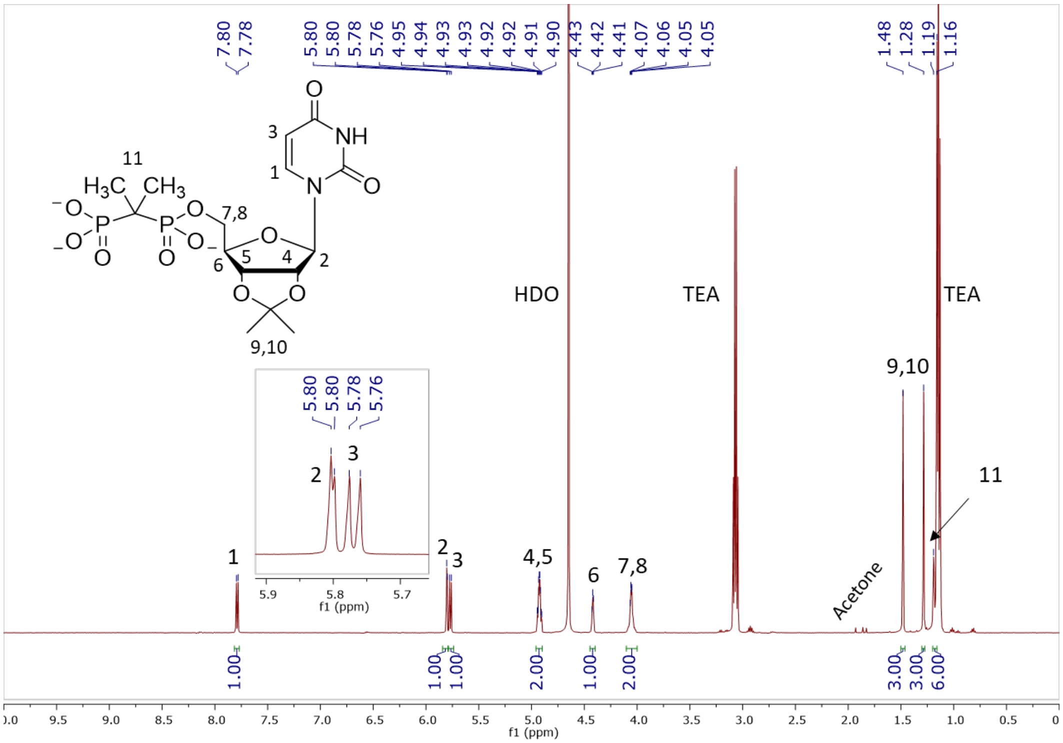

**Figure S55.** $^{1}H$ NMR (500 MHz, $D_2O$) of **24d**

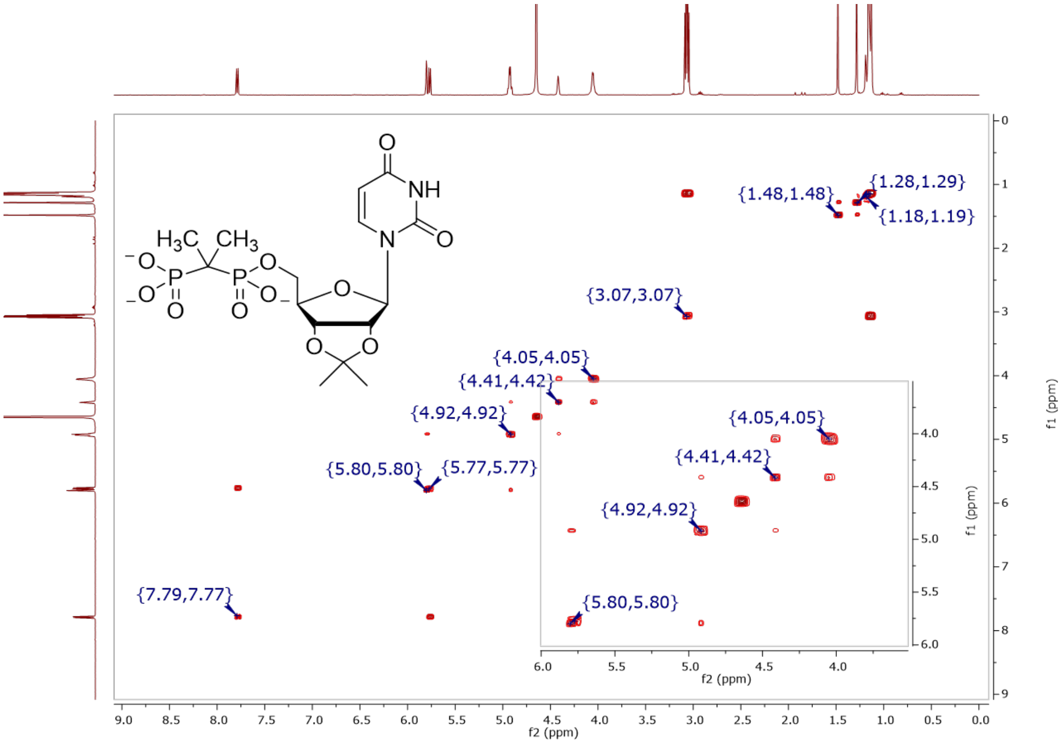

**Figure S56.** COSY (500 MHz, $D_2O$) of **24d**

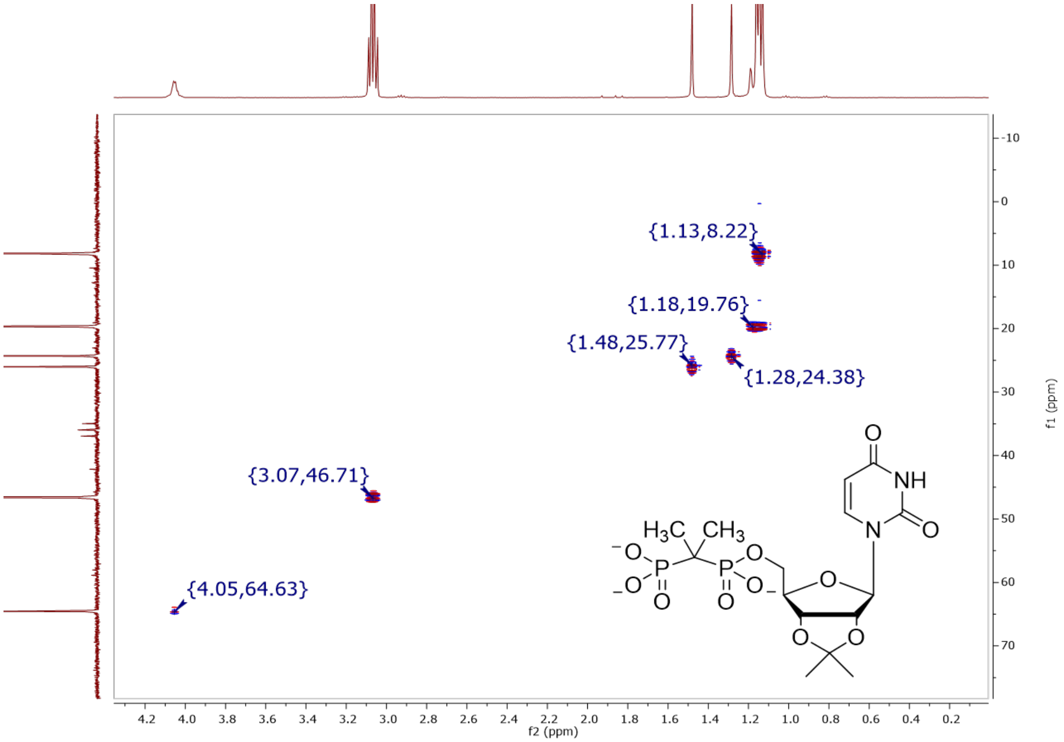

**Figure S57.** HSQCAD (500, 126 MHz, $D_2O$) of **24d**

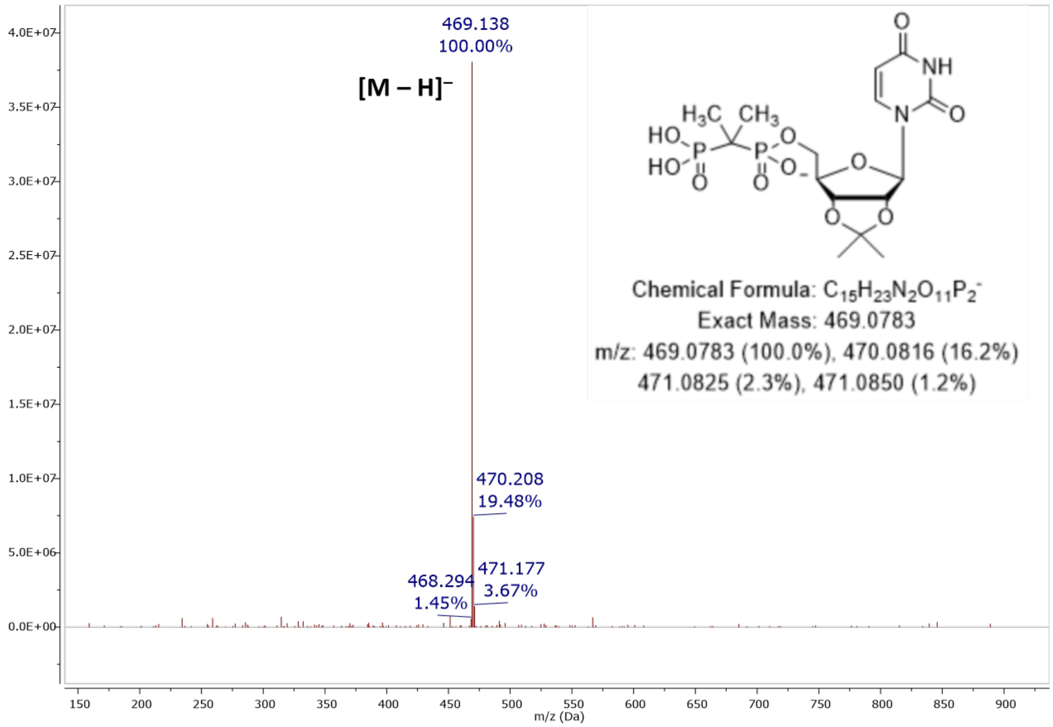

**Figure S58.** MS (ESI): [M – H]⁻ of **24d**

#### Elemental Composition Report

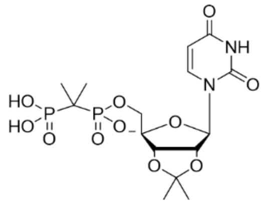

##### Single Mass Analysis

Tolerance = 5.0 PPM / DBE: min = -1.5, max = 50.0
Element prediction: Off
Number of isotope peaks used for i-FIT = 3

Monoisotopic Mass, Even Electron Ions
64 formula(e) evaluated with 1 results within limits (up to 50 closest results for each mass)
Elements Used:
C: 0-60 H: 0-80 N: 0-5 O: 10-12 P: 2-2

Pouya Haratipour USCMCK25
Synapt2_995 24 (0.482) Cm (21:24-5:11)

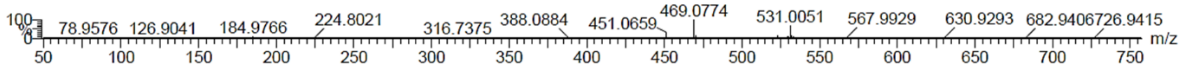

| | | | | | | | | |
|---|---|---|---|---|---|---|---|---|
| Minimum: | | | | -1.5 | | | | |
| Maximum: | | 5.0 | 5.0 | 50.0 | | | | |
| Mass | Calc. Mass | mDa | PPM | DBE | i-FIT | Norm | Conf(%) | Formula |
| 469.0774 | 469.0777 | -0.3 | -0.6 | 6.5 | 1045.7 | n/a | n/a | C15 H23 N2 O11 P2 |

**Figure S59.** Elemental composition report from HRMS (ESI-TOF) [M – H]⁻ of **24d**

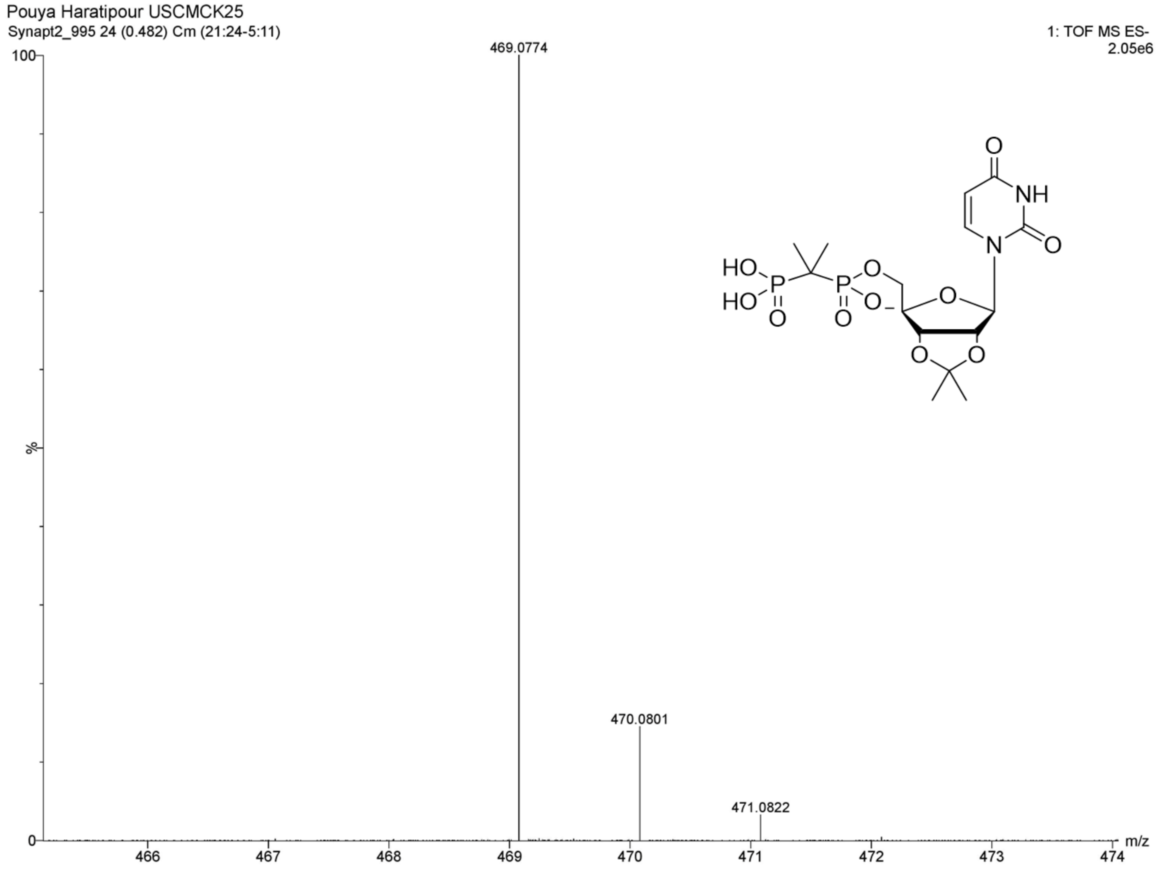

**Figure S60.** HRMS (ESI-TOF): $[M - H]^-$ of **24d**

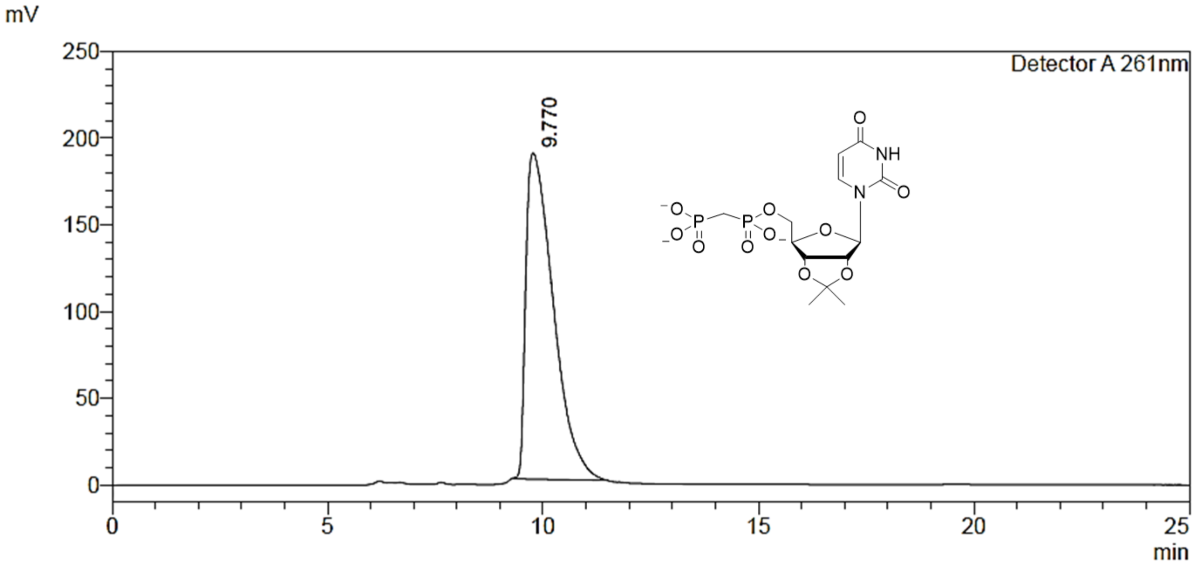

**Figure S61.** Preparative RP-HPLC purification of **24e**

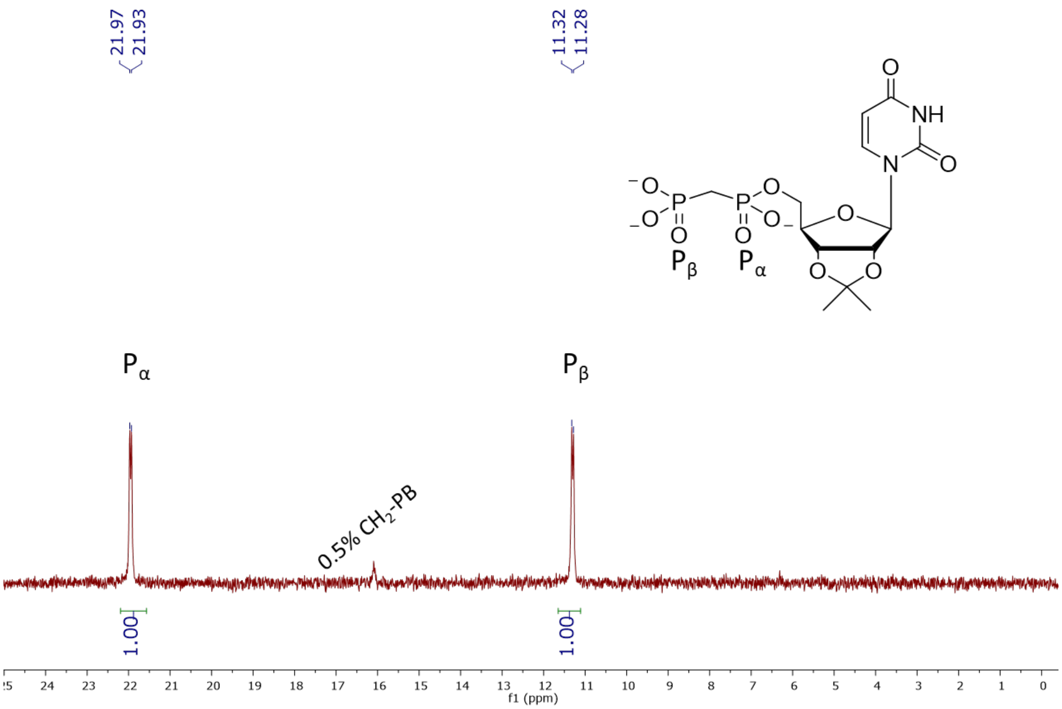

**Figure S62.** $^{31}$P NMR (202 MHz, $D_2O$, pH 7.5) of **24e**

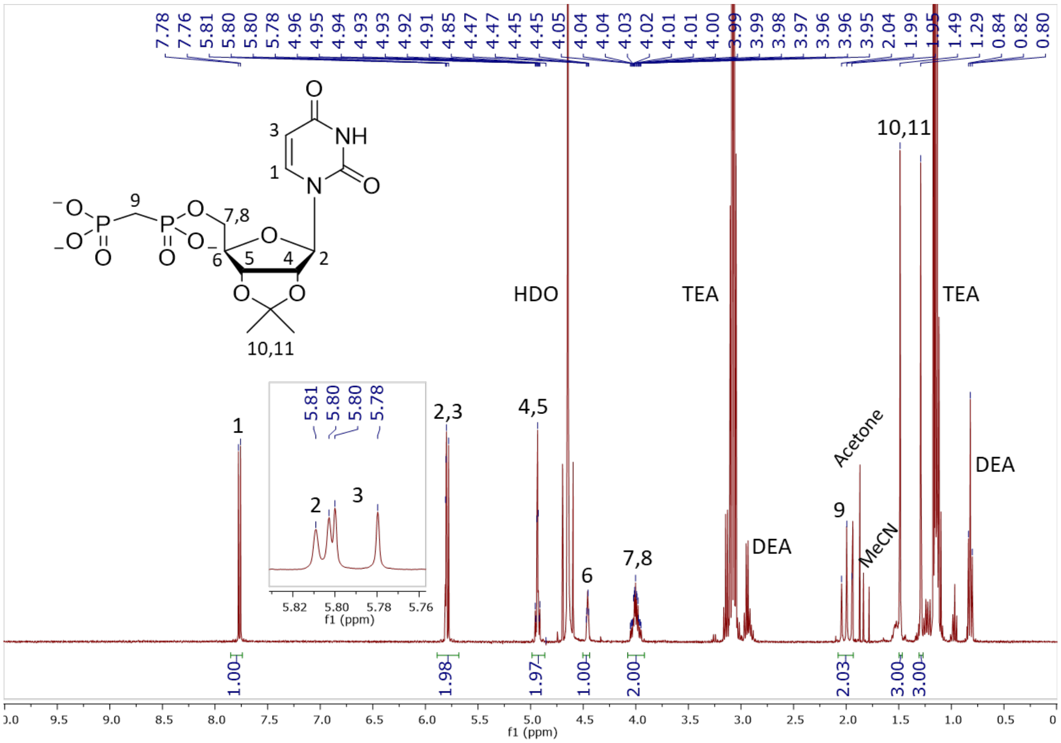

**Figure S63.** $^{1}$H NMR (400 MHz, $D_2O$) of **24e**

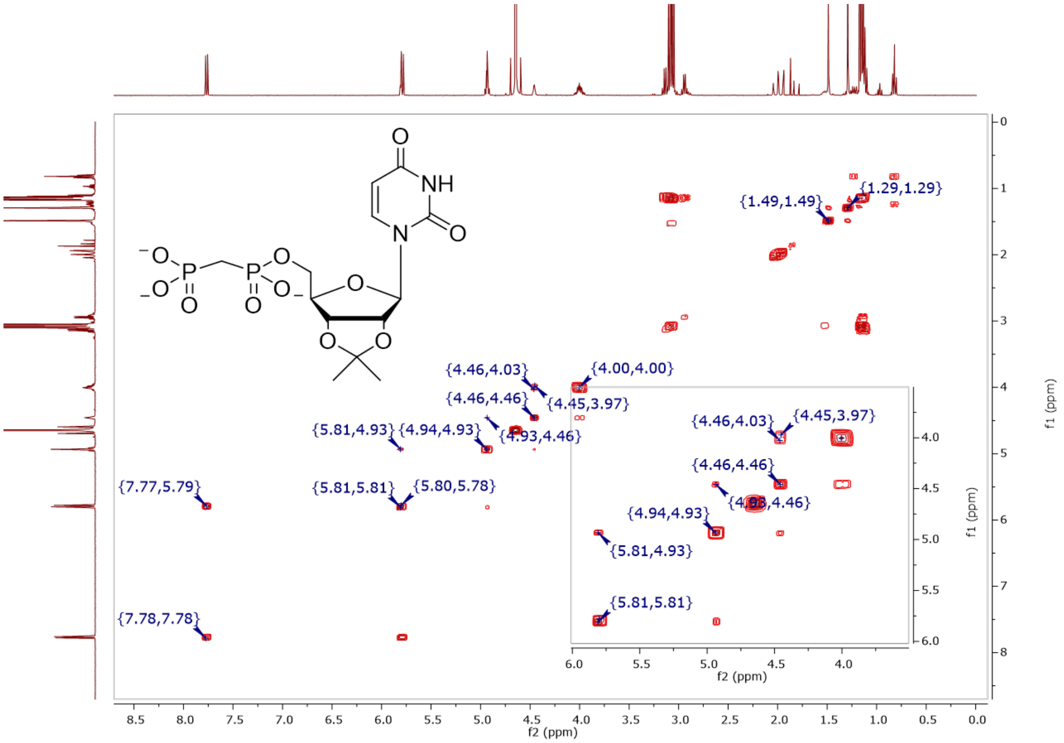

**Figure S64.** COSY (400 MHz, $D_2O$) of **24e**

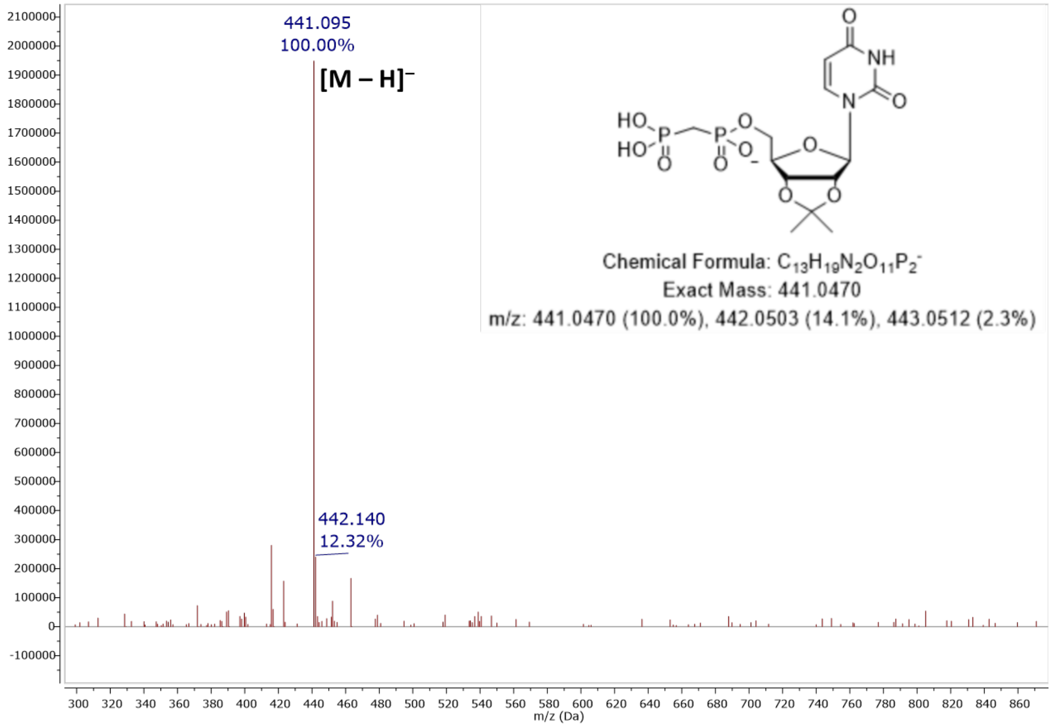

**Figure S65.** MS (ESI): $[M – H]^–$ of **24e**

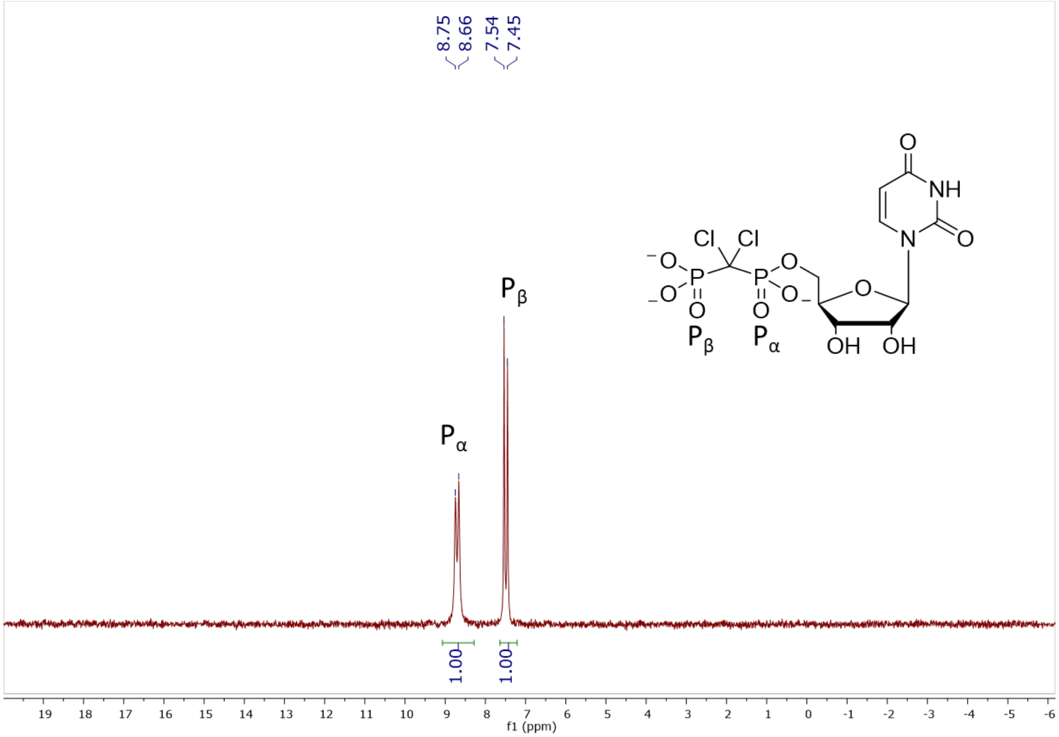

**Figure S66.** $^{31}P$ NMR (202 MHz, $D_2O$, pH 7.5) of **1**

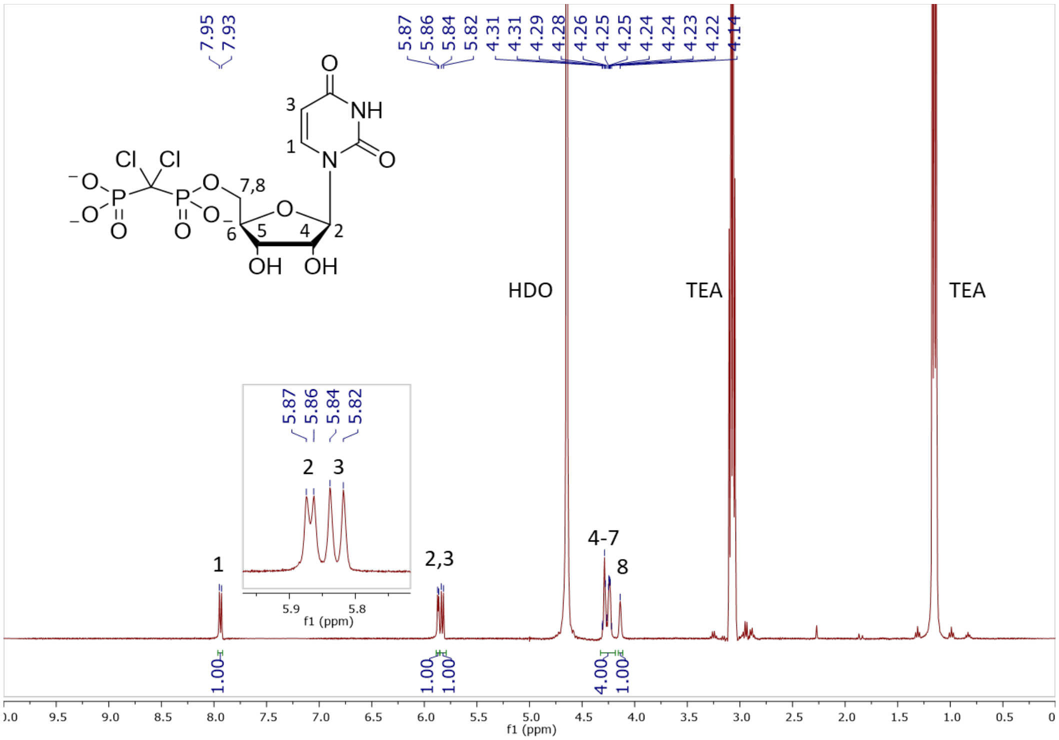

**Figure S67.** $^{1}H$ NMR (400 MHz, $D_2O$) of **1**

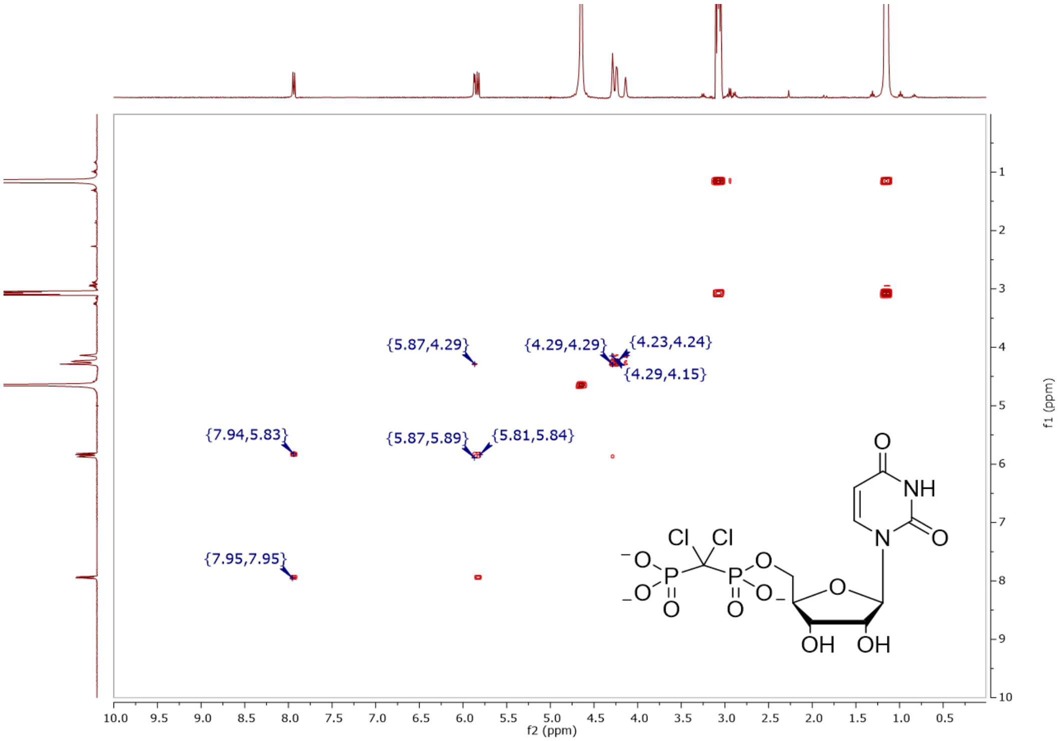

**Figure S68.** COSY (400 MHz, $D_2O$) of **1**

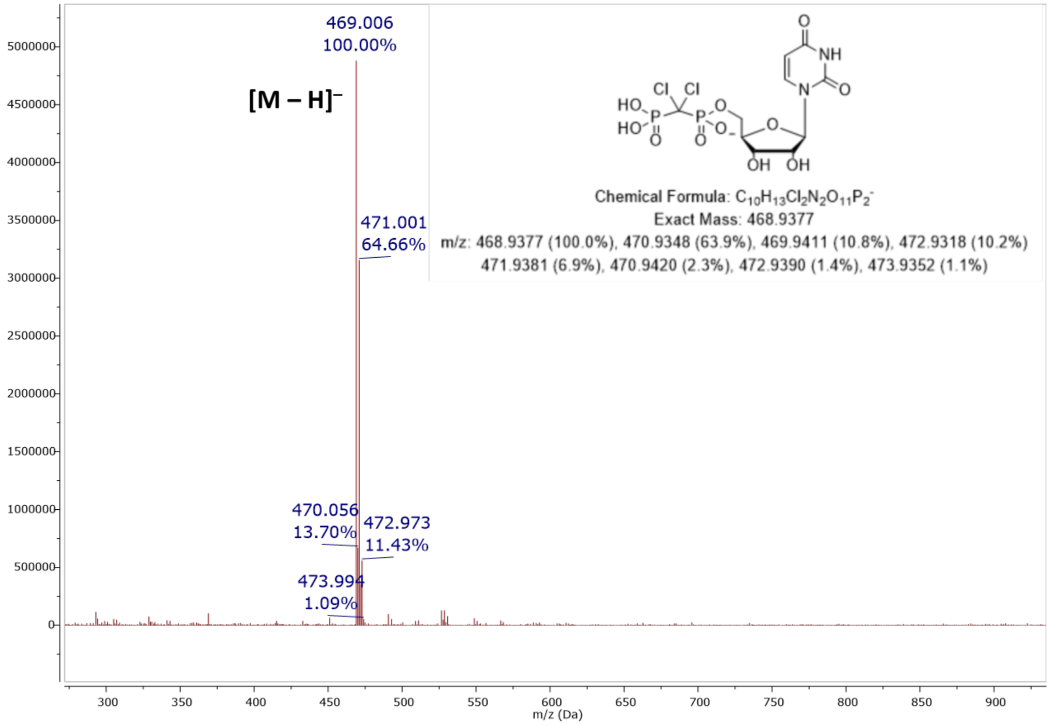

**Figure S69.** MS (ESI): $[M - H]^-$ of **1**

#### Elemental Composition Report

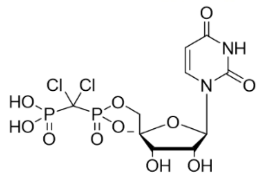

##### Single Mass Analysis

Tolerance = 5.0 PPM / DBE: min = -1.5, max = 100.0
Element prediction: Off
Number of isotope peaks used for i-FIT = 3

Monoisotopic Mass, Even Electron Ions
8 formula(e) evaluated with 1 results within limits (up to 50 closest results for each mass)
Elements Used:
C: 0-100 H: 0-100 N: 2-2 O: 10-12 P: 2-2 Cl: 2-2
Pouya Haratipour, USCMCK22
Synapt_26483b 53 (1.051) Cm (53-24)

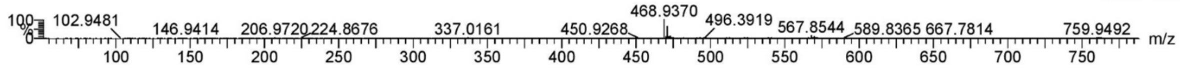

| | | | | | | | | |
|---|---|---|---|---|---|---|---|---|
| Minimum: | | | | -1.5 | | | | |
| Maximum: | | 5.0 | 5.0 | 100.0 | | | | |
| Mass | Calc. Mass | mDa | PPM | DBE | i-FIT | Norm | Conf(%) | Formula |
| 468.9370 | 468.9372 | -0.2 | -0.4 | 5.5 | 521.9 | n/a | n/a | C10 H13 N2 O11 P2 Cl2 |

**Figure S70.** Elemental composition report from HRMS (ESI-TOF) $[M - H]^-$ of **1**

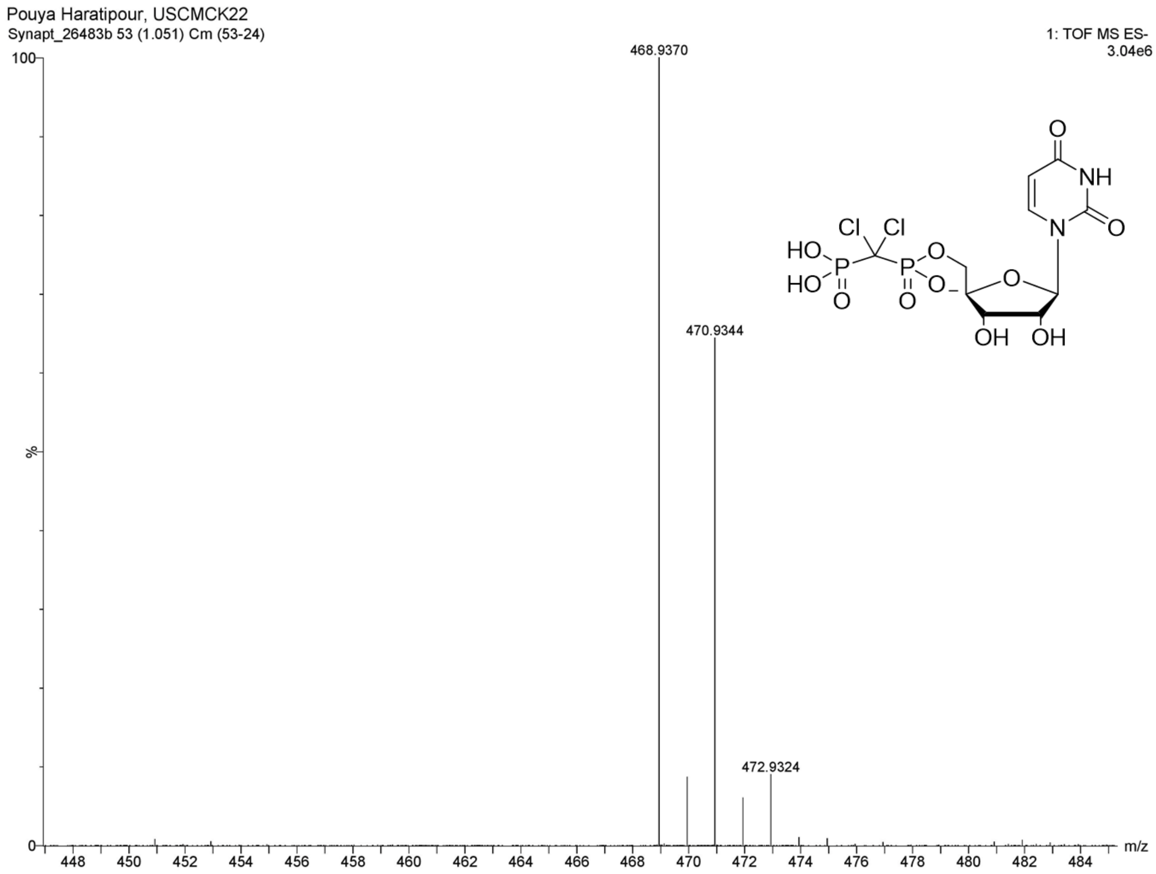

**Figure S71.** HRMS (ESI-TOF): $[M - H]^-$ of **1**

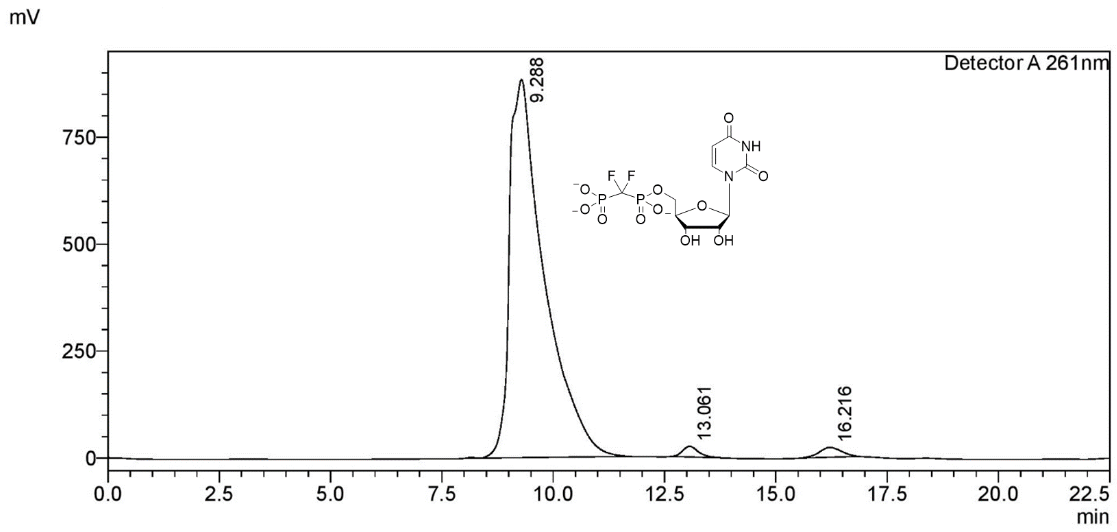

**Figure S72.** Preparative RP-HPLC purification of **3**

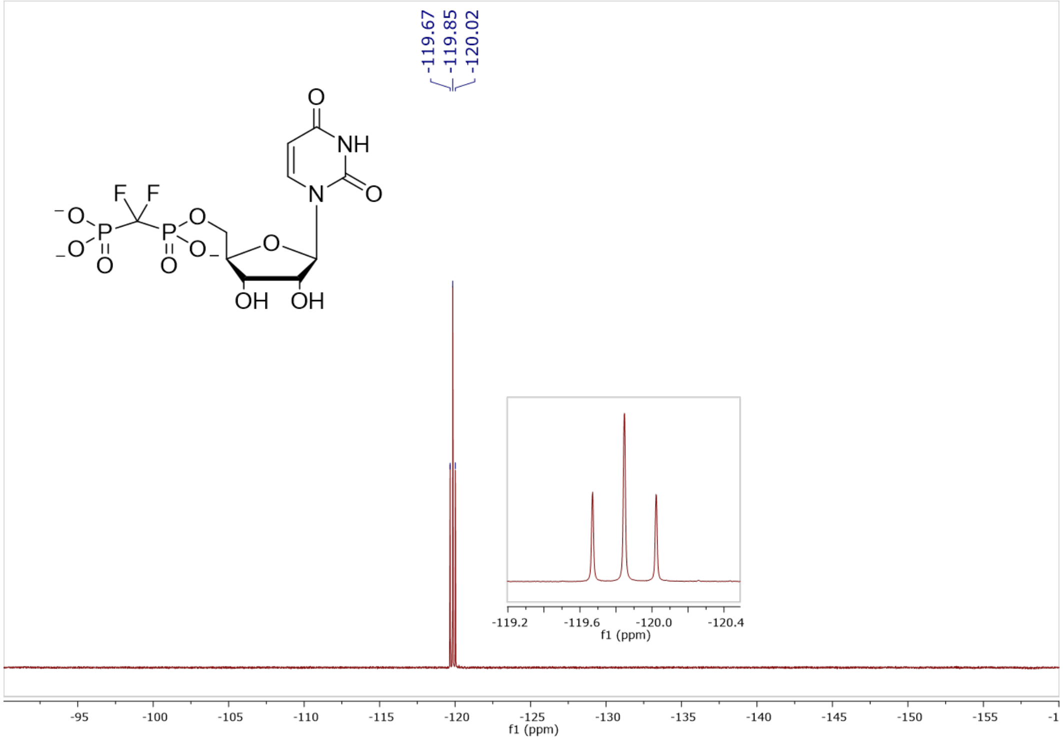

**Figure S73.** [19]F NMR (470 MHz, $D_2O$, pH 7.5) of **3**

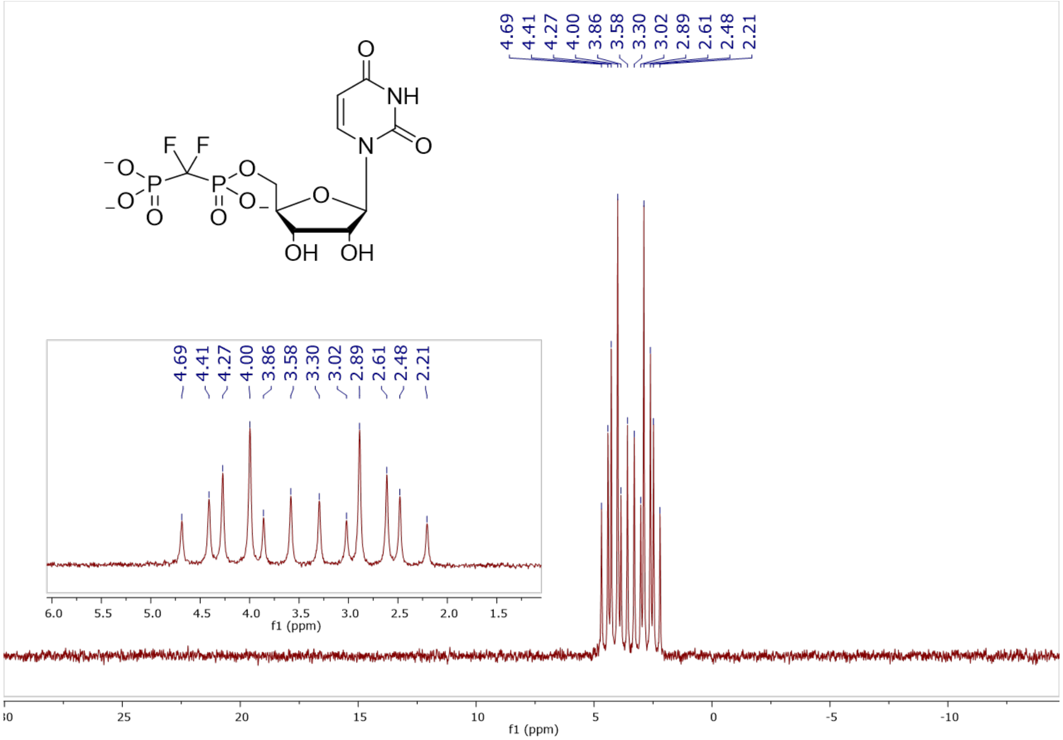

**Figure S74.** $^{31}P$ NMR (202 MHz, $D_2O$, pH 7.5) of **3**

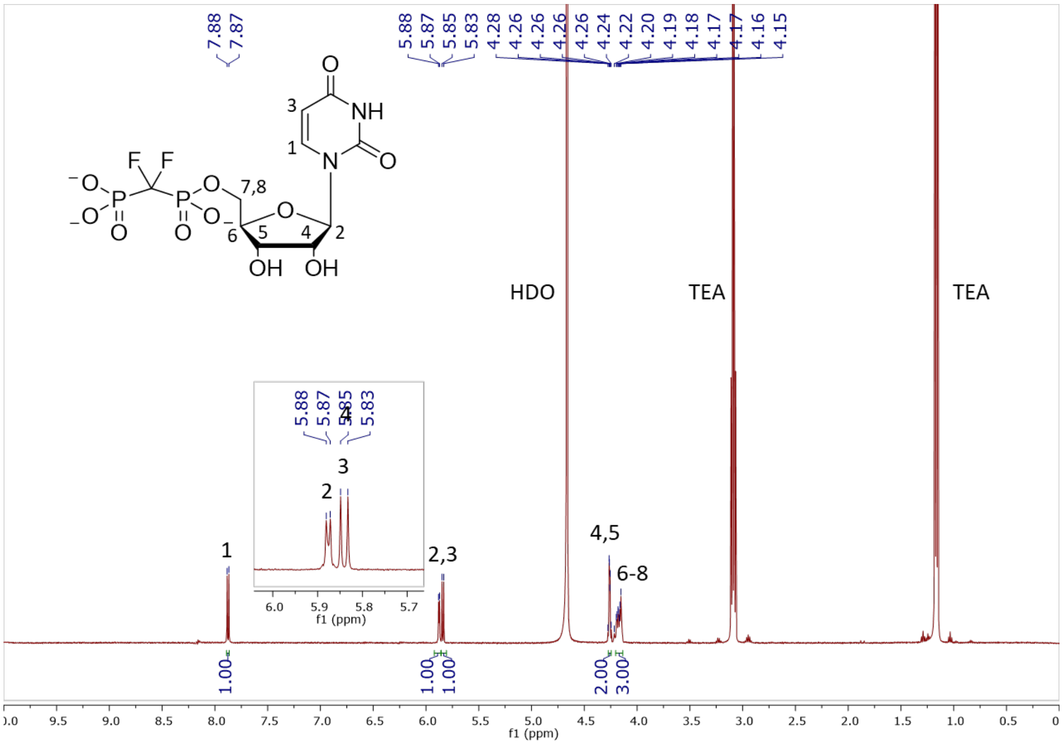

**Figure S75.** $^{1}H$ NMR (500 MHz, $D_2O$) of **3**

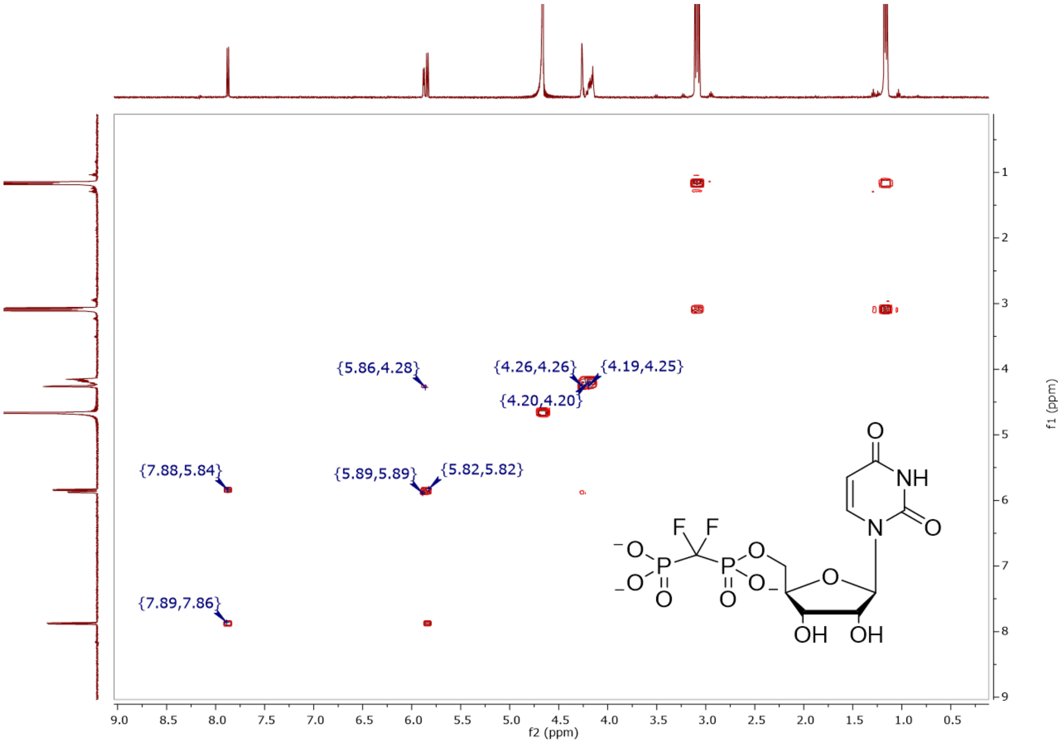

**Figure S76.** COSY (500 MHz, $D_2O$) of **3**

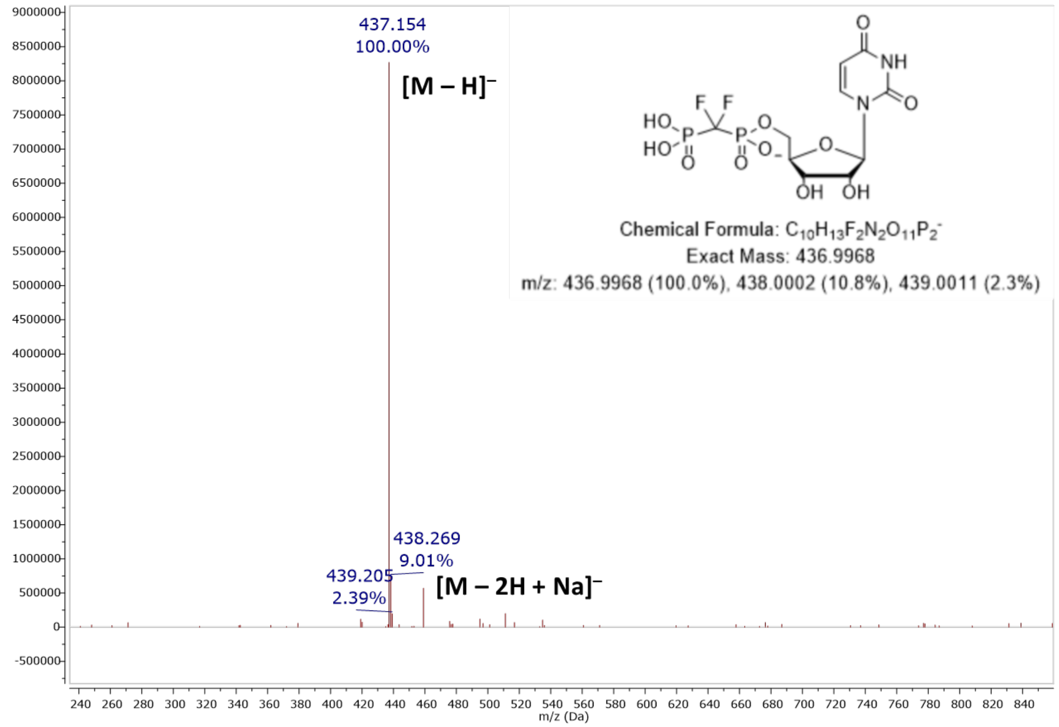

**Figure S77.** MS (ESI): $[M – H]^-$ of **3**

#### Elemental Composition Report

##### Single Mass Analysis

Tolerance = 5.0 PPM / DBE: min = -1.5, max = 50.0
Element prediction: Off
Number of isotope peaks used for i-FIT = 3

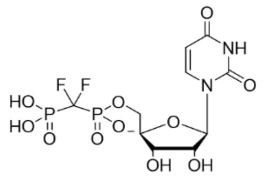

Monoisotopic Mass, Even Electron Ions
74 formula(e) evaluated with 1 results within limits (up to 50 closest results for each mass)
Elements Used:
C: 0-60 H: 0-100 N: 1-3 O: 10-12 P: 1-3 F: 2-2

Pouya Haratipour, USCMCK13
Synapt_26238 27 (0.552) Cm (27:29-5:7)

1: TOF MS ES-
2.33e+007

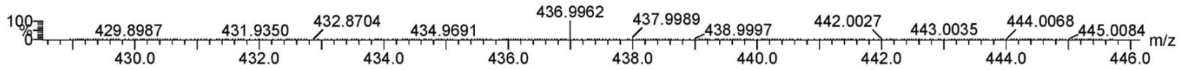

| | | | | | | | | |
|---|---|---|---|---|---|---|---|---|
| Minimum: | | | | -1.5 | | | | |
| Maximum: | | 5.0 | 5.0 | 50.0 | | | | |
| Mass | Calc. Mass | mDa | PPM | DBE | i-FIT | Norm | Conf(%) | Formula |
| 436.9962 | 436.9963 | -0.1 | -0.2 | 5.5 | 745.8 | n/a | n/a | C10 H13 N2 O11 P2 F2 |

**Figure S78.** Elemental composition report from HRMS (ESI-TOF) $[M - H]^-$ of **3**

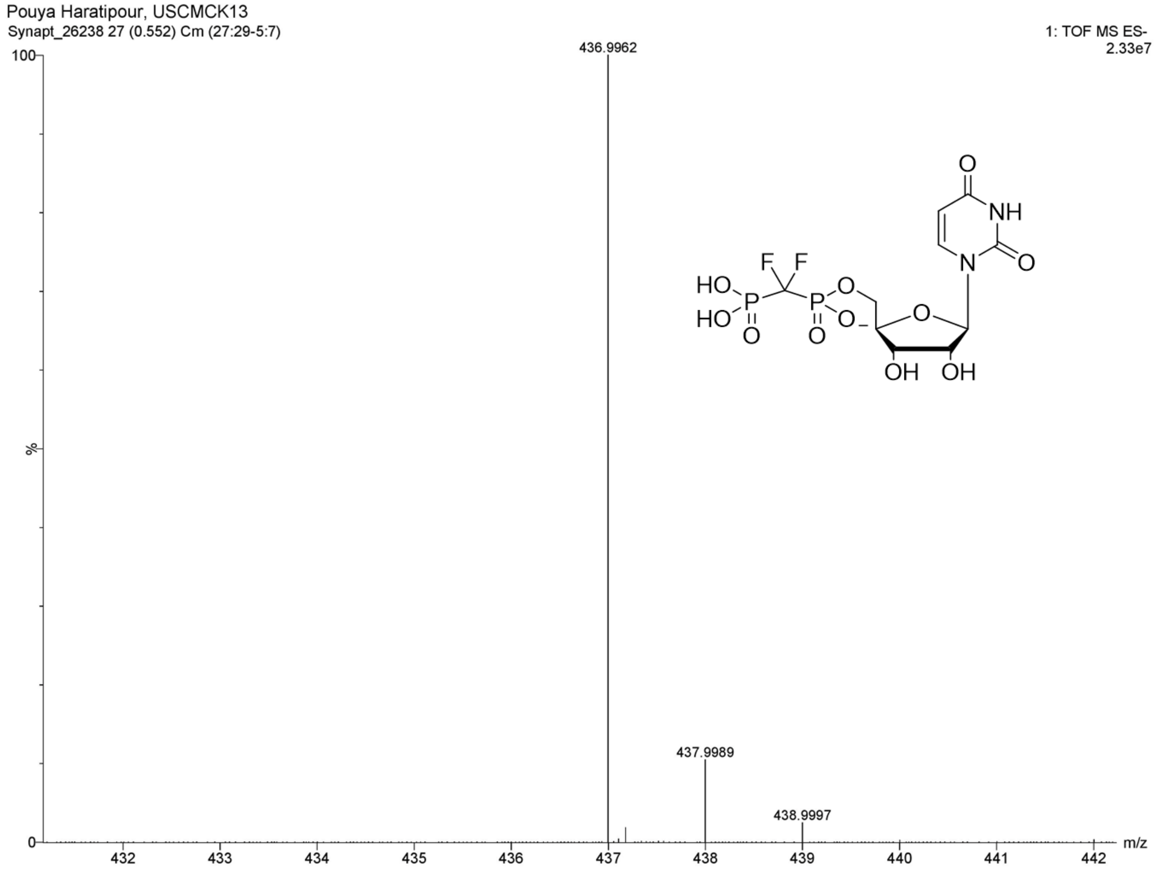

**Figure S79.** HRMS (ESI-TOF): $[M - H]^-$ of **3**

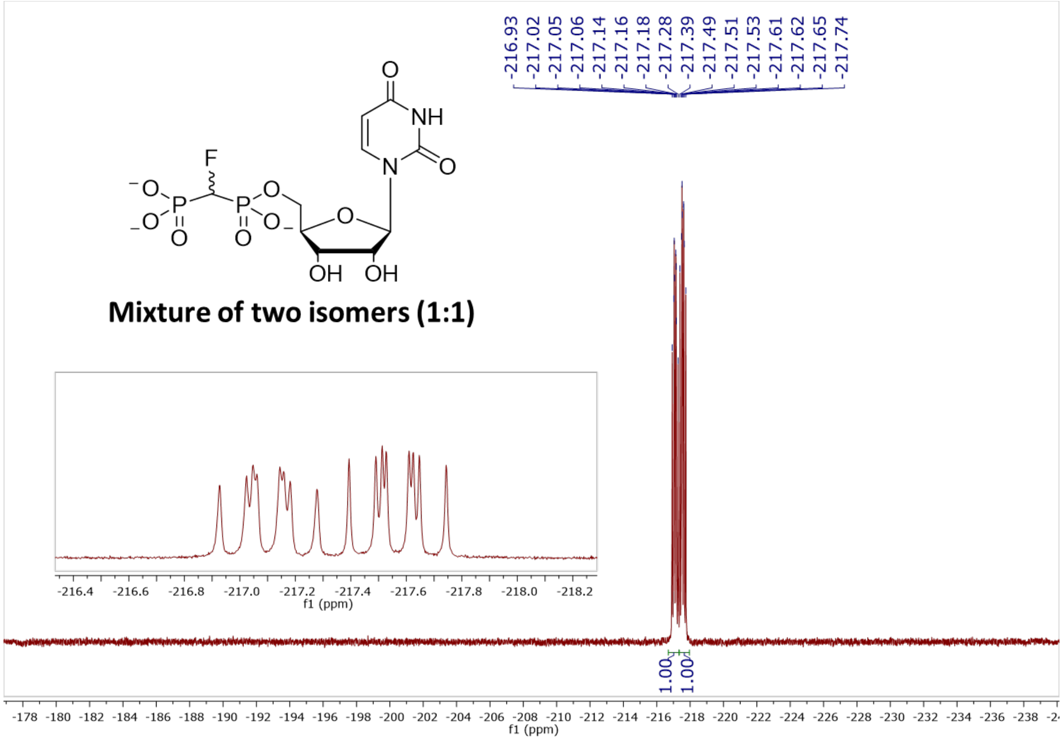

**Figure S80.** $^{19}$F NMR (470 MHz, $D_2O$, pH 7.5) of **6**

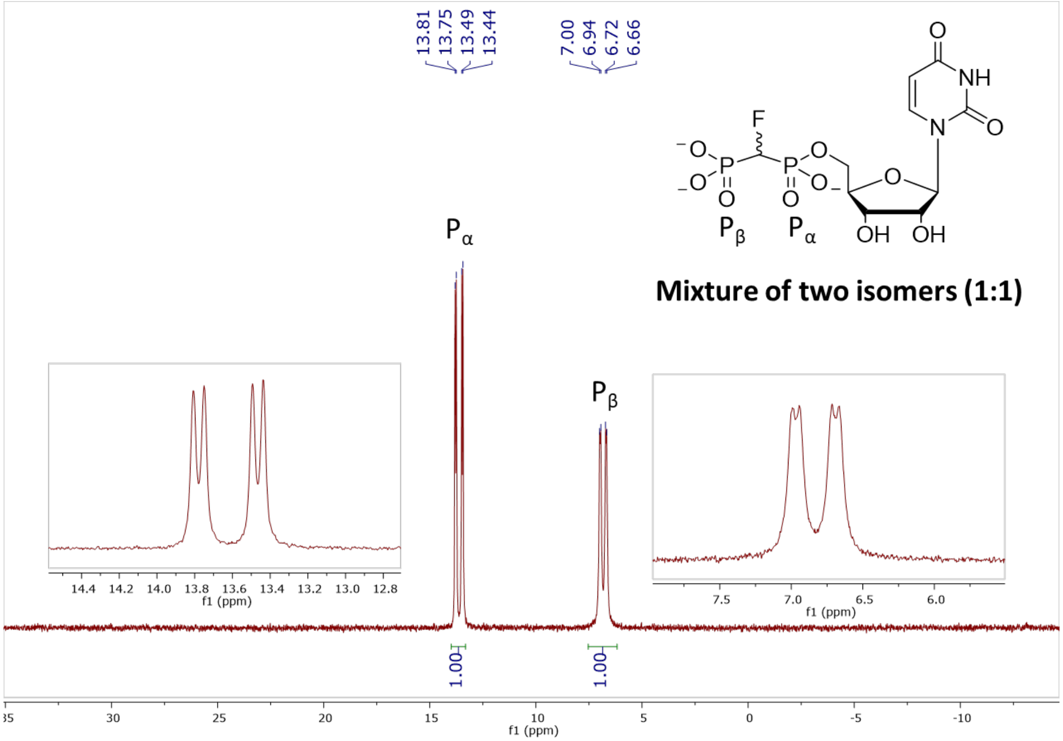

**Figure S81.** $^{31}$P NMR (202 MHz, $D_2O$, pH 7.5) of **6**

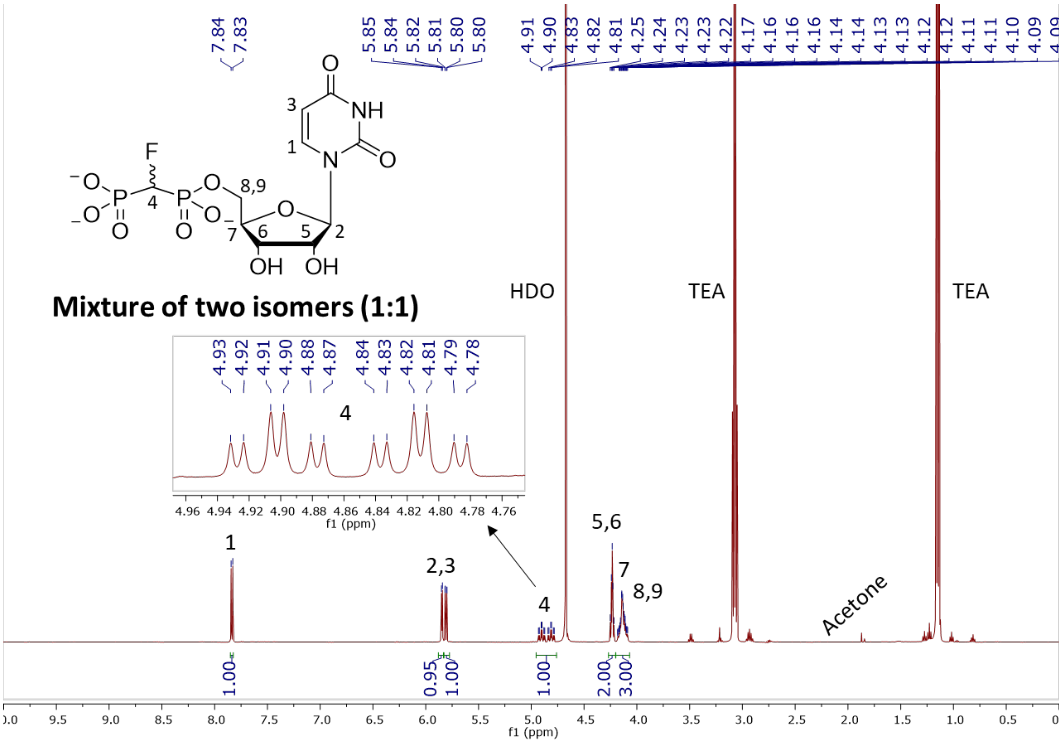

**Figure S82.** $^{1}$H NMR (500 MHz, $D_2O$) of **6**

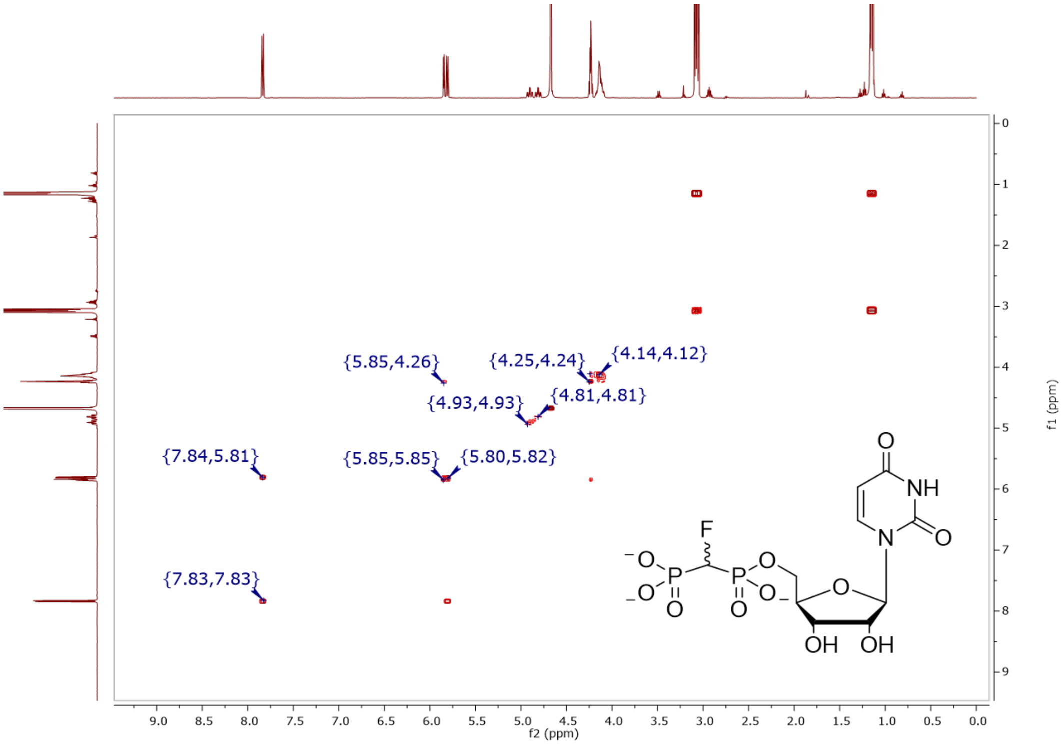

**Figure S83.** COSY (500 MHz, $D_2O$) of **6**

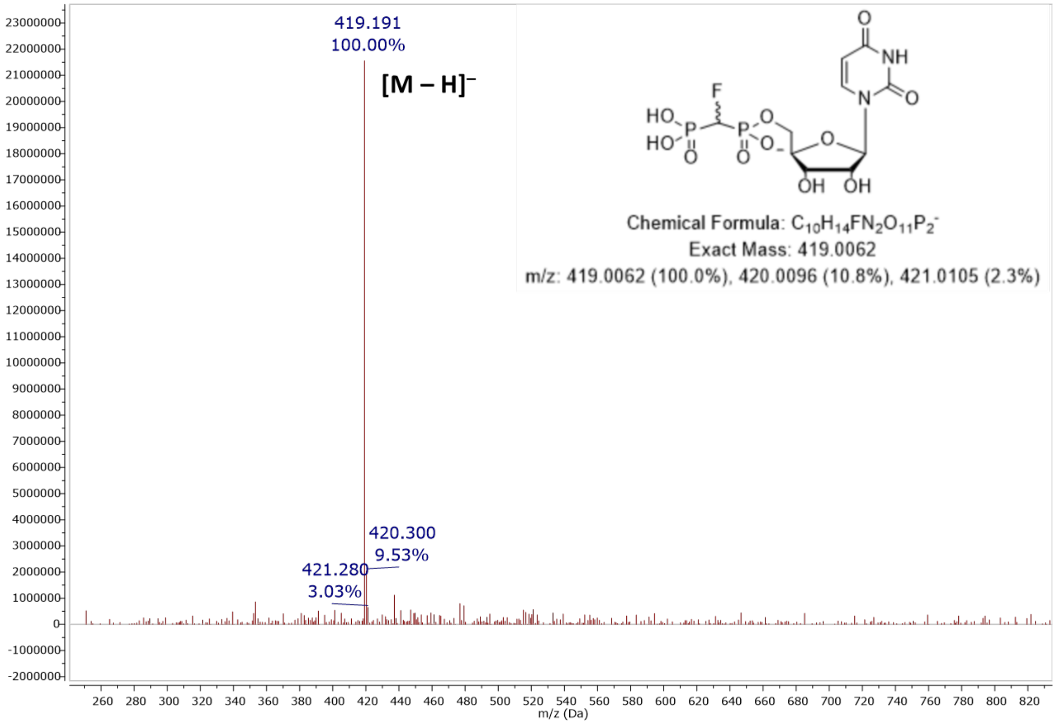

**Figure S84.** MS (ESI): [M – H]⁻ of **6**

#### Elemental Composition Report

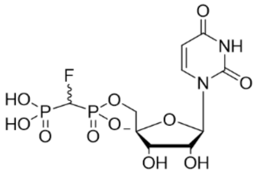

##### Single Mass Analysis

Tolerance = 5.0 PPM / DBE: min = -1.5, max = 50.0
Element prediction: Off
Number of isotope peaks used for i-FIT = 3

Monoisotopic Mass, Even Electron Ions
163 formula(e) evaluated with 2 results within limits (up to 50 closest results for each mass)
Elements Used:
C: 0-60 H: 0-100 N: 1-3 O: 10-12 F: 0-1 P: 1-3
Pouya Haratipour, USCMCK16
Synapt_26241 31 (0.620) Cm (31:35-6:8)

1: TOF MS ES-
1.77e+007

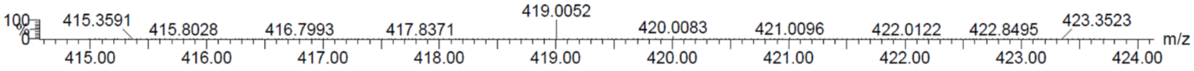

| | | | | | | | | |
|---|---|---|---|---|---|---|---|---|
| Minimum: | | | | -1.5 | | | | |
| Maximum: | | 5.0 | 5.0 | 50.0 | | | | |
| Mass | Calc. Mass | mDa | PPM | DBE | i-FIT | Norm | Conf(%) | Formula |
| 419.0052 | 419.0057 | -0.5 | -1.2 | 5.5 | 797.8 | 0.011 | 98.92 | C10 H14 N2 O11 F P2 |
| | 419.0045 | 0.7 | 1.7 | 9.5 | 802.4 | 4.533 | 1.08 | C13 H13 N2 O10 P2 |

**Figure S85.** Elemental composition report from HRMS (ESI-TOF) [M – H]⁻ of **6**

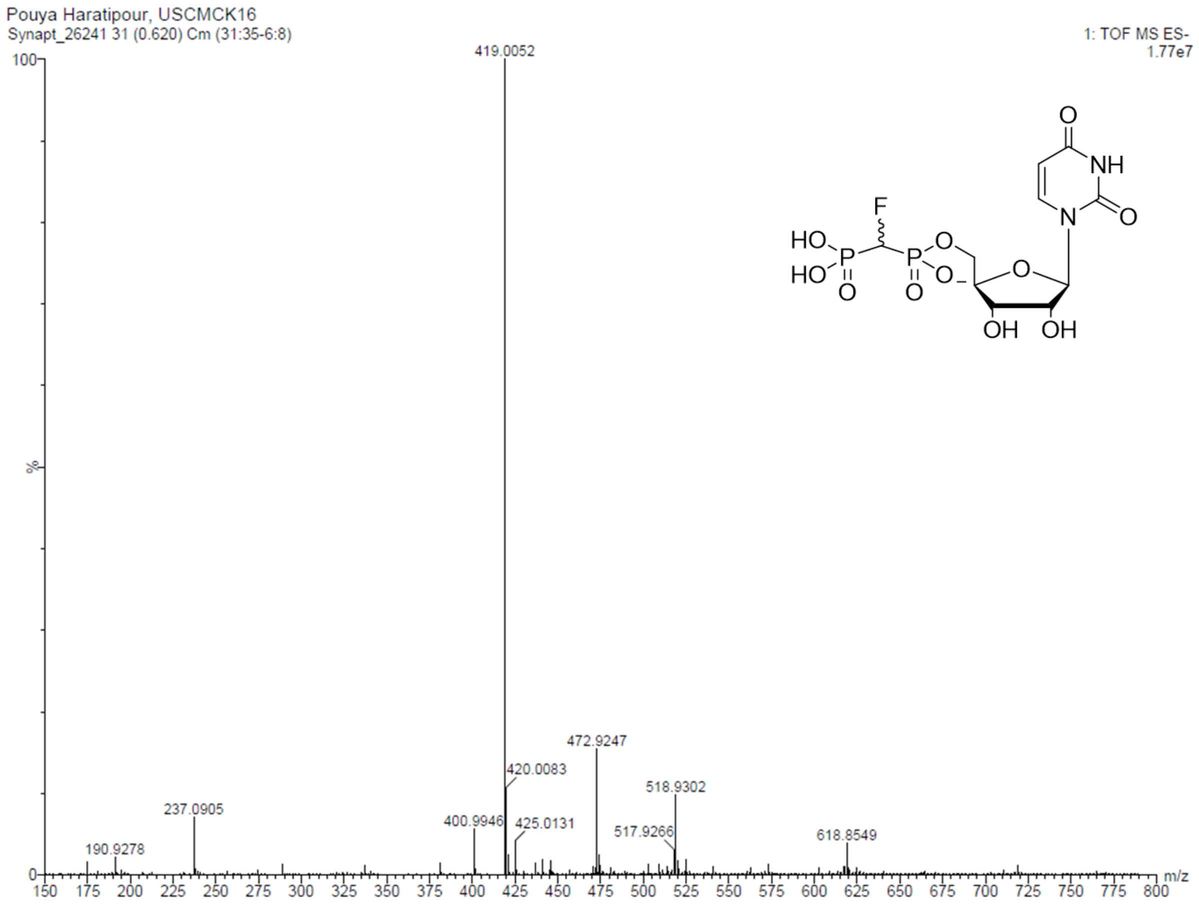

**Figure S86.** HRMS (ESI-TOF): [M – H]⁻ of **6**

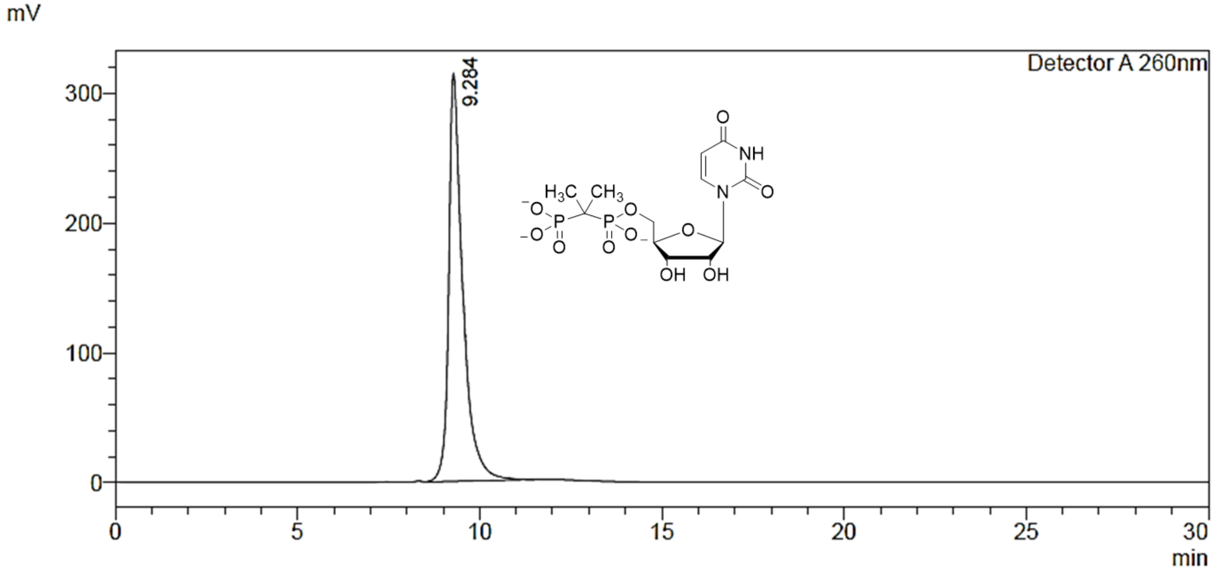

**Figure S87.** Preparative RP-HPLC purification of **9**

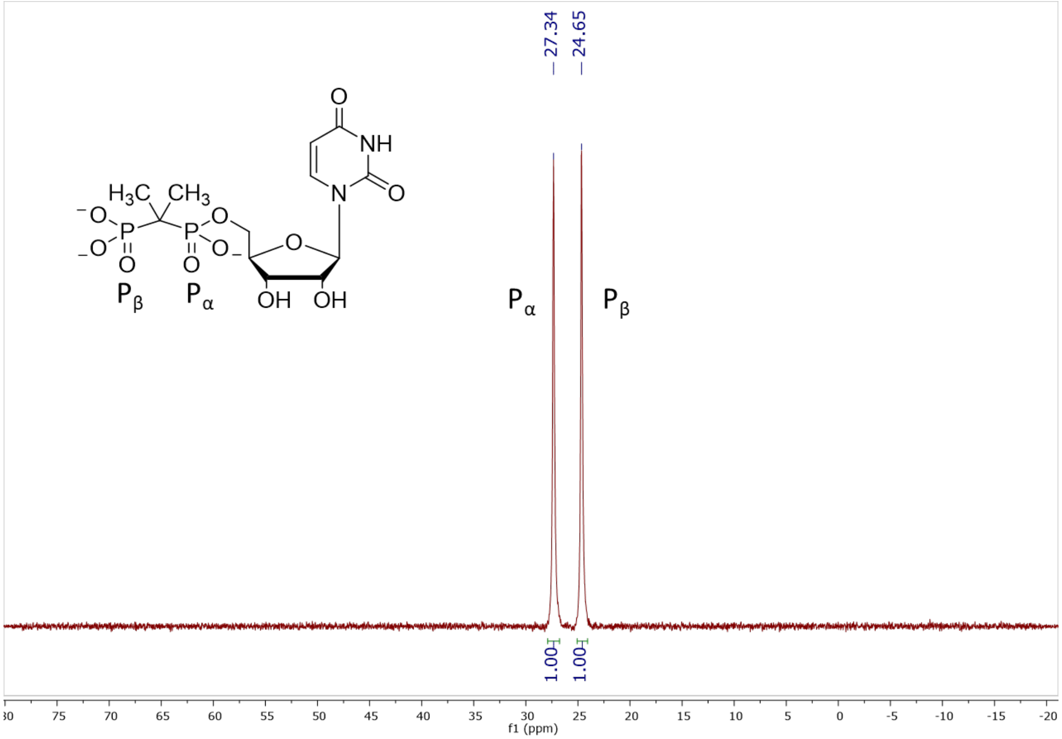

**Figure S88.** $^{31}P$ NMR (202 MHz, $D_2O$, pH 7.5) of **9**

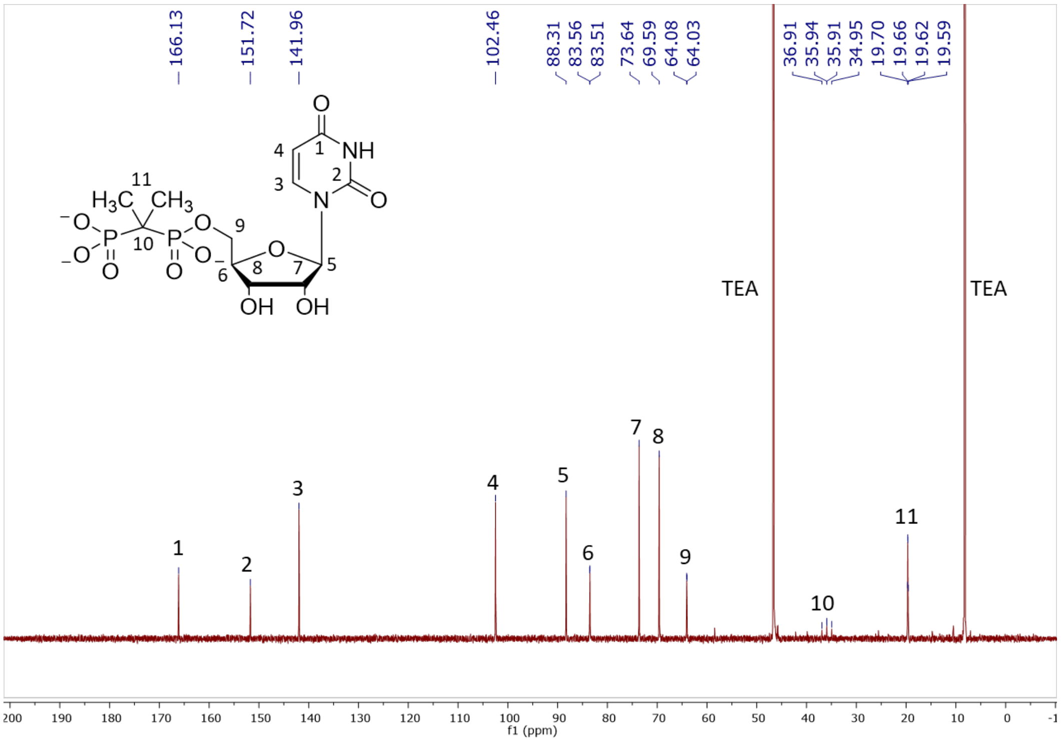

**Figure S89.** $^{13}C$ NMR (126 MHz, $D_2O$) of **9**

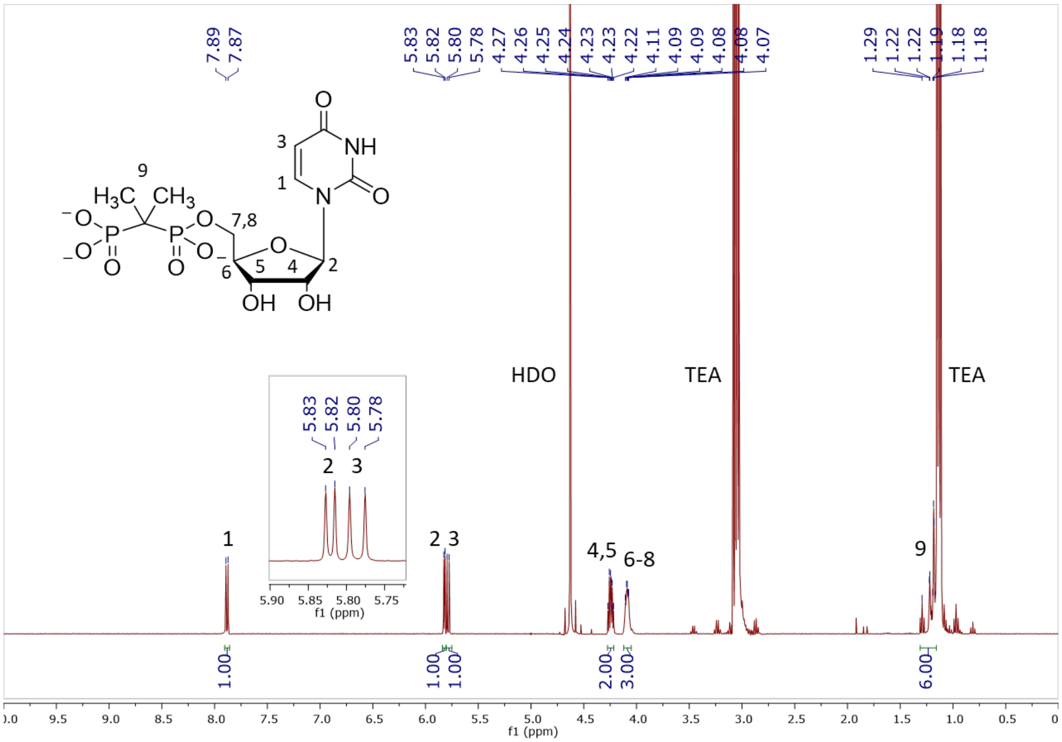

**Figure S90.** $^{1}H$ NMR (400 MHz, $D_2O$) of **9**

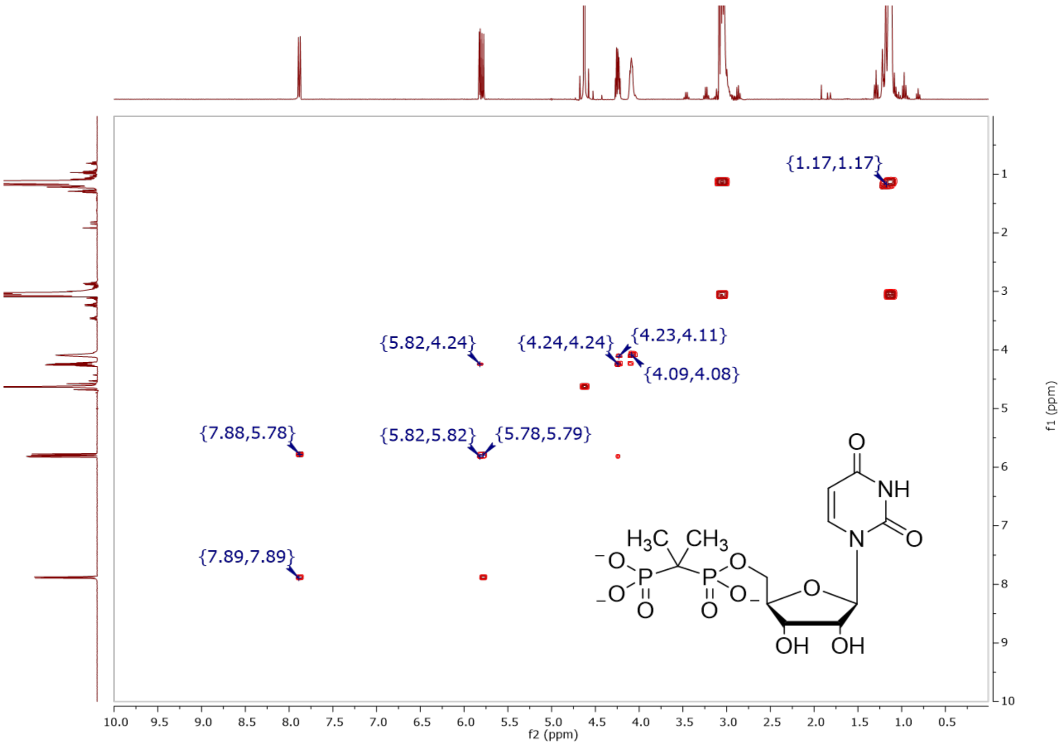

**Figure S91.** COSY (400 MHz, $D_2O$) of **9**

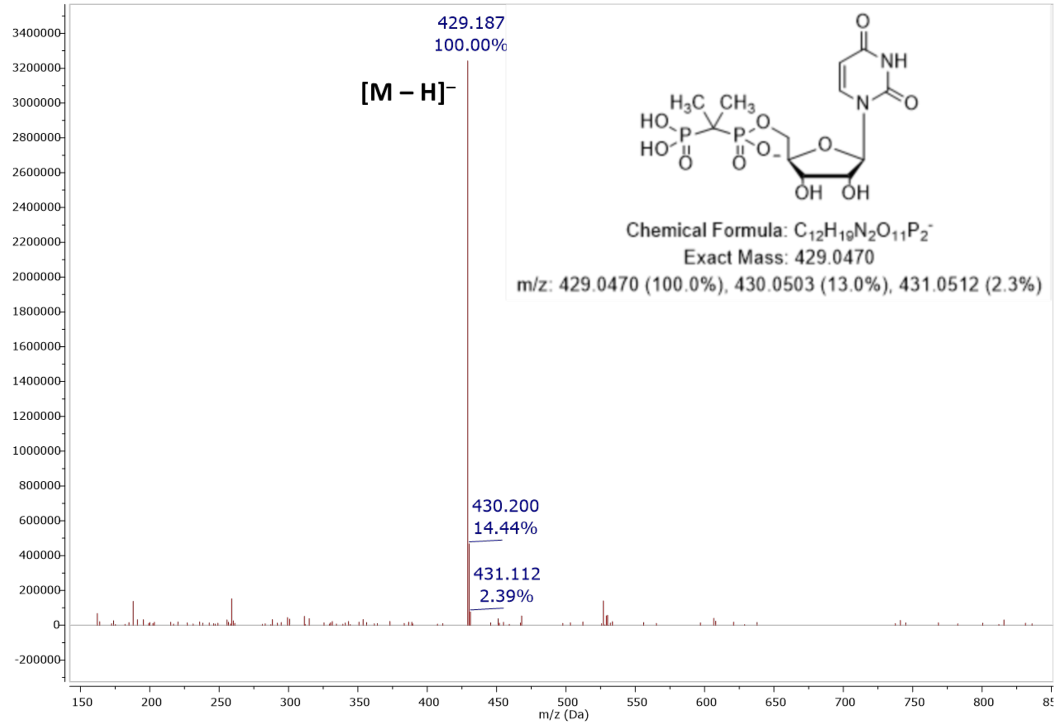

**Figure S92.** MS (ESI): [M – H]⁻ of **9**

#### Elemental Composition Report

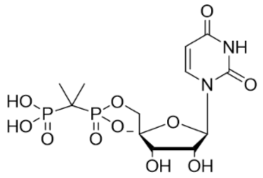

##### Single Mass Analysis

Tolerance = 5.0 PPM / DBE: min = -1.5, max = 50.0
Element prediction: Off
Number of isotope peaks used for i-FIT = 3

Monoisotopic Mass, Even Electron Ions
57 formula(e) evaluated with 1 results within limits (up to 50 closest results for each mass)
Elements Used:
C: 0-60 H: 0-80 N: 0-5 O: 10-12 P: 2-2
Pouya Haratipour USCMCK26
Synapt2_996 29 (0.586) Cm (25:30-4:10)

1: TOF MS ES-
6.45e+005

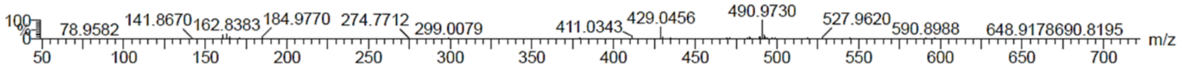

| Mass | Calc. Mass | mDa | PPM | DBE | i-FIT | Norm | Conf(%) | Formula |
|---|---|---|---|---|---|---|---|---|
| Minimum: | | | | -1.5 | | | | |
| Maximum: | | 5.0 | 5.0 | 50.0 | | | | |
| 429.0456 | 429.0464 | -0.8 | -1.9 | 5.5 | 1106.5 | n/a | n/a | C12 H19 N2 O11 P2 |

**Figure S93.** Elemental composition report from HRMS (ESI-TOF) [M – H]⁻ of **9**

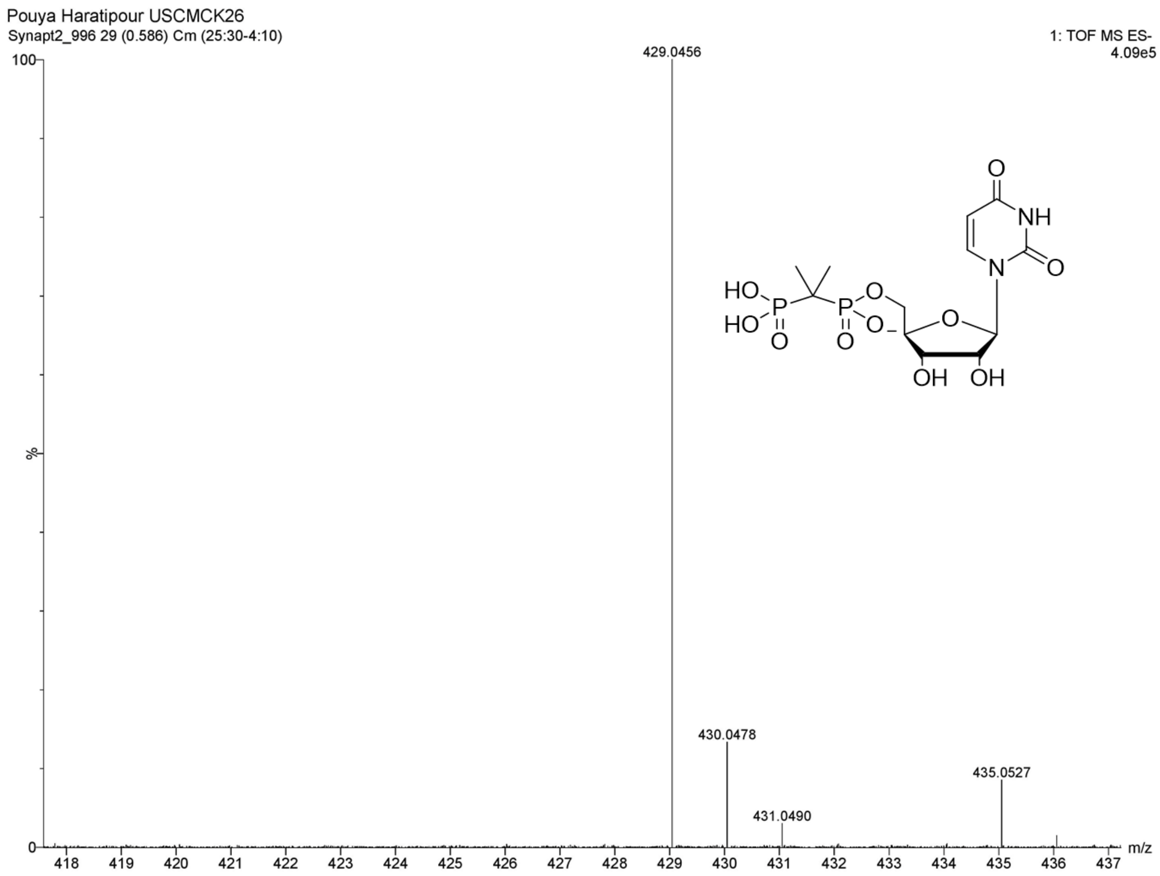

**Figure S94.** HRMS (ESI-TOF): [M – H]⁻ of **9**

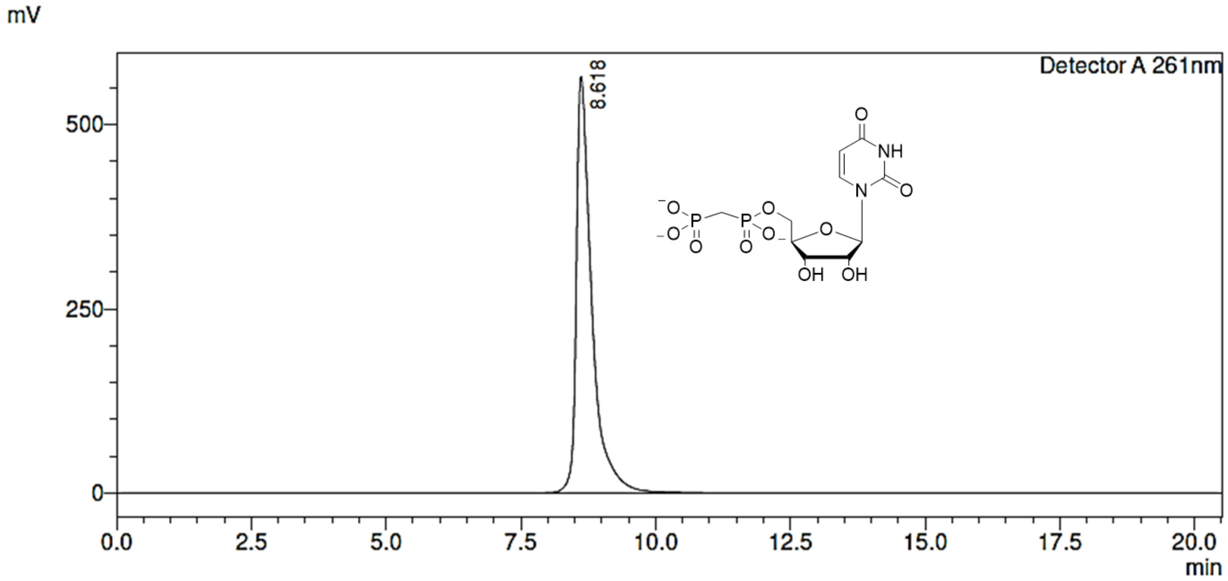

**Figure S95.** Preparative RP-HPLC purification of **10**

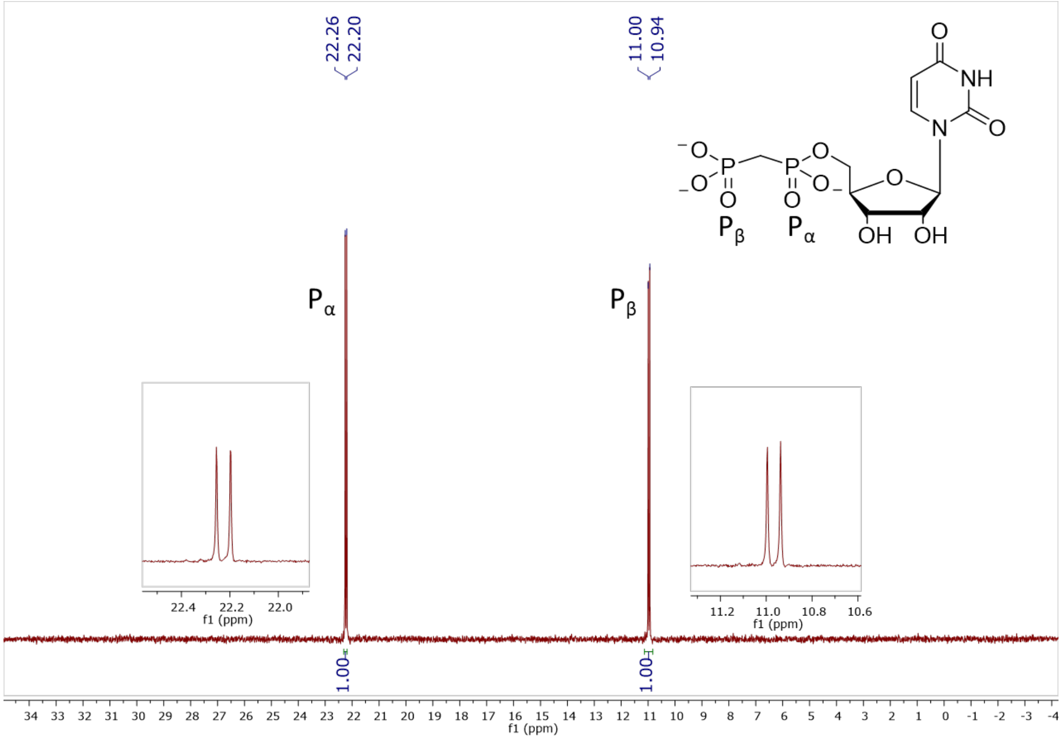

**Figure S96.** $^{31}P$ NMR (162 MHz, $D_2O$, pH 11.7) of **10**

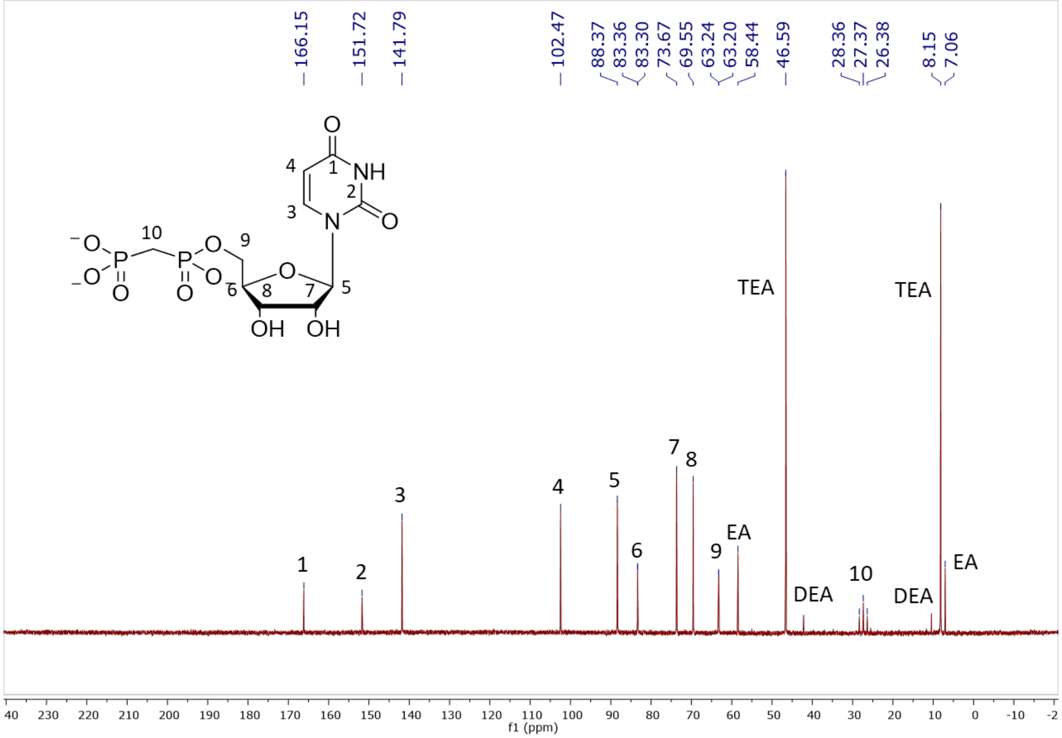

**Figure S97.** $^{13}C$ NMR (126 MHz, $D_2O$) of **10**

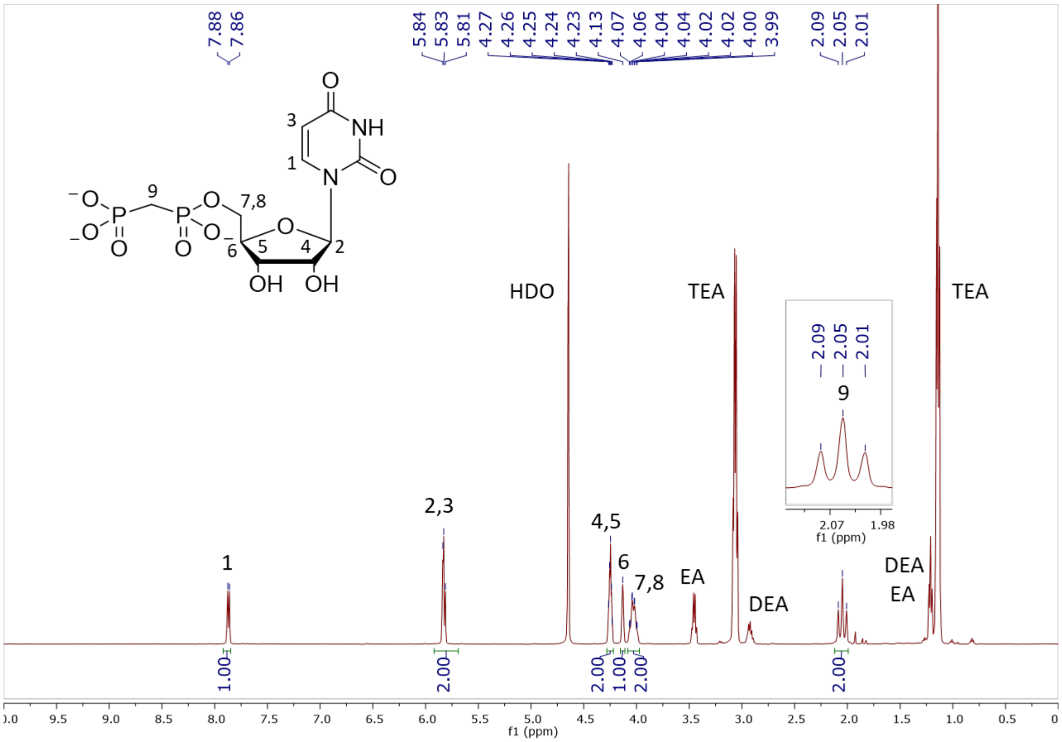

**Figure S98.** $^{1}H$ NMR (500 MHz, $D_2O$) of **10**

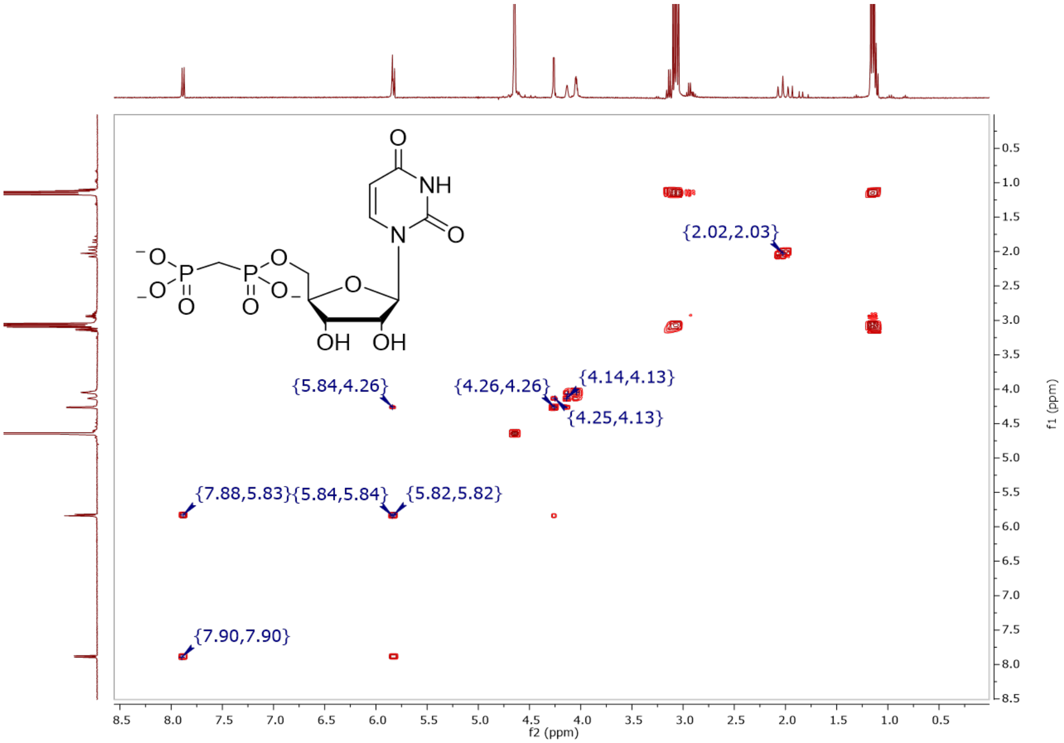

**Figure S99.** COSY (400 MHz, $D_2O$) of **10**

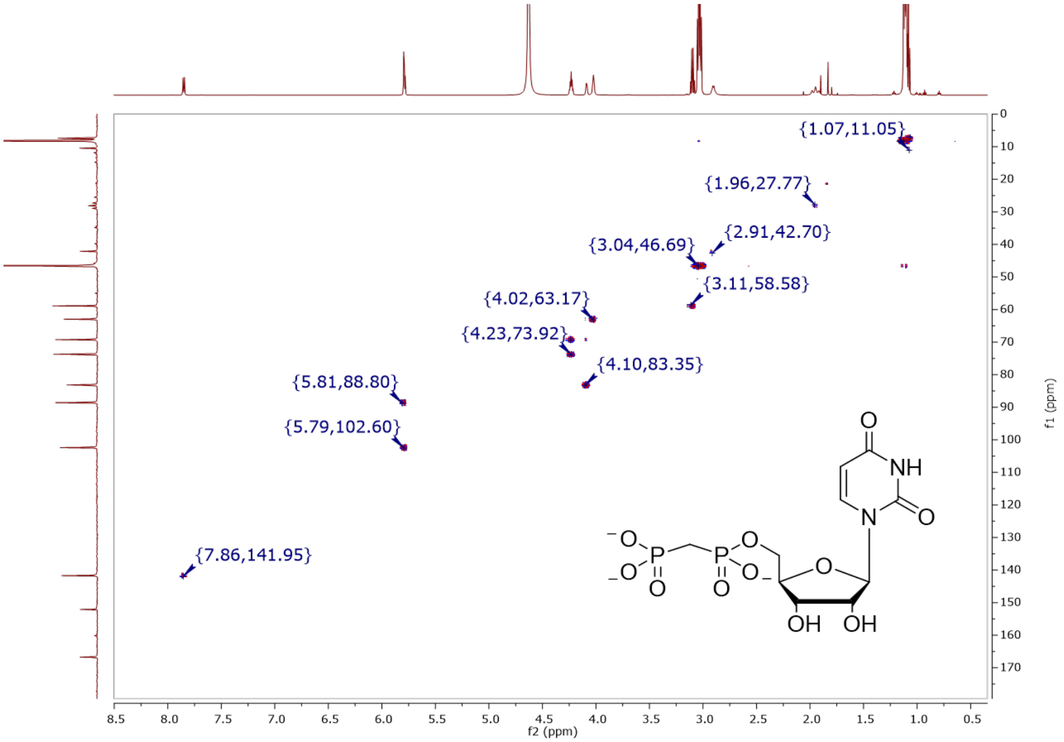

**Figure S100.** HSQCAD (600, 150 MHz, $D_2O$) of **10**

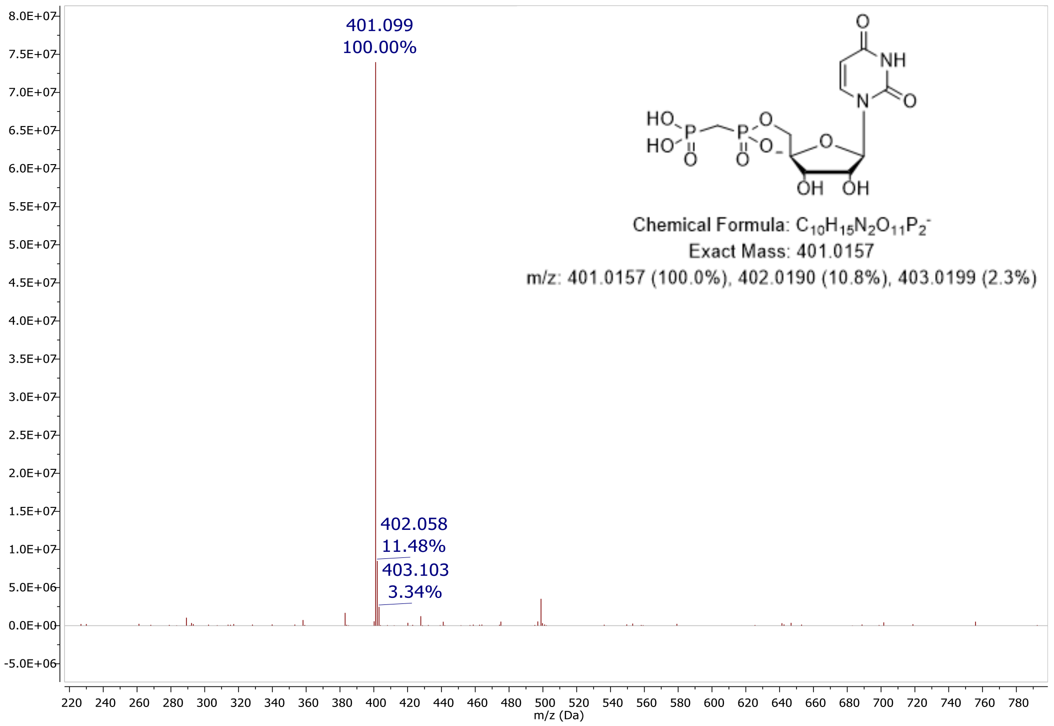

**Figure S101.** MS (ESI): $[M – H]^–$ of **10**

#### Elemental Composition Report

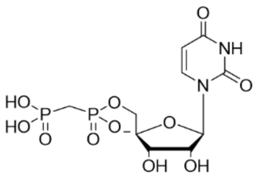

##### Single Mass Analysis

Tolerance = 5.0 PPM / DBE: min = -1.5, max = 50.0
Element prediction: Off
Number of isotope peaks used for i-FIT = 3

Monoisotopic Mass, Even Electron Ions
148 formula(e) evaluated with 2 results within limits (up to 50 closest results for each mass)
Elements Used:
C: 0-60 H: 0-100 N: 1-3 O: 10-12 F: 0-1 P: 1-3
Pouya Haratipour, USCMCK17
Synapt_26242 45 (0.896) Cm (45:47-5:8)

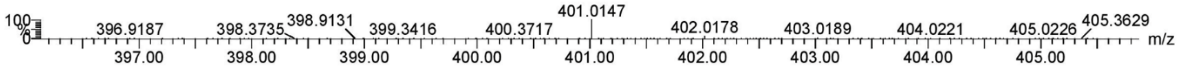

| | | | | | | | |
|---|---|---|---|---|---|---|---|
| Minimum: | | | | -1.5 | | | | |
| Maximum: | | 5.0 | 5.0 | 50.0 | | | | |
| Mass | Calc. Mass | mDa | PPM | DBE | i-FIT | Norm | Conf(%) | Formula |
| 401.0147 | 401.0151 | -0.4 | -1.0 | 5.5 | 574.1 | 0.013 | 98.69 | C10 H15 N2 O11 P2 |
| | 401.0163 | -1.6 | -4.0 | 1.5 | 578.4 | 4.335 | 1.31 | C7 H16 N2 O12 F P2 |

**Figure S102.** Elemental composition report from HRMS (ESI-TOF) $[M - H]^-$ of **10**

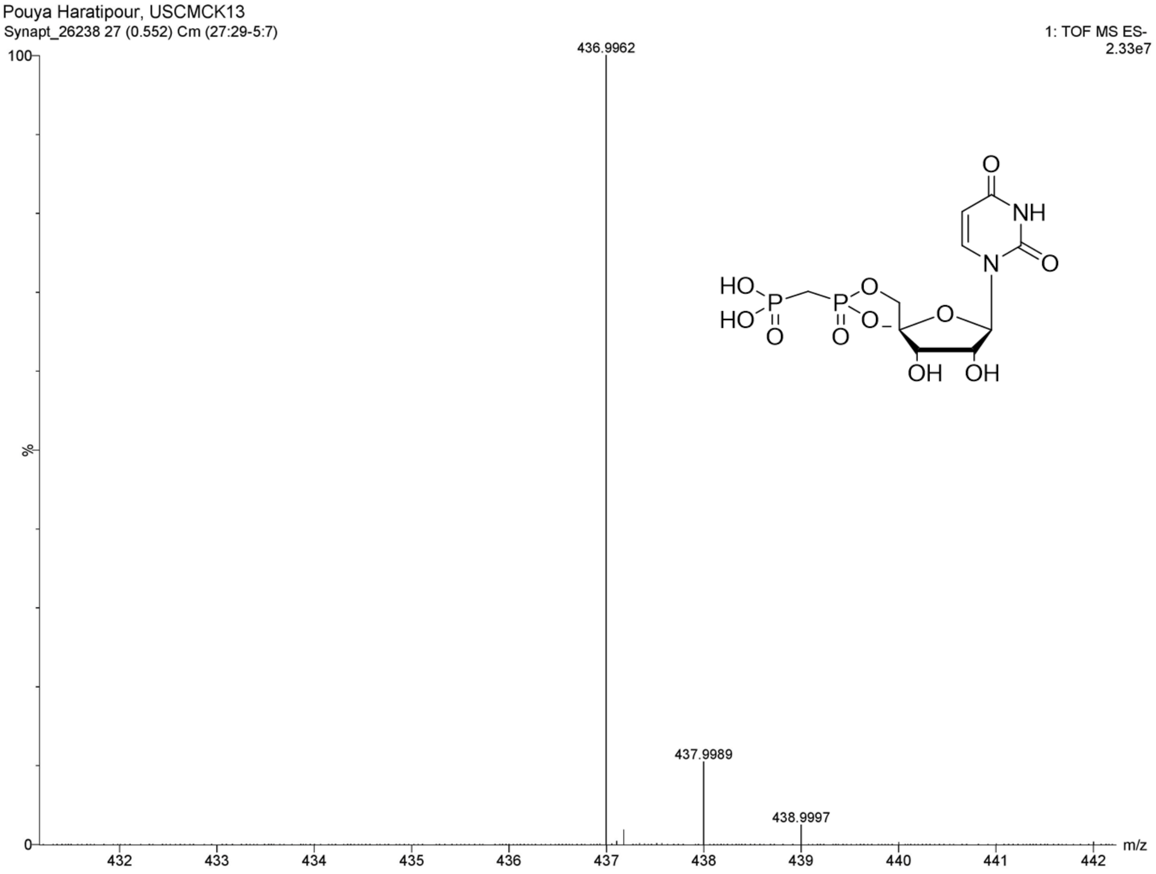

**Figure S103.** HRMS (ESI-TOF): $[M - H]^-$ of **10**

#### Spectra and chromatograms of GlcNAc-CXY-UBP analogues and intermediates

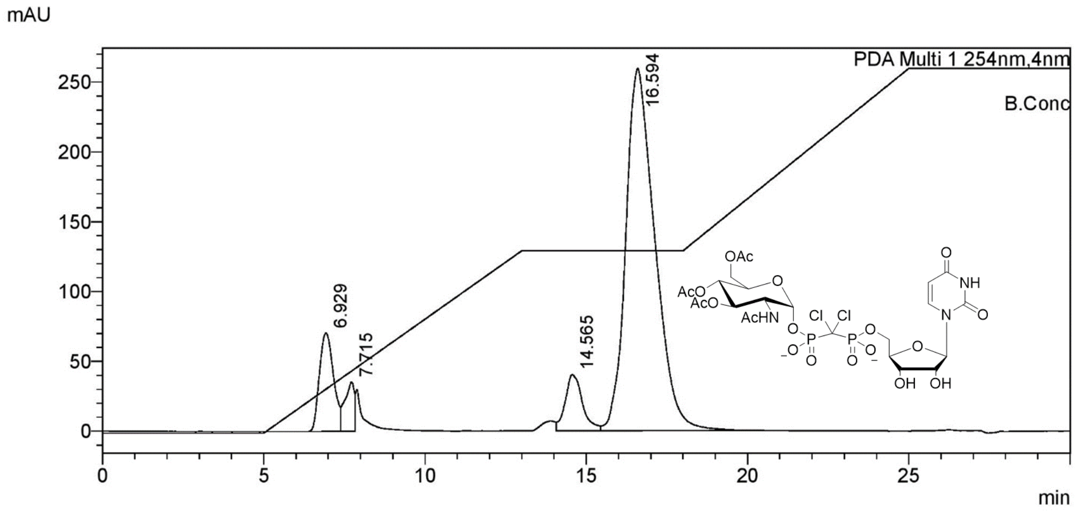

**Figure S104.** Preparative SAX-HPLC purification of **25a**

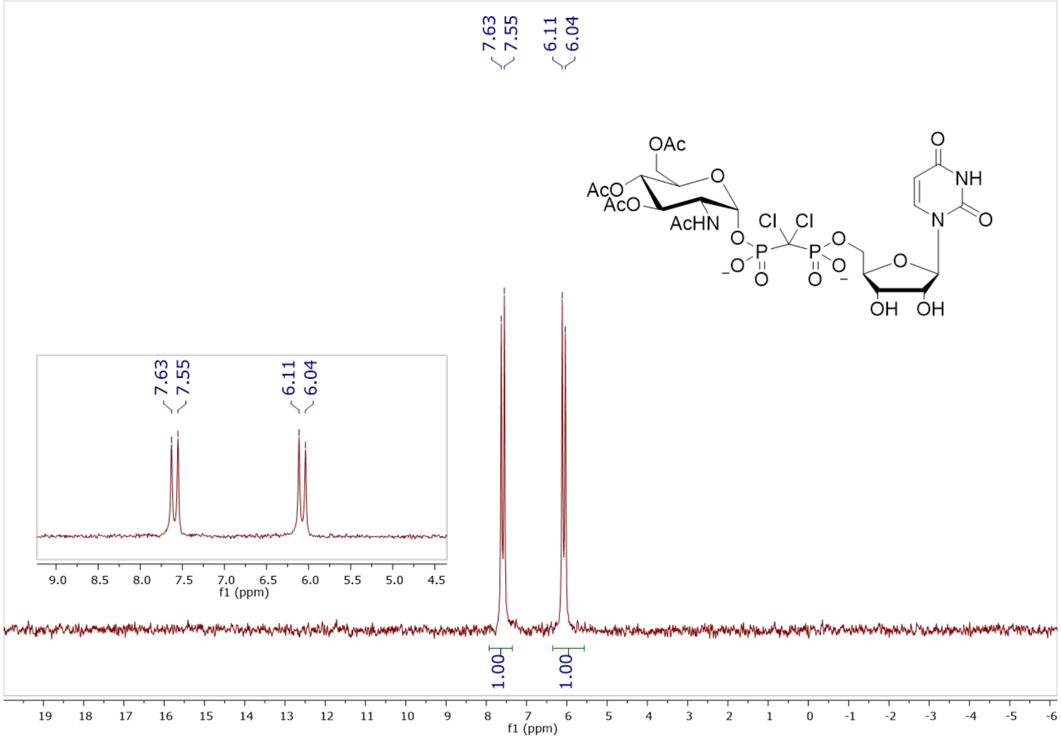

**Figure S105.** $^{31}$P NMR (202 MHz, $D_2O$, pH 7.5) of **25a**

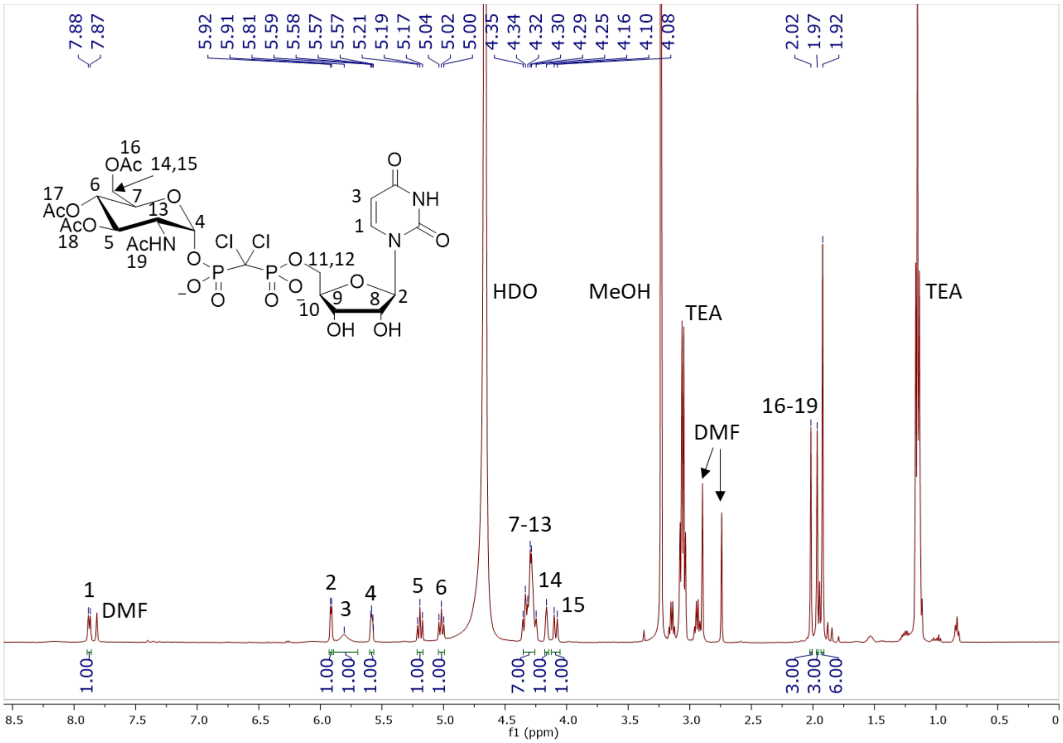

**Figure S106.** $^{1}H$ NMR (500 MHz, $D_2O$) of **25a**

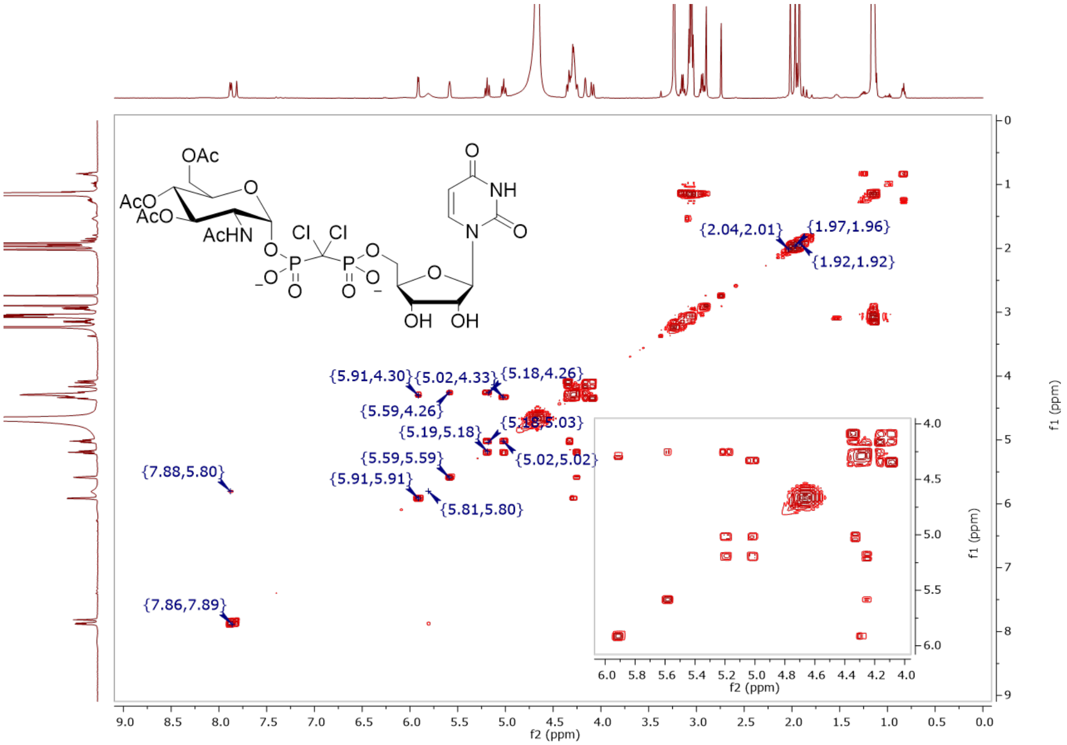

**Figure S107.** COSY (500 MHz, $D_2O$) of **25a**

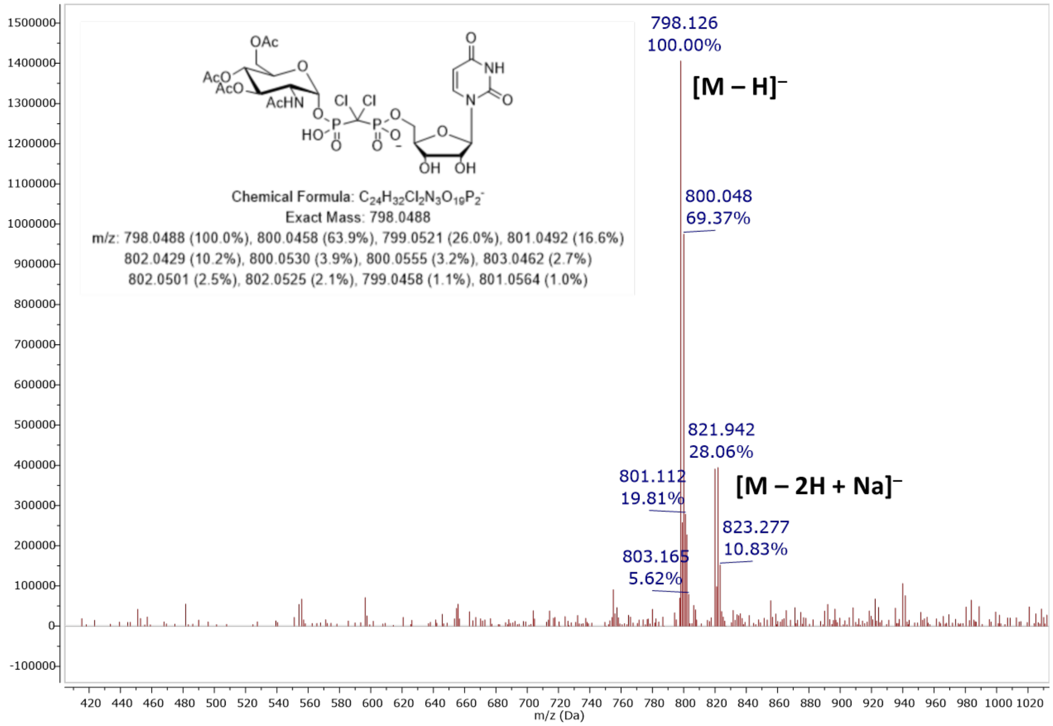

**Figure S108.** MS (ESI): [M – H]⁻ of **25a**

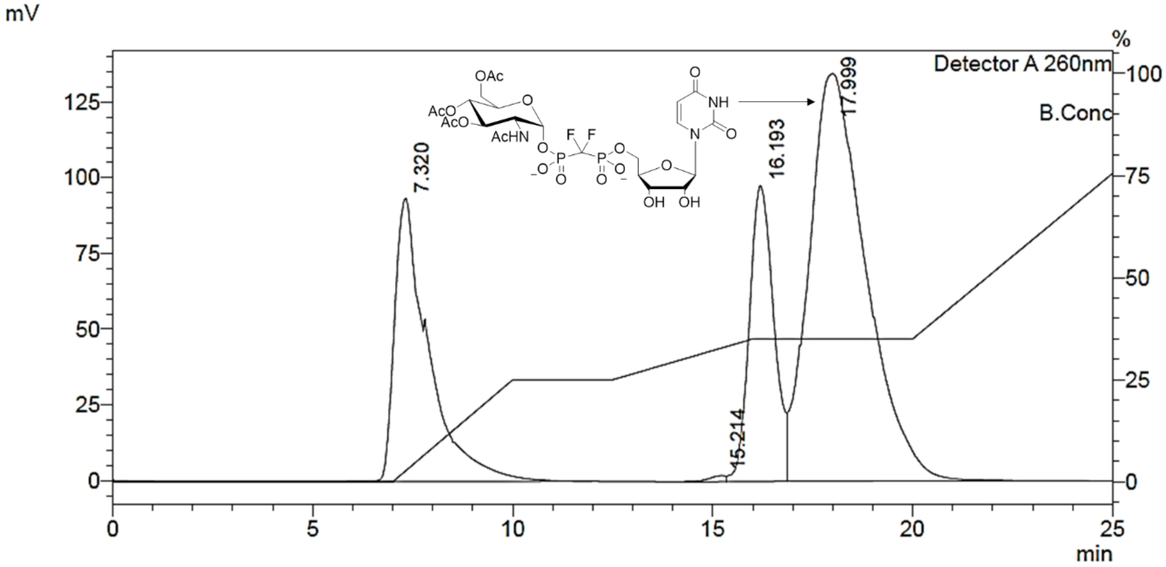

**Figure S109.** Preparative SAX-HPLC purification of **25b-α**

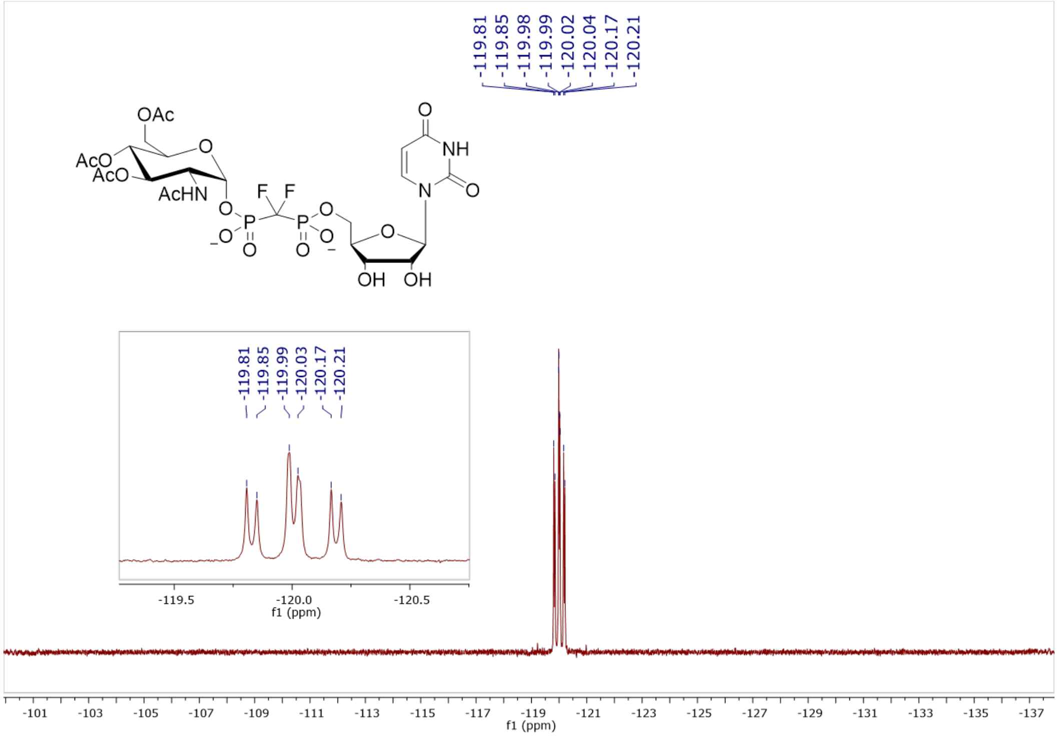

**Figure S110.** $^{19}F$ NMR (470 MHz, $D_2O$, pH 7.5) of **25b-α**

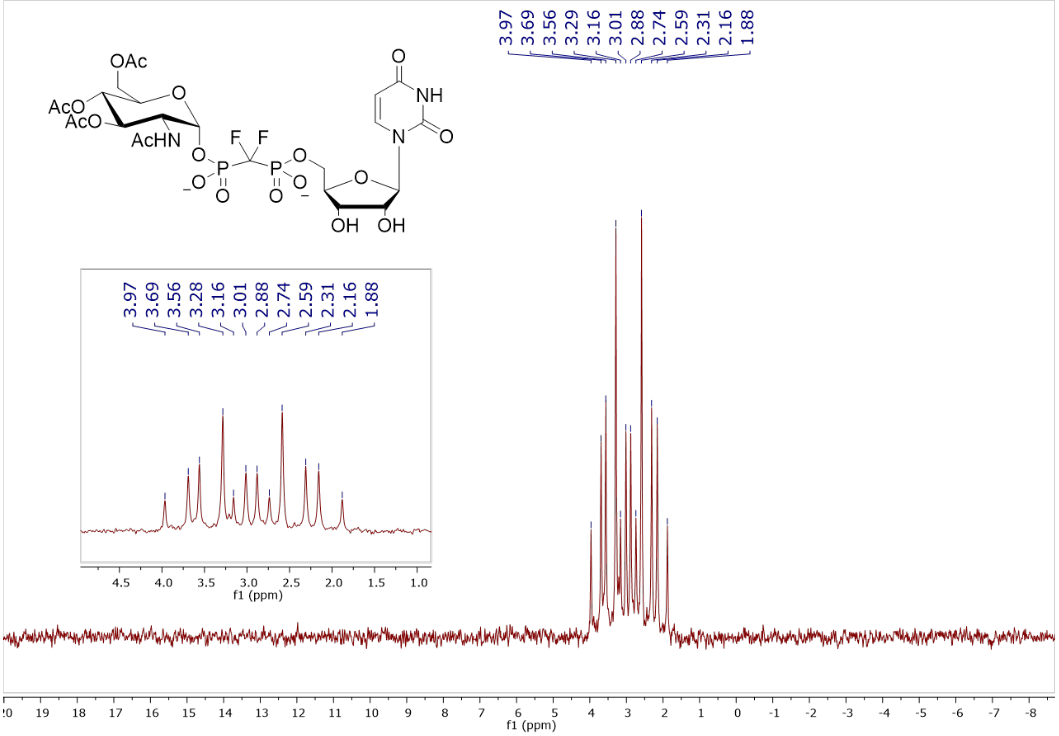

**Figure S111.** $^{31}P$ NMR (202 MHz, $D_2O$, pH 7.5) of **25b-α**

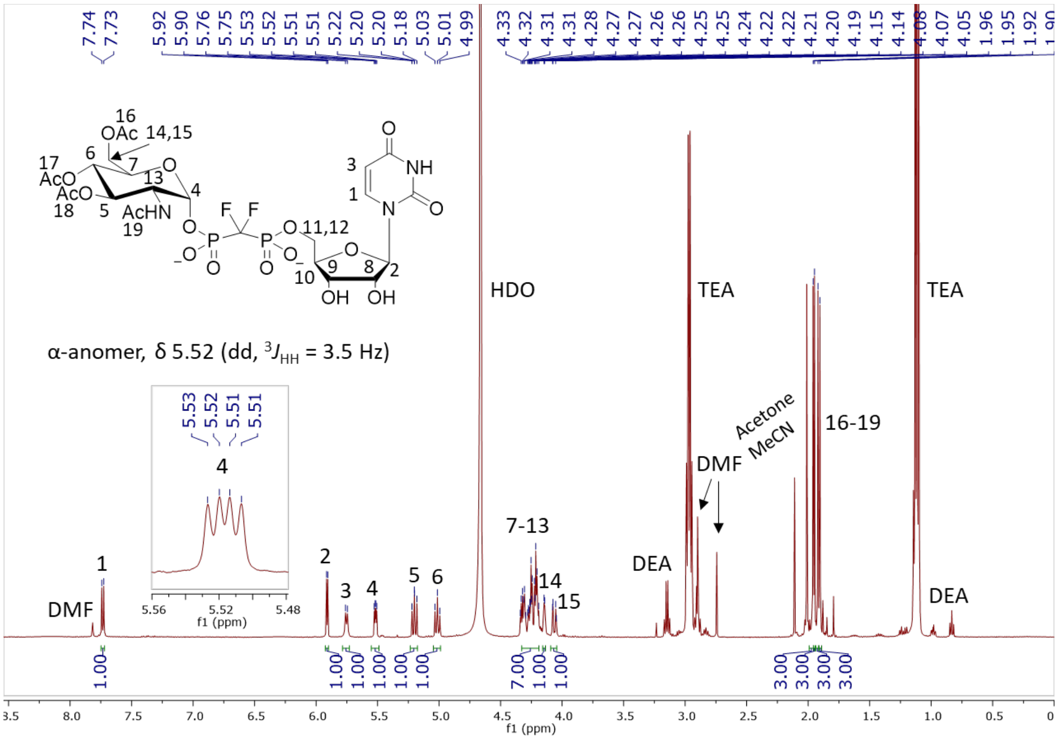

**Figure S112.** $^1$H NMR (500 MHz, $D_2O$) of **25b-α**

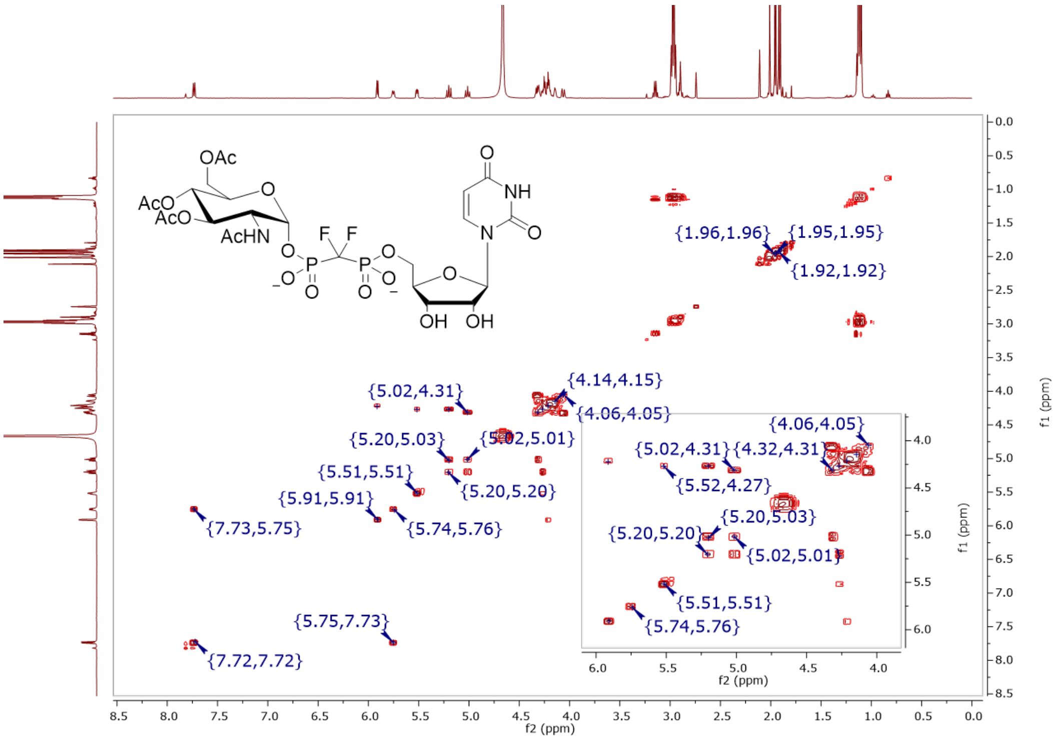

**Figure S113.** COSY (500 MHz, $D_2O$) of **25b-α**

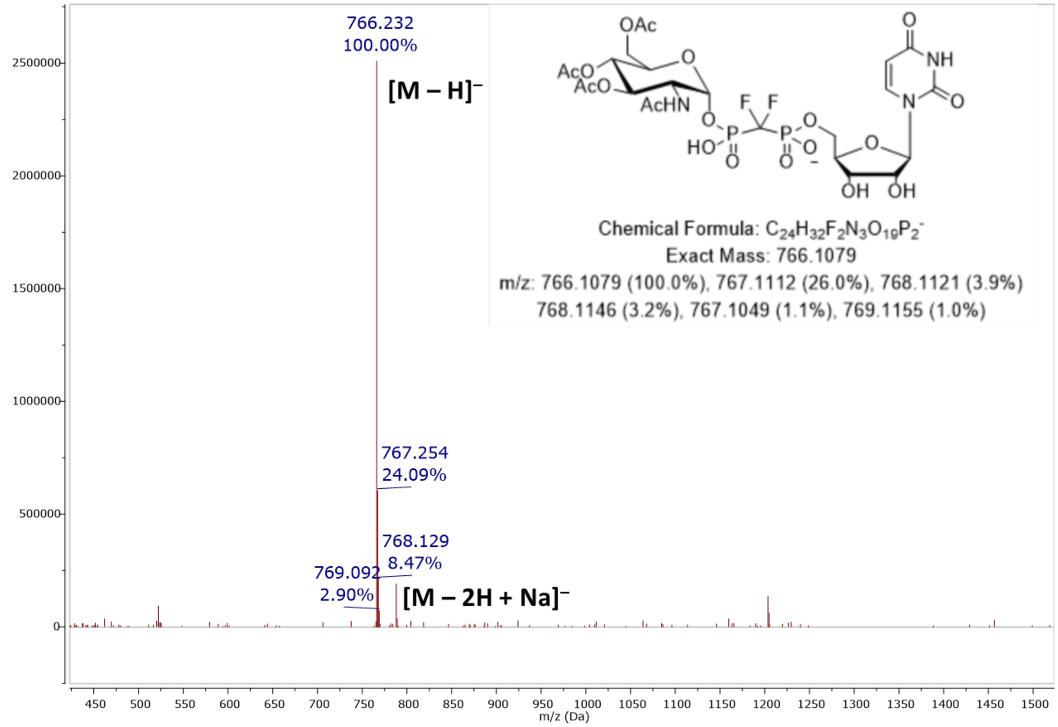

**Figure S114.** MS (ESI): [M – H]⁻ of **25b-α**

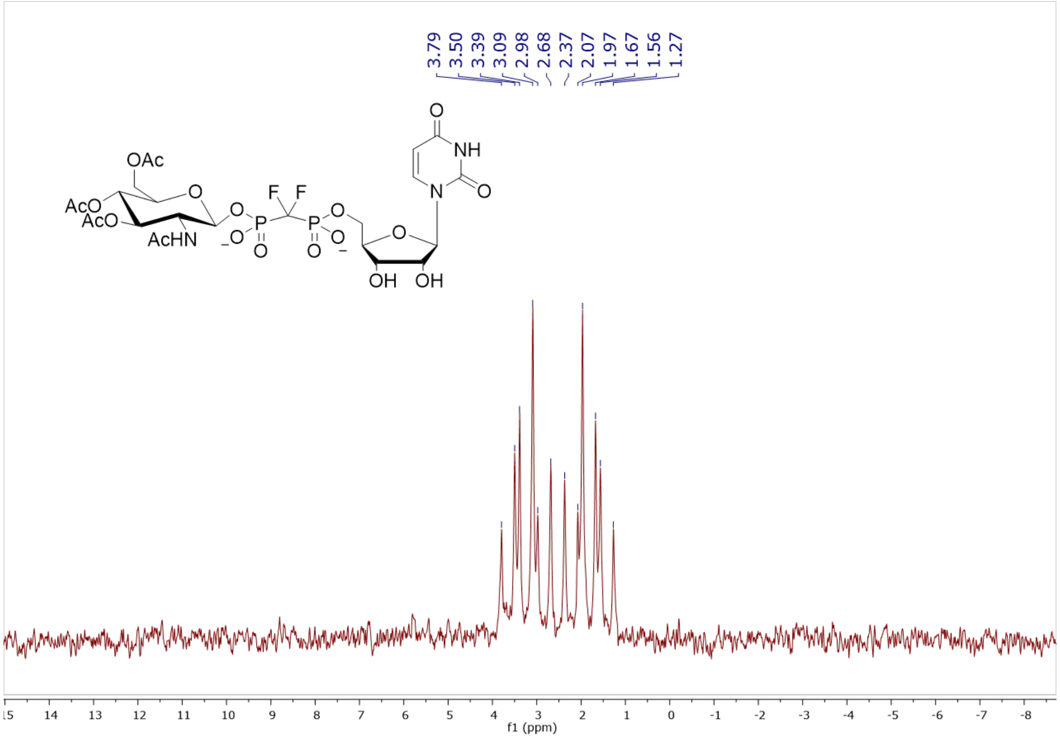

**Figure S115.** $^{31}$P NMR (202 MHz, $D_2O$, pH 7.5) of **25b-β**

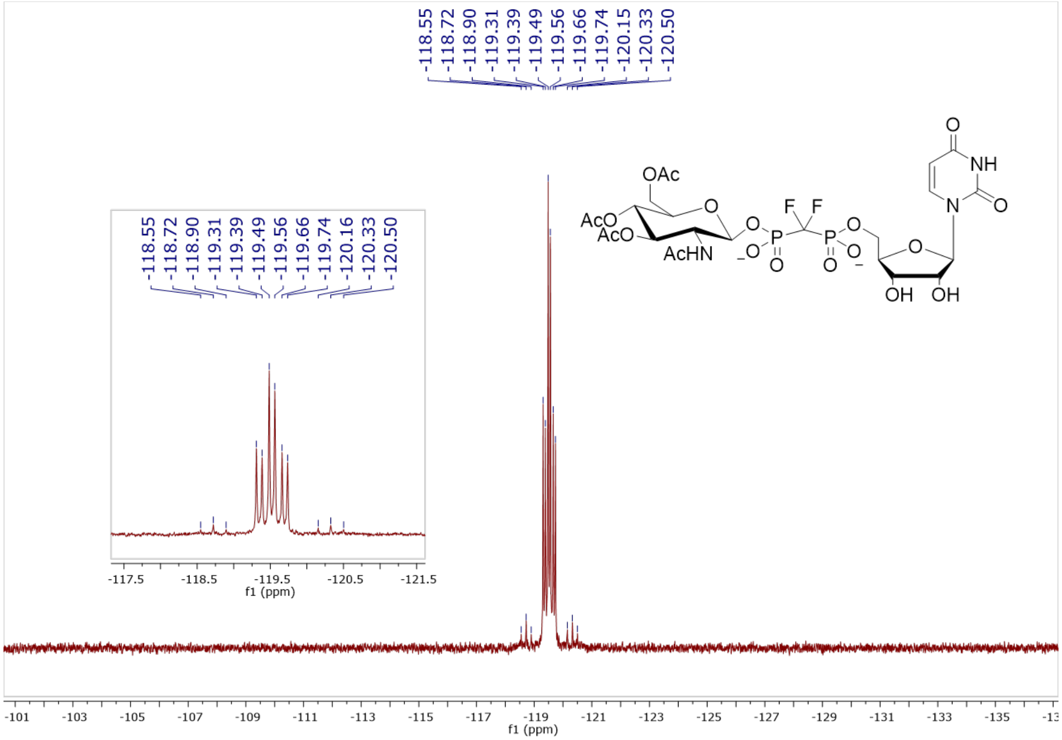

**Figure S116.** $^{19}F$ NMR (470 MHz, $D_2O$, pH 7.5) of **25b-β**

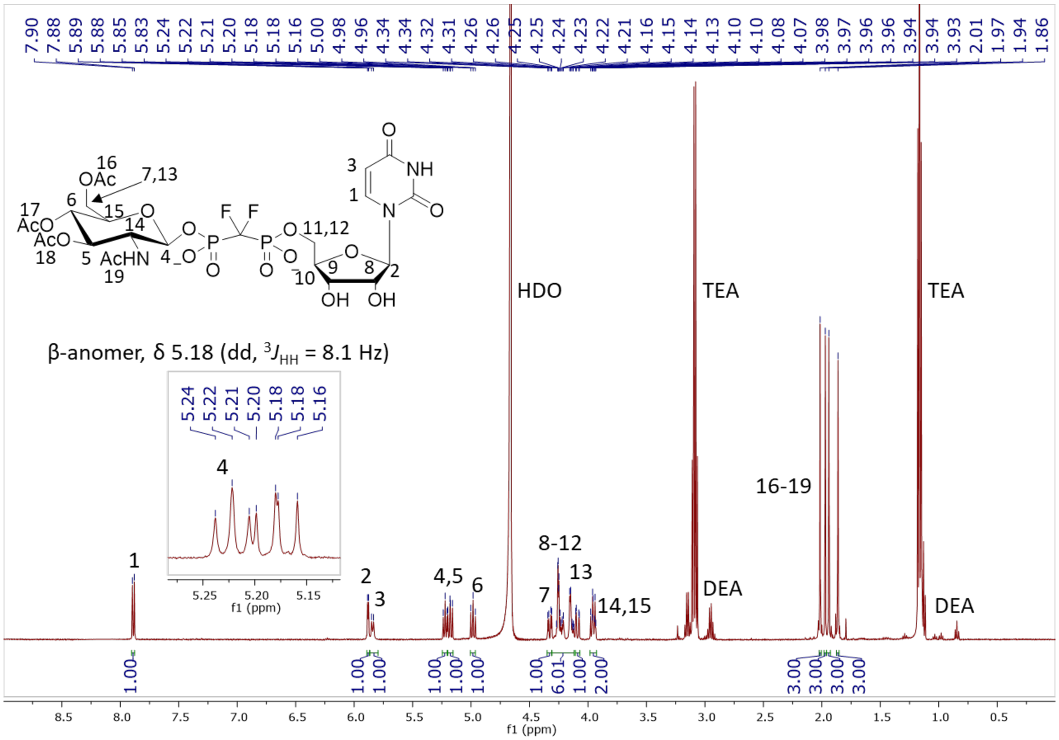

**Figure S117.** $^{1}H$ NMR (500 MHz, $D_2O$) of **25b-β**

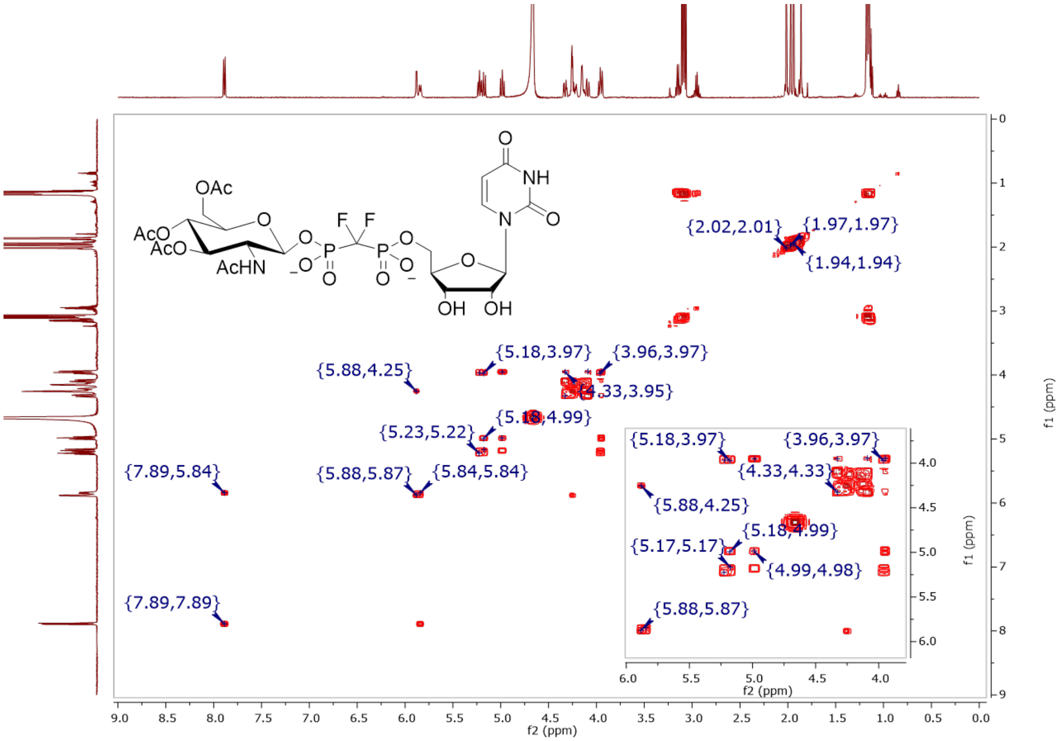

**Figure S118.** COSY (500 MHz, $D_2O$) of **25b-β**

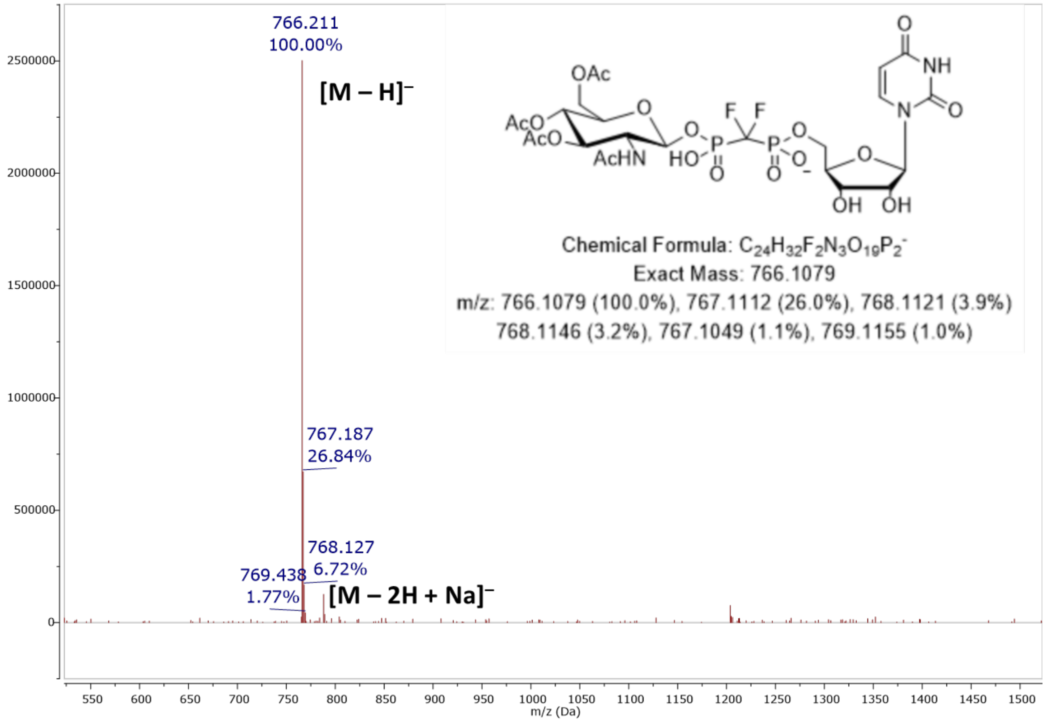

**Figure S119.** MS (ESI): $[M – H]^-$ of **25b-β**

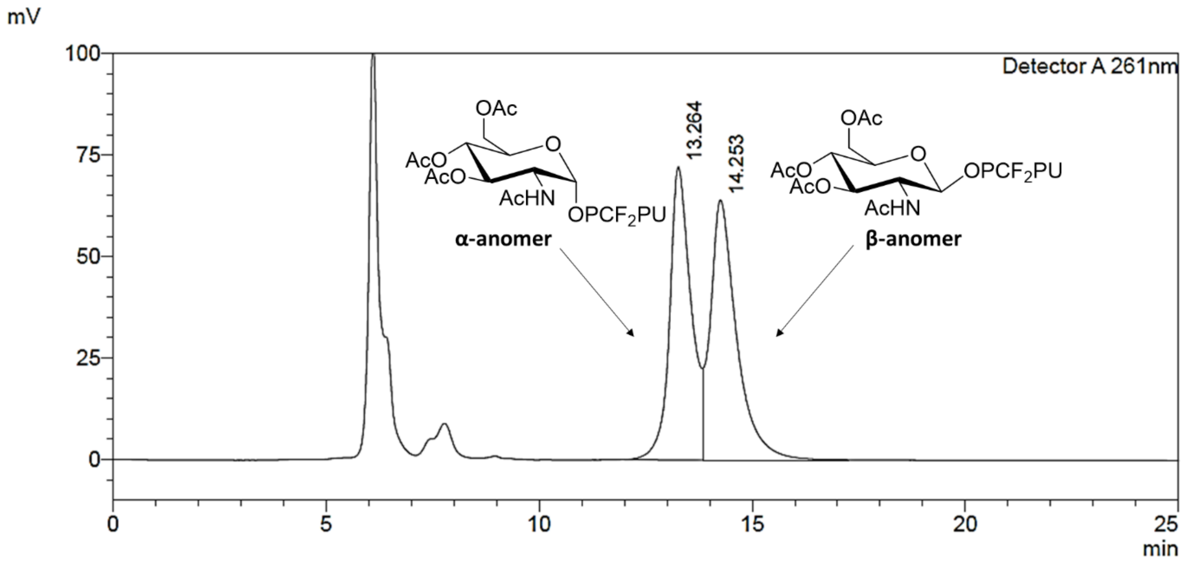

**Figure S120.** Preparative RP-HPLC separation of **25b-α** and **25b-β** (1:1)

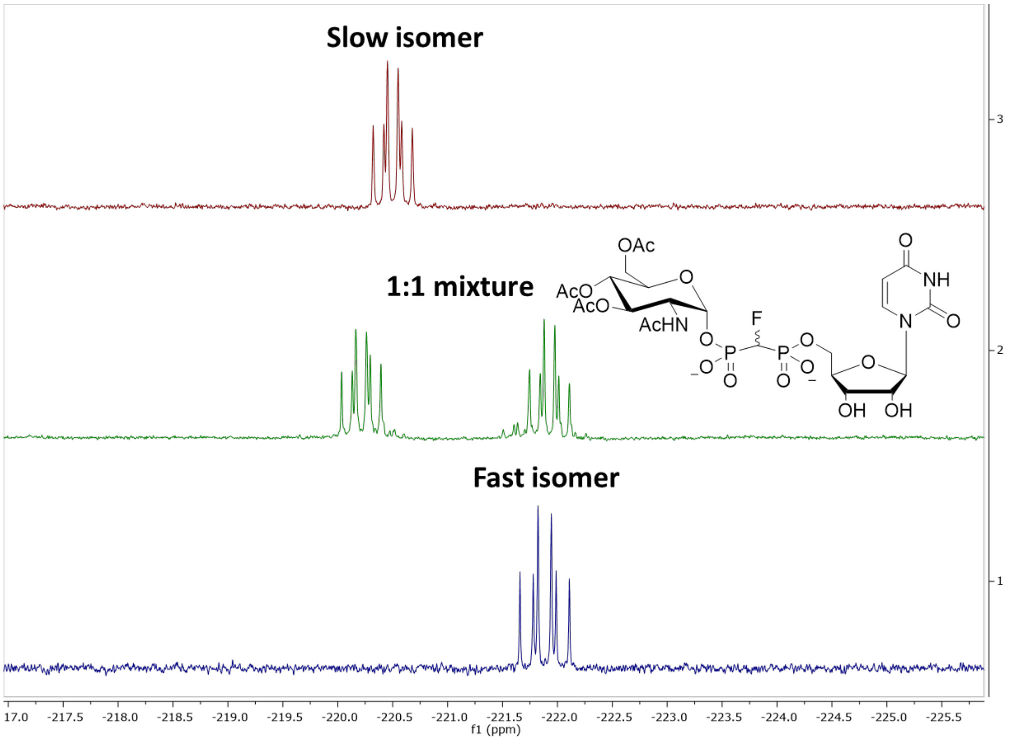

**Figure S121.** $^{19}$F NMR (spectra 2,3: 470 MHz, spectrum 1: 376 MHz, $D_2O$, pH 7.5) of **25c-α1** (fast isomer, (*S*)-CHF) and **25c-α2** (slow isomer, (*R*)-CHF)

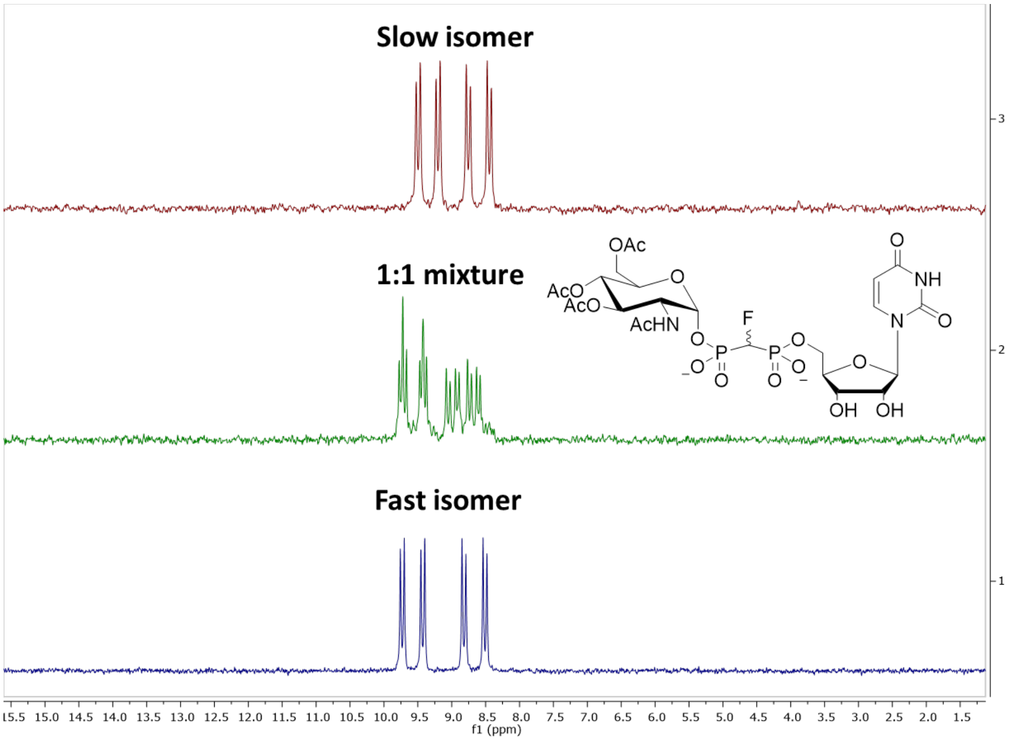

**Figure S122.** $^{31}P$ NMR (202 MHz, $D_2O$, pH 7.5) of **25c-α1** (fast eluting isomer, (*S*)-CHF) and **25c-α2** (slow eluting isomer, (*R*)-CHF)

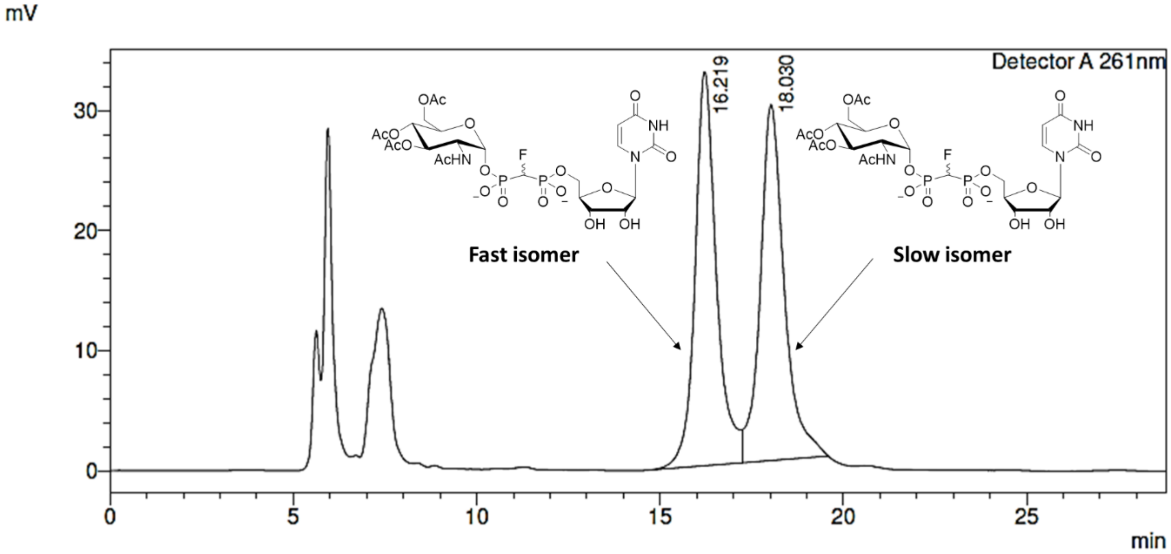

**Figure S123.** Preparative RP-HPLC separation of **25c-α1** (fast eluting isomer, (*S*)-CHF) and **25c-α2** (slow eluting isomer, (*R*)-CHF) (1:1)

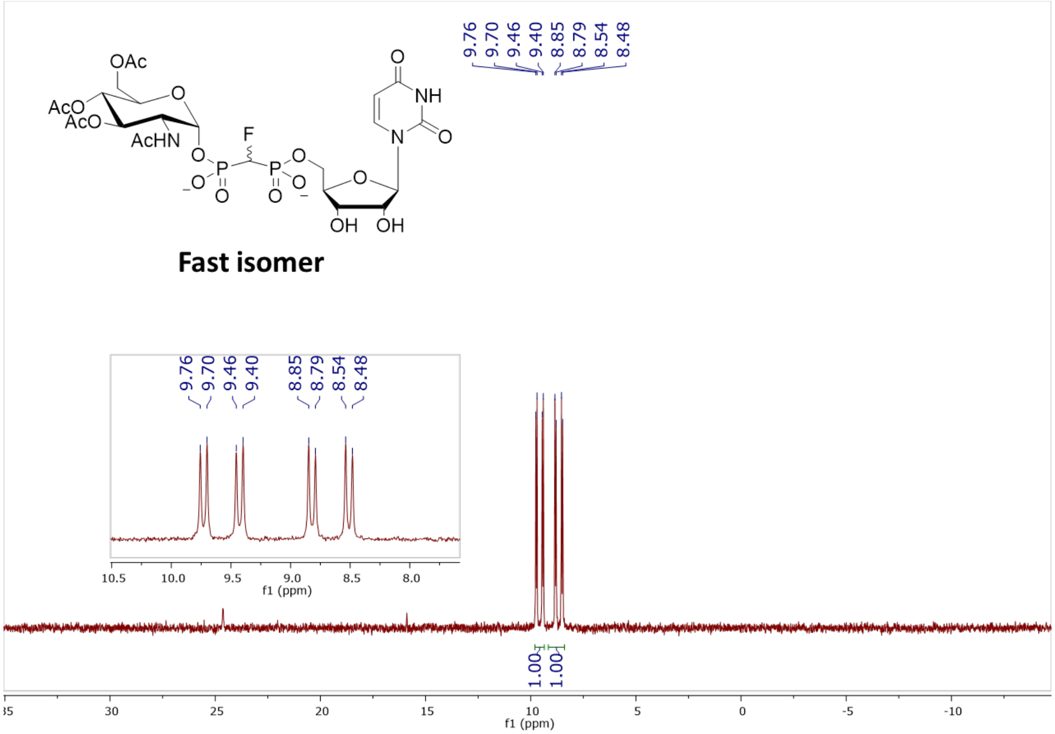

**Figure S124.** $^{31}$P NMR (202 MHz, $D_2O$, pH 7.5) of **25c-α1**, (*S*)-CHF isomer

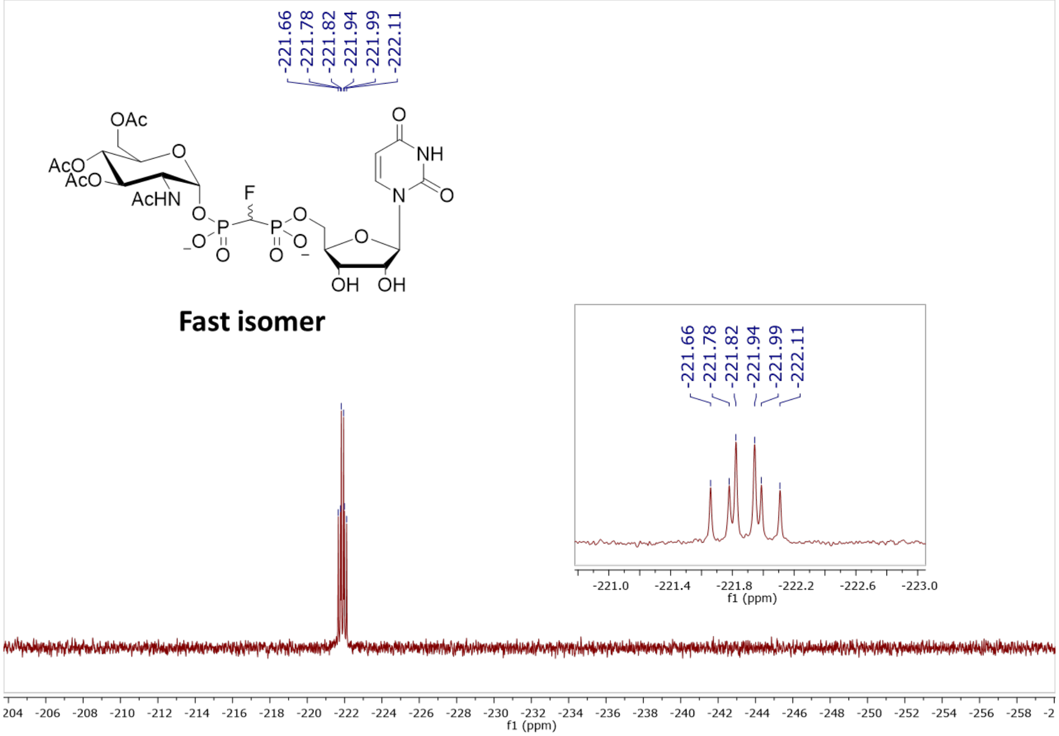

**Figure S125.** $^{19}$F NMR (376 MHz, $D_2O$, pH 7.5) of **25c-α1**, (*S*)-CHF isomer

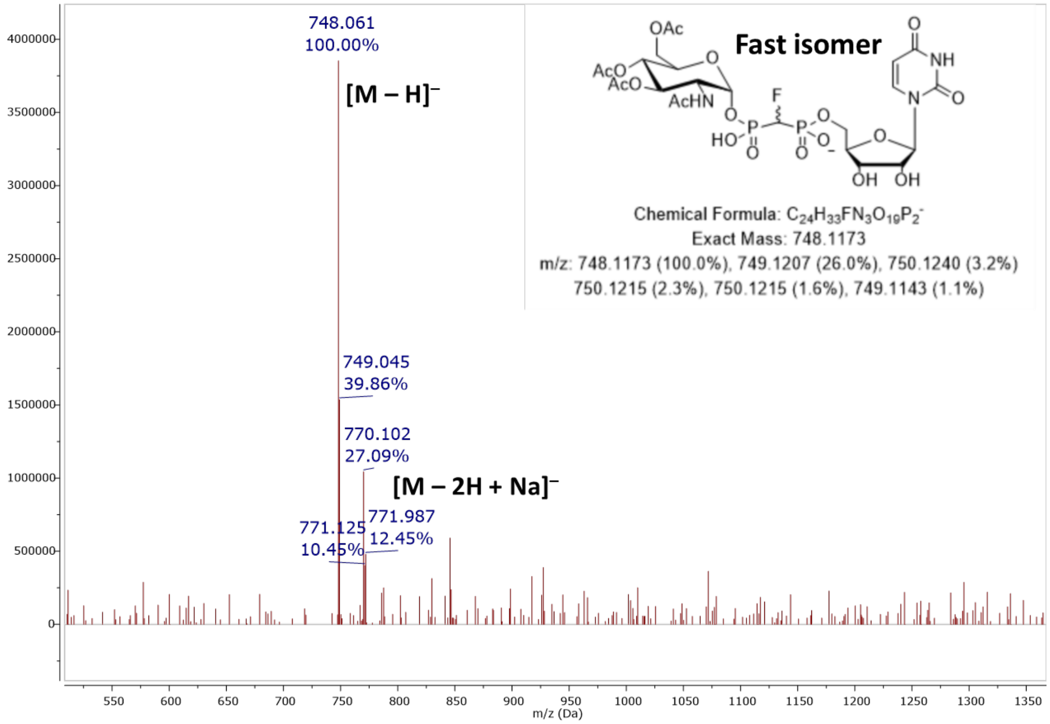

**Figure S126.** MS (ESI): [M – H]⁻ of **25c-α1**, (*S*)-CHF isomer

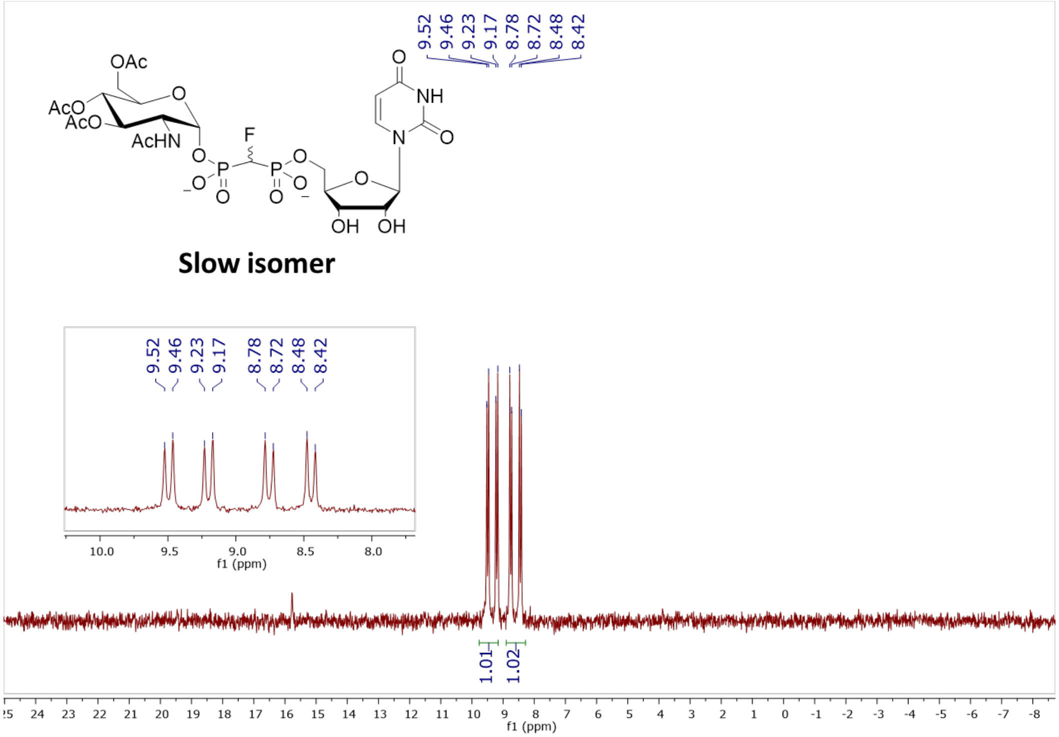

**Figure S127.** $^{31}$P NMR (202 MHz, $D_2O$, pH 7.5) of **25c-α2**, (*R*)-CHF isomer

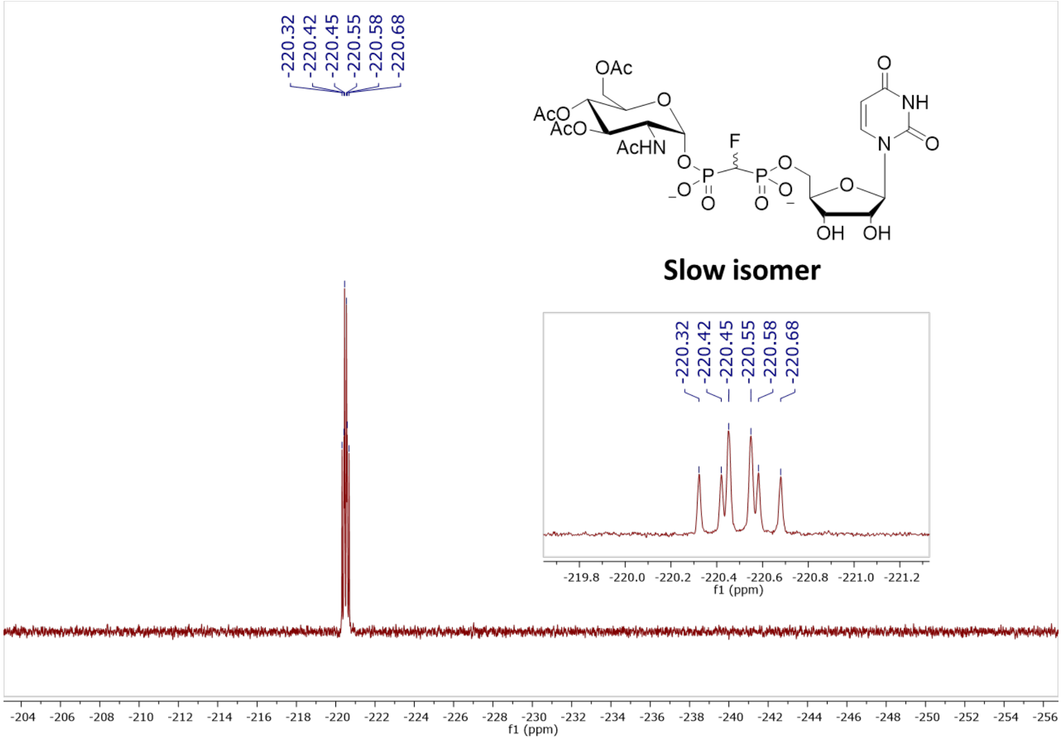

**Figure S128.** $^{19}F$ NMR (470 MHz, $D_2O$, pH 7.5) of **25c-α2**, (*R*)-CHF isomer

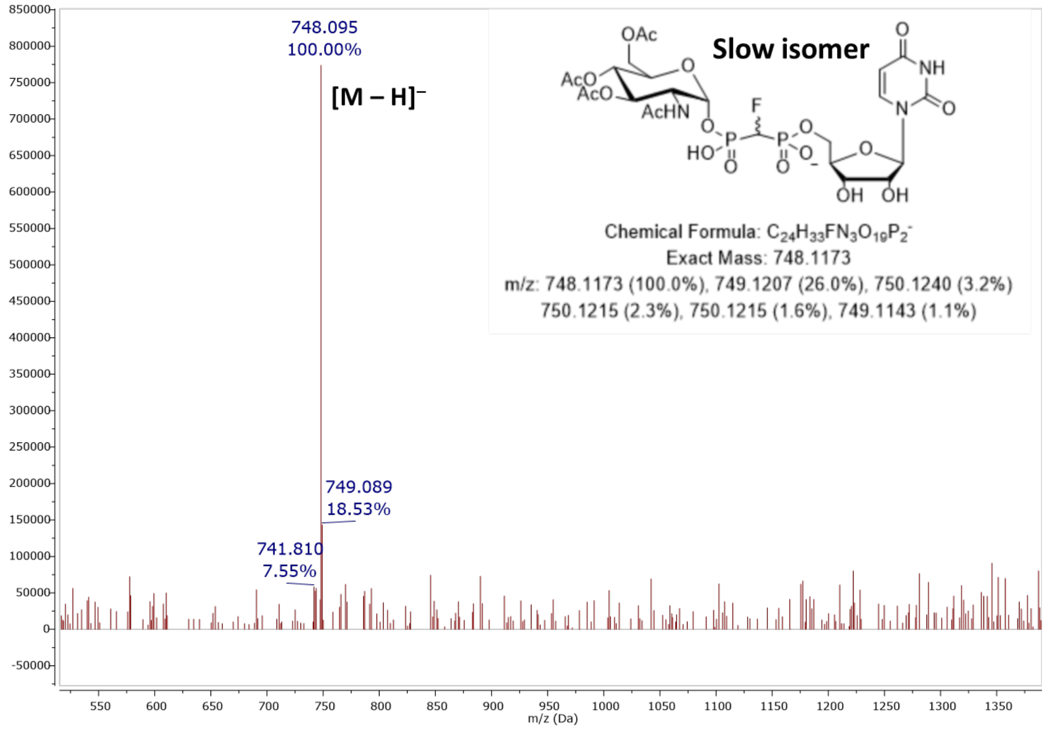

**Figure S129.** MS (ESI): [M – H]⁻ of **25c-α2**, (*R*)-CHF isomer

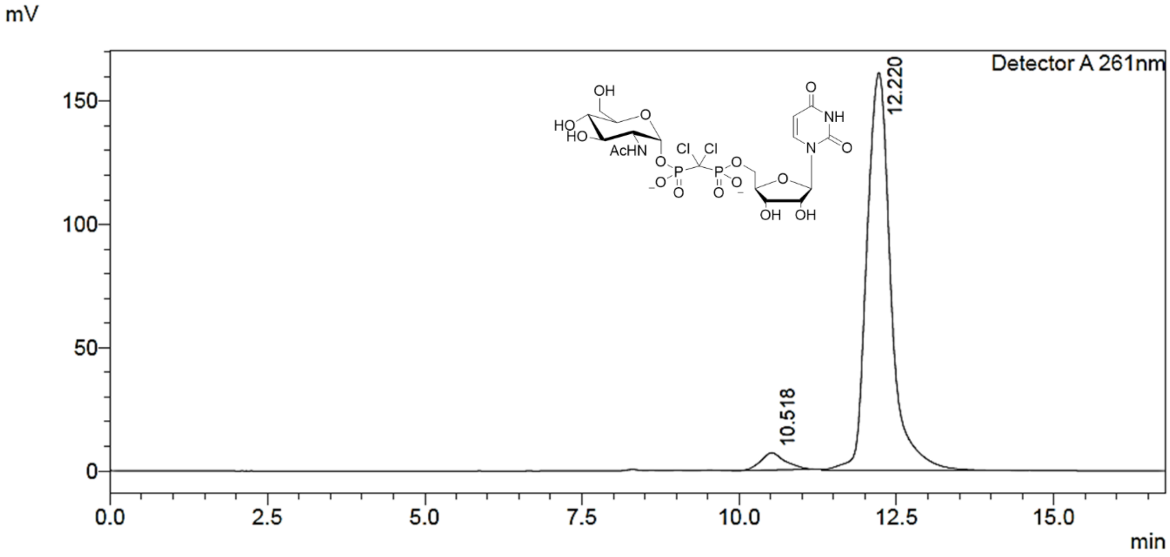

**Figure S130.** Preparative RP-HPLC purification of **2**

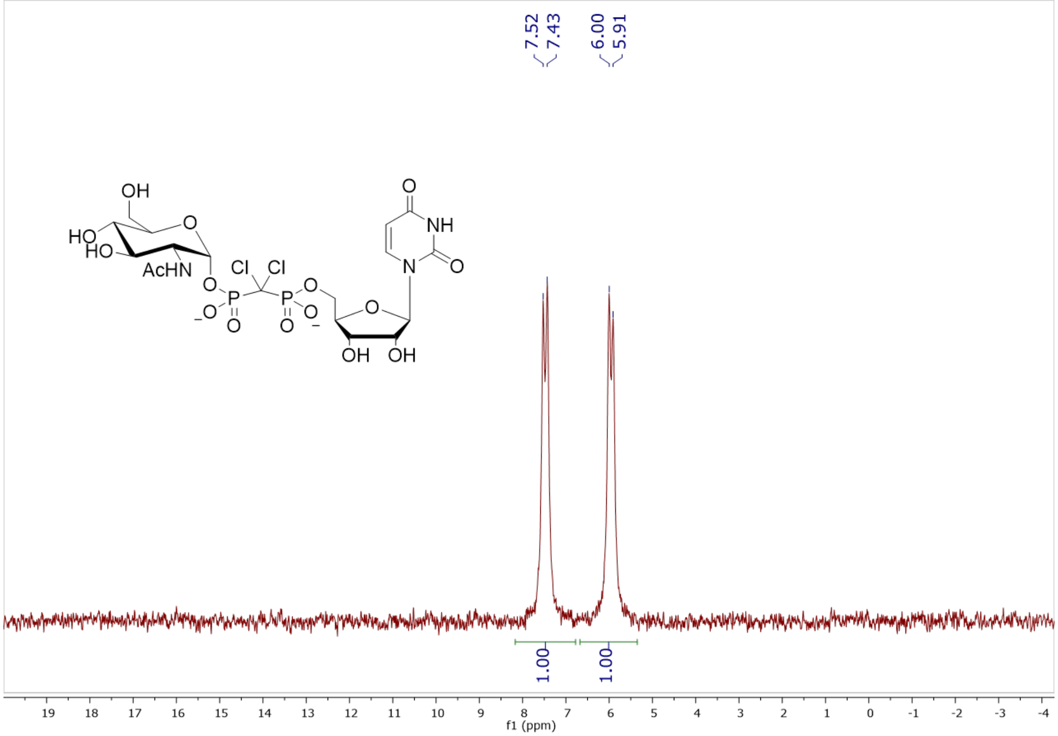

**Figure S131.** $^{31}$P NMR (162 MHz, $D_2O$, pH 7.5) of **2**

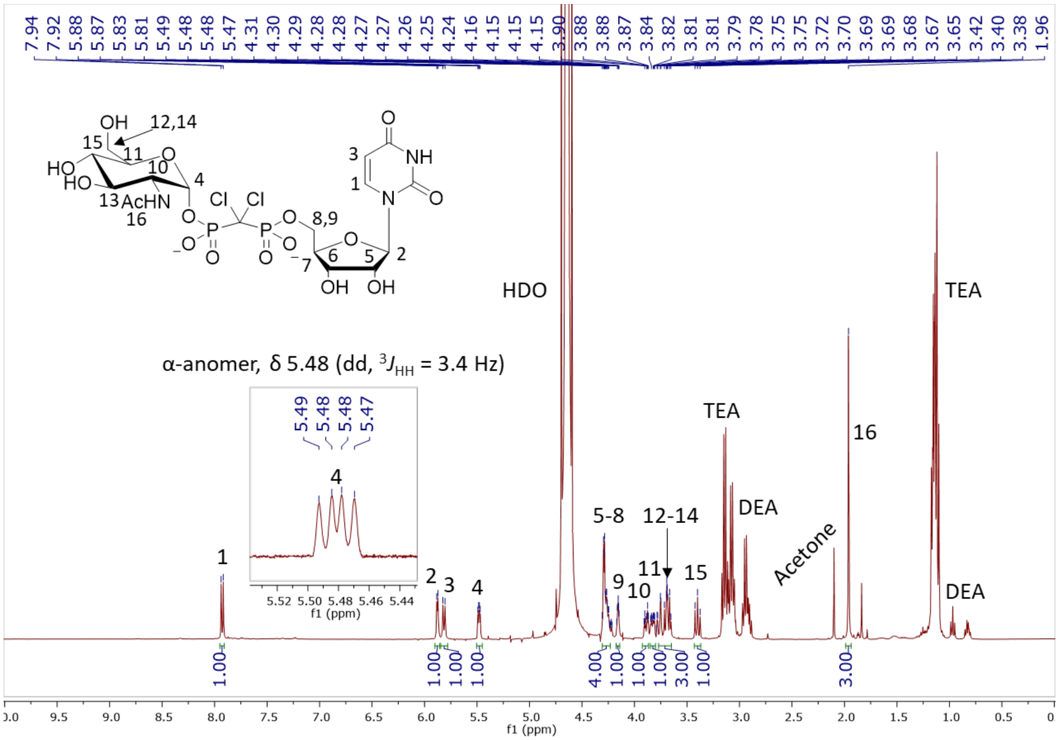

**Figure S132.** $^1H$ NMR (400 MHz, $D_2O$) of **2**

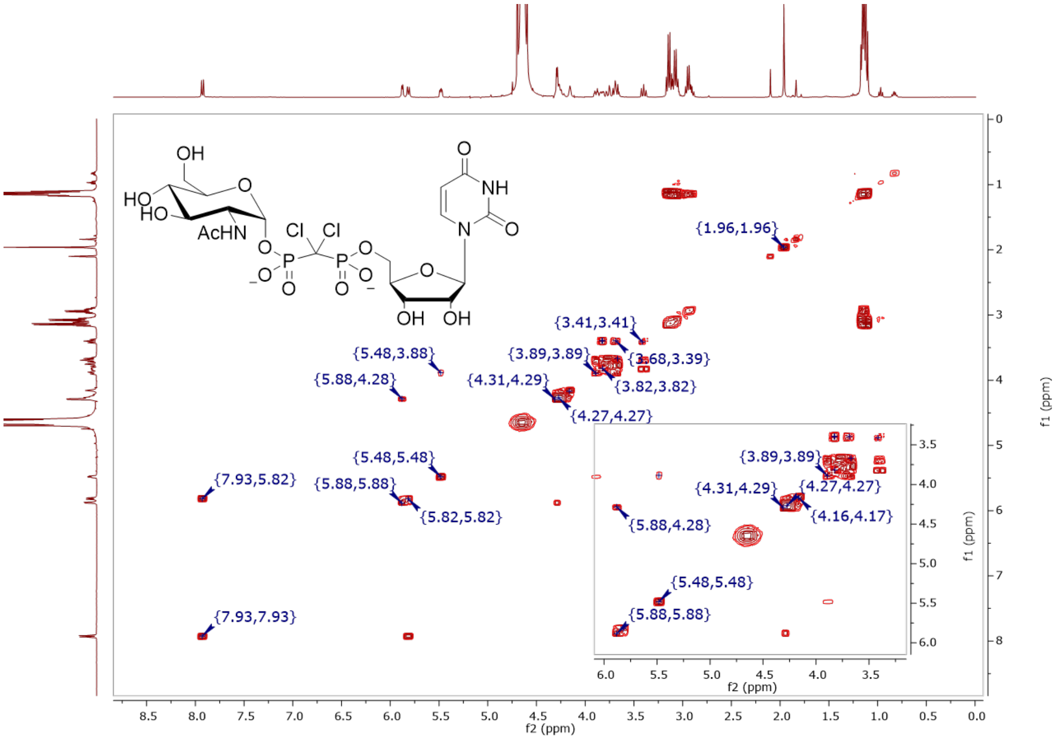

**Figure S133.** COSY (400 MHz, $D_2O$) of **2**

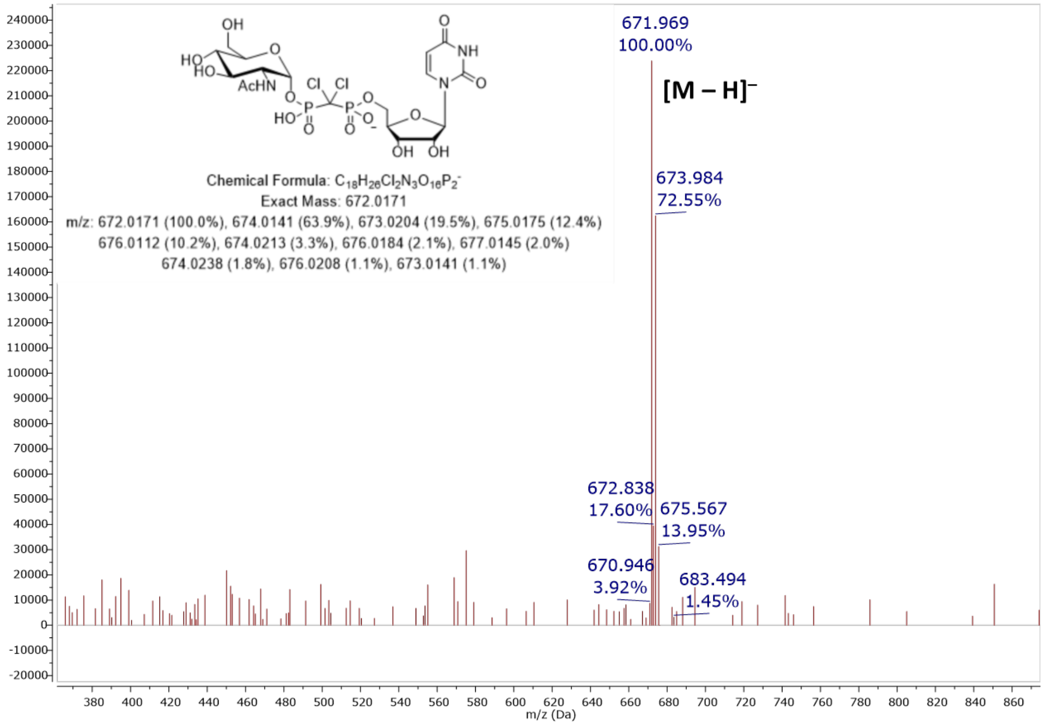

**Figure S134.** MS (ESI): $[M – H]^–$ of **2**

#### Elemental Composition Report

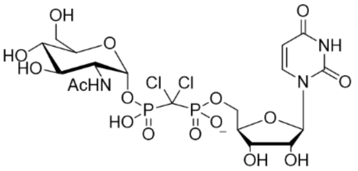

##### Single Mass Analysis

Tolerance = 5.0 PPM / DBE: min = -1.5, max = 100.0
Element prediction: Off
Number of isotope peaks used for i-FIT = 3

Monoisotopic Mass, Even Electron Ions
11 formula(e) evaluated with 1 results within limits (up to 50 closest results for each mass)
Elements Used:
C: 0-100 H: 0-100 N: 3-3 O: 15-17 P: 2-2 Cl: 2-2
Pouya Haratipour, USCMCK19
Synapt_26480a 24 (0.482) Cm (23:24-13:16)

1: TOF MS ES-
3.46e+005

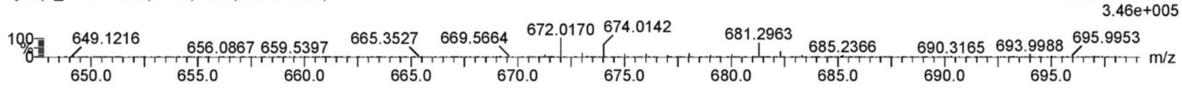

| | | | | | | | | |
|---|---|---|---|---|---|---|---|---|
| Minimum: | | | | -1.5 | | | | |
| Maximum: | | 5.0 | 5.0 | 100.0 | | | | |
| Mass | Calc. Mass | mDa | PPM | DBE | i-FIT | Norm | Conf(%) | Formula |
| 672.0170 | 672.0165 | 0.5 | 0.7 | 7.5 | 229.9 | n/a | n/a | C18 H26 N3 O16 P2 Cl2 |

**Figure S135.** Elemental composition report from HRMS (ESI-TOF) $[M – H]^–$ of **2**

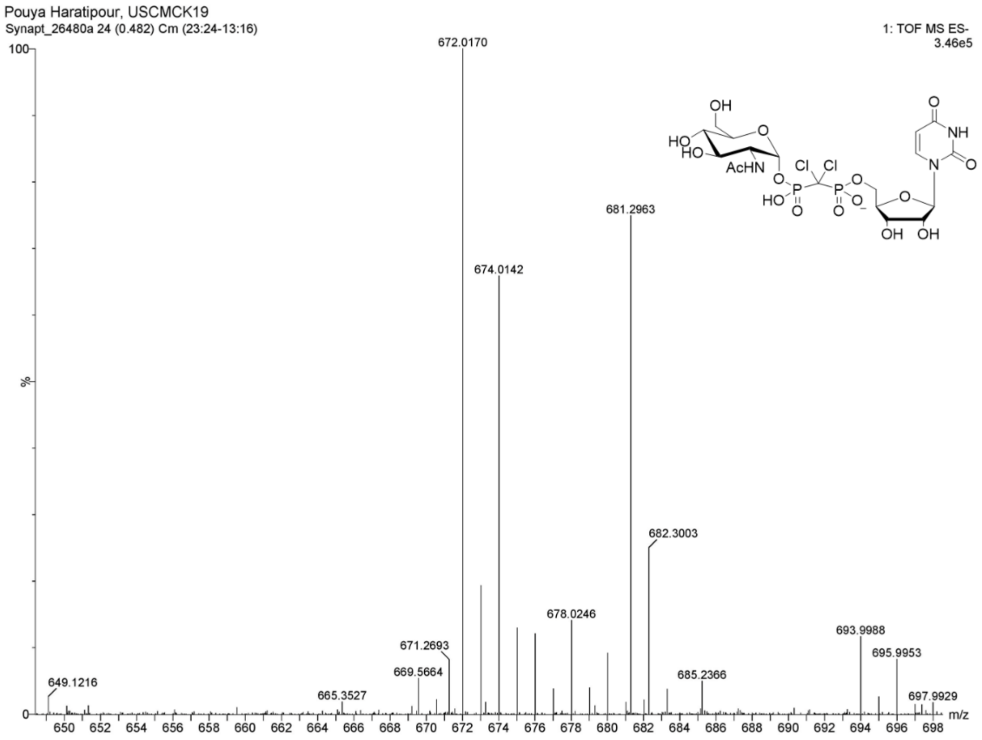

**Figure S136.** HRMS (ESI-TOF): [M – H]$^-$ of **2**

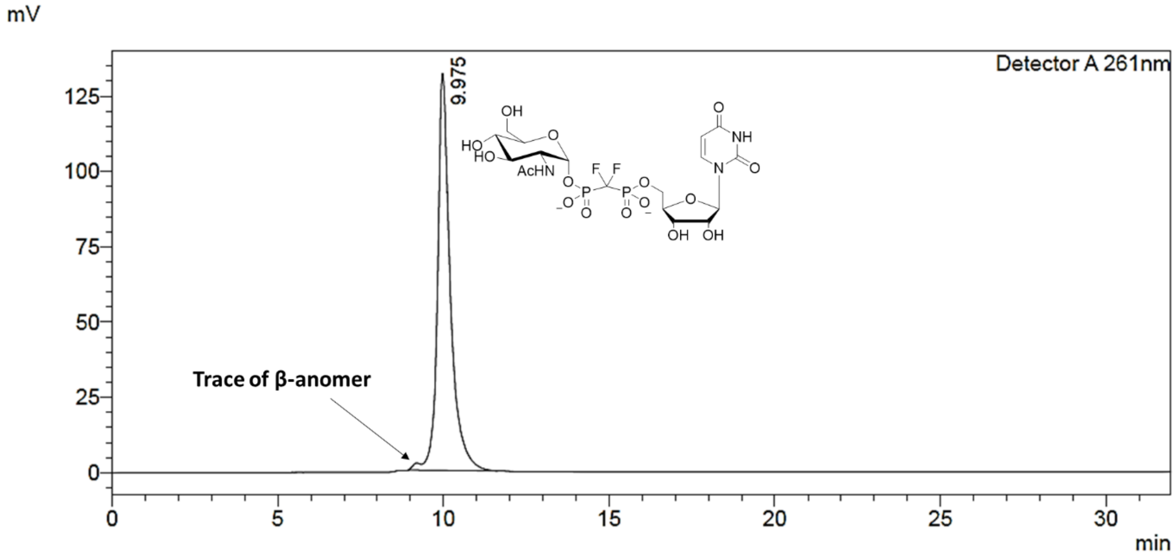

**Figure S137.** Preparative RP-HPLC purification of **4**

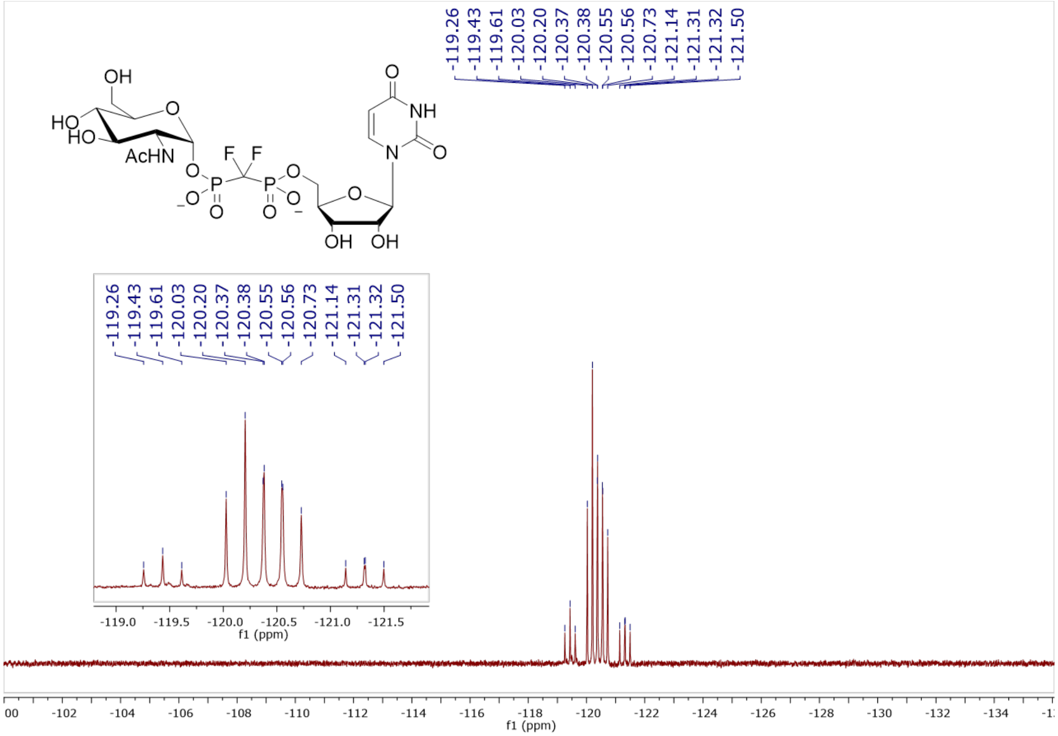

**Figure S138.** $^{19}F$ NMR (470 MHz, $D_2O$, pH 7.5) of **4**

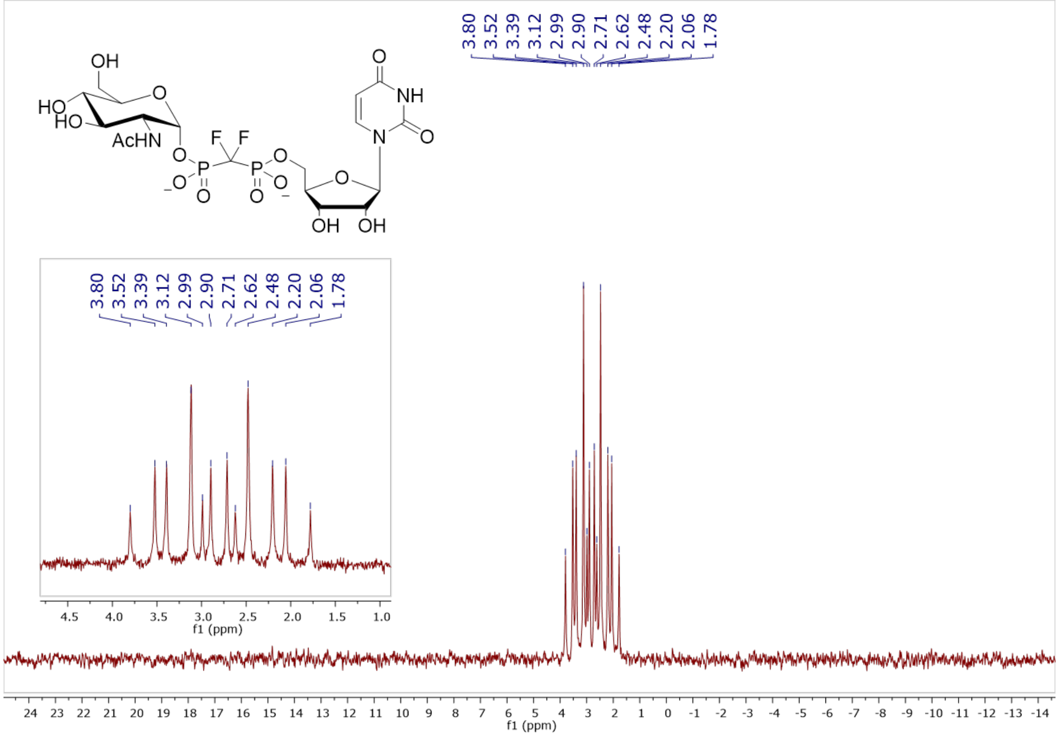

**Figure S139.** $^{31}P$ NMR (202 MHz, $D_2O$, pH 7.5) of **4**

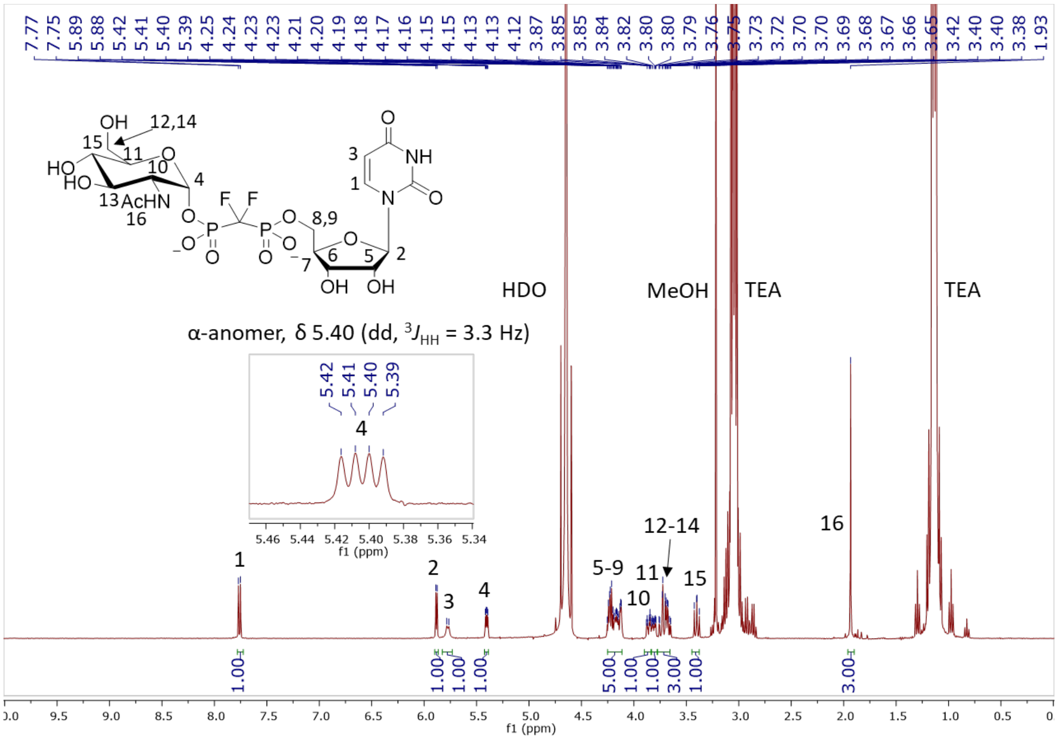

**Figure S140.** $^1$H NMR (400 MHz, $D_2O$) of **4**

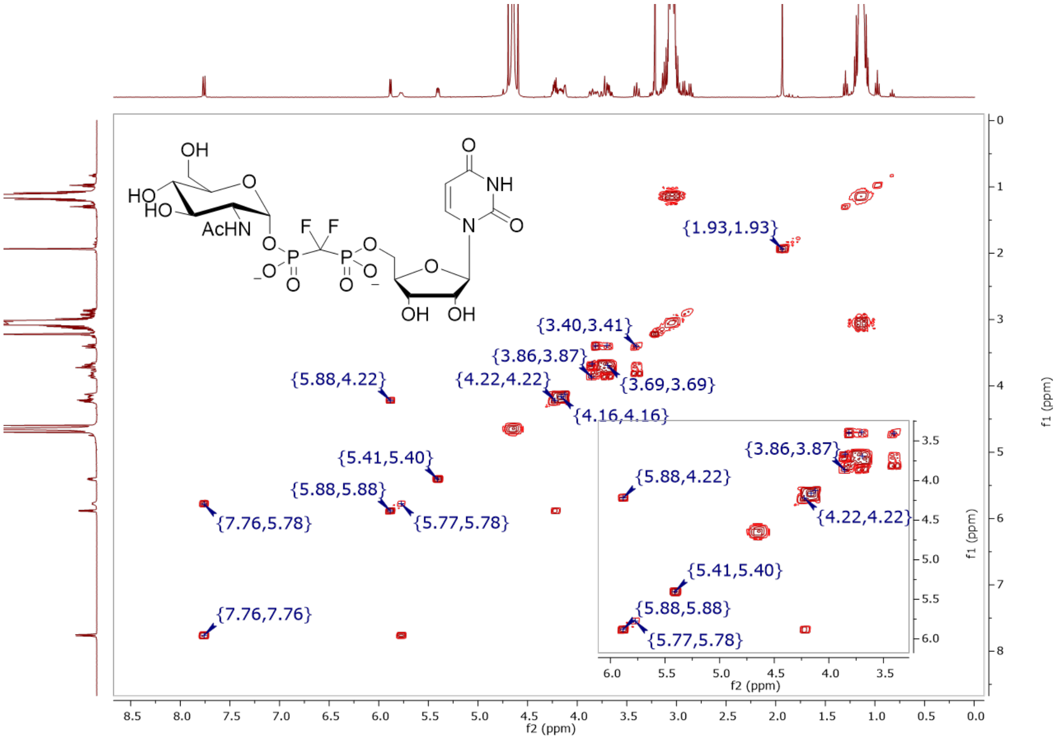

**Figure S141.** COSY (400 MHz, $D_2O$) of **4**

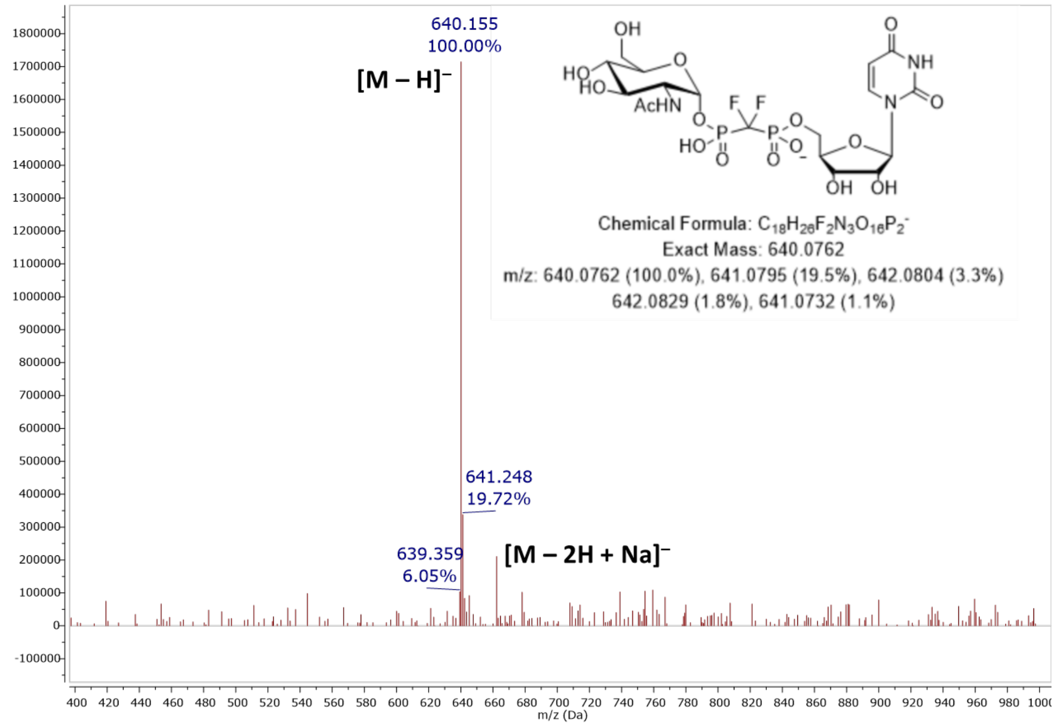

**Figure S142.** MS (ESI): [M – H]⁻ of **4**

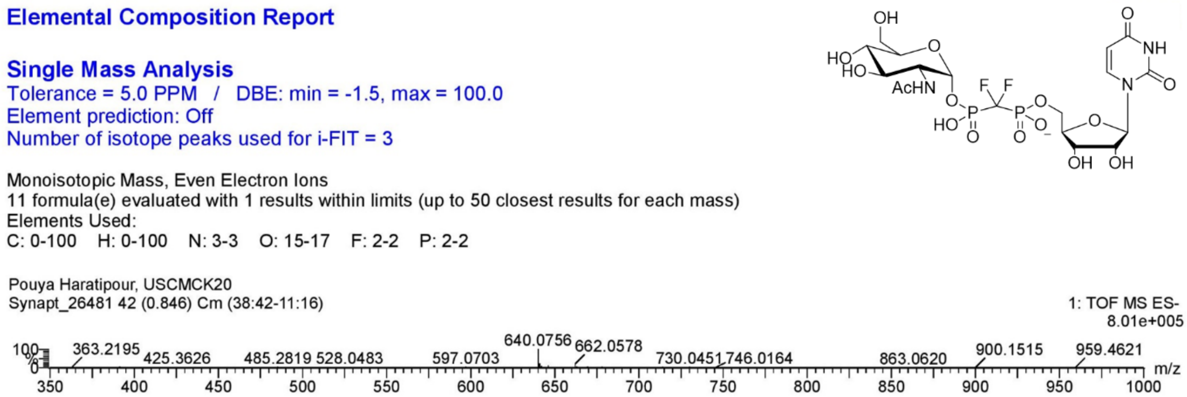

Elemental Composition Report

Single Mass Analysis
Tolerance = 5.0 PPM / DBE: min = -1.5, max = 100.0
Element prediction: Off
Number of isotope peaks used for i-FIT = 3

Monoisotopic Mass, Even Electron Ions
11 formula(e) evaluated with 1 results within limits (up to 50 closest results for each mass)
Elements Used:
C: 0-100 H: 0-100 N: 3-3 O: 15-17 F: 2-2 P: 2-2

Pouya Haratipour, USCMCK20
Synapt_26481 42 (0.846) Cm (38:42-11:16)

1: TOF MS ES-
8.01e+005

| | Mass | Calc. Mass | mDa | PPM | DBE | i-FIT | Norm | Conf(%) | Formula |
|---|---|---|---|---|---|---|---|---|---|
| Minimum: | | | | | -1.5 | | | | |
| Maximum: | | | 5.0 | 5.0 | 100.0 | | | | |
| | 640.0756 | 640.0756 | 0.0 | 0.0 | 7.5 | 470.8 | n/a | n/a | C18 H26 N3 O16 F2 P2 |

**Figure S143.** Elemental composition report from HRMS (ESI-TOF) [M – H]⁻ of **4**

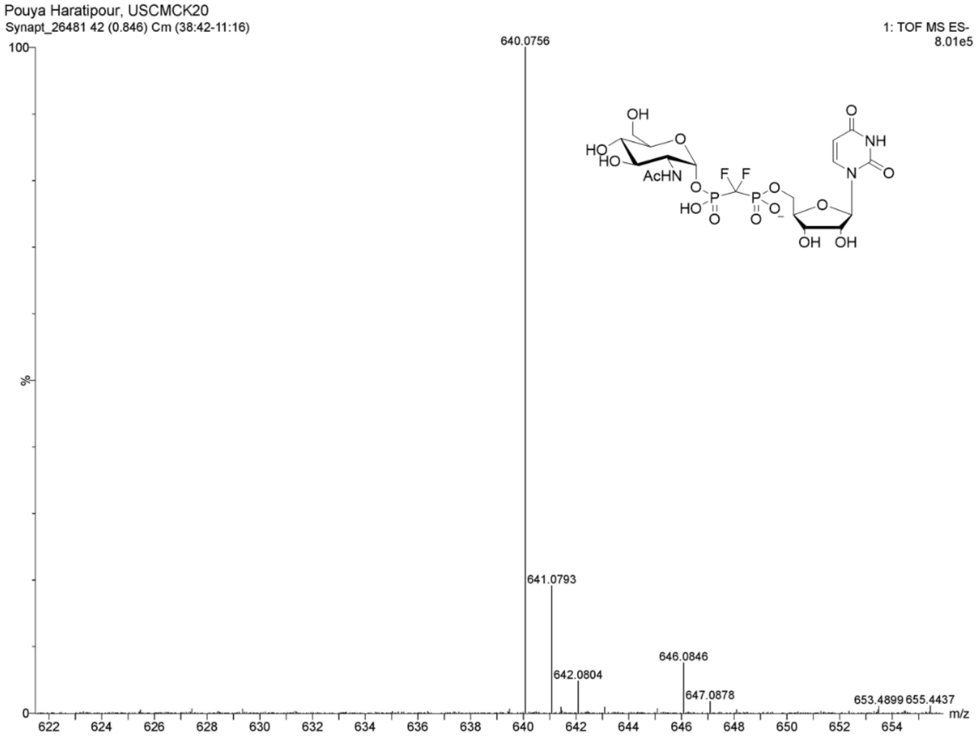

**Figure S144.** HRMS (ESI-TOF): [M – H]⁻ of **4**

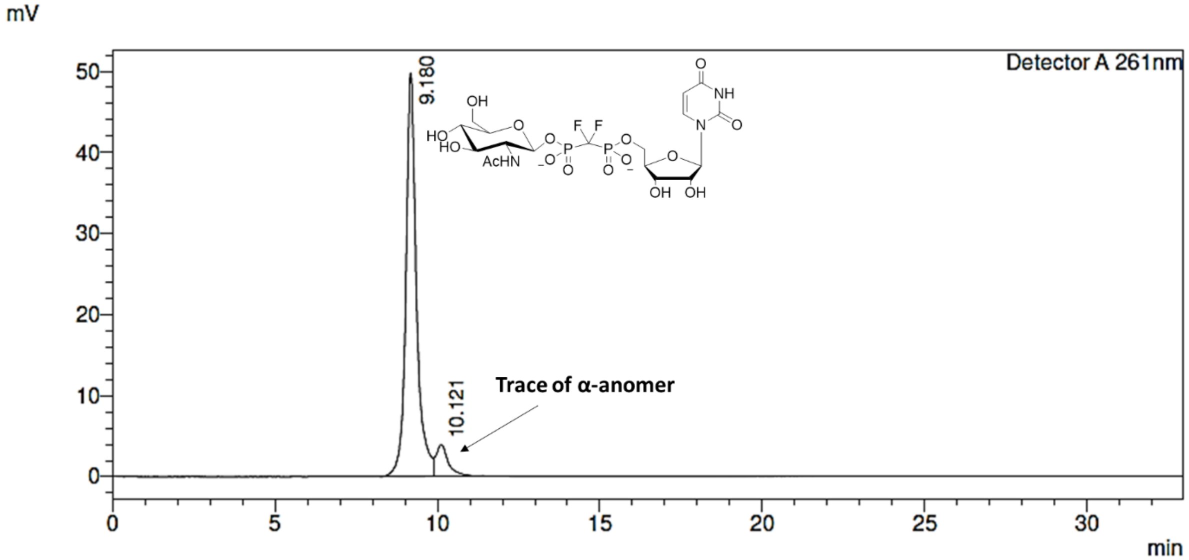

**Figure S145.** Preparative RP-HPLC purification of **5**

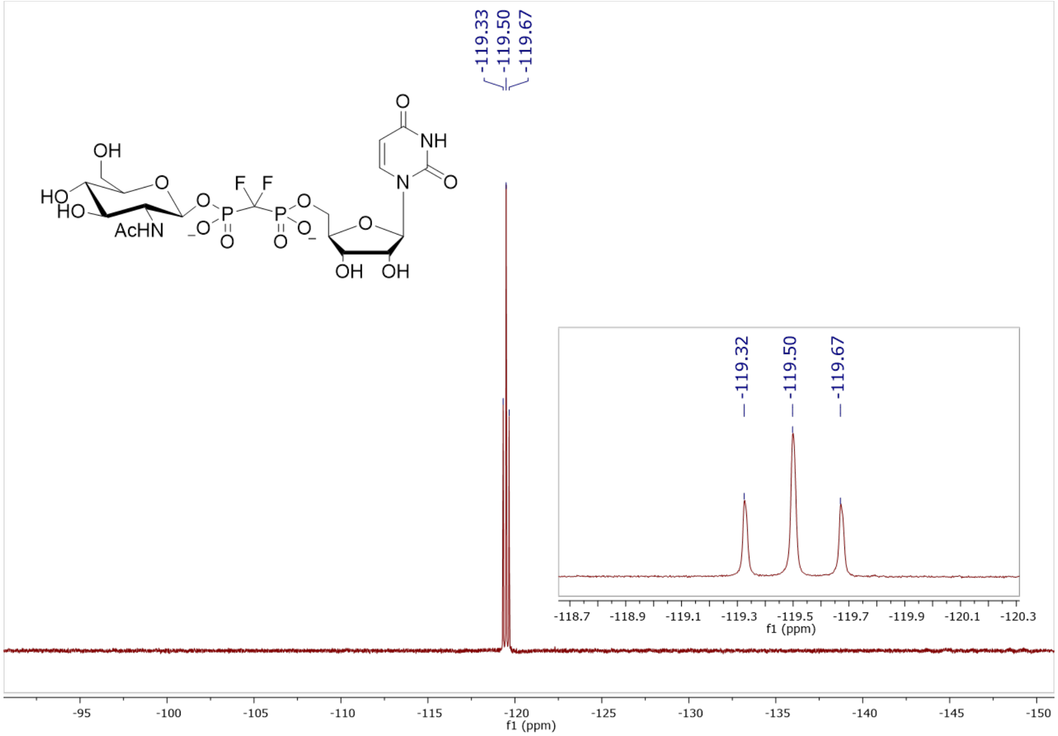

**Figure S146.** $^{19}F$ NMR (470 MHz, $D_2O$, pH 7.5) of **5**

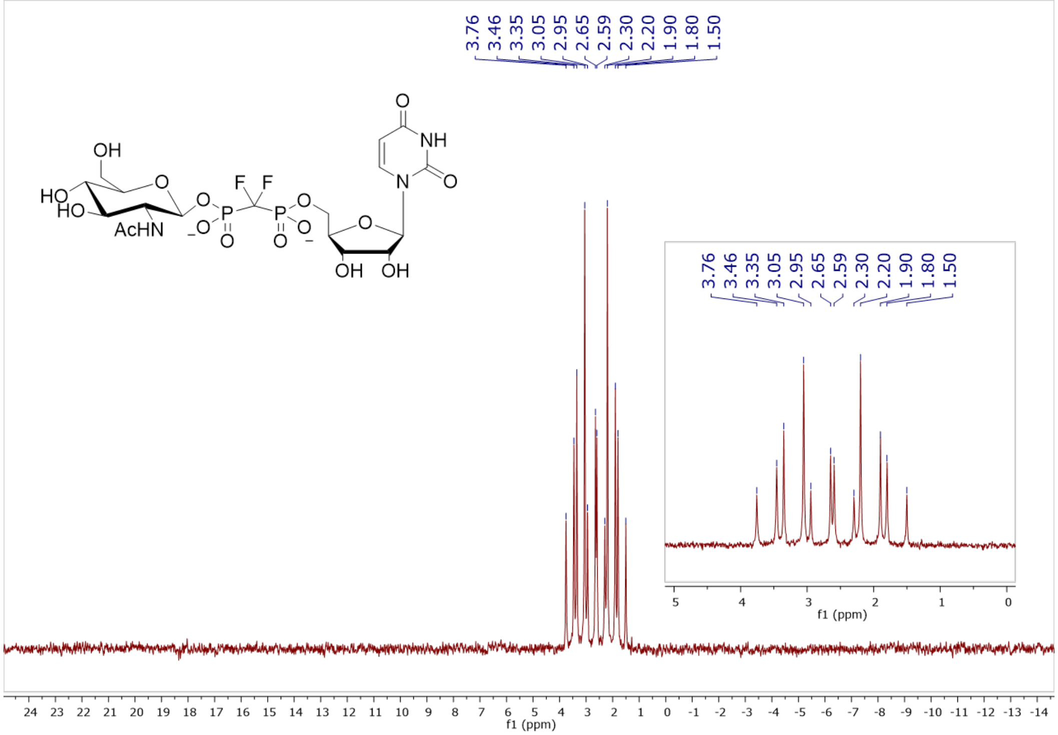

**Figure S147.** $^{31}P$ NMR (202 MHz, $D_2O$, pH 7.5) of **5**

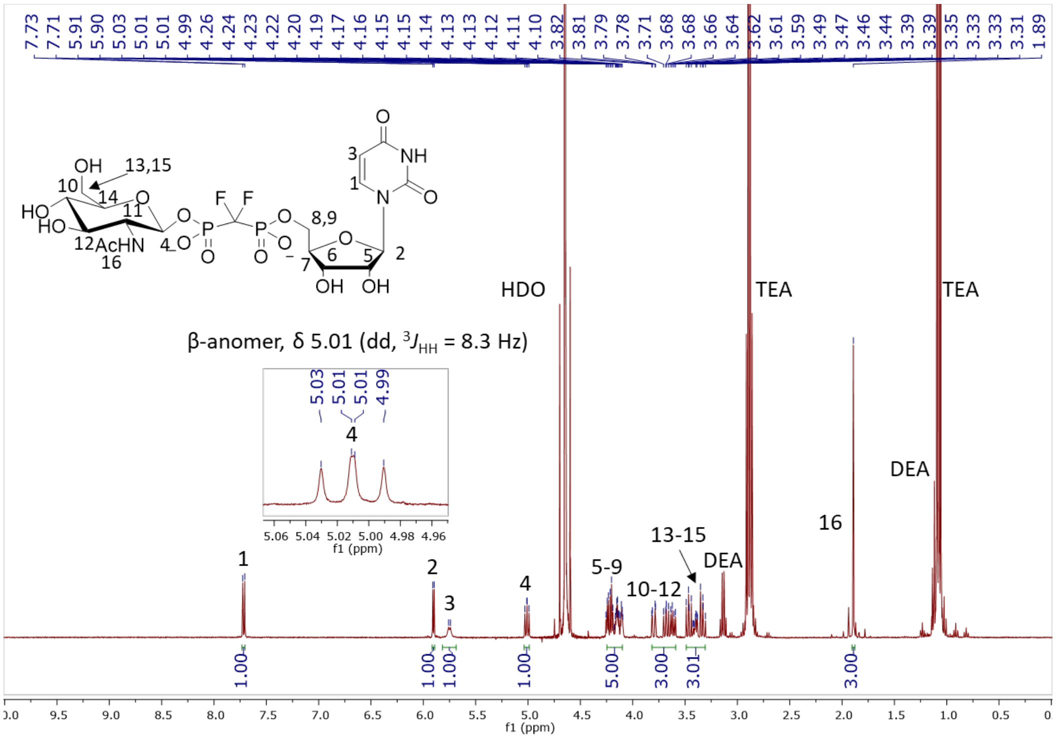

**Figure S148.** $^{1}H$ NMR (500 MHz, $D_2O$) of **5**

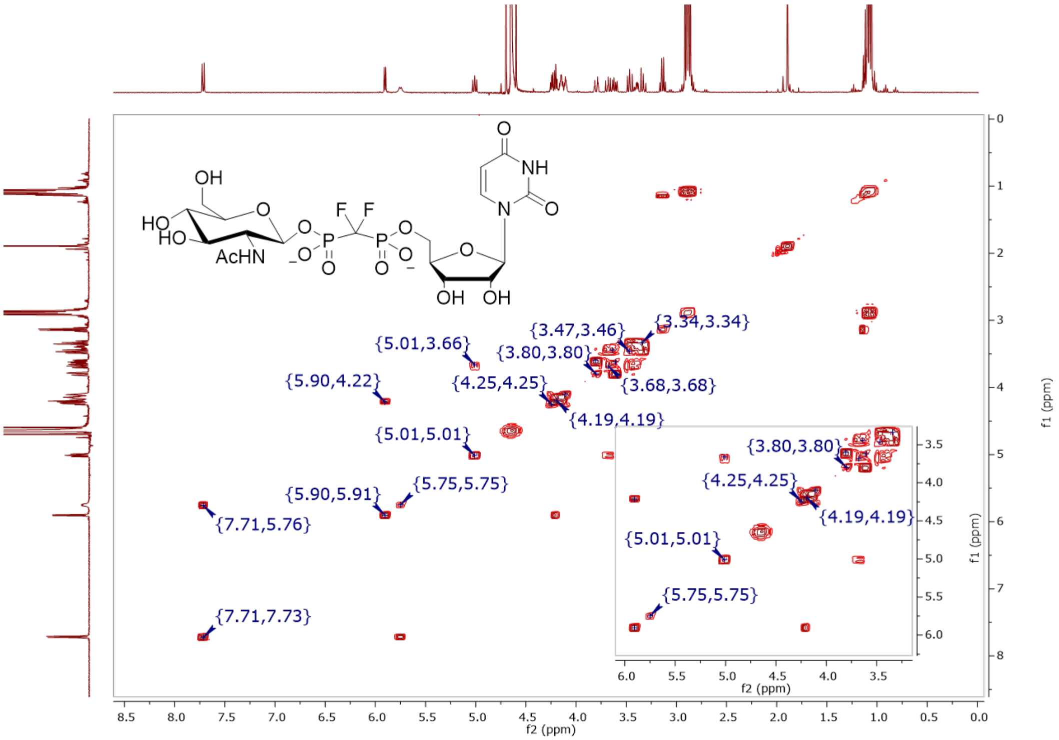

**Figure S149.** COSY (400 MHz, $D_2O$) of **5**

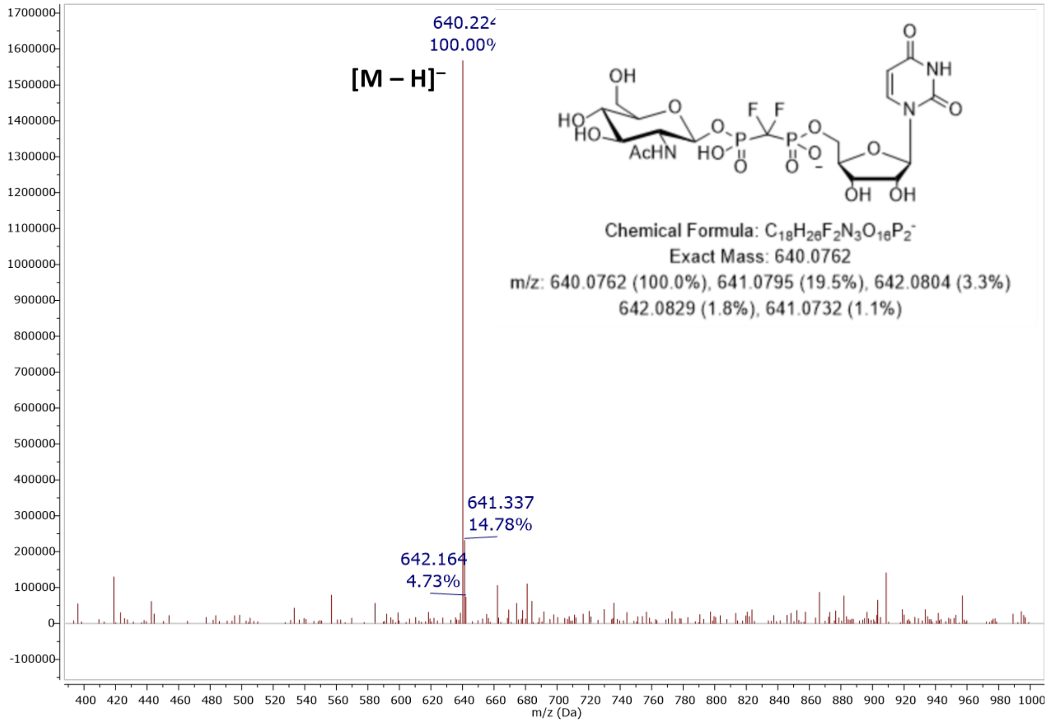

**Figure S150.** MS (ESI): $[M - H]^-$ of **5**

#### Elemental Composition Report

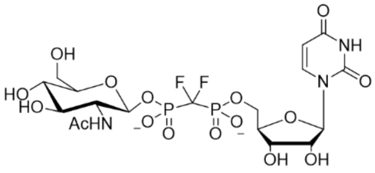

##### Single Mass Analysis

Tolerance = 5.0 PPM / DBE: min = -1.5, max = 100.0
Element prediction: Off
Number of isotope peaks used for i-FIT = 3

Monoisotopic Mass, Even Electron Ions
11 formula(e) evaluated with 1 results within limits (up to 50 closest results for each mass)
Elements Used:
C: 0-100 H: 0-100 N: 3-3 O: 15-17 P: 2-2 F: 2-2

Pouya Haratipour, USCMCK21
Synapt_26482_LC 40 (0.393) Cm (40-62)
1: TOF MS ES-
1.73e+005

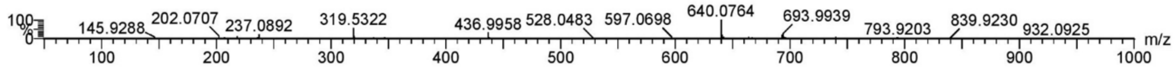

| Mass | Calc. Mass | mDa | PPM | DBE | i-FIT | Norm | Conf(%) | Formula |
|---|---|---|---|---|---|---|---|---|
| Minimum: | | | | -1.5 | | | | |
| Maximum: | | 5.0 | 5.0 | 100.0 | | | | |
| 640.0764 | 640.0756 | 0.8 | 1.2 | 7.5 | 250.2 | n/a | n/a | C18 H26 N3 O16 P2 F2 |

**Figure S151.** Elemental composition report from HRMS (ESI-TOF) $[M - H]^-$ of **5**

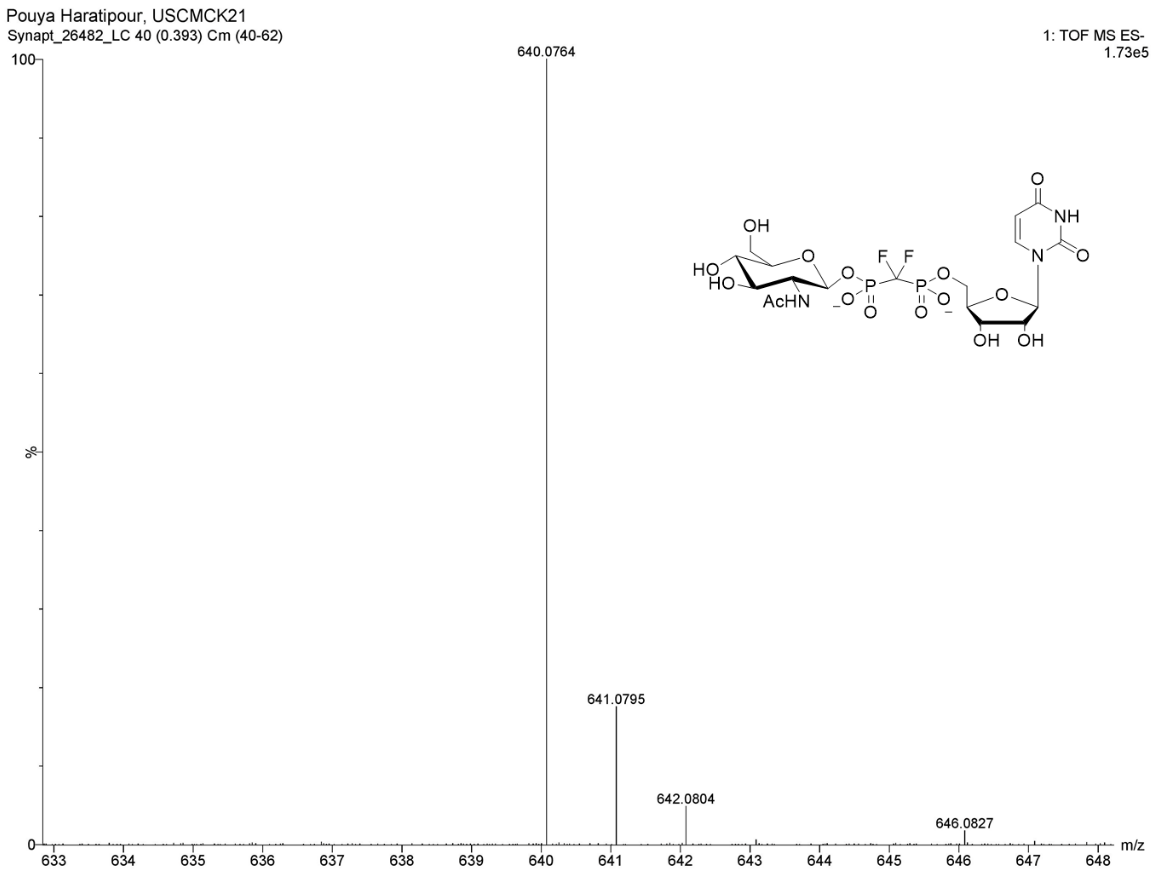

**Figure S152.** HRMS (ESI-TOF): $[M – H]^-$ of **5**

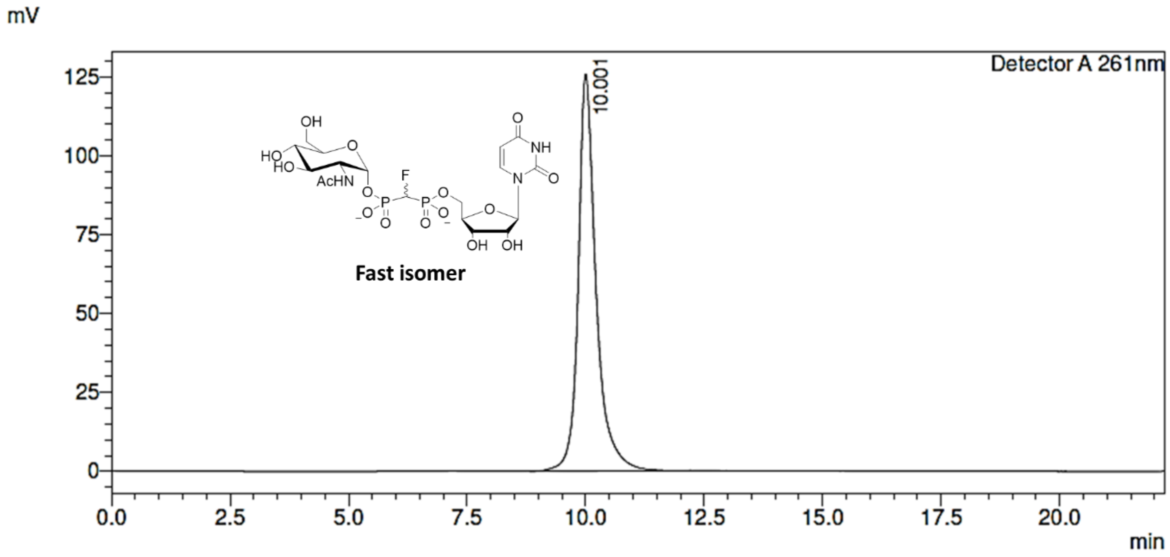

**Figure S153.** Preparative RP-HPLC purification of **7-α1** (*S*)-CHF isomer

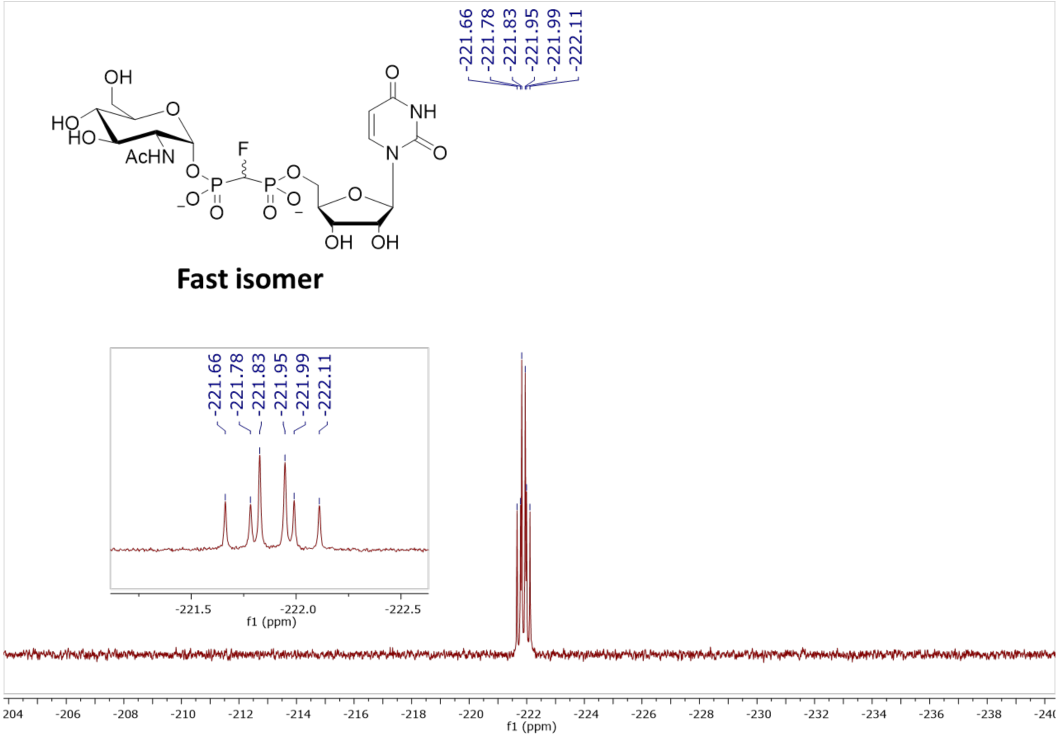

**Figure S154.** $^{19}F$ NMR (376 MHz, $D_2O$, pH 7.5) of **7-α1** (*S*)-CHF isomer

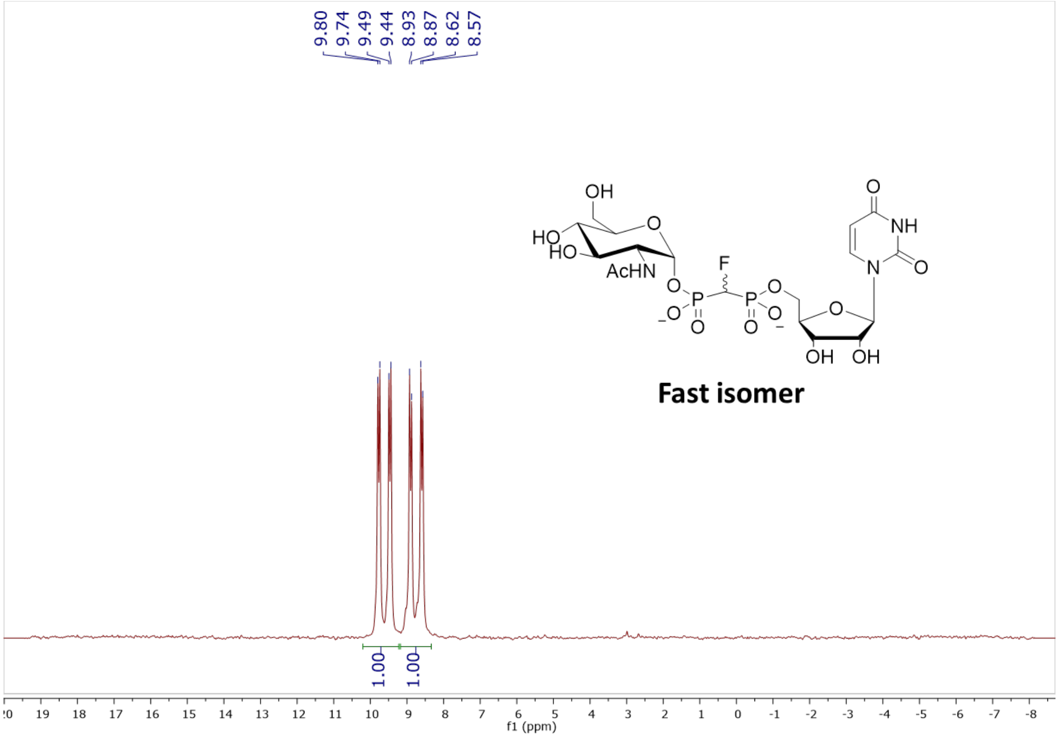

**Figure S155.** $^{31}P$ NMR (202 MHz, $D_2O$, pH 7.5) of **7-α1** (*S*)-CHF isomer

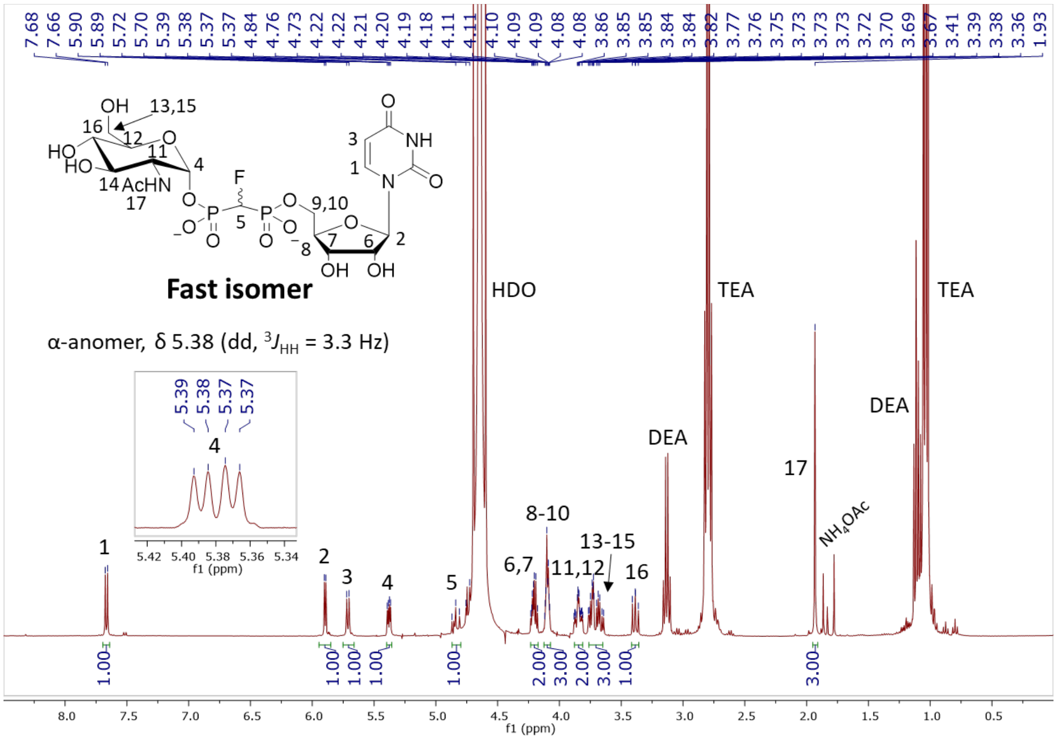

**Figure S156.** $^1$H NMR (400 MHz, $D_2O$) of **7-α1** (*S*)-CHF isomer

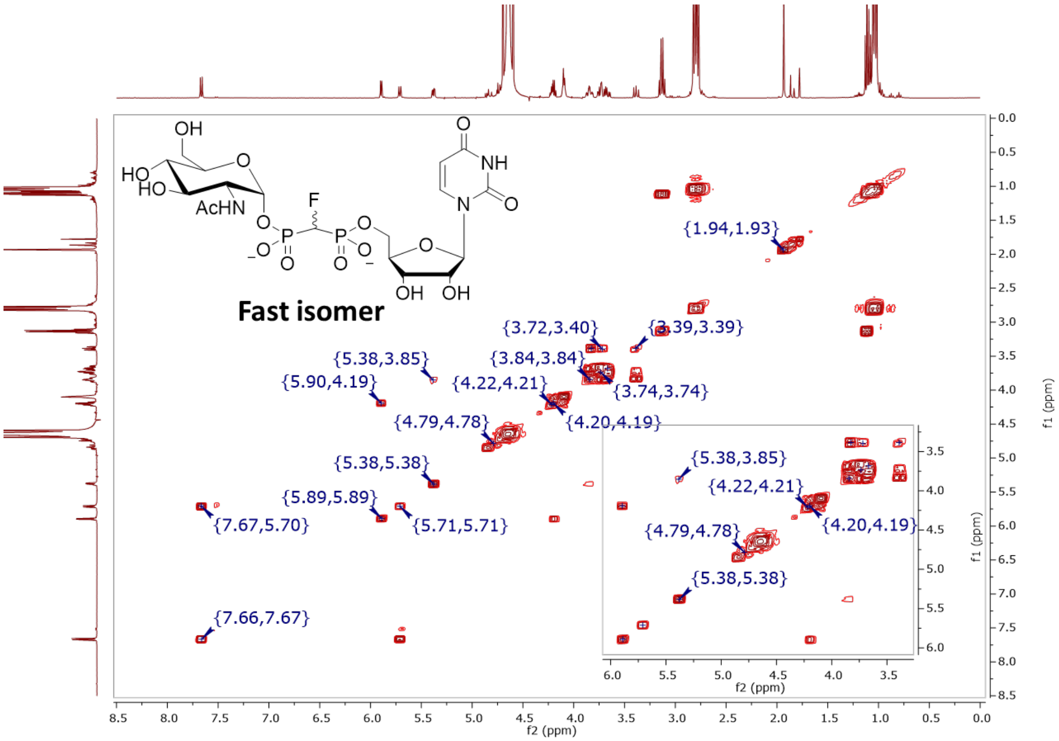

**Figure S157.** COSY (400 MHz, $D_2O$) of **7-α1** (*S*)-CHF isomer

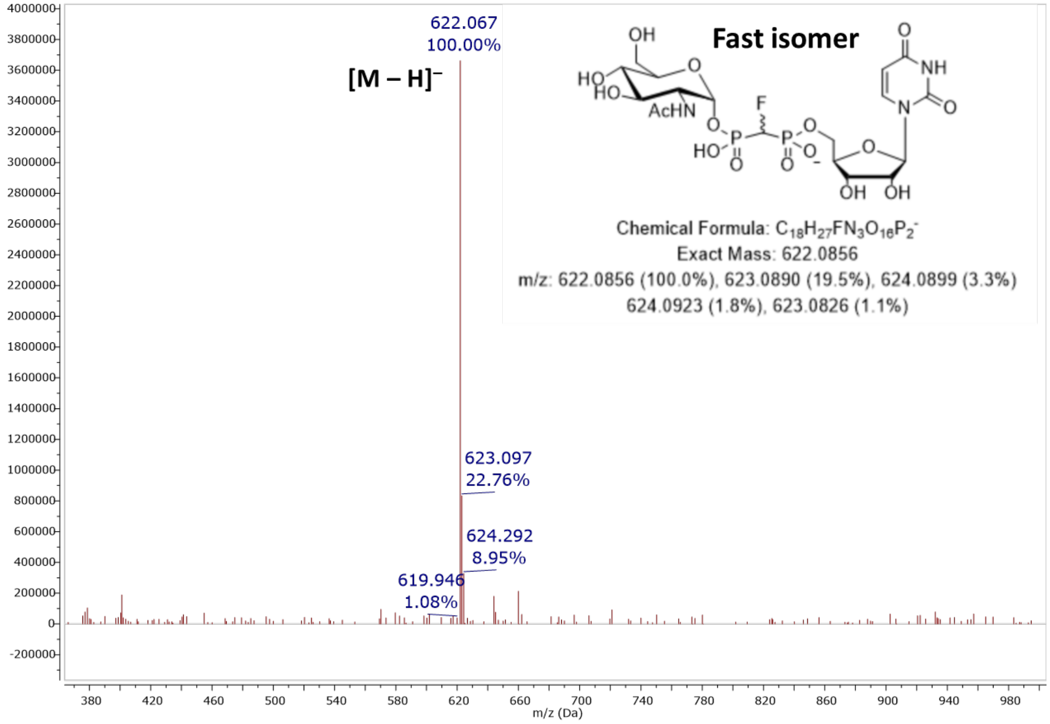

**Figure S158.** MS (ESI): $[M – H]^-$ of **7-α1** (*S*)-CHF isomer

**Elemental Composition Report**

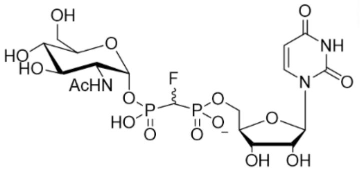

**Single Mass Analysis**
Tolerance = 5.0 PPM / DBE: min = -1.5, max = 50.0
Element prediction: Off
Number of isotope peaks used for i-FIT = 3

Monoisotopic Mass, Even Electron Ions
234 formula(e) evaluated with 2 results within limits (up to 50 closest results for each mass)
Elements Used:
C: 0-60 H: 0-100 N: 2-4 O: 15-17 F: 0-1 P: 1-3
Pouya Haratipour, USCMCK15
Synapt_26240 25 (0.519) Cm (25:27-4:6)

1: TOF MS ES-
3.27e+006

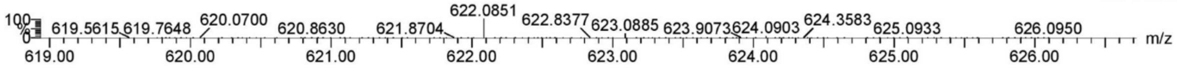

| | | | | | | | | |
|---|---|---|---|---|---|---|---|---|
| Minimum: | | | | -1.5 | | | | |
| Maximum: | | 5.0 | 5.0 | 50.0 | | | | |
| Mass | Calc. Mass | mDa | PPM | DBE | i-FIT | Norm | Conf(%) | Formula |
| 622.0851 | 622.0851 | 0.0 | 0.0 | 7.5 | 667.0 | 0.007 | 99.32 | C18 H27 N3 O16 F P2 |
| | 622.0839 | 1.2 | 1.9 | 11.5 | 671.9 | 4.992 | 0.68 | C21 H26 N3 O15 P2 |

**Figure S159.** Elemental composition report from HRMS (ESI-TOF) $[M – H]^-$ of **7-α1** (*S*)-CHF isomer

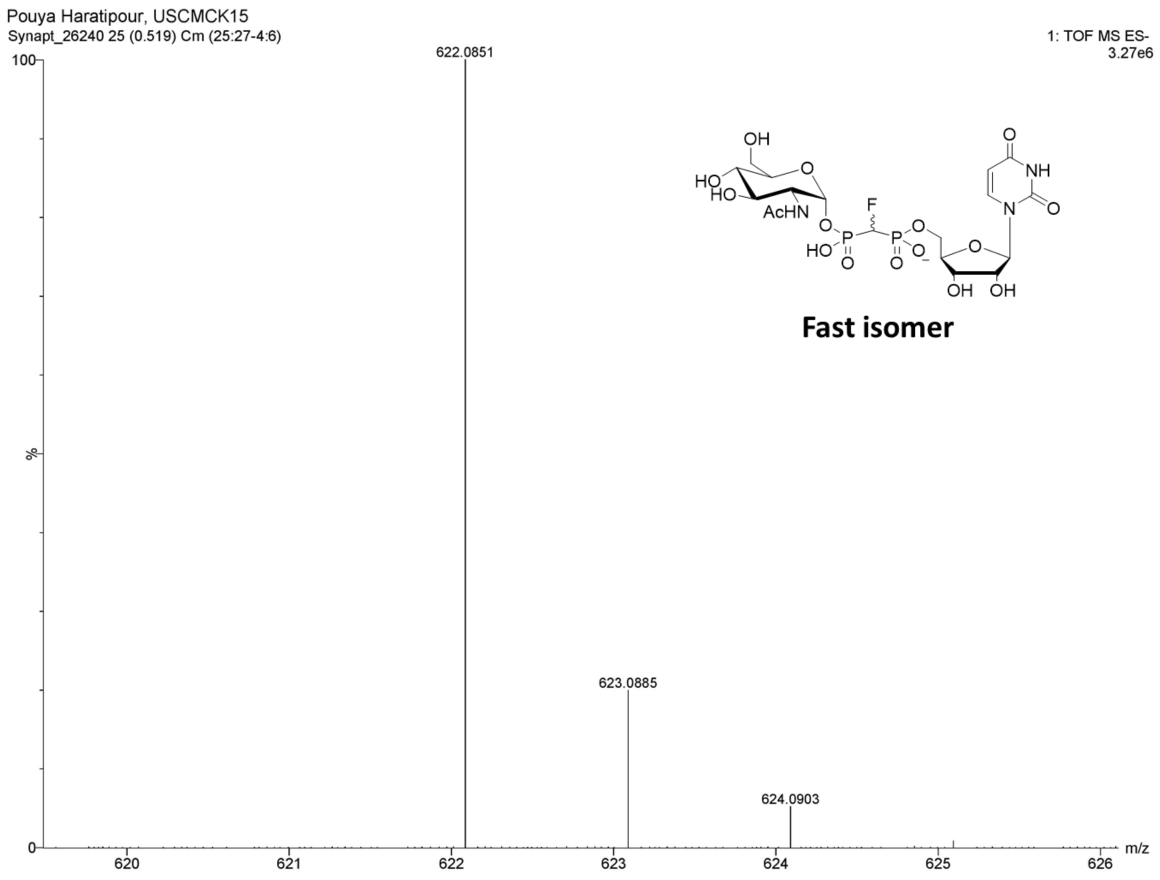

**Figure S160.** HRMS (ESI-TOF): $[M – H]^{-}$ of **7-α1** (*S*)-CHF isomer

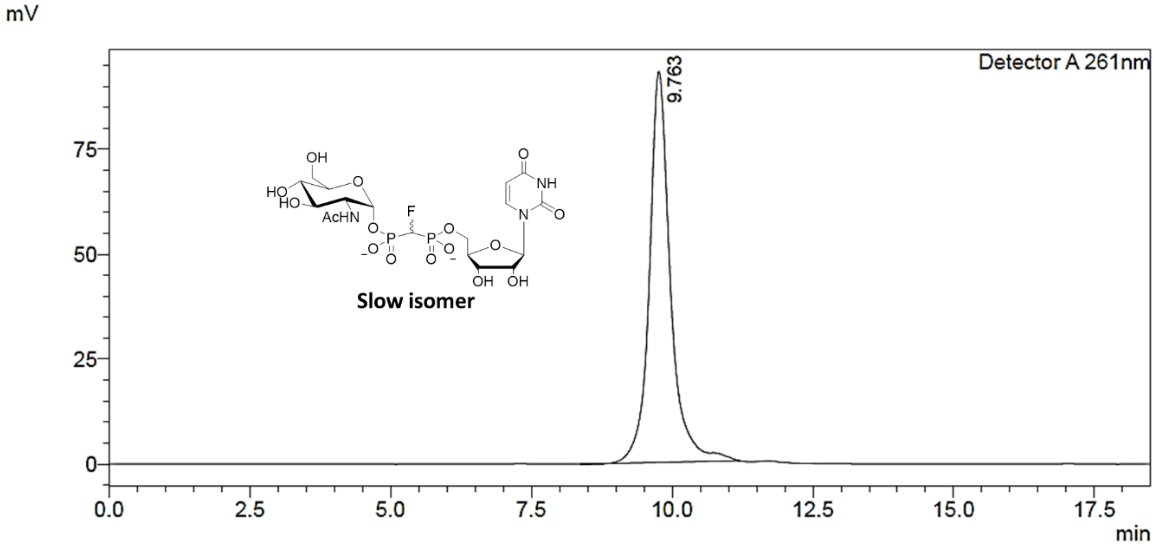

**Figure S161.** Preparative RP-HPLC purification of **8-α2** (*R*)-CHF isomer

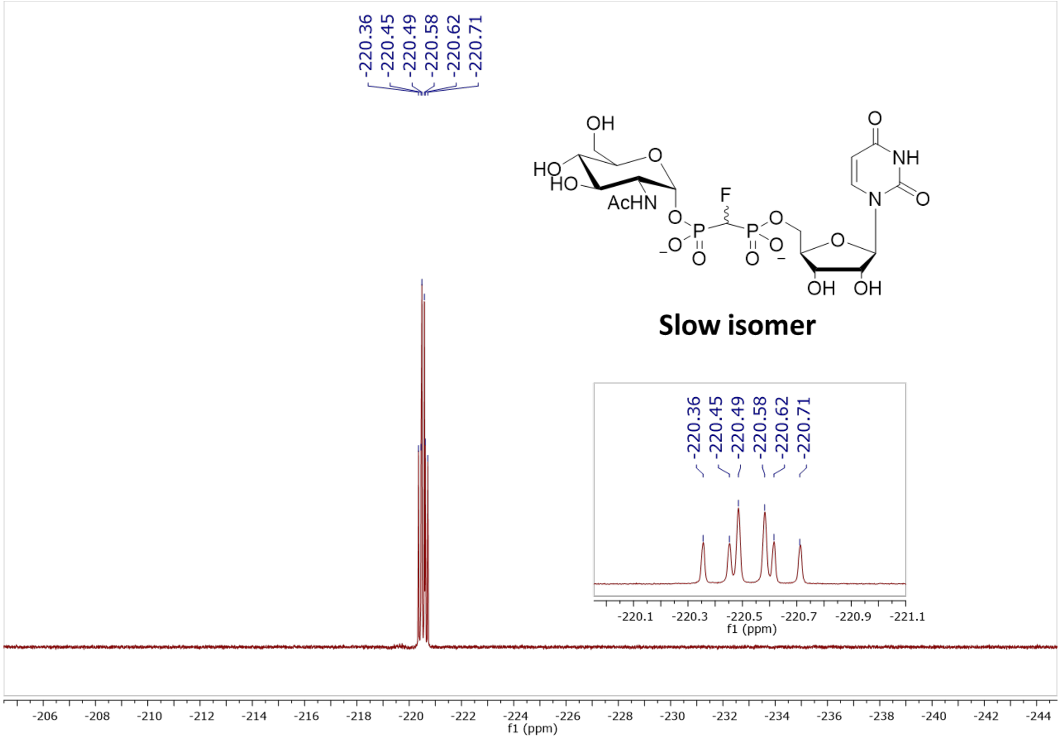

**Figure S162.** $^{19}$F NMR (470 MHz, $D_2O$, pH 7.5) of **8-α2** (*R*)-CHF isomer

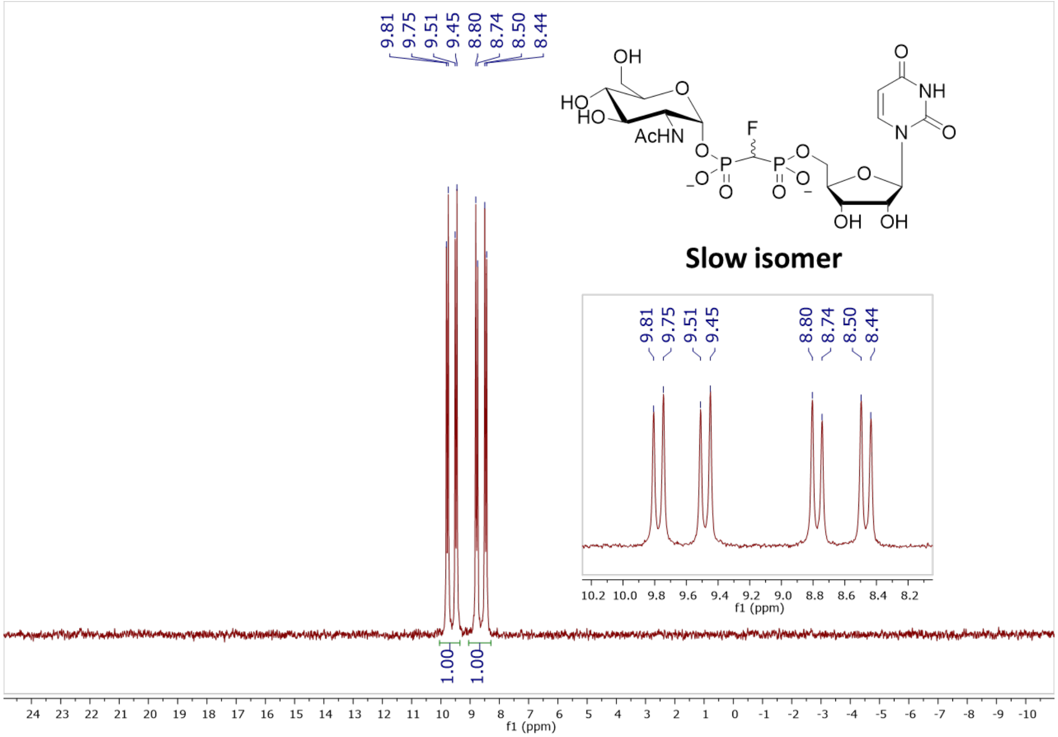

**Figure S163.** $^{31}$P NMR (202 MHz, $D_2O$, pH 7.5) of **8-α2** (*R*)-CHF isomer

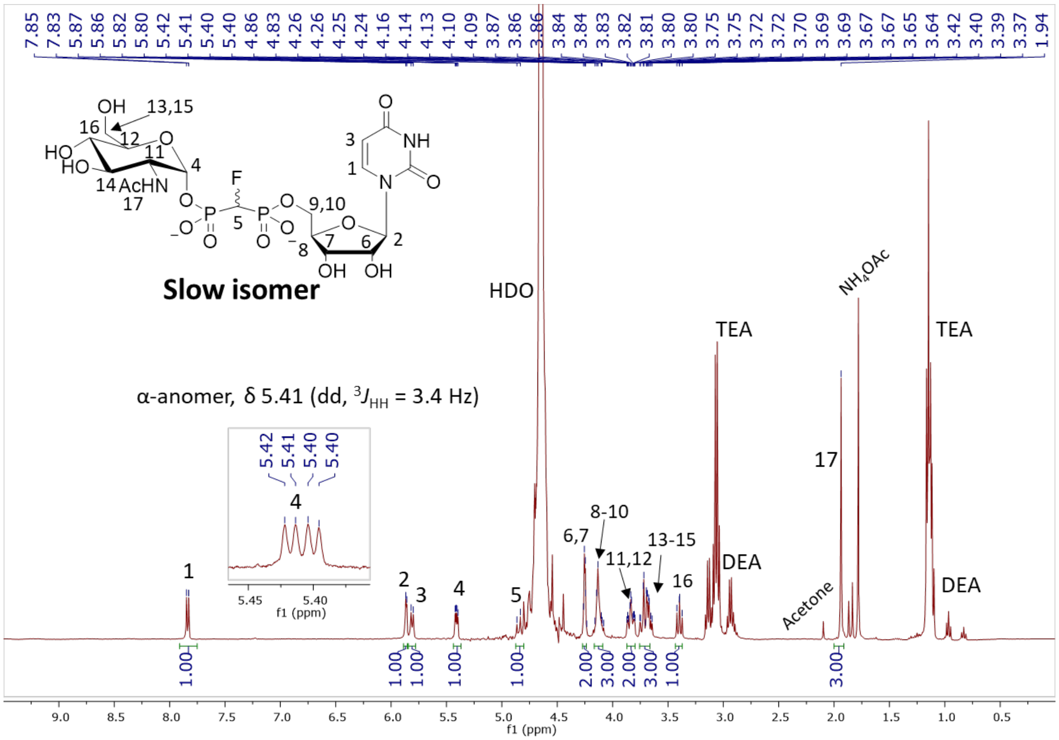

**Figure S164.** $^{1}H$ NMR (400 MHz, $D_2O$) of **8-α2** (*R*)-CHF isomer

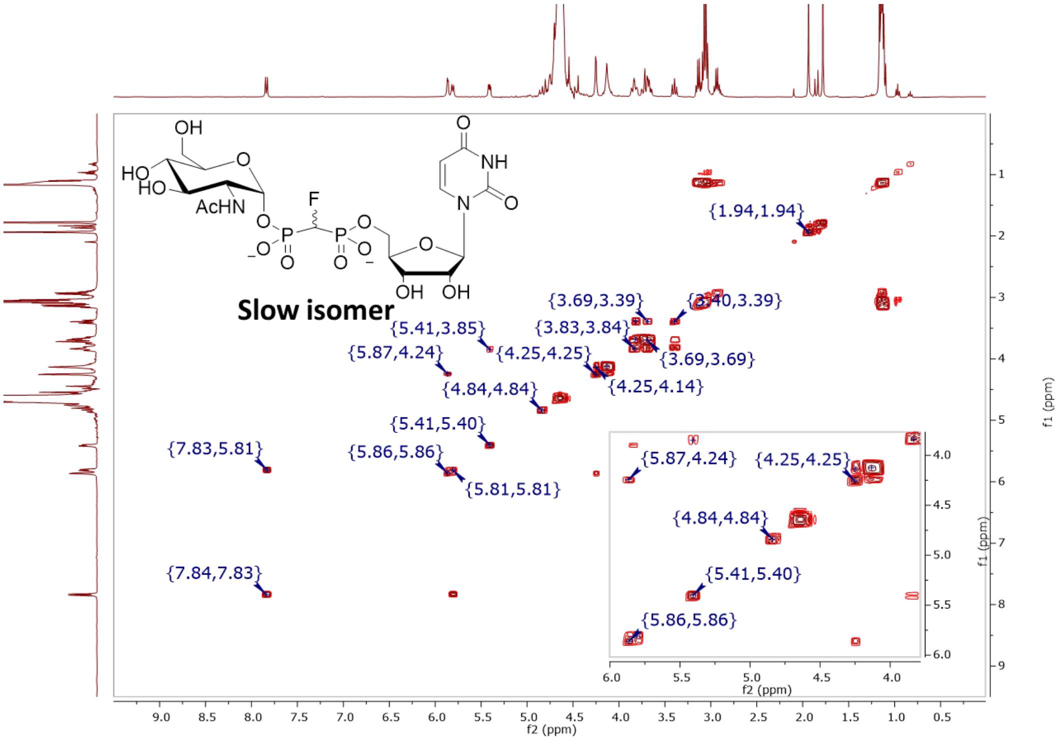

**Figure S165.** COSY (400 MHz, $D_2O$) of **8-α2** (*R*)-CHF isomer

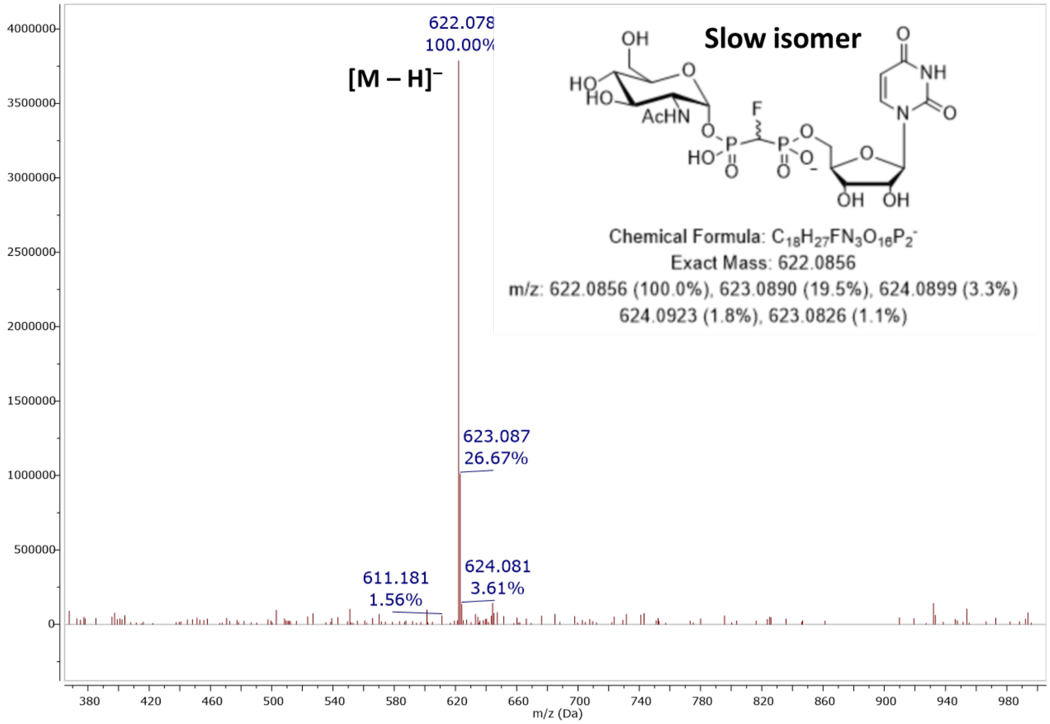

**Figure S166.** MS (ESI): [M – H]⁻ of **8-α2** (*R*)-CHF isomer

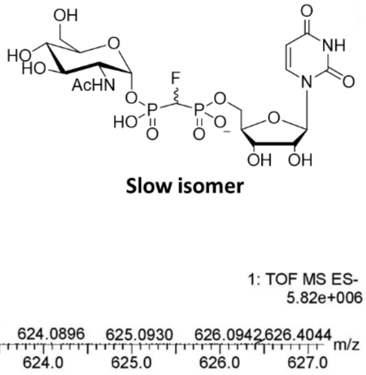

**Elemental Composition Report**

**Single Mass Analysis**
Tolerance = 5.0 PPM / DBE: min = -1.5, max = 50.0
Element prediction: Off
Number of isotope peaks used for i-FIT = 3

Monoisotopic Mass, Even Electron Ions
234 formula(e) evaluated with 2 results within limits (up to 50 closest results for each mass)
Elements Used:
C: 0-60 H: 0-100 N: 2-4 O: 15-17 F: 0-1 P: 1-3
Pouya Haratipour, USCMCK14
Synapt_26239 25 (0.519) Cm (25:27-6:8)

1: TOF MS ES-
5.82e+006

| Mass | Calc. Mass | mDa | PPM | DBE | i-FIT | Norm | Conf(%) | Formula |
|---|---|---|---|---|---|---|---|---|
| Minimum: | | | | -1.5 | | | | |
| Maximum: | | 5.0 | 5.0 | 50.0 | | | | |
| 622.0850 | 622.0851 | -0.1 | -0.2 | 7.5 | 696.2 | 0.015 | 98.47 | C18 H27 N3 O16 F P2 |
| | 622.0839 | 1.1 | 1.8 | 11.5 | 700.4 | 4.183 | 1.53 | C21 H26 N3 O15 P2 |

**Figure S167.** Elemental composition report from HRMS (ESI-TOF) [M – H]⁻ of **8-α2** (*R*)-CHF isomer

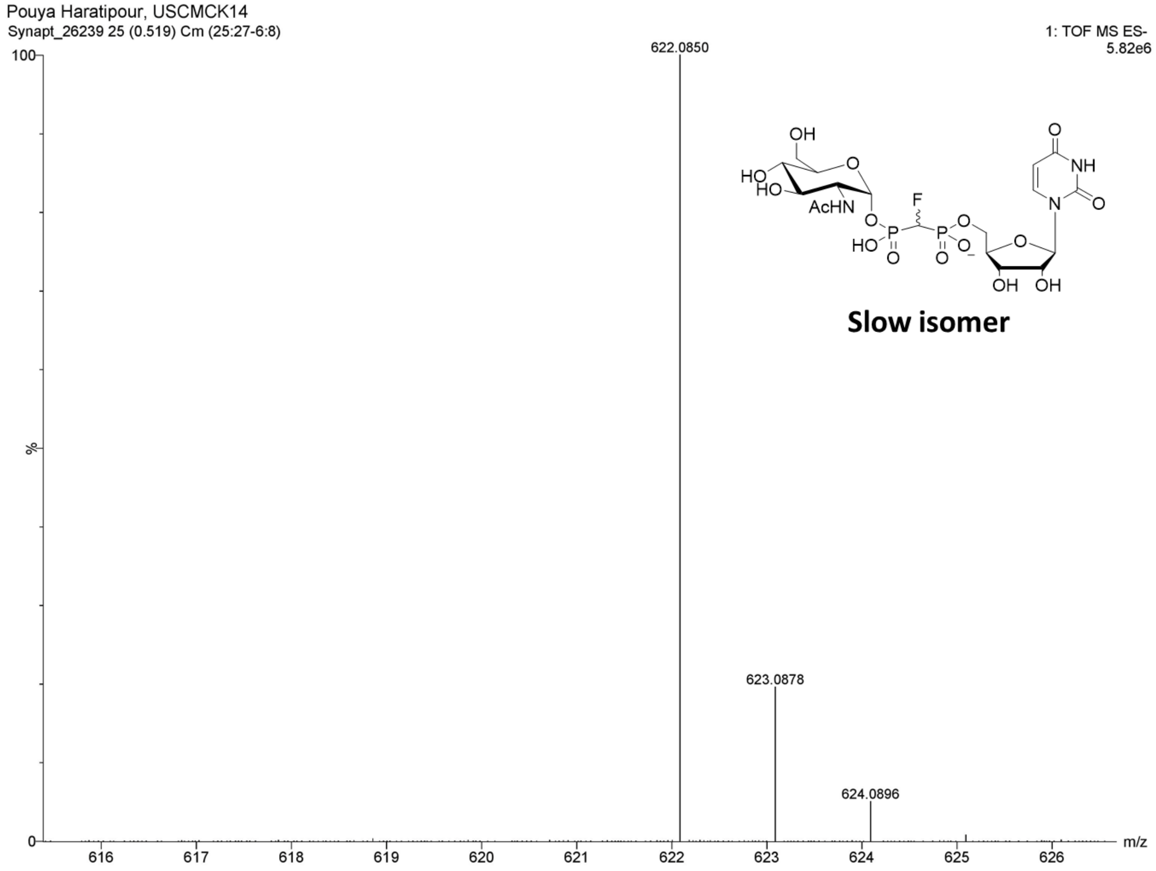

**Figure S168.** HRMS (ESI-TOF): [M – H]$^-$ of **8-α2** (*R*)-CHF isomer

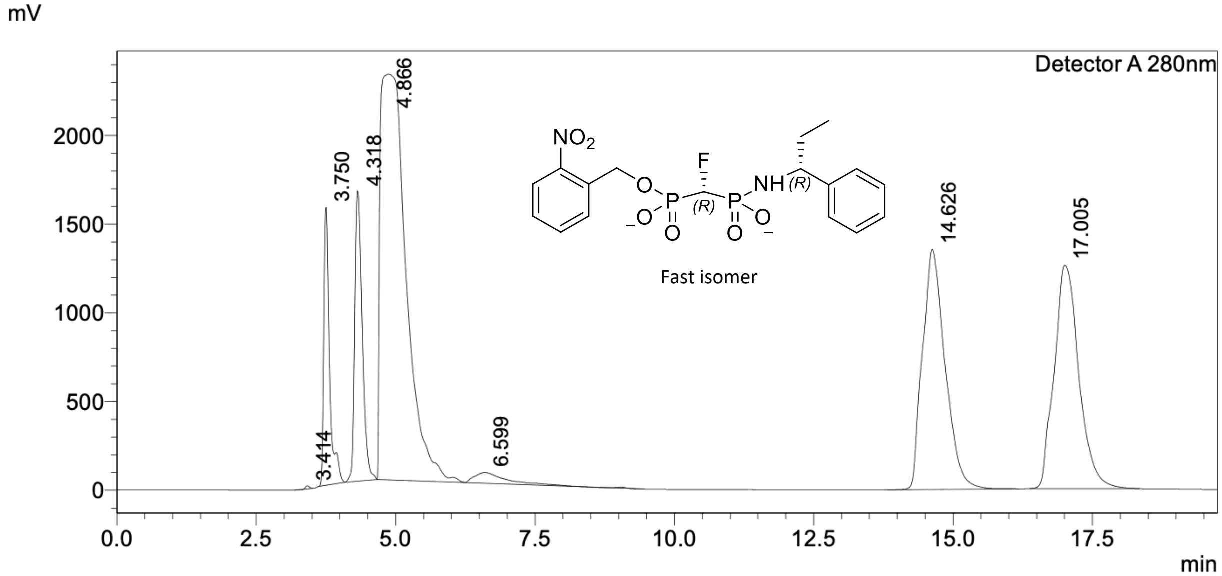

**Figure S169.** Semi-Preparative RP-HPLC separation of **26** (*R*)-CHF) and (*S*)-CHF (1:1)

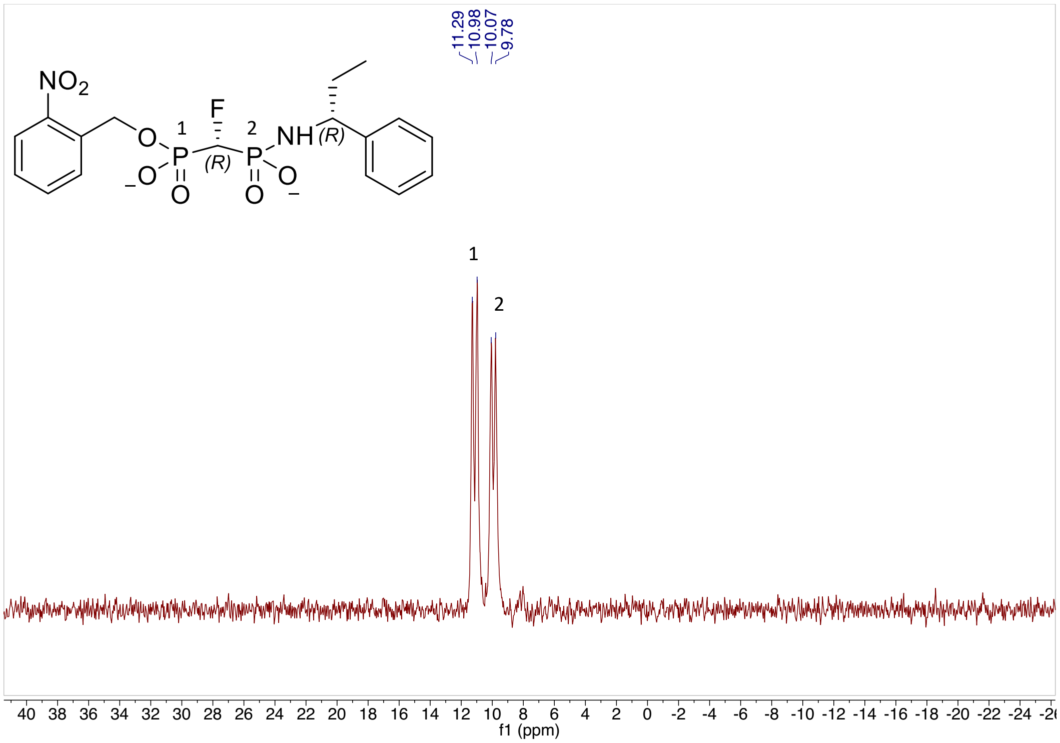

**Figure S170.** $^{1}$H NMR (500MHz, $CD_3OD$) of (*R*)-CHF isomer **26**

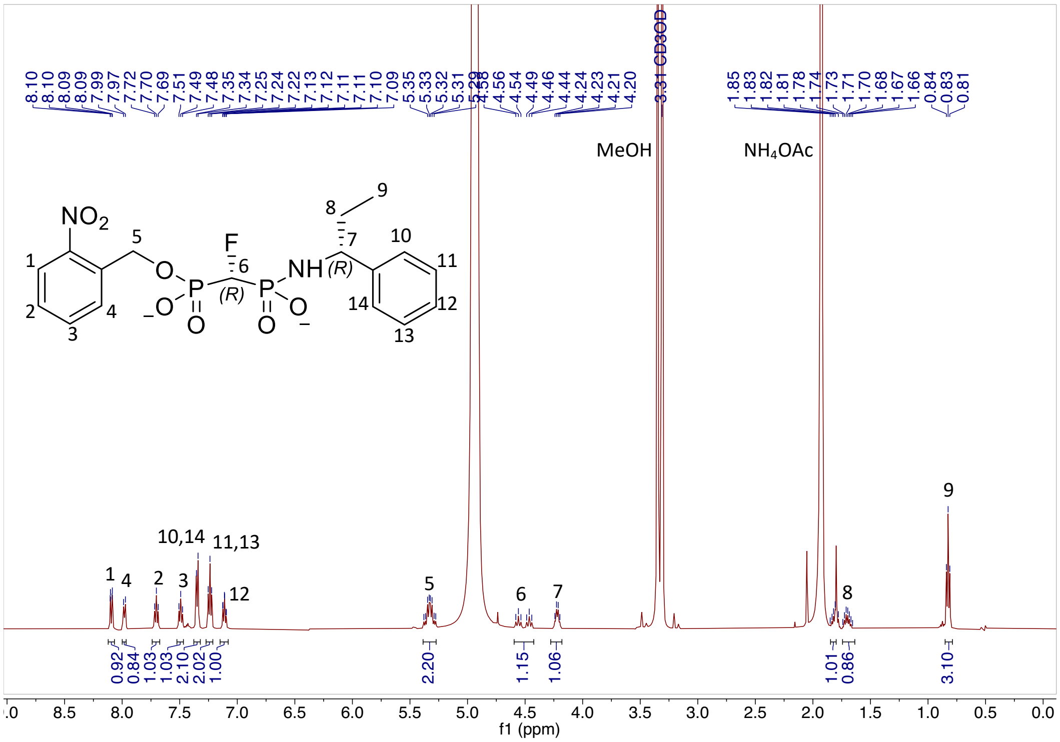

**Figure S171.** $^{31}$P NMR (202MHz, $CD_3OD$) of (*R*)-CHF isomer **26**

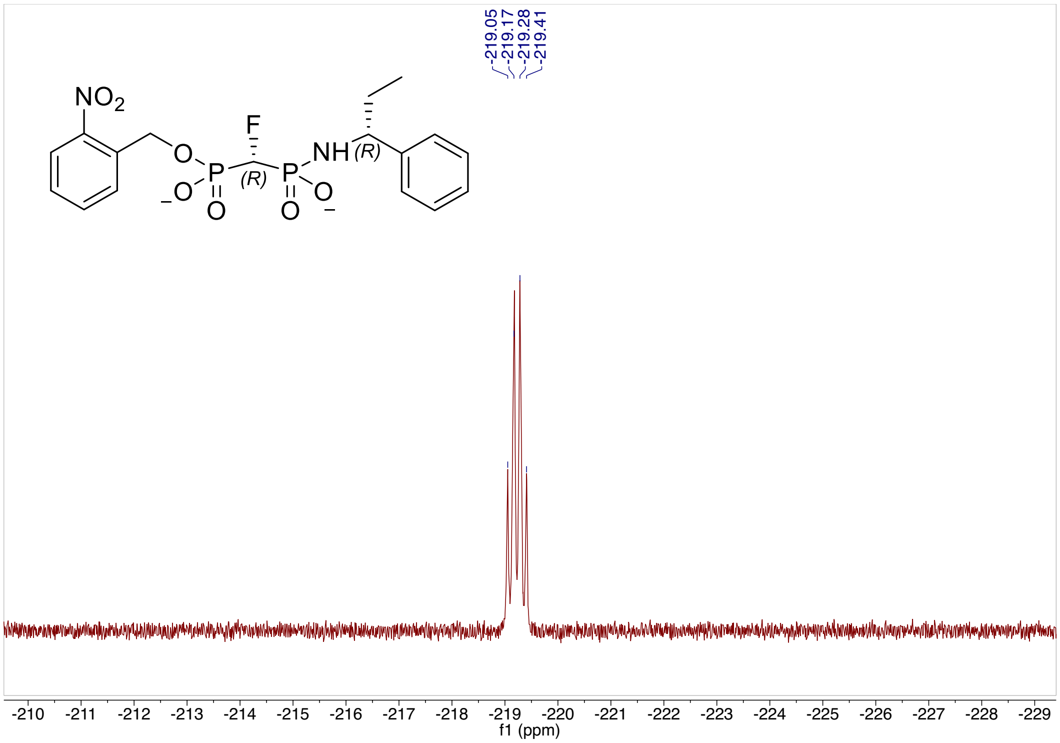

**Figure S172.** $^{19}F$ NMR (470MHz, $CD_3OD$) of (*R*)-CHF isomer **26**

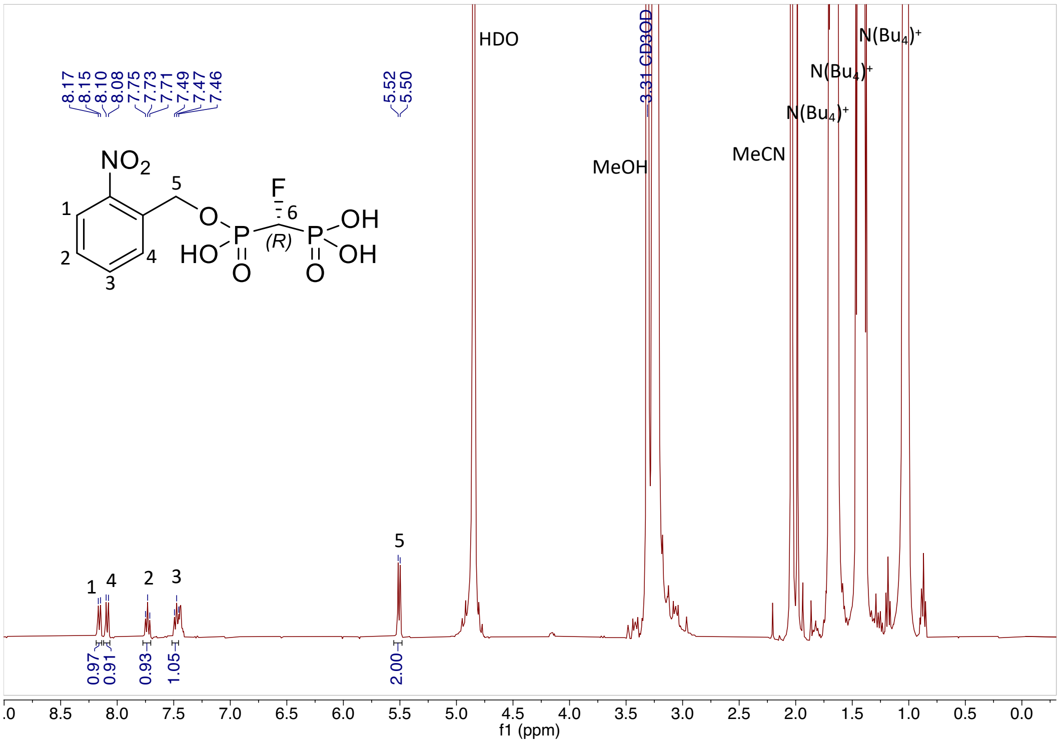

**Figure S173.** $^{1}H$ NMR (500MHz, $CD_3OD$) of (*R*)-CHF isomer **27**

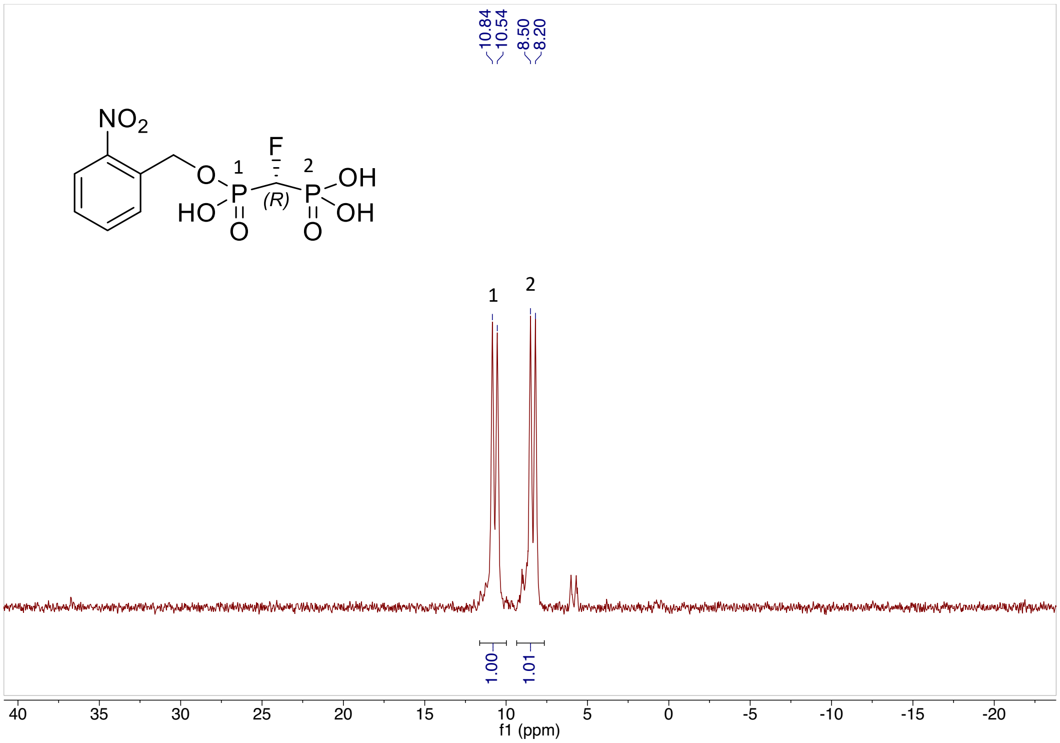

**Figure S174.** $^{31}P$ NMR (202MHz, $D_2O$) of (*R*)-CHF isomer **27**

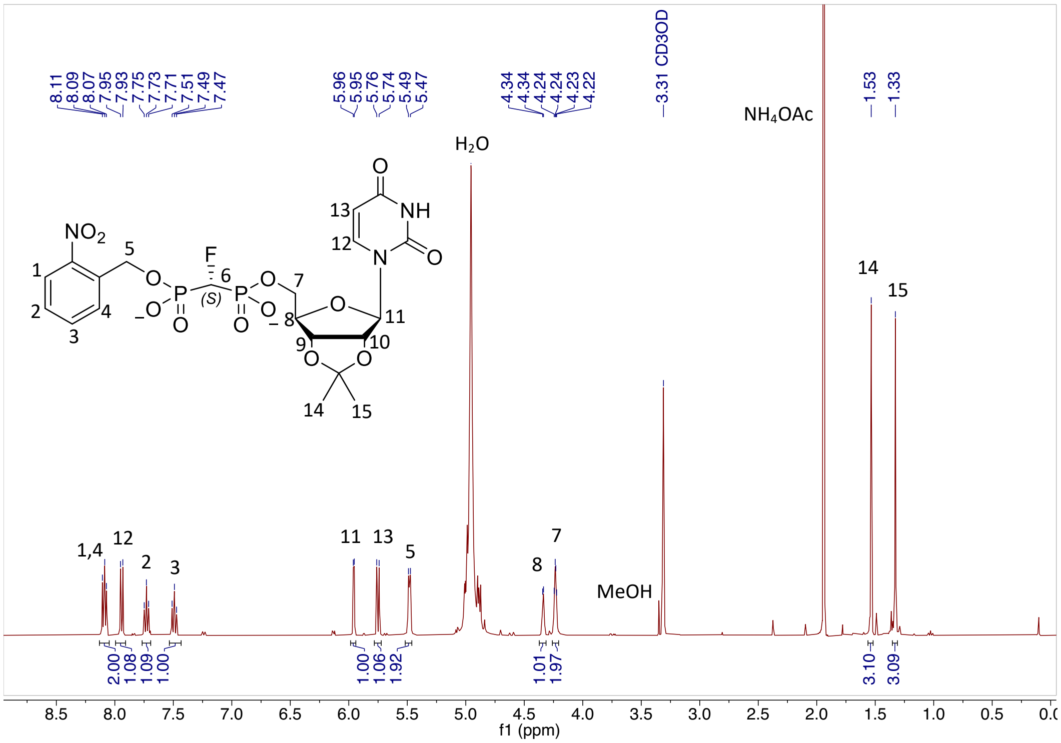

**Figure S175.** $^{1}H$ NMR (400MHz, $CD_3OD$) of (*R*)-CHF isomer **28**

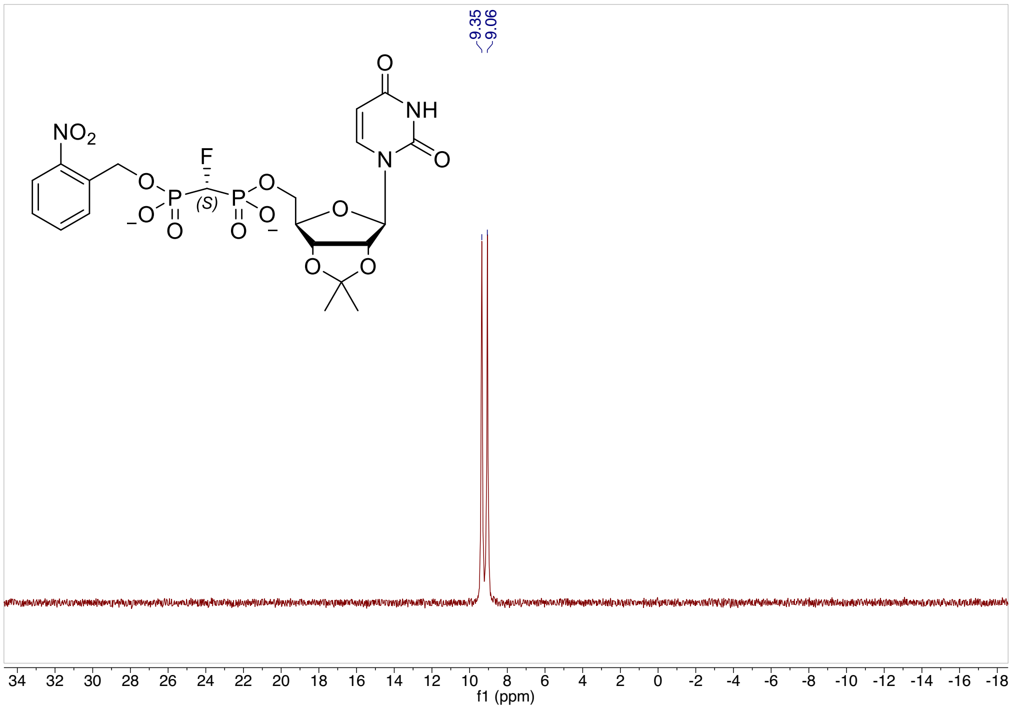

**Figure S176.** $^{31}$P NMR (202MHz, $CD_3OD$) of (*R*)-CHF isomer **28**

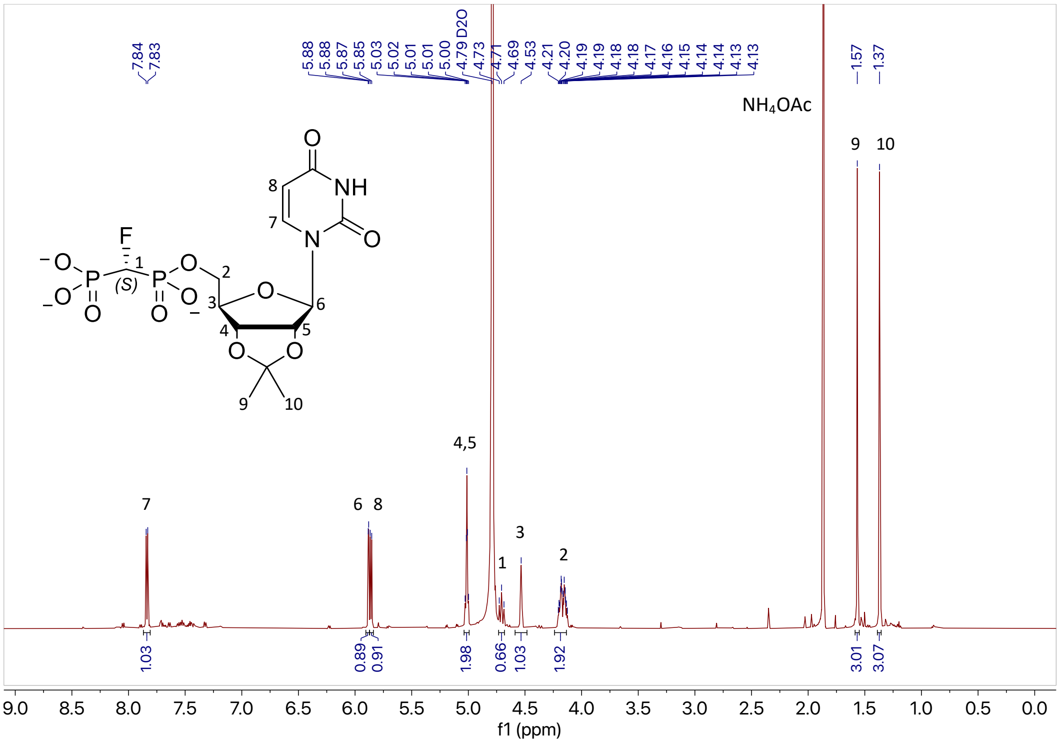

**Figure S177.** $^{1}$H NMR (600MHz, $D_2O$) of (*R*)-CHF isomer **29**

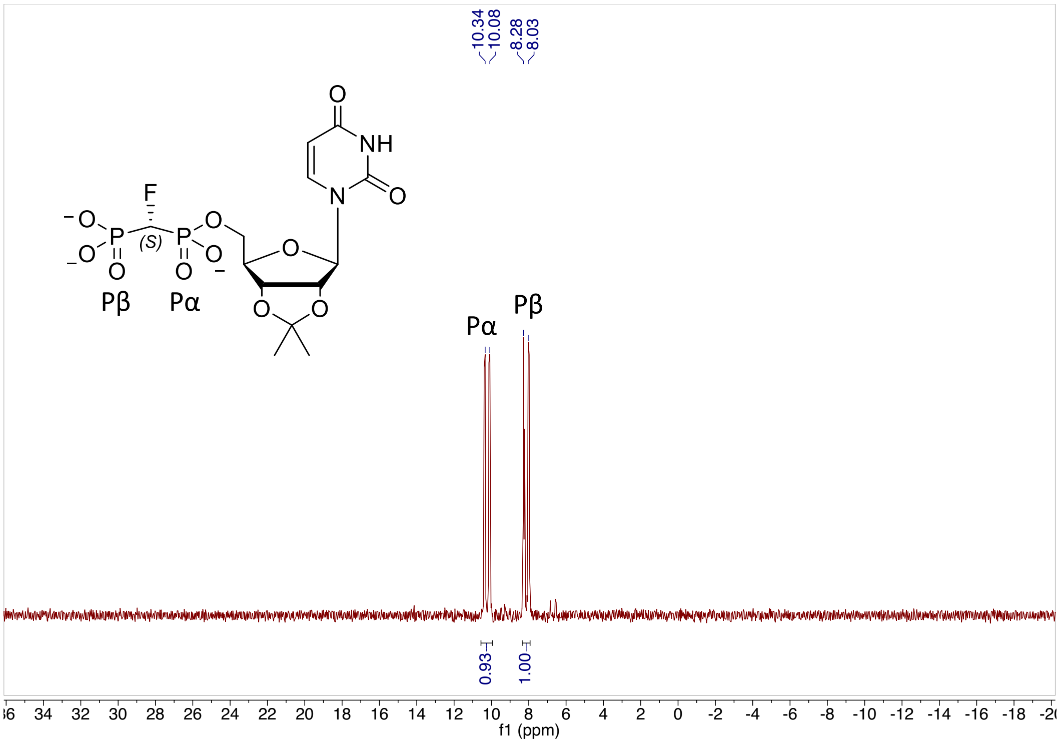

**Figure S178.** $^{31}P$ NMR (243MHz, $D_2O$) of (*R*)-CHF isomer **29**

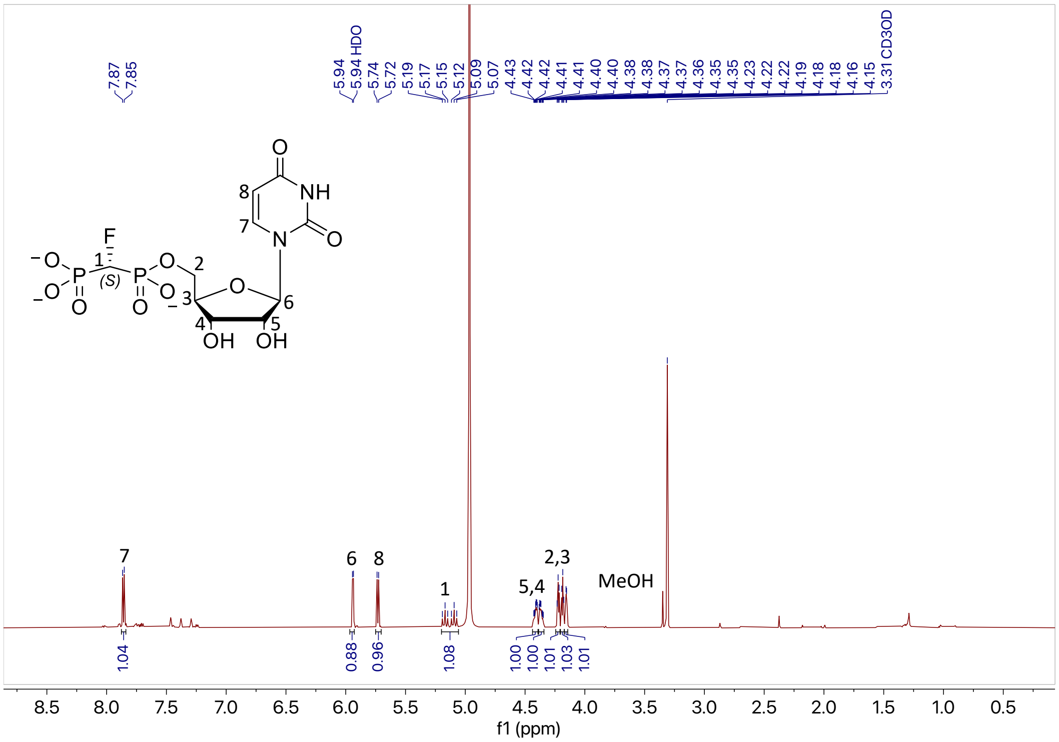

**Figure S179.** $^{1}H$ NMR (600MHz, $CD_3OD$) of (*R*)-CHF isomer **30**

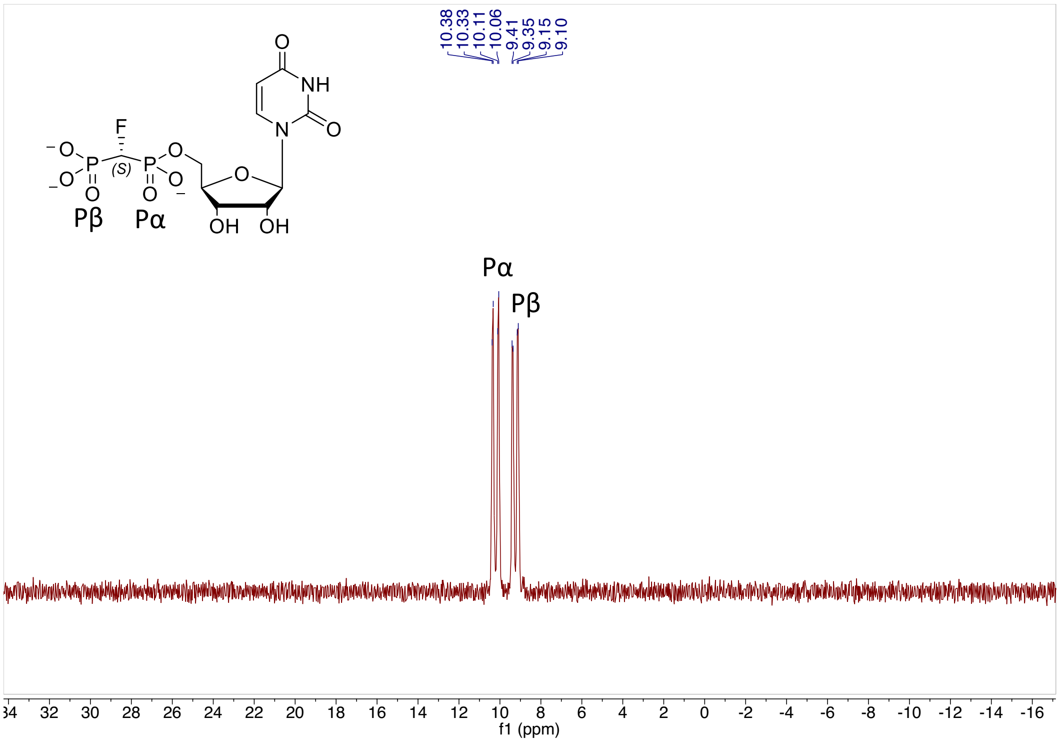

**Figure S180.** $^{31}$P NMR (243MHz, $D_2O$) of (*R*)-CHF isomer **30**

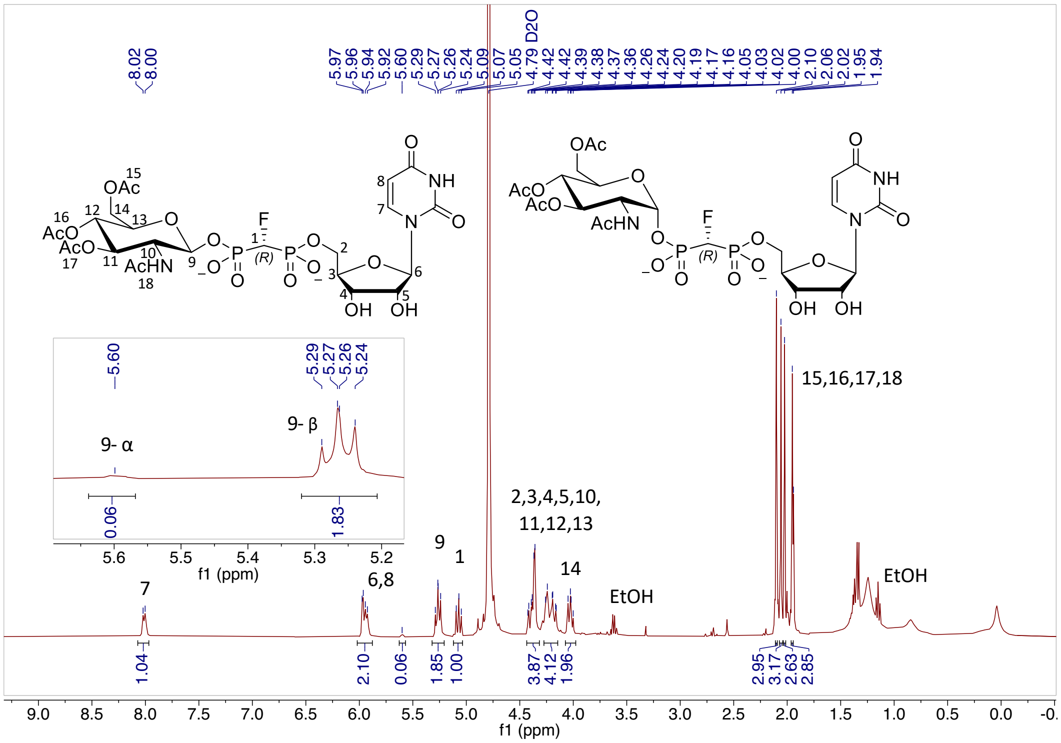

**Figure S181.** $^{1}$H NMR (400MHz, $D_2O$) of (*R*)-CHF anomers **31α** and **31β**

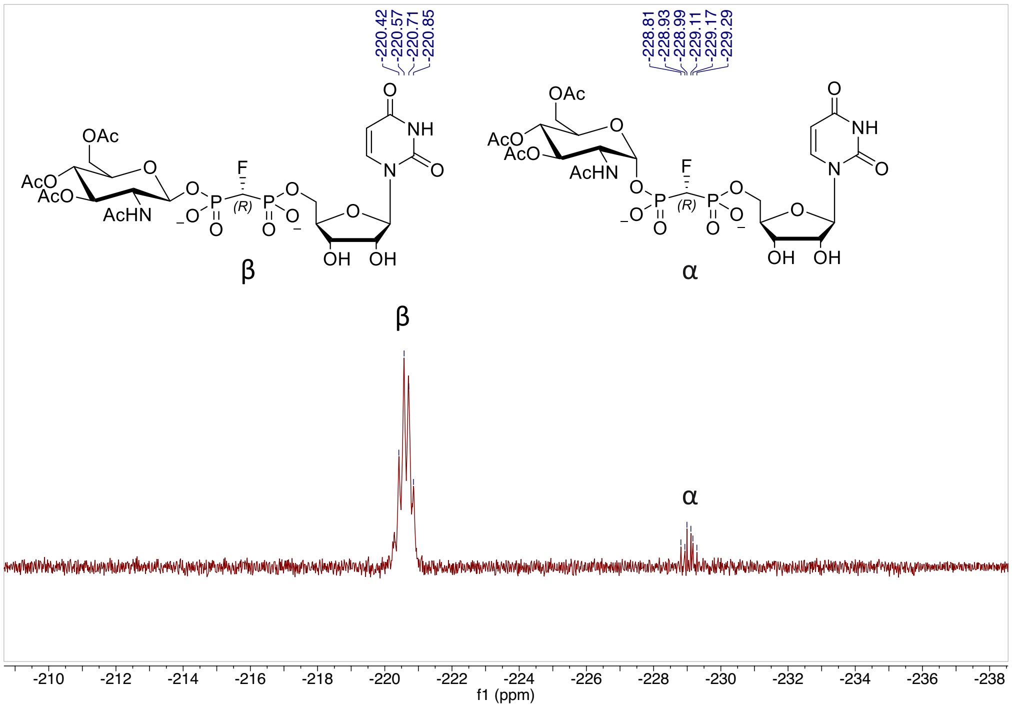

**Figure S182.** $^{19}F$ NMR (376MHz, $D_2O$) of (*R*)-CHF anomers **31α** and **31β**

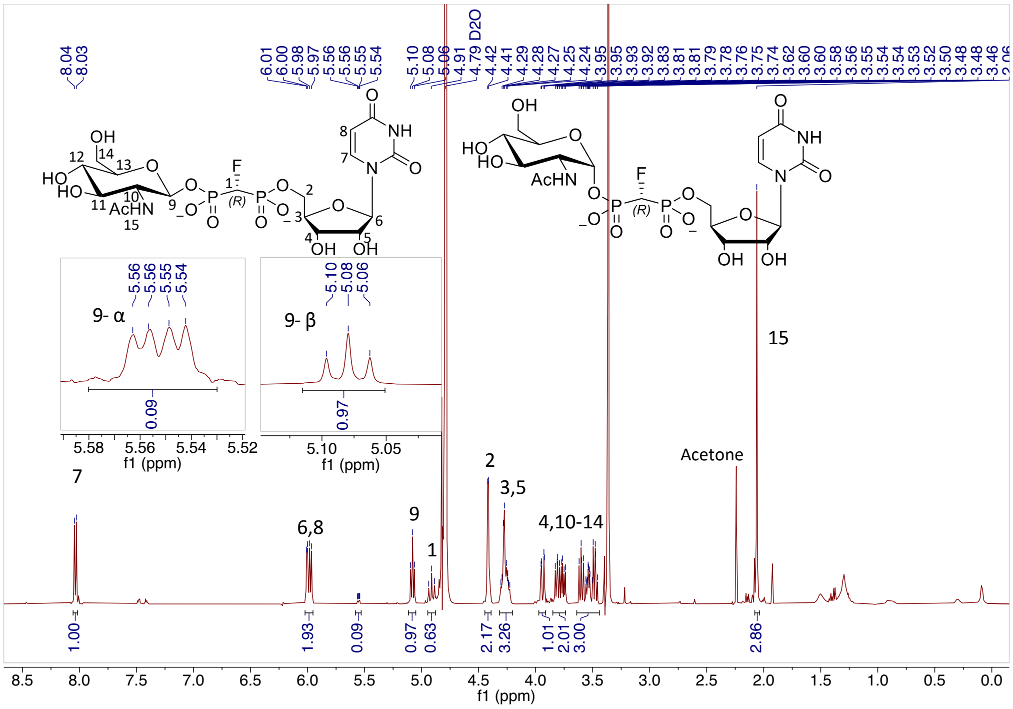

**Figure S183.** $^{1}H$ NMR (500MHz, $D_2O$) of (*R*)-CHF anomers **8α** (R) and **8β** (R)

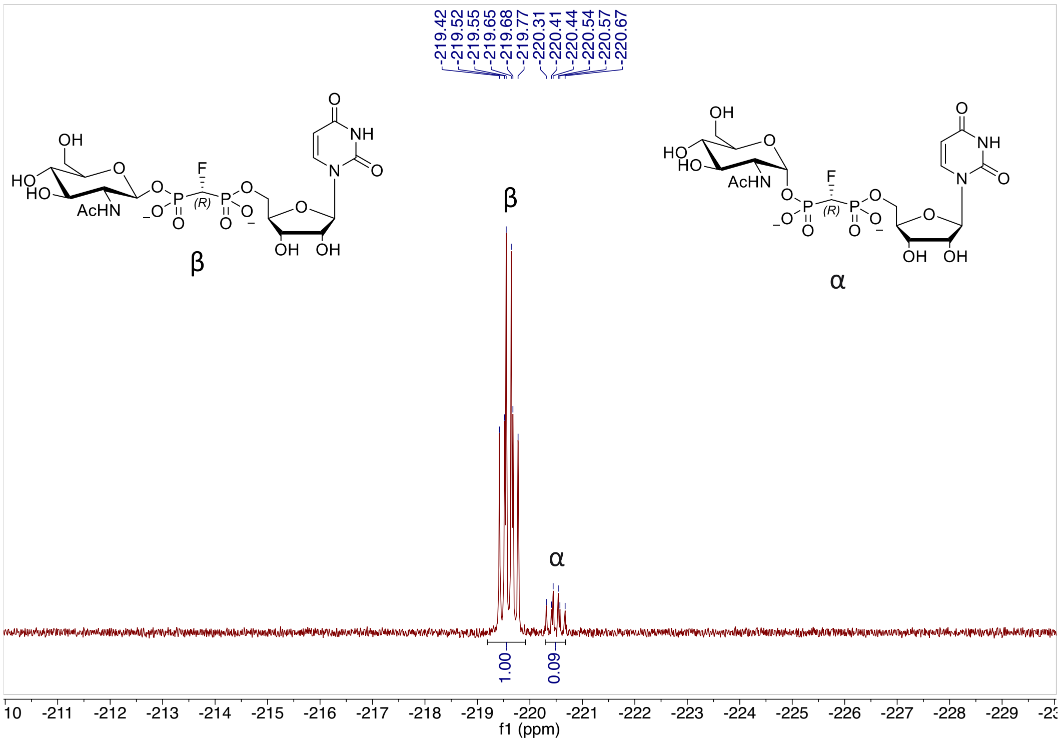

**Figure S184.** $^{19}F$ NMR (470MHz, $D_2O$) of (*R*)-CHF anomers **8α** (R) and **8β** (R)

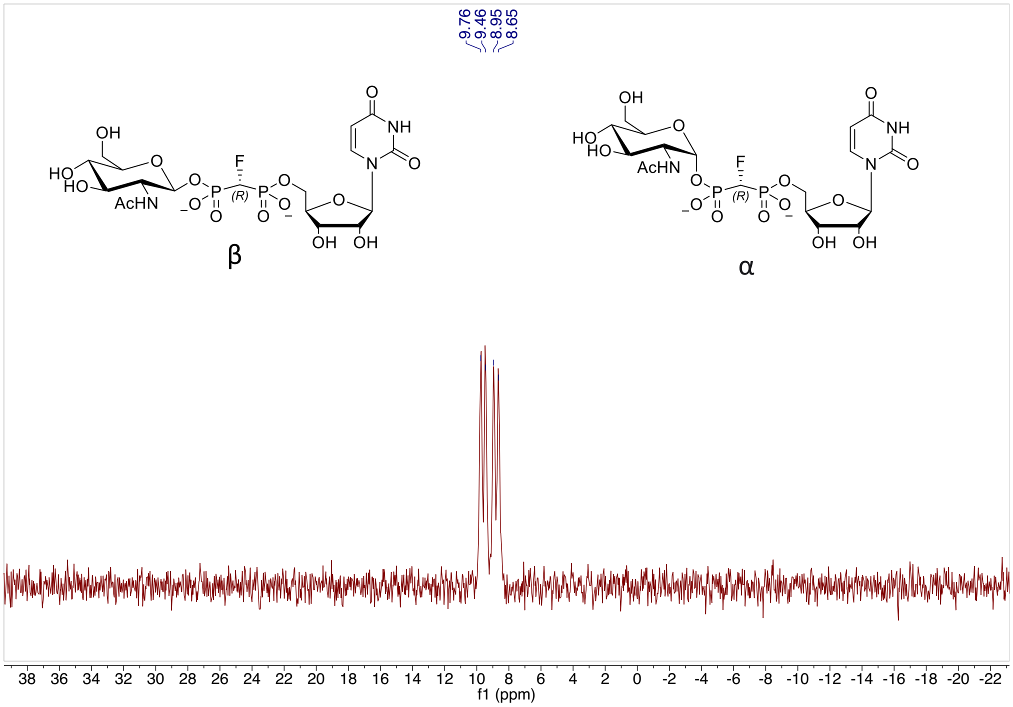

**Figure S185.** $^{31}P$ NMR (202MHz, $D_2O$) of (*R*)-CHF anomers **8α** (R) and **8β** (R)

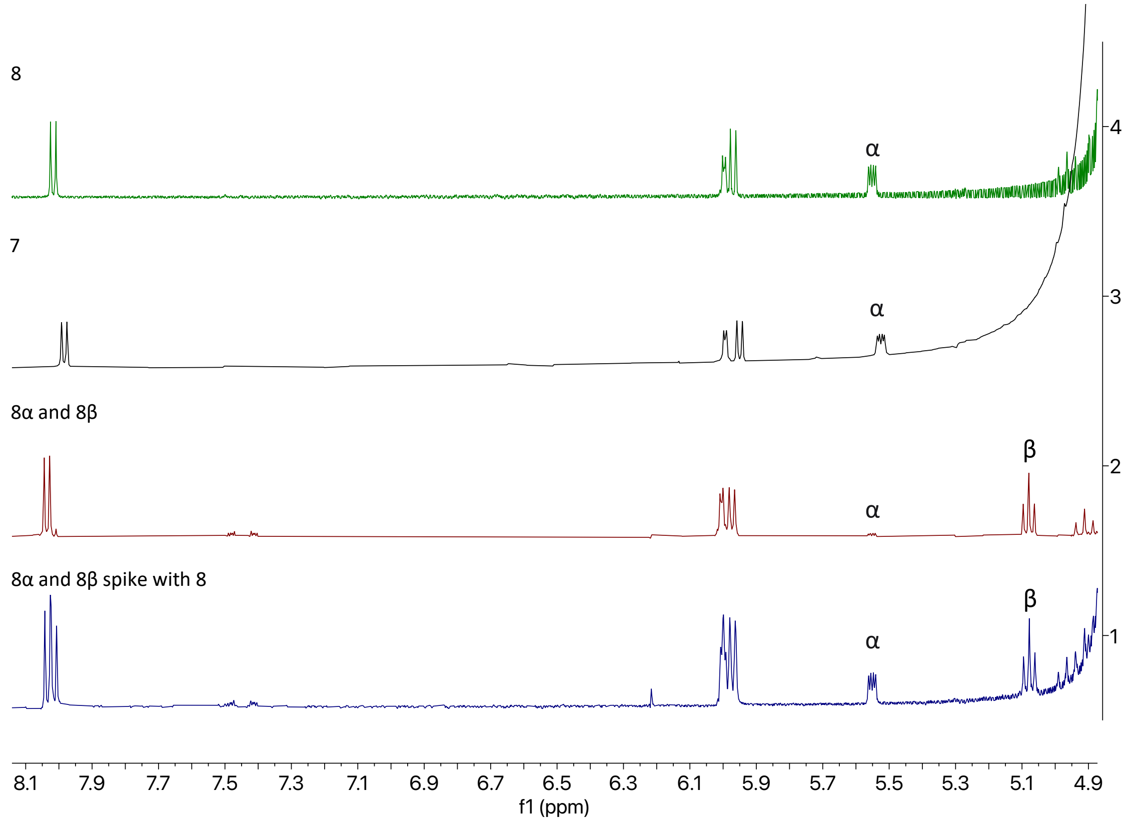

**Figure S186.** [1]H NMR of (*R*)-CHF anomers **8α** (*R*) and **8β** (*R*) stacked with (*S*)-CHF isomer **7** and (*R*)-CHF isomer **8**.

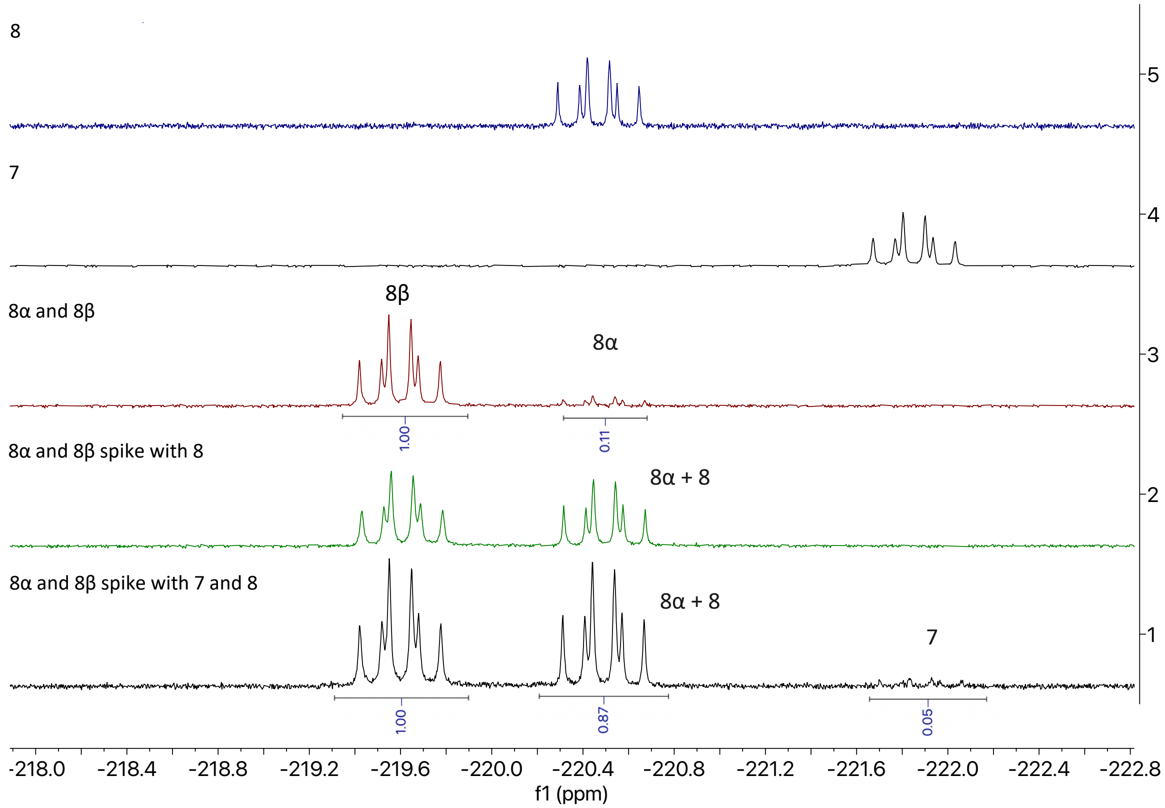

**Figure S187.** $^{19}$F NMR of (*R*)-CHF anomers **8α** and **8β** stacked with (*S*)-CHF isomer **7** and (*R*)-CHF isomer **8**.

#### Summary of nuclear Overhauser Effect (NOE) Studies

We investigated the solution-state conformations of UDP-GlcNAc, **(*S*)-7,** and **(*R*)-8** in the presence of $MgCl_2$. In all three compounds, the uracil protons are in NOE range to many of the GlcNAc protons, indicating a collapsed structure. The NOEs are similar amongst all three compounds with the exception of the *N*-acetyl group protons on the GlcNAc. UDP-GlcNAc and **(*S*)-7** show the acetyl group within detectable range of NOEs. The **(*R*)-8** isomer also has the *N*-acetyl group within range for an NOE effect, albeit with a much weaker signal. This difference is significant enough to conclude that **(*S*)-7** and **(*R*)-8** have different favored conformations in solution, and that **(*S*)-7** behaves similarly to UDP-GlcNAc in solution with this data.

Overall, the signals are too weak to calculate meaningful distances from these NOEs, but we can make assumptions on the conformation of the molecule based on the relative sizes of the NOE peaks. NOTE: Additional large peaks are HEPES buffer.

**NMR summary**

$^1H$ -$^1H$ NOESY spectra were recorded on a Bruker 401 MHz and 600 MHz NMR spectrometers. Each sample was dissolved with $D_2O$ with 0.95 equiv of $Mg^{2+}$. Overall, the enhancement of the uracil H5 resonance (5.85 ppm) is somewhat larger in the **(*S*)-7** isomer than in the **(*R*)-8** isomer but the most notable difference is in the enhancement of the *N*-acetyl methyl signals. Confirmation of this comes from $^1H$-$^1H$ NOE experiments, inverting the (major) methyl resonance (1.985 ppm) enhances the resonances of both uracil protons of the UDP-GlcNAc and the **(*S*)-7**, but does not show the same enhancement for the **(*R*)-8**; inverting the uracil H6 resonance (7.856 pm) likewise enhances the methyl resonance only for UDP-GlcNAc and **(*S*)-7** but not for **(*R*)-8.**

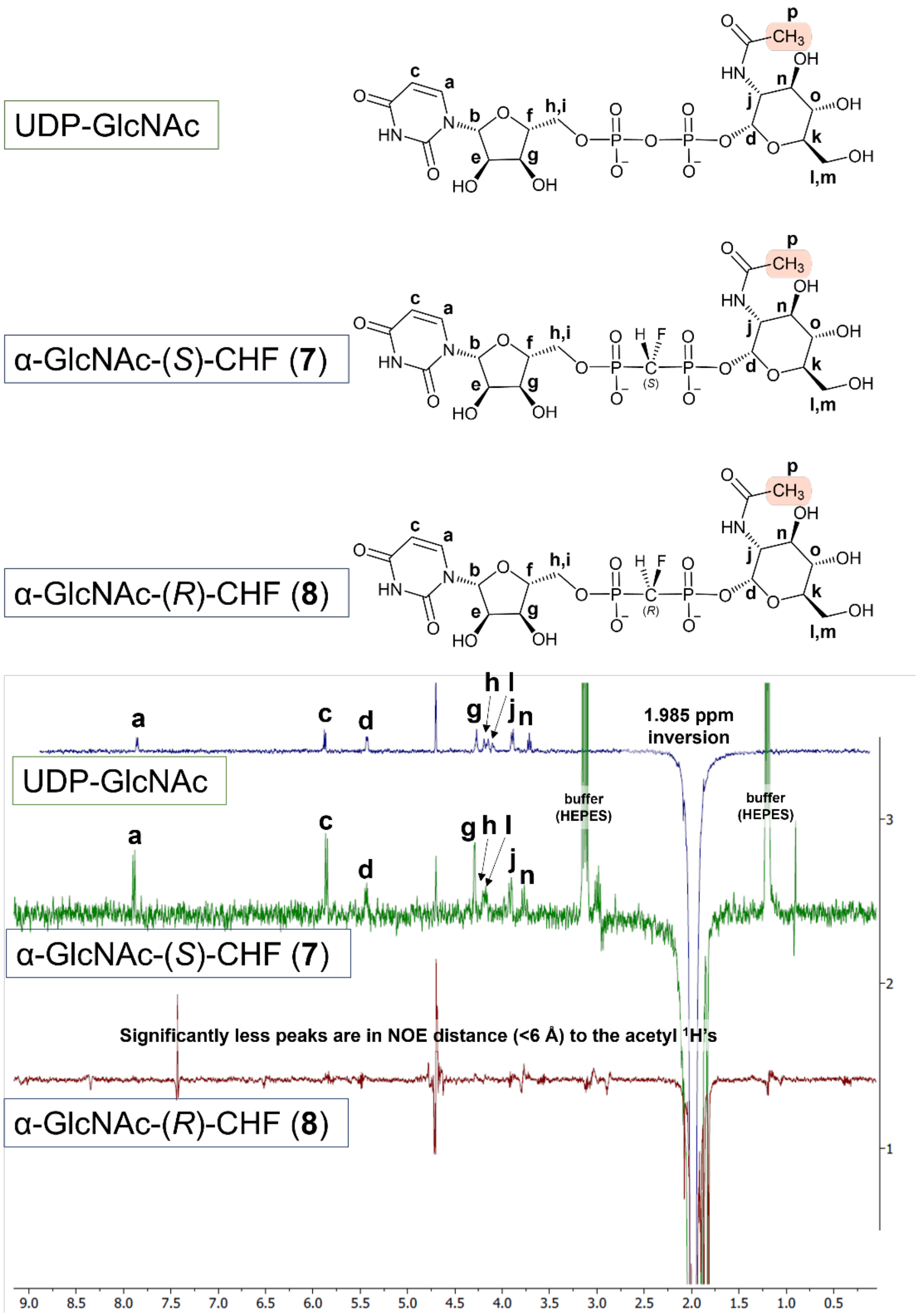

**Figure S188.** Top UDP GlcNAc, middle **(*S*)-7**, bottom is **(*R*)-8**

A 1-D selective NOE experiment was carried out, the *N*-acetyl protons (**p**) were selectively irradiated (1.985 ppm) and the spectra inverted. The positively phased peaks (**a**, **c**, **d**, **g**, **h**, **l**, **j** and **n**) are correlated within NOE distance constraints to the *N*-acetyl-protons.

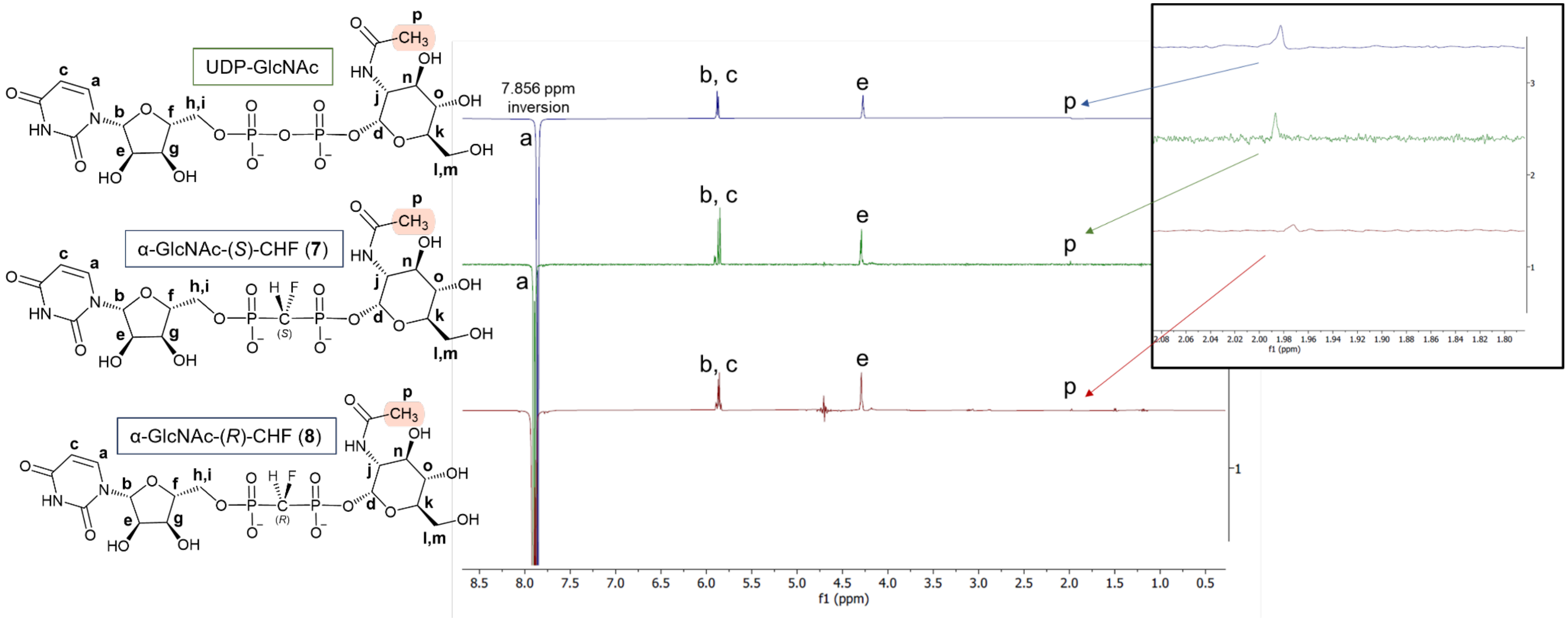

**Figure S189.** Top UDP-GlcNAc, middle **(*S*)-7**, bottom **(*R*)-8**

A 1-D selective NOE experiment was carried out, a select uracil proton (**a**) was selectively irradiated (7.856 ppm) and the spectra inverted. The positively phased peaks (**b**, **c**, **e** for all three compounds. The N-acetyl protons (**p**) only correlates strongly with UDP-GlcNAc and **(S)-7**) are correlated within NOE distance constraints to the *N*-acetyl-protons.

**Table S1.** Exact chemical shifts have been recorded for (*S*) and (*R*) and changes in shifts (δ) in ppm are recorded. This is meant to show how similar each compound is via $^{1}H$ and $^{13}C$ chemical shifts. (r = ribose; u = uracil, g = GlcNAc)

| | **(*S*) 7** | **$^{1}H$** (ppm) | **$^{13}C$** (ppm) | | **(*R*) 8** | **$^{1}H$** (ppm) | **$^{13}C$** (ppm) | **δH** (ppm) | **δC** (ppm) |
|---|---|---|---|---|---|---|---|---|---|
| u6 | **a** | 7.89, d | 141.7 | u6 | **a** | 7.92, d | 141.7 | -0.03 | 0 |
| r1 | **b** | 5.90, d | 88.4 | r1 | **b** | 5.90, d | 88.3 | 0 | 0.1 |
| u5 | **c** | 5.86, d | 102.5 | u5 | **c** | 5.88, d | 102.4 | -0.02 | 0.1 |
| g1 | **d** | 5.44, dd | 94.2 | g1 | **d** | 5.46, dd | 93.5 | -0.02 | 0.7 |
| r2 | **e** | 4.30, m | 73.9 | r2 | **e** | 4.30, m | 73.7 | 0 | 0.2 |
| r4 | **f** | 4.20, br | 83.5 | r4 | **f** | 4.19, m | 83.4 | 0.01 | 0.1 |
| r3 | **g** | 4.28,m | 69.8 | r3 | **g** | 4.30, m | 69.7 | -0.02 | 0.1 |
| r5 | **h** | 4.17, 2H, m | 64.4 | r5 | **h** | 4.22, m | 64.3 | -0.05 | 0.1 |
| r5' | **i** | 4.17, 2H, m | 64.4 | r5' | **i** | 4.15, m | 64.3 | 0.02 | 0.1 |
| g2 | **j** | 3.92, m | 53.8 | g2 | **j** | 3.90, dm | 53.6 | -0.02 | 0.2 |
| g5 | **k** | 3.88, m | 72.9 | g5 | **k** | 3.87, m | 72.9 | 0.01 | 0 |
| g6' | **l** | 3.72, dd | 60.5 | g6' | **l** | 3.71, dd | 60.4 | 0.01 | 0.1 |
| g6 | **m** | 3.80, dd | 60.5 | g6 | **m** | 3.79, dd | 60.4 | 0.01 | 0.1 |
| g3 | **n** | 3.77, t | 70.9 | g3 | **n** | 3.74, m | 70.9 | 0.03 | 0 |
| g4 | **o** | 3.44, t | 69.6 | g4 | **o** | 3.45, t | 69.6 | -0.01 | 0 |
| g7 | **p** | 1.98, s | 23.6 | g7 | **p** | 1.98, s | 23.6 | 0 | 0.1 |
| CHF | **q** | 4.85, dt | 86.8 | CHF | **q** | 4.83, dt | 87.4 | 0.02 | -0.6 |

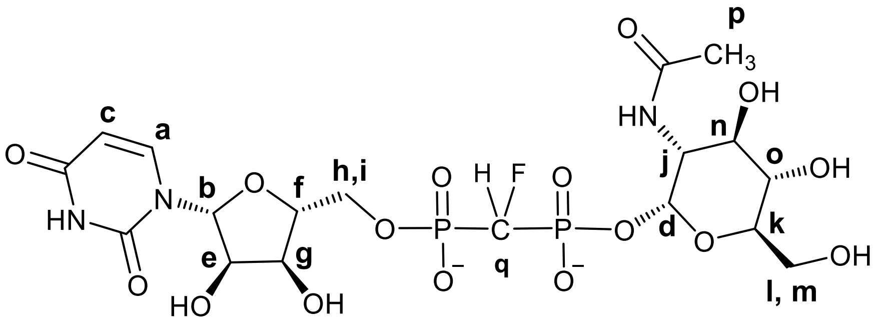

#### Sequence alignments of polyPGTs

**Figure S190**. Sequence alignment showing DD motif that is proposed to be associated with catalysis.

```
CLUSTAL O(1.2.4) multiple sequence alignment

sp|Q9X1N5|WECA_THEMA      --------------------------MWEAIISFFLTSVLS------------VFAKKTE  22
sp|Q9H3H5|GPT_HUMAN       ------------MWAFSELPMPLLINLIVSLLGFVATVTLIPAFRGHFIA-ARLCGQDLN  47
sp|Q03521|MRAY_BACSU      --------------------MLEQVILFTILMGFLISVLLSPILIPFLRRL--KFGQ---  35
sp|O66465|MRAY_AQUAE      MLYQLALLLKDYWFAFNVLKYITFRSFTAVLIAFFLTLVLSPSFINRLRKIQRLFGG---  57
                                                    :   ::.*. :  *               .

sp|Q9X1N5|WECA_THEMA      FLDRPDSRKSHG--RAVPPVGGVSIFLTLLIF----------ERDNPF-----------F  59
sp|Q9H3H5|GPT_HUMAN       KTSRQQIPESQGVISG-----AVFLIILFCFIPFPFLNCFVKEQCKAFPHHEFVALIGAL  102
sp|Q03521|MRAY_BACSU      -SIREEGPKSHQKKSGTPTMGGVMIILSIIVTTIVMTQKF----SEISPE-----MVLLL  85
sp|O66465|MRAY_AQUAE      -YVREYTPESHEVKKYTPTMGGIVILIVVTLSTLLLMRW-----DIKY-------TWVVL  104
                             *    :*:          .: ::: . .                            :

sp|Q9X1N5|WECA_THEMA      LFSIPLFLLGLLDDLFDLSYRIKLAVTALV----------------AVWFSTAVTI-E--  100
sp|Q9H3H5|GPT_HUMAN       LAICCMIFLGFADDVLNLRWRHKLLLPTAASL----------PLLMVYFTNFGNTTIVVP  152
sp|Q03521|MRAY_BACSU      FVTLGYGLLGFLDDYIKVVMKRNLGLTSKQKLIGQIIIAVVFYA-VYHYYNFAT---DIR  141
sp|O66465|MRAY_AQUAE      LSFLSFGTIGFWDDYVKLKNKKGISIKTK--FLLQVLSASLISVLIYYWADIDT---ILY  159
                          :       :*: ** ..:  :  : : :                    : .

sp|Q9X1N5|WECA_THEMA      --------VSIFGARIHPVFFVIWFVGMVNAFNVVDGLDGLLSGISLFSSLMIGER----  148
sp|Q9H3H5|GPT_HUMAN       KPFRPILGLHLDLGILYYVYMGLLAVFCTNAINILAGINGLEAGQSLVISASIIVFNLVE  212
sp|Q03521|MRAY_BACSU      IP---GTDLSFDLGWAYFILVLFMLVGGSNAVNLTDGLDGLLSGTAAIAFGAFAILAWN-  197
sp|O66465|MRAY_AQUAE      FPF--FKELYVDLGVLYLPFAVFVIVGSANAVNLTDGLDGLAIGPAMTTATALGVVAYAV  217
                                  : .  .  :     :  *   **.*:  *::**  * :      :

sp|Q9X1N5|WECA_THEMA      ---------------------------SLAFSIIGFLPWNLPDAKVFLGNSGSFLLGAYL  181
sp|Q9H3H5|GPT_HUMAN       LEGDCRDDHV----------FSLYFMIPFFFTTLGLLYHNWYPSRVFVGDTFCYFAGMTF  262
sp|Q03521|MRAY_BACSU      ----------------QSQYDVAIFSVAVVGAVLGFLVFNAHPAKVFMGDTGSLALGGAI  241
sp|O66465|MRAY_AQUAE      --GHSKIAQYLNIPYVPYAGELTVFCFALVGAGLGFLWFNSFPAQMFMGDVGSLSIGASL  275
                                                      .  : :*:*  *   :::*:*:  .   *  :

sp|Q9X1N5|WECA_THEMA      STASVVFFEGDLGYATLFLGFPFYEIVFSFV-------------R-----------RLVV  217
sp|Q9H3H5|GPT_HUMAN       AVVGILGHF---SKT---MLLFFMPQVFNFLYSLPQLLHIIPCPRHRIPRLNIKTGKLEM  316
sp|Q03521|MRAY_BACSU      VTIAILTKL---EIL---LV--IIGGVFVI-ETLSVILQVISFKT---------TGKRIF  283
sp|O66465|MRAY_AQUAE      ATVALLTKS---EFI---FA--VAAGVFVF-ETISVILQIIYFRW--------TGGKRLF  318
                           . .::            :   .   ** :                          :  .

sp|Q9X1N5|WECA_THEMA      KKNPFSPDE----------------KHTHHVFSRKIGKWKTLLILVSFSLMFNLLGLSQK  261
sp|Q9H3H5|GPT_HUMAN       SYSKFKTKSLSFLGTFILKVAESLQLVTVHQSETEDGEFTECNNMTLINLLLKVLGPIHE  376
sp|Q03521|MRAY_BACSU      KMSPLHHH---------------YEL-------VGWSEWRVVVTFWAAGLLLAVLGIYIE  321
sp|O66465|MRAY_AQUAE      KRAPFHHH---------------LEL-------NGLPEPKIVVRMWIISILLAIIAISML  356
                          .   :  .                             :      :   .::: ::.

sp|Q9X1N5|WECA_THEMA      FYFIFLYVVLC---CVLLFTYCVLQRGNGNLKL---- 291
sp|Q9H3H5|GPT_HUMAN       RNLTLLLLLLQILGSAITFSI-----RYQLVRLFYDV 408
sp|Q03521|MRAY_BACSU      VWL---------------------------------- 324
sp|O66465|MRAY_AQUAE      KLR---------------------------------- 359
```

**Sequences and information on published crystal structures with ligands**

GPT (*H. sapiens*) with UDP-GlcNAc and Mg(II) - PDB: 6FWZ

GPT (*H. sapiens*) with tunicamycin – PDB: 6BW5

MraY (*A. aeolicus*) with Mg(II) - PDB: 4J72

WecA (*B. subtilis*) structure ND

#### Structural analysis of polyPGTs

**Figure S191**. Series of PyMOL structures of overlays between **6FWZ [**GPT with UDP-GlcNAc and Mg(II)] **and 6BW5 [**GPT with tunicamycin]

**A**. Overlay between: **6FWZ** (GPT with UDP-GlcNAc and Mg(II) - GPT pale blue, Mg(II) green, UDP-GlcNAc: C-white/O-red/N-blue, P-orange. **6BW5** (GPT with tunicamycin) – GPT wheat, tunicamycin: C-yellow/O-red/N-blue, P-orange.

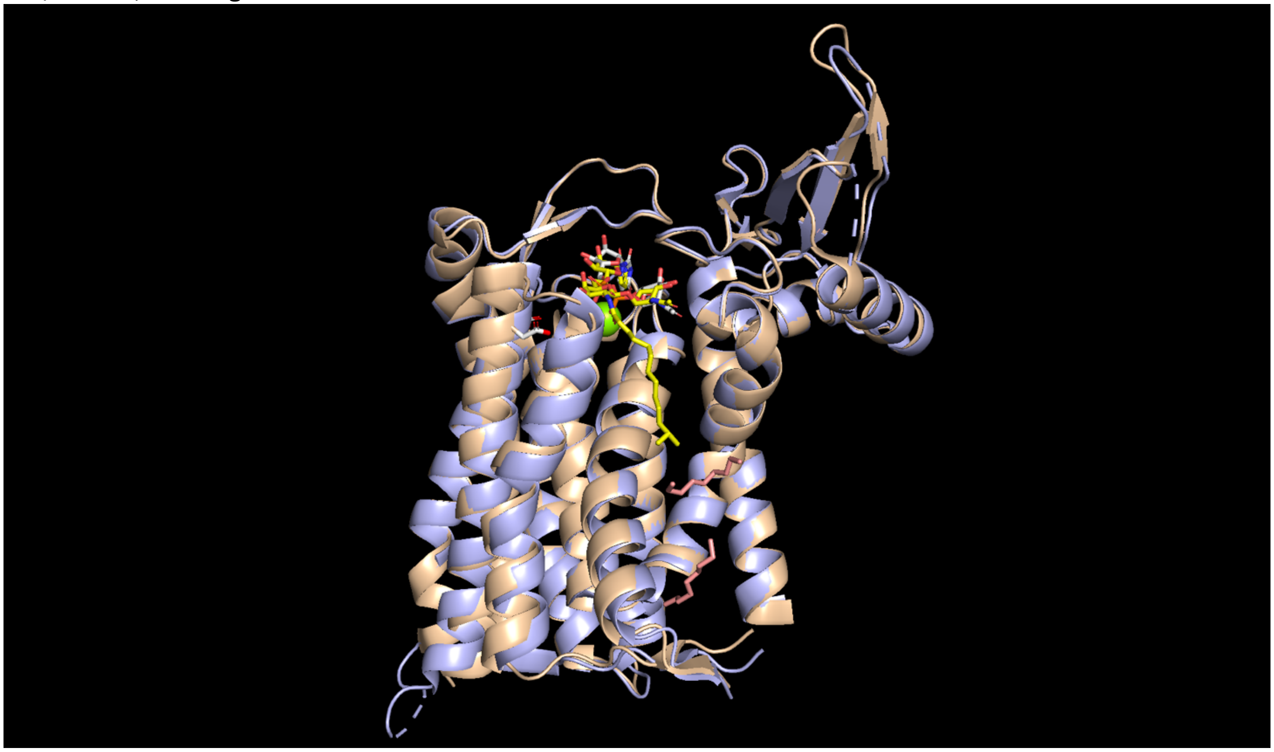

**B.** Close up of **Figure S191A** demonstrating the conserved DD motif that is proposed to be associated with catalysis, D115 and D116.

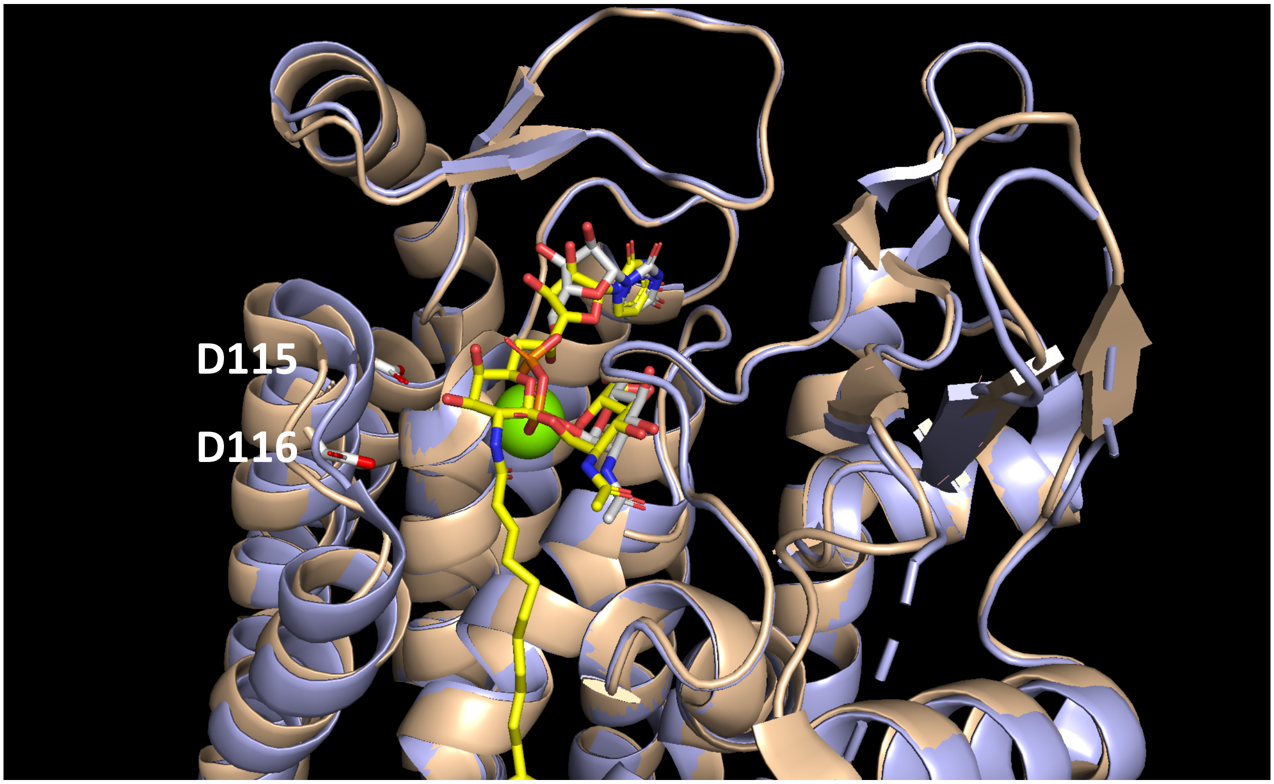

**C. 6FWZ**, GPT shown with UDP-GlcNAc and Mg(II).

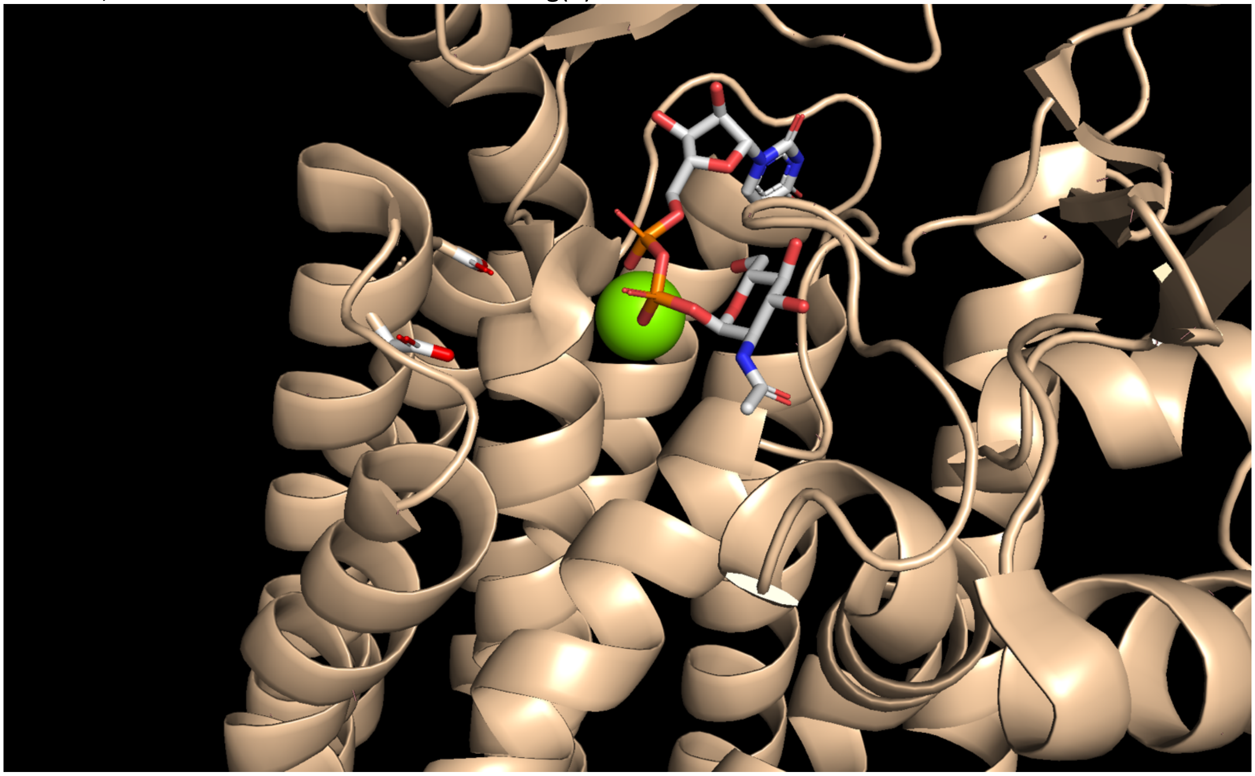

**D.** 6BW5, GPT shown with tunicamycin**.**

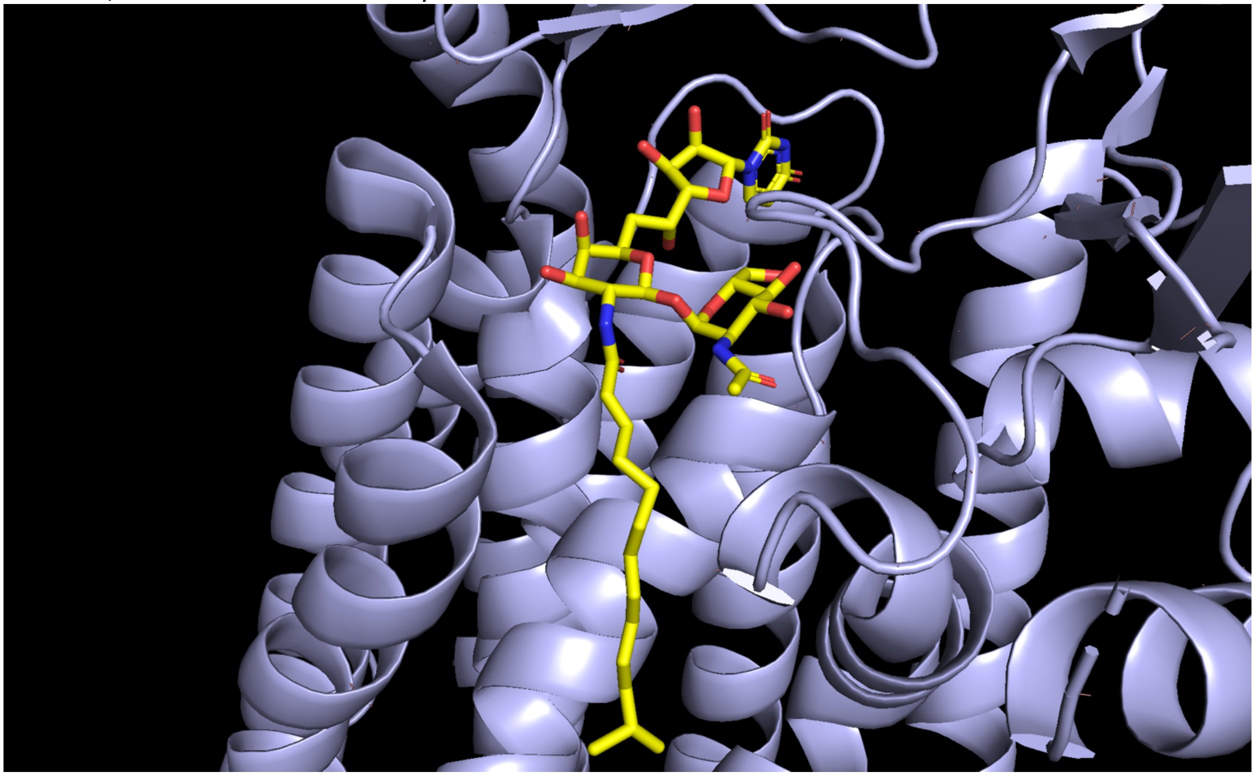
